# Analysis of the potential impact of genomic variants in SARS-CoV-2 genomes from India on molecular diagnostic assays

**DOI:** 10.1101/2020.08.05.238618

**Authors:** Abhinav Jain, Mercy Rophina, Saurabh Mahajan, Bhavya Balaji Krishnan, Manasa Sharma, Sreya Mandal, Teresa Fernandez, Sumayra Sultanji, Samatha Mathew, Sridhar Sivasubbu, Vinod Scaria

## Abstract

An isolated epidemic of Severe Acute Respiratory Syndrome Coronavirus 2 (SARS-CoV-2) causing Coronavirus Diseases (COVID-19) originating in Wuhan, China has now rapidly emerged into a global pandemic affecting millions of people worldwide. Molecular detection of SARS-CoV-2 using reverse transcription polymerase chain reaction (RT-PCR) forms the mainstay in screening, diagnosis and epidemiology of disease. The virus has been evolving through base substitutions. The recent availability of genomes of SARS-CoV-2 isolates from different countries including India motivated us to assess the presence and potential impact of variations in target sites for the oligonucleotide primers and probes used in molecular diagnosis. We catalogued a total of 132 primers or probes sequences from the literature and the public domain. Our analysis revealed a total of 125 unique genetic variants in 80 either primers or probes binding sites. A total of 13 unique variants had allele frequency of ≥ 1% in Indian SARS-CoV-2 genomes mapped to the primers or probes binding sites. A total of 15 primers or probes binding sites had cumulative variant frequency of ≥ 1% in the SARS-CoV-2 genomes. These included primers or probes sites which are widely used in India and across the world for molecular diagnosis as well as approved by national and international agencies. This highlights the need for sequencing genomes of emerging pathogens to make evidence based policies for development and approval of diagnostics. To the best of our knowledge, ours is the most comprehensive analysis of genomic variants in genomes of SARS-CoV-2 isolates from India and their potential impact on efficacy of molecular diagnostics.

---

Coronavirus Disease 2019 (COVID-19) has now rapidly emerged as a global pandemic. Reverse transcription Polymerase Chain Reaction (RT-PCR) based assays have been the mainstay for the diagnosis and screening of COVID-19 due the high sensitivity and specificity (Shen et al. 2020). These assays utilize oligonucleotide primers and probes specific to the viral nucleic acid. The SARS-CoV-2 has been continuously evolving and has an estimated substitution rate of 1.19 to 1.31 × 10^-3^ per site per year (Li et al. 2020). Recent reports that suggest genetic variation in viruses at the primers or probes binding site could decrease its sensitivity (Yang et al. 2014). Motivated by the availability of a large number of genomes of SARS-CoV-2 isolates from India, we attempted to understand the genomic variants and their potential impact on molecular assays.

We analysed genomic sequences of SARS-CoV-2 isolates from India in GISAID (Shu and McCauley 2017) as on 23rd of June 2020. sequences with <99% alignment and ≥1% gaps were not considered for analysis. The genomes were re-aligned to the reference SARS-CoV-2 genome Wuhan-Hu-1 (Wu et al. 2020) using EMBOSS needle (Rice et al. 2000) and parsed for variants using bespoke scripts. The primer/probe sequences were compiled using extensive literature searches as well as databases (COVID-19 Primer, 2020) and were mto apped to the reference genome using BLAST (Rice et al. 2000). The SARS-CoV-2 genomic variant coordinates were overlapped with the primer/probe binding sites. Tm and Gibbs free energy (ΔG) was calculated. We also evaluated the internal single mismatch as well as terminal mismatch which could have an impact on the thermodynamics stability of the nucleic acid secondary structure as well as on Tm **Supplementary Methods 1**.

Of the 938 genomes of SARS-CoV-2 isolates from India, a total of 717 were of high quality and were further considered for the variant calling **Supplementary Data 1**. This analysis revealed a total of 1,523 single nucleotide variants (SNVS) as well as 27 indels. We could compile a total of 132 primers or probe sequences **Supplementary Data 2**. A total of 123 SNVS and 2 indels mapped to at least one of the 80 odd primer/probe sites in the genome **Supplementary Data 3** of which, a total of 13 unique variants had allele frequency ≥ 1% Table 1 and Figure 1. Of significant note were three primers/probes for the *N* gene, which has variants mapping to the target sites in over 10% of Indian isolates. Variants with >1% frequency were also found in primer / probes encompassing *S, E, RdRP, ORF1a*, and *ORF3a* genes Table 1. One of the variant, 15451:G:A had a high frequency of 0.6% in Indian isolates and mapping to the WHO and ICMR-India recommended RdRP_SARSr-F2 “GTGARATGGTCATGTGTGGCGG” primer. Additionally, two indels with 3’ end terminal mismatch were in (N=6) 0.8% in Indian SARS-CoV-2 genomes. One of the primers is a part of WHO protocol E_sarbeco_R2 (Corman et al. 2020) and widely used (Eurofins Genomics, 2020). Out of these indels, one involves a 3 nt deletion from 3’ end of the primers while the other indel deletes 41 nt that encompasses the whole E_sarbeco_R2 primer.

**Table 1:**
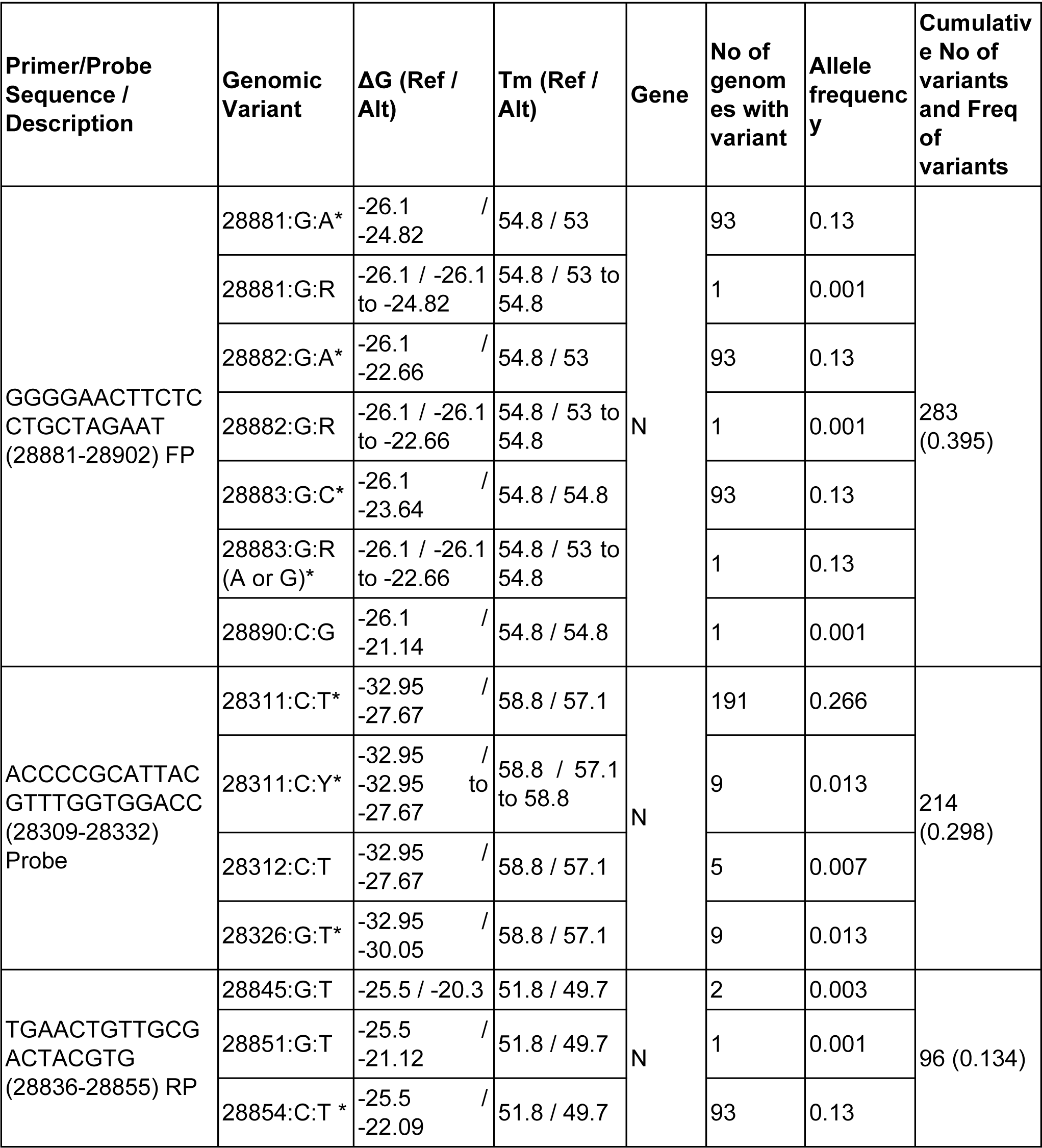
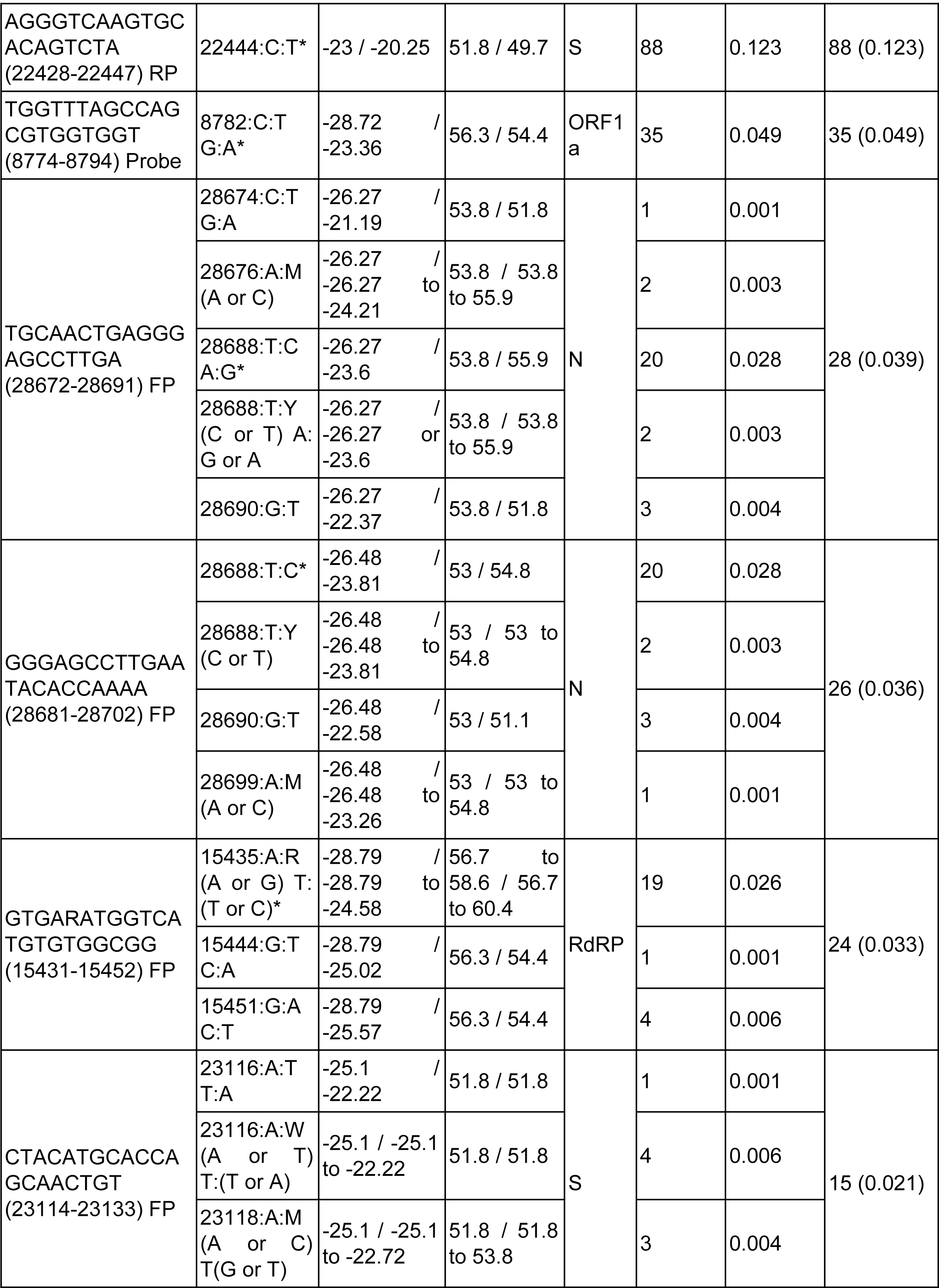
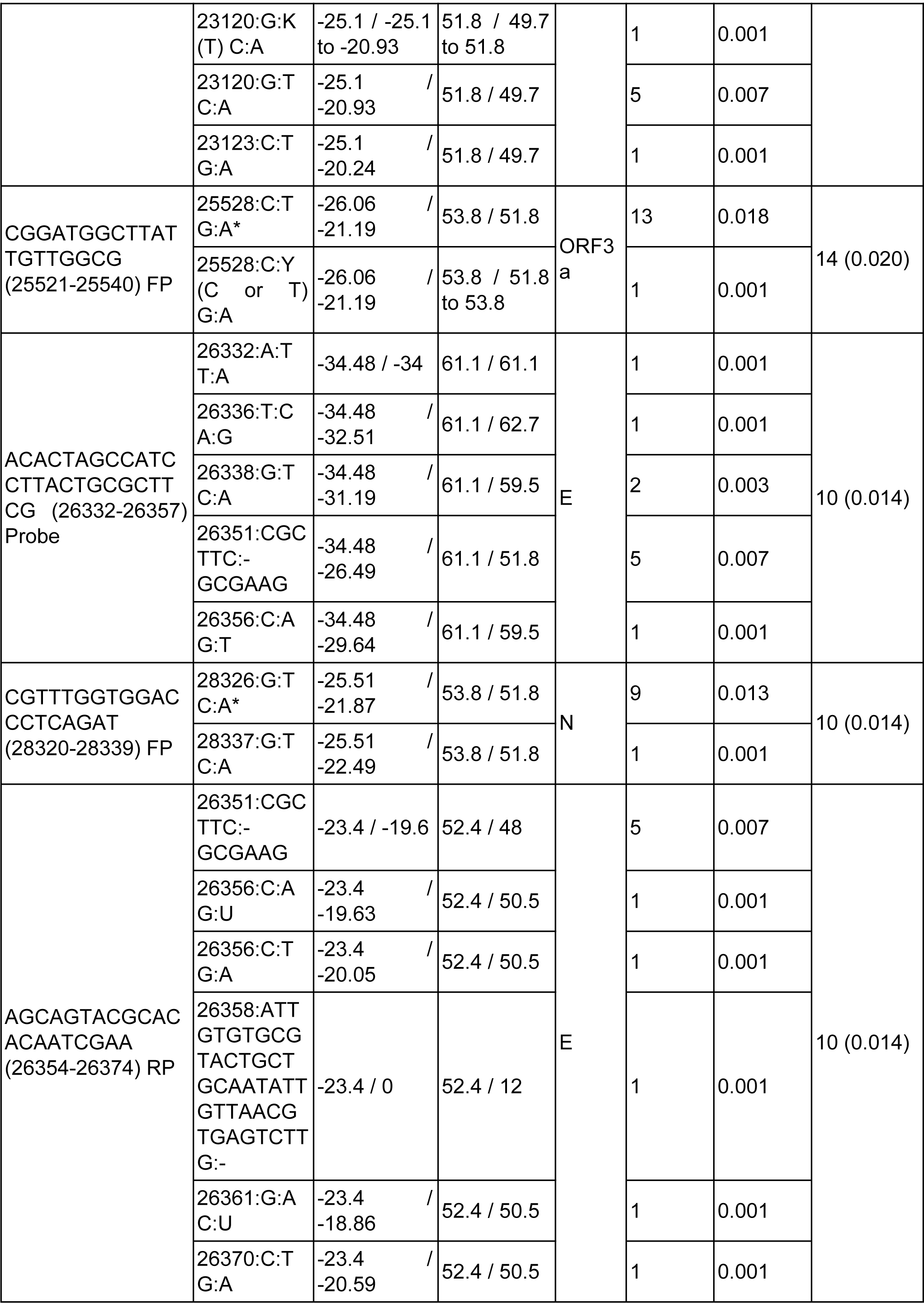
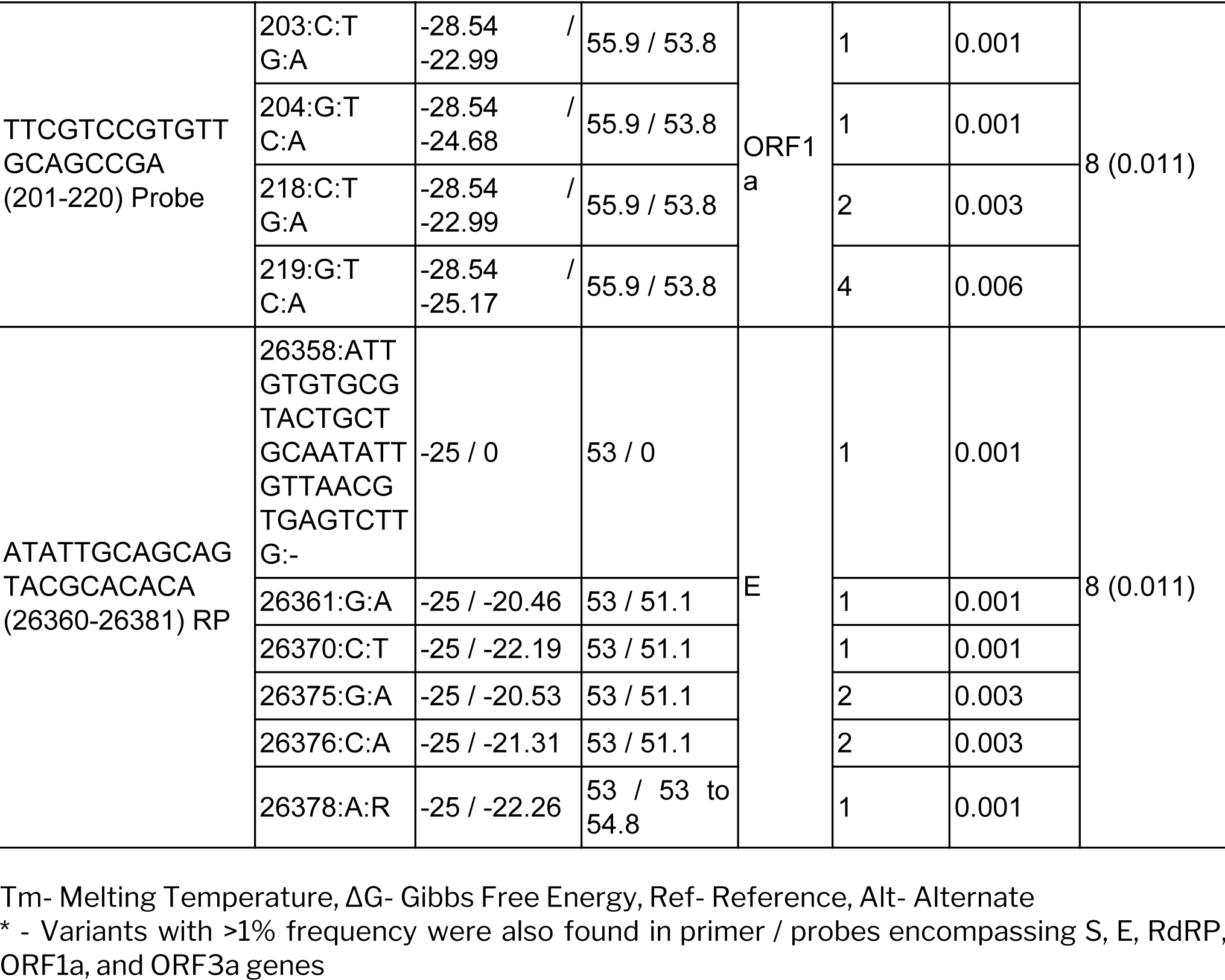
summary of Primer and Probe sequences and genomic variants analysis.

**Figure 1.**
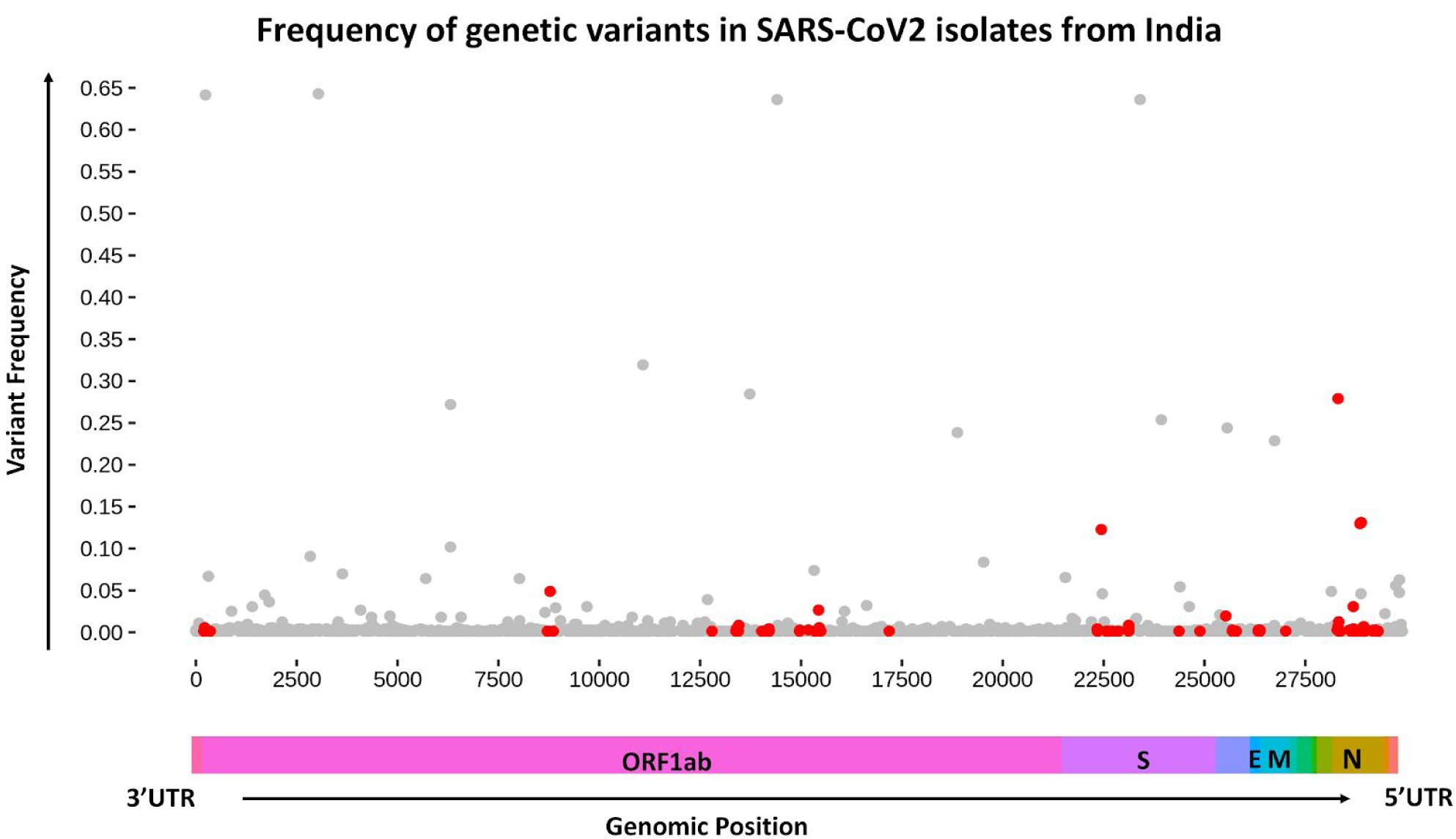
The frequency of genetic variants in SARS-CoV-2 isolates from India. The variants mapping to oligonucleotide primer/probe sites are marked in red and others in grey. The panel at the bottom depicts the SARS-CoV-2 genome with genomic annotation.

Our analysis suggests that genome sequencing of isolates in an epidemic could provide useful insights into assessing the diagnostic efficacies as also suggested by previous authors (Khan and Cheung 2020). We surmise that this could possibly drive policies on evaluation and approvals of the assays for screening and diagnosis. It has not escaped our attention that a number of genomic loci had significantly low variability in Indian isolates of SARS-CoV-2 isolates suggesting an opportunity to develop better molecular assays. To the best of our knowledge, this report is the most comprehensive report of the assessment of genomic variants and their impact on molecular assays for Indian isolates of SARS-CoV-2. The study highlights the need to widely share genome sequences of isolates as well as molecular probe information during epidemics.

## ACKNOWLEDGEMENTS

We acknowledge the researchers who have made the SARS-CoV-2 genomes available in the public domain. A comprehensive list of genomes, contributing laboratories, and acknowledgement is available in **Supplementary Data 1.** Authors acknowledge Paras Sehgal for constructive comments which enriched the manuscript.

Authors acknowledge funding from CSIR India. AJ and SM acknowledge a research fellowship from CSIR India. The funders had no role in the preparation of the manucsript or decision to publish.

## AUTHOR CONTRIBUTIONS

MR performed the genome analysis and variant calls. AJ and SM1 co-ordinated the compendium of primers and probes with help of Bhavya Balaji Krishnan, Manasa Sharma, Sreya Mandal, Teresa Fernandez and Sumayra Sultanji. SM2 contributed to mapping the primers to the genomic loci. AJ performed the analysis of variants mapping to the probe-target sites and was assisted by MR. VS and SS provided the conceptual overview to the analysis. VS, MR and AJ wrote the manuscript, the content and analysis which was read and agreed upon by all authors.

## SUPPLEMENTARY DATASETS

**Supplementary Methods 1:** Methodology for calculating primers/probes melting temperature and Gibbs free energy with variants impact.

**Supplementary Data 1.** Summary of the genomes of Indian isolates of SARS-CoV-2 available in public domain

**Supplementary Data 2.** Curated primers and probes sequence and their genomic coordinate used in the molecular assays for detection of SARS-CoV-2.

**Supplementary Data 3. Summary of Primer and Probe sequences and genomic variants** Calculated variant frequency, cumulative variant frequency, and melting temperature (Tm) for reference and alternate in the Indian SARS-CoV-2 isolates in the primers and probes binding sites.

