## Supplementary Data 1 for "Analysis of the potential impact of genomic variants in SARS-CoV-2 genomes from India on molecular diagnostic assays"

| Virus_name | GISAID ID | Collection_date | Originating_lab | Submitting_lab | Authors |
|---|---|---|---|---|---|
| hCoV-19/India/1-27/2020 | EPI_ISL_413522 | 1/27/2020 | Indian Council of Medical Research - National Institute of Virology | National Influenza Center, Indian Council of Medical Research - National Institute of Virology | Potdar V, Yadav PD, Choudhary ML, Shete-Aich A |
| hCoV-19/India/1-31/2020 | EPI_ISL_413523 | 1/31/2020 | Indian Council of Medical Research-National Institute of Virology | National Influenza Center, Indian Council of Medical Research-National Institute of Virology | Potdar V, Yadav PD, Choudhary ML, Shete-Aich A |
| hCoV-19/India/763/2020 | EPI_ISL_420543 | 3/3/2020 | National Influenza Center, Indian Council of Medical Research - National Institute of Virology | Indian Council of Medical Research-National Institute of Virology, Microbial Containment Complex | Pragya D. Yadav. Savita Patil, Varsha Potdar, Prasad Sarkale, Dimpal A. Nyayanit, Gajanan Sapkal, Anita M. Shete, Atanu Basu, Lalit Dar, M Choudhary, Amita Jain, Bharati Malhotra, Pranita Gawande, Sarah Cherian, Priya Abraham |
| hCoV-19/India/2020763/2020 | EPI_ISL_420544 | 2020 | Indian Council of Medical Research-National Institute of Virology, Microbial Containment Complex | Indian Council of Medical Research-National Institute of Virology, Microbial Containment Complex | Pragya D. Yadav. Savita Patil, Varsha Potdar, Prasad Sarkale, Dimpal A. Nyayanit, Gajanan Sapkal, Anita M. Shete, Atanu Basu, Lalit Dar, M Choudhary, Amita Jain, Bharati Malhotra, Pranita Gawande, Sarah Cherian, Priya Abraham |
| hCoV-19/India/770/2020 | EPI_ISL_420545 | 3/3/2020 | National Influenza Center, Indian Council of Medical Research - National Institute of Virology | Indian Council of Medical Research-National Institute of Virology, Microbial Containment Complex | Pragya D. Yadav. Savita Patil, Varsha Potdar, Prasad Sarkale, Dimpal A. Nyayanit, Gajanan Sapkal, Anita M. Shete, Atanu Basu, Lalit Dar, M Choudhary, Amita Jain, Bharati Malhotra, Pranita Gawande, Sarah Cherian, Priya Abraham |
| hCoV-19/India/2020770/2020 | EPI_ISL_420546 | 2020 | Indian Council of Medical Research-National Institute of Virology, Microbial Containment Complex | Indian Council of Medical Research-National Institute of Virology, Microbial Containment Complex | Pragya D. Yadav. Savita Patil, Varsha Potdar, Prasad Sarkale, Dimpal A. Nyayanit, Gajanan Sapkal, Anita M. Shete, Atanu Basu, Lalit Dar, M Choudhary, Amita Jain, Bharati Malhotra, Pranita Gawande, Sarah Cherian, Priya Abraham |
| hCoV-19/India/772/2020 | EPI_ISL_420547 | 3/3/2020 | National Influenza Center, Indian Council of Medical Research - National Institute of Virology | Indian Council of Medical Research-National Institute of Virology, Microbial Containment Complex | Pragya D. Yadav. Savita Patil, Varsha Potdar, Prasad Sarkale, Dimpal A. Nyayanit, Gajanan Sapkal, Anita M. Shete, Atanu Basu, Lalit Dar, M Choudhary, Amita Jain, Bharati Malhotra, Pranita Gawande, Sarah Cherian, Priya Abraham |
| hCoV-19/India/2020772/2020 | EPI_ISL_420548 | 2020 | Indian Council of Medical Research-National Institute of Virology, Microbial Containment Complex | Indian Council of Medical Research-National Institute of Virology, Microbial Containment Complex | Pragya D. Yadav. Savita Patil, Varsha Potdar, Prasad Sarkale, Dimpal A. Nyayanit, Gajanan Sapkal, Anita M. Shete, Atanu Basu, Lalit Dar, M Choudhary, Amita Jain, Bharati Malhotra, Pranita Gawande, Sarah Cherian, Priya Abraham |
| hCoV-19/India/773/2020 | EPI_ISL_420549 | 3/3/2020 | National Influenza Center, Indian Council of Medical Research - National Institute of Virology | Indian Council of Medical Research-National Institute of Virology, Microbial Containment Complex | Pragya D. Yadav. Savita Patil, Varsha Potdar, Prasad Sarkale, Dimpal A. Nyayanit, Gajanan Sapkal, Anita M. Shete, Atanu Basu, Lalit Dar, M Choudhary, Amita Jain, Bharati Malhotra, Pranita Gawande, Sarah Cherian, Priya Abraham |
| hCoV-19/India/2020773/2020 | EPI_ISL_420550 | 2020 | Indian Council of Medical Research-National Institute of Virology, Microbial Containment Complex | Indian Council of Medical Research-National Institute of Virology, Microbial Containment Complex | Pragya D. Yadav. Savita Patil, Varsha Potdar, Prasad Sarkale, Dimpal A. Nyayanit, Gajanan Sapkal, Anita M. Shete, Atanu Basu, Lalit Dar, M Choudhary, Amita Jain, Bharati Malhotra, Pranita Gawande, Sarah Cherian, Priya Abraham |
| hCoV-19/India/777/2020 | EPI_ISL_420551 | 3/3/2020 | National Influenza Center, Indian Council of Medical Research - National Institute of Virology | Indian Council of Medical Research-National Institute of Virology, Microbial Containment Complex | Pragya D. Yadav. Savita Patil, Varsha Potdar, Prasad Sarkale, Dimpal A. Nyayanit, Gajanan Sapkal, Anita M. Shete, Atanu Basu, Lalit Dar, M Choudhary, Amita Jain, Bharati Malhotra, Pranita Gawande, Sarah Cherian, Priya Abraham |
| hCoV-19/India/2020777/2020 | EPI_ISL_420552 | 2020 | Indian Council of Medical Research-National Institute of Virology, Microbial Containment Complex | Indian Council of Medical Research-National Institute of Virology, Microbial Containment Complex | Pragya D. Yadav. Savita Patil, Varsha Potdar, Prasad Sarkale, Dimpal A. Nyayanit, Gajanan Sapkal, Anita M. Shete, Atanu Basu, Lalit Dar, M Choudhary, Amita Jain, Bharati Malhotra, Pranita Gawande, Sarah Cherian, Priya Abraham |
| hCoV-19/India/781/2020 | EPI_ISL_420553 | 3/3/2020 | National Influenza Center, Indian Council of Medical Research - National Institute of Virology | Indian Council of Medical Research-National Institute of Virology, Microbial Containment Complex | Pragya D. Yadav. Savita Patil, Varsha Potdar, Prasad Sarkale, Dimpal A. Nyayanit, Gajanan Sapkal, Anita M. Shete, Atanu Basu, Lalit Dar, M Choudhary, Amita Jain, Bharati Malhotra, Pranita Gawande, Sarah Cherian, Priya Abraham |
| hCoV-19/India/2020781/2020 | EPI_ISL_420554 | 2020 | Indian Council of Medical Research-National Institute of Virology, Microbial Containment Complex | Indian Council of Medical Research-National Institute of Virology, Microbial Containment Complex | Pragya D. Yadav. Savita Patil, Varsha Potdar, Prasad Sarkale, Dimpal A. Nyayanit, Gajanan Sapkal, Anita M. Shete, Atanu Basu, Lalit Dar, M Choudhary, Amita Jain, Bharati Malhotra, Pranita Gawande, Sarah Cherian, Priya Abraham |
| hCoV-19/India/c32/2020 | EPI_ISL_420555 | 3/3/2020 | National Influenza Center, Indian Council of Medical Research - National Institute of Virology | Indian Council of Medical Research-National Institute of Virology, Microbial Containment Complex | Pragya D. Yadav. Savita Patil, Varsha Potdar, Prasad Sarkale, Dimpal A. Nyayanit, Gajanan Sapkal, Anita M. Shete, Atanu Basu, Lalit Dar, M Choudhary, Amita Jain, Bharati Malhotra, Pranita Gawande, Sarah Cherian, Priya Abraham |
| hCoV-19/India/2020c32/2020 | EPI_ISL_420556 | 2020 | Indian Council of Medical Research-National Institute of Virology, Microbial Containment Complex | Indian Council of Medical Research-National Institute of Virology, Microbial Containment Complex | Pragya D. Yadav. Savita Patil, Varsha Potdar, Prasad Sarkale, Dimpal A. Nyayanit, Gajanan Sapkal, Anita M. Shete, Atanu Basu, Lalit Dar, M Choudhary, Amita Jain, Bharati Malhotra, Pranita Gawande, Sarah Cherian, Priya Abraham |
| hCoV-19/India/1073/2020 | EPI_ISL_421662 | 3/10/2020 | National Influenza Center, Indian Council of Medical Research - National Institute of Virology | Indian Council of Medical Research-National Institute of Virology, Microbial Containment Complex | Pragya D. Yadav, Varsha Potdar, Savita Patil, Dimpal A. Nyayanit, Triparna Majumdar, Manohar. L. Chaudhary, Gururaj Deshpande, Padinjaremattathil Thankappan Ullas, Anita Shete-Aich, Hitesh Dighe, Sreelekshmy Mohandas, Gajanan Sapkal, Atanu Basu, Amita Jain, Bharti Malhotra, Deepika Chaudhary, Sarah Cherian, Priya Abraham |
| hCoV-19/India/1093/2020 | EPI_ISL_421663 | 3/10/2020 | National Influenza Center, Indian Council of Medical Research - National Institute of Virology | Indian Council of Medical Research-National Institute of Virology, Microbial Containment Complex | Pragya D. Yadav, Varsha Potdar, Savita Patil, Dimpal A. Nyayanit, Triparna Majumdar, Manohar. L. Chaudhary, Gururaj Deshpande, Padinjaremattathil Thankappan Ullas, Anita Shete-Aich, Hitesh Dighe, Sreelekshmy Mohandas, Gajanan Sapkal, Atanu Basu, Amita Jain, Bharti Malhotra, Deepika Chaudhary, Sarah Cherian, Priya Abraham |
| hCoV-19/India/1100/2020 | EPI_ISL_421664 | 3/10/2020 | National Influenza Center, Indian Council of Medical Research - National Institute of Virology | Indian Council of Medical Research-National Institute of Virology, Microbial Containment Complex | Pragya D. Yadav, Varsha Potdar, Savita Patil, Dimpal A. Nyayanit, Triparna Majumdar, Manohar. L. Chaudhary, Gururaj Deshpande, Padinjaremattathil Thankappan Ullas, Anita Shete-Aich, Hitesh Dighe, Sreelekshmy Mohandas, Gajanan Sapkal, Atanu Basu, Amita Jain, Bharti Malhotra, Deepika Chaudhary, Sarah Cherian, Priya Abraham |
| hCoV-19/India/1104/2020 | EPI_ISL_421665 | 3/10/2020 | National Influenza Center, Indian Council of Medical Research - National Institute of Virology | Indian Council of Medical Research-National Institute of Virology, Microbial Containment Complex | Pragya D. Yadav, Varsha Potdar, Savita Patil, Dimpal A. Nyayanit, Triparna Majumdar, Manohar. L. Chaudhary, Gururaj Deshpande, Padinjaremattathil Thankappan Ullas, Anita Shete-Aich, Hitesh Dighe, Sreelekshmy Mohandas, Gajanan Sapkal, Atanu Basu, Amita Jain, Bharti Malhotra, Deepika Chaudhary, Sarah Cherian, Priya Abraham |
| hCoV-19/India/1111/2020 | EPI_ISL_421666 | 3/10/2020 | National Influenza Center, Indian Council of Medical Research - National Institute of Virology | Indian Council of Medical Research-National Institute of Virology, Microbial Containment Complex | Pragya D. Yadav, Varsha Potdar, Savita Patil, Dimpal A. Nyayanit, Triparna Majumdar, Manohar. L. Chaudhary, Gururaj Deshpande, Padinjaremattathil Thankappan Ullas, Anita Shete-Aich, Hitesh Dighe, Sreelekshmy Mohandas, Gajanan Sapkal, Atanu Basu, Amita Jain, Bharti Malhotra, Deepika Chaudhary, Sarah Cherian, Priya Abraham |
| hCoV-19/India/1115/2020 | EPI_ISL_421667 | 3/10/2020 | National Influenza Center, Indian Council of Medical Research - National Institute of Virology | Indian Council of Medical Research-National Institute of Virology, Microbial Containment Complex | Pragya D. Yadav, Varsha Potdar, Savita Patil, Dimpal A. Nyayanit, Triparna Majumdar, Manohar. L. Chaudhary, Gururaj Deshpande, Padinjaremattathil Thankappan Ullas, Anita Shete-Aich, Hitesh Dighe, Sreelekshmy Mohandas, Gajanan Sapkal, Atanu Basu, Amita Jain, Bharti Malhotra, Deepika Chaudhary, Sarah Cherian, Priya Abraham |
| hCoV-19/India/1125/2020 | EPI_ISL_421668 | 3/10/2020 | National Influenza Center, Indian Council of Medical Research - National Institute of Virology | Indian Council of Medical Research-National Institute of Virology, Microbial Containment Complex | Pragya D. Yadav, Varsha Potdar, Savita Patil, Dimpal A. Nyayanit, Triparna Majumdar, Manohar. L. Chaudhary, Gururaj Deshpande, Padinjaremattathil Thankappan Ullas, Anita Shete-Aich, Hitesh Dighe, Sreelekshmy Mohandas, Gajanan Sapkal, Atanu Basu, Amita Jain, Bharti Malhotra, Deepika Chaudhary, Sarah Cherian, Priya Abraham |
| hCoV-19/India/1616/2020 | EPI_ISL_421669 | 3/12/2020 | National Influenza Center, Indian Council of Medical Research - National Institute of Virology | Indian Council of Medical Research-National Institute of Virology, Microbial Containment Complex | Pragya D. Yadav, Varsha Potdar, Savita Patil, Dimpal A. Nyayanit, Triparna Majumdar, Manohar. L. Chaudhary, Gururaj Deshpande, Padinjaremattathil Thankappan Ullas, Anita Shete-Aich, Hitesh Dighe, Sreelekshmy Mohandas, Gajanan Sapkal, Atanu Basu, Amita Jain, Bharti Malhotra, Deepika Chaudhary, Sarah Cherian, Priya Abraham |
| hCoV-19/India/1617/2020 | EPI_ISL_421670 | 3/12/2020 | National Influenza Center, Indian Council of Medical Research - National Institute of Virology | Indian Council of Medical Research-National Institute of Virology, Microbial Containment Complex | Pragya D. Yadav, Varsha Potdar, Savita Patil, Dimpal A. Nyayanit, Triparna Majumdar, Manohar. L. Chaudhary, Gururaj Deshpande, Padinjaremattathil Thankappan Ullas, Anita Shete-Aich, Hitesh Dighe, Sreelekshmy Mohandas, Gajanan Sapkal, Atanu Basu, Amita Jain, Bharti Malhotra, Deepika Chaudhary, Sarah Cherian, Priya Abraham |
| hCoV-19/India/1621/2020 | EPI_ISL_421671 | 3/12/2020 | National Influenza Center, Indian Council of Medical Research - National Institute of Virology | Indian Council of Medical Research-National Institute of Virology, Microbial Containment Complex | Pragya D. Yadav, Varsha Potdar, Savita Patil, Dimpal A. Nyayanit, Triparna Majumdar, Manohar. L. Chaudhary, Gururaj Deshpande, Padinjaremattathil Thankappan Ullas, Anita Shete-Aich, Hitesh Dighe, Sreelekshmy Mohandas, Gajanan Sapkal, Atanu Basu, Amita Jain, Bharti Malhotra, Deepika Chaudhary, Sarah Cherian, Priya Abraham |
| hCoV-19/India/1644/2020 | EPI_ISL_421672 | 3/12/2020 | National Influenza Center, Indian Council of Medical Research - National Institute of Virology | Indian Council of Medical Research-National Institute of Virology, Microbial Containment Complex | Pragya D. Yadav, Varsha Potdar, Savita Patil, Dimpal A. Nyayanit, Triparna Majumdar, Manohar. L. Chaudhary, Gururaj Deshpande, Padinjaremattathil Thankappan Ullas, Anita Shete-Aich, Hitesh Dighe, Sreelekshmy Mohandas, Gajanan Sapkal, Atanu Basu, Amita Jain, Bharti Malhotra, Deepika Chaudhary, Sarah Cherian, Priya Abraham |
| hCoV-19/India/1063/2020 | EPI_ISL_424361 | 3/10/2020 | National Influenza Center, Indian Council of Medical Research - National Institute of Virology | Indian Council of Medical Research-National Institute of Virology, Microbial Containment Complex | Pragya D. Yadav, Varsha Potdar, Savita Patil, Dimpal A. Nyayanit, Triparna Majumdar, Manohar. L. Chaudhary, Gururaj Deshpande, Padinjaremattathil Thankappan Ullas, Anita Shete-Aich, Hitesh Dighe, Sreelekshmy Mohandas, Gajanan Sapkal, Atanu Basu, Amita Jain, Bharti Malhotra, Deepika Chaudhary, Sarah Cherian, Priya Abraham |
| hCoV-19/India/1135/2020 | EPI_ISL_424362 | 3/10/2020 | National Influenza Center, Indian Council of Medical Research - National Institute of Virology | Indian Council of Medical Research-National Institute of Virology, Microbial Containment Complex | Pragya D. Yadav, Varsha Potdar, Savita Patil, Dimpal A. Nyayanit, Triparna Majumdar, Manohar. L. Chaudhary, Gururaj Deshpande, Padinjaremattathil Thankappan Ullas, Anita Shete-Aich, Hitesh Dighe, Sreelekshmy Mohandas, Gajanan Sapkal, Atanu Basu, Amita Jain, Bharti Malhotra, Deepika Chaudhary, Sarah Cherian, Priya Abraham |
| hCoV-19/India/1652/2020 | EPI_ISL_424363 | 3/12/2020 | National Influenza Center, Indian Council of Medical Research - National Institute of Virology | Indian Council of Medical Research-National Institute of Virology, Microbial Containment Complex | Pragya D. Yadav, Varsha Potdar, Savita Patil, Dimpal A. Nyayanit, Triparna Majumdar, Manohar. L. Chaudhary, Gururaj Deshpande, Padinjaremattathil Thankappan Ullas, Anita Shete-Aich, Hitesh Dighe, Sreelekshmy Mohandas, Gajanan Sapkal, Atanu Basu, Amita Jain, Bharti Malhotra, Deepika Chaudhary, Sarah Cherian, Priya Abraham |
| hCoV-19/India/3118/2020 | EPI_ISL_424364 | 3/17/2020 | National Influenza Center, Indian Council of Medical Research - National Institute of Virology | Indian Council of Medical Research-National Institute of Virology, Microbial Containment Complex | Pragya D. Yadav, Varsha Potdar, Savita Patil, Dimpal A. Nyayanit, Triparna Majumdar, Manohar. L. Chaudhary, Gururaj Deshpande, Padinjaremattathil Thankappan Ullas, Anita Shete-Aich, Hitesh Dighe, Sreelekshmy Mohandas, Gajanan Sapkal, Atanu Basu, Amita Jain, Bharti Malhotra, Deepika Chaudhary, Sarah Cherian, Priya Abraham |
| hCoV-19/India/3239/2020 | EPI_ISL_424365 | 3/17/2020 | National Influenza Center, Indian Council of Medical Research - National Institute of Virology | Indian Council of Medical Research-National Institute of Virology, Microbial Containment Complex | Pragya D. Yadav, Varsha Potdar, Savita Patil, Dimpal A. Nyayanit, Triparna Majumdar, Manohar. L. Chaudhary, Gururaj Deshpande, Padinjaremattathil Thankappan Ullas, Anita Shete-Aich, Hitesh Dighe, Sreelekshmy Mohandas, Gajanan Sapkal, Atanu Basu, Amita Jain, Bharti Malhotra, Deepika Chaudhary, Sarah Cherian, Priya Abraham |
| hCoV-19/India/c31/2020 | EPI_ISL_426179 | 3/2/2020 | National Influenza Center, Indian Council of Medical Research - National Institute of Virology | Indian Council of Medical Research-National Institute of Virology, Microbial Containment Complex | Pragya D. Yadav, Varsha Potdar, Savita Patil, Dimpal A. Nyayanit, Triparna Majumdar, Manohar. L. Chaudhary, Gururaj Deshpande, Padinjaremattathil Thankappan Ullas, Anita Shete-Aich, Hitesh Dighe, Sreelekshmy Mohandas, Gajanan Sapkal, Atanu Basu, Amita Jain, Bharti Malhotra, Deepika Chaudhary, Sarah Cherian, Priya Abraham |
| hCoV-19/India/GBRC1/2020 | EPI_ISL_426414 | 4/5/2020 | Sir M P Shah Government Medical College | Gujarat Biotechnology Research Centre | Ramesh Pandit, Tejas Shah, Ankit Hinsu, Pritesh Sabara, Apurvasinh Puvar, Janvi Raval, Monika Gandhi, Pinal Trivedi, Maharshi Pandya, Amit Kanani, Akanksha Verma, Nitin Savaliya, Raghawendra Kumar, Dinesh Kumar, Zubair Saiyed, Dipa Kinariwala, Disha Patel, Binita Aring, Geeta Vaghela, Sonia Barve, Bhavesh Modi, Kairavi Joshi, Nidhi Sood, Pranay Shah, R D Dixit, Snehal Bagatharia, Madhvi Joshi, C |
| hCoV-19/India/GBRC1s/2020 | EPI_ISL_426415 | 4/5/2020 | Sir M P Shah Government Medical College, Jamnagar | Gujarat Biotechnology Research Centre, Gandhinagar | Ramesh Pandit, Tejas Shah, Ankit Hinsu, Pritesh Sabara, Apurvasinh Puvar, Janvi Raval, Monika Gandhi, Pinal Trivedi, Maharshi Pandya, Amit Kanani, Akanksha Verma, Nitin Savaliya, Raghvendra Kumar, Dinesh Kumar, Zuber Saiyed, Dipa Kinariwala, Disha Patel, Binita Aring, Geeta Vaghela, Sonia Barve, Bhavesh Modi, Kairavi Joshi, Nidhi Sood, Pranay Shah, R D Dixit, Snehal Bagatharia, Madhvi Joshi, Cha |
| hCoV-19/India/S3/2020 | EPI_ISL_430464 | 3/21/2020 | ICMR-National Institute of Cholera and Enteric Diseases | National Institute of Biomedical Genomics | Arindam Maitra, Mamta Chawla Sarkar, Sreedhar Chinnaswamy, Hasina Banu, Ananya Chatterjee, Shanta Dutta, Saumitra Das |
| hCoV-19/India/S5/2020 | EPI_ISL_430465 | 3/28/2020 | ICMR-National Institute of Cholera and Enteric Diseases | National Institute of Biomedical Genomics | Arindam Maitra, Mamta Chawla Sarkar, Sreedhar Chinnaswamy, Hasina Banu, Ananya Chatterjee, Shanta Dutta, Saumitra Das |
| hCoV-19/India/S6/2020 | EPI_ISL_430466 | 3/26/2020 | ICMR-National Institute of Cholera and Enteric Diseases | National Institute of Biomedical Genomics | Arindam Maitra, Mamta Chawla Sarkar, Sreedhar Chinnaswamy, Hasina Banu, Ananya Chatterjee, Shanta Dutta, Saumitra Das |
| hCoV-19/India/S11/2020 | EPI_ISL_430467 | 4/3/2020 | ICMR-National Institute of Cholera and Enteric Diseases | National Institute of Biomedical Genomics | Arindam Maitra, Mamta Chawla Sarkar, Sreedhar Chinnaswamy, Hasina Banu, Ananya Chatterjee, Shanta Dutta, Saumitra Das |
| hCoV-19/India/S2/2020 | EPI_ISL_430468 | 3/21/2020 | ICMR-National Institute of Cholera and Enteric Diseases | National Institute of Biomedical Genomics | Arindam Maitra, Mamta Chawla Sarkar, Sreedhar Chinnaswamy, Hasina Banu, Ananya Chatterjee, Shanta Dutta, Saumitra Das |
| hCoV-19/India/GMC-RK100/2020 | EPI_ISL_431101 | 3/1/2020 | Department of Microbiology,Gandhi Medical College and Hospital | Virus Research Laboratory, Department of Zoology, Osmania University,Hyderabad,India | Muttineni Radhakrishna, Nagamani K, Thrilok Chander B, Raja Rao M, Kalyani Putty, Ravikumar P, Sunitha P, Pankaj Singh D, Anand Kumar K, Amit A. Upadhyay Steven E. Bosinger, Rama Amara |
| hCoV-19/India/GMC-KN318/2020 | EPI_ISL_431102 | 3/11/2020 | Department of MicroBiology,Gandhi Medical College and Hospital,Secendrabad,Hyderabad,India | Department of Microbiology, Gandhi Medical College and Hospital, Secendrabad, Hyderabad | Nagamani K, Muttineni Radhakrishna, Thrilok Chander B, Raja Rao M, Kalyani Putty, Ravikumar P, Sunitha P, Pankaj Singh D, Anand Kumar K, Amit A. Upadhyay, Steven E. Bosinger, Rama Amara |
| hCoV-19/India/GMC-KN443/2020 | EPI_ISL_431103 | 3/16/2020 | Department of Microbiology, Gandhi Medical College and Hospital, Secendrabad, Hyderabad, India | Department of Microbiology, Gandhi Medical College and Hospital, Secendrabad, Hyderabad, India | Nagamani K, Muttineni Radhakrishna, Thrilok Chander B, Raja Rao M, Kalyani Putty, Ravikumar P, Sunitha P, Pankaj Singh D, Anand Kumar K, Amit A. Upadhyay, Steven E. Bosinger, Rama Amara |
| hCoV-19/India/GMC-TC469/2020 | EPI_ISL_431117 | 3/20/2020 | Department of Microbiology, Gandhi Medical College and Hospital, Secendrabad, Hyderabad, India | Department of Microbiology, Gandhi Medical College and Hospital, Secendrabad, Hyderabad, India | Thrilok Chander B, Muttineni Radhakrishna, Nagamani K, Raja Rao M, Kalyani Putty, Ravikumar P, Sunitha P, Pankaj Singh D, Anand Kumar K, Amit A. Upadhyay, Steven E.Bosinger, Rama Amara |
| hCoV-19/India/GBRC2/2020 | EPI_ISL_435049 | 4/13/2020 | B.J. Medical College and Civil hospital | Gujarat Biotechnology Research Centre | Pinal Trivedi, Maharshi Pandya, Amit Kanani, Akanksha Verma, Nitin Savaliya, Raghawendra Kumar, Dinesh Kumar, Zuber Saiyed, Dipa Kinariwala, Disha Patel, Binita Aring, Geeta Vaghela, Sonia Barve, Bhavesh Modi, Kairavi Joshi, Gaurishankar Shrimali, Nidhi Sood, Pranay Shah, R D Dixit, Snehal Bagatharia, Kamlesh J Upadhyay, Ramesh Pandit, Tejas Shah, Ankit Hinsu, Pritesh Sabara, Apurvasinh Puvar, |
| hCoV-19/India/GBRC3/2020 | EPI_ISL_435050 | 4/13/2020 | B.J. Medical College and Civil hospital | Gujarat Biotechnology Research Centre | Ankit Hinsu, Pritesh Sabara, Apurvasinh Puvar, Janvi Raval, Monika Gandhi, Pinal Trivedi, Maharshi Pandya, Amit Kanani, Akanksha Verma, Nitin Savaliya, Raghawendra Kumar, Dinesh Kumar, Zuber Saiyed, Dipa Kinariwala, Disha Patel, Binita Aring, Geeta Vaghela, Sonia Barve, Bhavesh Modi, Kairavi Joshi, Gaurishankar Shrimali, Nidhi Sood, Pranay Shah, R D Dixit, Snehal Bagatharia, Kamlesh J Upadhyay, |
| hCoV-19/India/GBRC4/2020 | EPI_ISL_435051 | 4/13/2020 | B.J. Medical College and Civil hospital | Gujarat Biotechnology Research Centre | Pritesh Sabara, Apurvasinh Puvar, Janvi Raval, Monika Gandhi, Pinal Trivedi, Maharshi Pandya, Amit Kanani, Akanksha Verma, Nitin Savaliya, Raghawendra Kumar, Dinesh Kumar, Zuber Saiyed, Dipa Kinariwala, Disha Patel, Binita Aring, Geeta Vaghela, Sonia Barve, Bhavesh Modi, Kairavi Joshi, Gaurishankar Shrimali, Nidhi Sood, Pranay Shah, R D Dixit, Snehal Bagatharia, Kamlesh J Upadhyay, Ramesh Pa |
| hCoV-19/India/GBRC6/2020 | EPI_ISL_435053 | 4/7/2020 | B.J. Medical College and Civil hospital | Gujarat Biotechnology Research Centre | Janvi Raval, Monika Gandhi, Pinal Trivedi, Maharshi Pandya, Amit Kanani, Akanksha Verma, Nitin Savaliya, Raghawendra Kumar, Dinesh Kumar, Zuber Saiyed, Dipa Kinariwala, Disha Patel, Binita Aring, Geeta Vaghela, Sonia Barve, Bhavesh Modi, Kairavi Joshi, Gaurishankar Shrimali, Nidhi Sood, Pranay Shah, R D Dixit, Snehal Bagatharia, Kamlesh J Upadhyay, Ramesh Pandit, Tejas Shah, Ankit Hinsu, Prites |
| hCoV-19/India/GBRC7/2020 | EPI_ISL_435054 | 4/14/2020 | B.J. Medical College and Civil hospital | Gujarat Biotechnology Research Centre | Monika Gandhi, Pinal Trivedi, Maharshi Pandya, Amit Kanani, Akanksha Verma, Nitin Savaliya, Raghawendra Kumar, Dinesh Kumar, Zuber Saiyed, Dipa Kinariwala, Disha Patel, Binita Aring, Geeta Vaghela, Sonia Barve, Bhavesh Modi, Kairavi Joshi, Gaurishankar Shrimali, Nidhi Sood, Pranay Shah, R D Dixit, Snehal Bagatharia, Kamlesh J Upadhyay, Ramesh Pandit, Tejas Shah, Ankit Hinsu, Pritesh Sabara, A |
| hCoV-19/India/GBRC8/2020 | EPI_ISL_435055 | 4/22/2020 | Gujarat Biotechnology Research Centre | Gujarat Biotechnology Research Centre | Tejas Shah, Ankit Hinsu, Pritesh Sabara, Apurvasinh Puvar, Janvi Raval, Monika Gandhi, Pinal Trivedi, Maharshi Pandya, Amit Kanani, Akanksha Verma, Nitin Savaliya, Raghawendra Kumar, Dinesh Kumar, Zuber Saiyed, Dipa Kinariwala, Disha Patel, Binita Aring, Geeta Vaghela, Sonia Barve, Bhavesh Modi, Kairavi Joshi, Gaurishankar Shrimali, Nidhi Sood, Pranay Shah, R D Dixit, Snehal Bagatharia, Kamlesh |
| hCoV-19/India/GBRC9/2020 | EPI_ISL_435056 | 4/21/2020 | Gujarat Biotechnology Research Centre | Gujarat Biotechnology Research Centre | Maharshi Pandya, Amit Kanani, Akanksha Verma, Nitin Savaliya, Raghawendra Kumar, Dinesh Kumar, Zuber Saiyed, Dipa Kinariwala, Disha Patel, Binita Aring, Geeta Vaghela, Sonia Barve, Bhavesh Modi, Kairavi Joshi, Gaurishankar Shrimali, Nidhi Sood, Pranay Shah, R D Dixit, Snehal Bagatharia, Kamlesh J Upadhyay, Ramesh Pandit, Tejas Shah, Ankit Hinsu, Pritesh Sabara, Apurvasinh Puvar, Janvi Raval, |
| hCoV-19/India/NCDC-01257/2020 | EPI_ISL_435060 | 3/12/2020 | National Centre for Disease control (NCDC), CSIR-Institute of Genomics and Integrative Biology (CSIR-IGIB) | NCDC/CSIR-IGIB | Pramod Kumar, Rajesh Pandey, Pooja Sharma, Mahesh Dhar, Vivekanand A, Bharathram Uppili, Himanshu Vashisht, Saruchi Wadhwa, Nishu Tyagi, Uma Sharma, Priyanka Singh, Hemlata Lall, Meena Datta, Poonam Gupta, Nidhi Saini, Aarti Tewari, Bibhash Nandi, Dhirendra Kumar, Satyabrata Bag, Varun Jaiswal, Hema Gogia, Preeti Madan, Simrita Singh, Prateek Singh, Debasis Dash, Mitali Mukerji, Manju |
| hCoV-19/India/NCDC-01537/2020 | EPI_ISL_435061 | 3/16/2020 | National Centre for Disease control (NCDC), CSIR-Institute of Genomics and Integrative Biology (CSIR-IGIB) | NCDC/CSIR-IGIB | Pramod Kumar, Rajesh Pandey, Pooja Sharma, Mahesh Dhar, Vivekanand A, Bharathram Uppili, Himanshu Vashisht, Saruchi Wadhwa, Nishu Tyagi, Uma Sharma, Priyanka Singh, Hemlata Lall, Meena Datta, Poonam Gupta, Nidhi Saini, Aarti Tewari, Bibhash Nandi, Dhirendra Kumar, Satyabrata Bag, Varun Jaiswal, Hema Gogia, Preeti Madan, Simrita Singh, Prateek Singh, Debasis Dash, Mitali Mukerji, Manju |
| hCoV-19/India/NCDC-01538/2020 | EPI_ISL_435062 | 3/16/2020 | National Centre for Disease control (NCDC), CSIR-Institute of Genomics and Integrative Biology (CSIR-IGIB) | NCDC/CSIR-IGIB | Pramod Kumar, Rajesh Pandey, Pooja Sharma, Mahesh Dhar, Vivekanand A, Bharathram Uppili, Himanshu Vashisht, Saruchi Wadhwa, Nishu Tyagi, Uma Sharma, Priyanka Singh, Hemlata Lall, Meena Datta, Poonam Gupta, Nidhi Saini, Aarti Tewari, Bibhash Nandi, Dhirendra Kumar, Satyabrata Bag, Varun Jaiswal, Hema Gogia, Preeti Madan, Simrita Singh, Prateek Singh, Debasis Dash, Mitali Mukerji, Manju |
| hCoV-19/India/NCDC-01326/2020 | EPI_ISL_435063 | 3/13/2020 | National Centre for Disease control (NCDC), CSIR-Institute of Genomics and Integrative Biology (CSIR-IGIB) | NCDC/CSIR-IGIB | Pramod Kumar, Rajesh Pandey, Pooja Sharma, Mahesh Dhar, Vivekanand A, Bharathram Uppili, Himanshu Vashisht, Saruchi Wadhwa, Nishu Tyagi, Uma Sharma, Priyanka Singh, Hemlata Lall, Meena Datta, Poonam Gupta, Nidhi Saini, Aarti Tewari, Bibhash Nandi, Dhirendra Kumar, Satyabrata Bag, Varun Jaiswal, Hema Gogia, Preeti Madan, Simrita Singh, Prateek Singh, Debasis Dash, Mitali Mukerji, Manju |
| hCoV-19/India/NCDC-01705/2020 | EPI_ISL_435064 | 3/18/2020 | National Centre for Disease control (NCDC), CSIR-Institute of Genomics and Integrative Biology (CSIR-IGIB) | NCDC/CSIR-IGIB | Pramod Kumar, Rajesh Pandey, Pooja Sharma, Mahesh Dhar, Vivekanand A, Bharathram Uppili, Himanshu Vashisht, Saruchi Wadhwa, Nishu Tyagi, Uma Sharma, Priyanka Singh, Hemlata Lall, Meena Datta, Poonam Gupta, Nidhi Saini, Aarti Tewari, Bibhash Nandi, Dhirendra Kumar, Satyabrata Bag, Varun Jaiswal, Hema Gogia, Preeti Madan, Simrita Singh, Prateek Singh, Debasis Dash, Mitali Mukerji, Manju |
| hCoV-19/India/NCDC-01475/2020 | EPI_ISL_435065 | 3/15/2020 | National Centre for Disease control (NCDC), CSIR-Institute of Genomics and Integrative Biology (CSIR-IGIB) | NCDC/CSIR-IGIB | Pramod Kumar, Rajesh Pandey, Pooja Sharma, Mahesh Dhar, Vivekanand A, Bharathram Uppili, Himanshu Vashisht, Saruchi Wadhwa, Nishu Tyagi, Uma Sharma, Priyanka Singh, Hemlata Lall, Meena Datta, Poonam Gupta, Nidhi Saini, Aarti Tewari, Bibhash Nandi, Dhirendra Kumar, Satyabrata Bag, Varun Jaiswal, Hema Gogia, Preeti Madan, Simrita Singh, Prateek Singh, Debasis Dash, Mitali Mukerji, Manju |
| hCoV-19/India/NCDC-01711/2020 | EPI_ISL_435066 | 3/18/2020 | National Centre for Disease control (NCDC), CSIR-Institute of Genomics and Integrative Biology (CSIR-IGIB) | NCDC/CSIR-IGIB | Pramod Kumar, Rajesh Pandey, Pooja Sharma, Mahesh Dhar, Vivekanand A, Bharathram Uppili, Himanshu Vashisht, Saruchi Wadhwa, Nishu Tyagi, Uma Sharma, Priyanka Singh, Hemlata Lall, Meena Datta, Poonam Gupta, Nidhi Saini, Aarti Tewari, Bibhash Nandi, Dhirendra Kumar, Satyabrata Bag, Varun Jaiswal, Hema Gogia, Preeti Madan, Simrita Singh, Prateek Singh, Debasis Dash, Mitali Mukerji, Manju |
| hCoV-19/India/NCDC-01712/2020 | EPI_ISL_435067 | 3/18/2020 | National Centre for Disease control (NCDC), CSIR-Institute of Genomics and Integrative Biology (CSIR-IGIB) | NCDC/CSIR-IGIB | Pramod Kumar, Rajesh Pandey, Pooja Sharma, Mahesh Dhar, Vivekanand A, Bharathram Uppili, Himanshu Vashisht, Saruchi Wadhwa, Nishu Tyagi, Uma Sharma, Priyanka Singh, Hemlata Lall, Meena Datta, Poonam Gupta, Nidhi Saini, Aarti Tewari, Bibhash Nandi, Dhirendra Kumar, Satyabrata Bag, Varun Jaiswal, Hema Gogia, Preeti Madan, Simrita Singh, Prateek Singh, Debasis Dash, Mitali Mukerji, Manju |
| hCoV-19/India/NCDC-01757/2020 | EPI_ISL_435068 | 3/18/2020 | National Centre for Disease control (NCDC), CSIR-Institute of Genomics and Integrative Biology (CSIR-IGIB) | NCDC/CSIR-IGIB | Pramod Kumar, Rajesh Pandey, Pooja Sharma, Mahesh Dhar, Vivekanand A, Bharathram Uppili, Himanshu Vashisht, Saruchi Wadhwa, Nishu Tyagi, Uma Sharma, Priyanka Singh, Hemlata Lall, Meena Datta, Poonam Gupta, Nidhi Saini, Aarti Tewari, Bibhash Nandi, Dhirendra Kumar, Satyabrata Bag, Varun Jaiswal, Hema Gogia, Preeti Madan, Simrita Singh, Prateek Singh, Debasis Dash, Mitali Mukerji, Manju |
| hCoV-19/India/NCDC-01744/2020 | EPI_ISL_435069 | 3/18/2020 | National Centre for Disease control (NCDC), CSIR-Institute of Genomics and Integrative Biology (CSIR-IGIB) | NCDC/CSIR-IGIB | Pramod Kumar, Rajesh Pandey, Pooja Sharma, Mahesh Dhar, Vivekanand A, Bharathram Uppili, Himanshu Vashisht, Saruchi Wadhwa, Nishu Tyagi, Uma Sharma, Priyanka Singh, Hemlata Lall, Meena Datta, Poonam Gupta, Nidhi Saini, Aarti Tewari, Bibhash Nandi, Dhirendra Kumar, Satyabrata Bag, Varun Jaiswal, Hema Gogia, Preeti Madan, Simrita Singh, Prateek Singh, Debasis Dash, Mitali Mukerji, Manju |
| hCoV-19/India/NCDC-01710/2020 | EPI_ISL_435070 | 3/18/2020 | National Centre for Disease control (NCDC), CSIR-Institute of Genomics and Integrative Biology (CSIR-IGIB) | NCDC/CSIR-IGIB | Pramod Kumar, Rajesh Pandey, Pooja Sharma, Mahesh Dhar, Vivekanand A, Bharathram Uppili, Himanshu Vashisht, Saruchi Wadhwa, Nishu Tyagi, Uma Sharma, Priyanka Singh, Hemlata Lall, Meena Datta, Poonam Gupta, Nidhi Saini, Aarti Tewari, Bibhash Nandi, Dhirendra Kumar, Satyabrata Bag, Varun Jaiswal, Hema Gogia, Preeti Madan, Simrita Singh, Prateek Singh, Debasis Dash, Mitali Mukerji, Manju |
| hCoV-19/India/NCDC-02105/2020 | EPI_ISL_435071 | 3/25/2020 | National Centre for Disease control (NCDC), CSIR-Institute of Genomics and Integrative Biology (CSIR-IGIB) | NCDC/CSIR-IGIB | Pramod Kumar, Rajesh Pandey, Pooja Sharma, Mahesh Dhar, Vivekanand A, Bharathram Uppili, Himanshu Vashisht, Saruchi Wadhwa, Nishu Tyagi, Uma Sharma, Priyanka Singh, Hemlata Lall, Meena Datta, Poonam Gupta, Nidhi Saini, Aarti Tewari, Bibhash Nandi, Dhirendra Kumar, Satyabrata Bag, Varun Jaiswal, Hema Gogia, Preeti Madan, Simrita Singh, Prateek Singh, Debasis Dash, Mitali Mukerji, Manju |
| hCoV-19/India/NCDC-02250/2020 | EPI_ISL_435074 | 3/28/2020 | National Centre for Disease control (NCDC), CSIR-Institute of Genomics and Integrative Biology (CSIR-IGIB) | NCDC/CSIR-IGIB | Pramod Kumar, Rajesh Pandey, Pooja Sharma, Mahesh Dhar, Vivekanand A, Bharathram Uppili, Himanshu Vashisht, Saruchi Wadhwa, Nishu Tyagi, Uma Sharma, Priyanka Singh, Hemlata Lall, Meena Datta, Poonam Gupta, Nidhi Saini, Aarti Tewari, Bibhash Nandi, Dhirendra Kumar, Satyabrata Bag, Varun Jaiswal, Hema Gogia, Preeti Madan, Simrita Singh, Prateek Singh, Debasis Dash, Mitali Mukerji, Manju |
| hCoV-19/India/NCDC-02311/2020 | EPI_ISL_435078 | 3/29/2020 | National Centre for Disease control (NCDC), CSIR-Institute of Genomics and Integrative Biology (CSIR-IGIB) | NCDC/CSIR-IGIB | Pramod Kumar, Rajesh Pandey, Pooja Sharma, Mahesh Dhar, Vivekanand A, Bharathram Uppili, Himanshu Vashisht, Saruchi Wadhwa, Nishu Tyagi, Uma Sharma, Priyanka Singh, Hemlata Lall, Meena Datta, Poonam Gupta, Nidhi Saini, Aarti Tewari, Bibhash Nandi, Dhirendra Kumar, Satyabrata Bag, Varun Jaiswal, Hema Gogia, Preeti Madan, Simrita Singh, Prateek Singh, Debasis Dash, Mitali Mukerji, Manju |
| hCoV-19/India/NCDC-02248/2020 | EPI_ISL_435081 | 3/28/2020 | National Centre for Disease control (NCDC), CSIR-Institute of Genomics and Integrative Biology (CSIR-IGIB) | NCDC/CSIR-IGIB | Pramod Kumar, Rajesh Pandey, Pooja Sharma, Mahesh Dhar, Vivekanand A, Bharathram Uppili, Himanshu Vashisht, Saruchi Wadhwa, Nishu Tyagi, Uma Sharma, Priyanka Singh, Hemlata Lall, Meena Datta, Poonam Gupta, Nidhi Saini, Aarti Tewari, Bibhash Nandi, Dhirendra Kumar, Satyabrata Bag, Varun Jaiswal, Hema Gogia, Preeti Madan, Simrita Singh, Prateek Singh, Debasis Dash, Mitali Mukerji, Manju |
| hCoV-19/India/NCDC-02242/2020 | EPI_ISL_435083 | 3/28/2020 | National Centre for Disease control (NCDC), CSIR-Institute of Genomics and Integrative Biology (CSIR-IGIB) | NCDC/CSIR-IGIB | Pramod Kumar, Rajesh Pandey, Pooja Sharma, Mahesh Dhar, Vivekanand A, Bharathram Uppili, Himanshu Vashisht, Saruchi Wadhwa, Nishu Tyagi, Uma Sharma, Priyanka Singh, Hemlata Lall, Meena Datta, Poonam Gupta, Nidhi Saini, Aarti Tewari, Bibhash Nandi, Dhirendra Kumar, Satyabrata Bag, Varun Jaiswal, Hema Gogia, Preeti Madan, Simrita Singh, Prateek Singh, Debasis Dash, Mitali Mukerji, Manju |
| hCoV-19/India/NCDC-02309/2020 | EPI_ISL_435084 | 3/29/2020 | National Centre for Disease control (NCDC), CSIR-Institute of Genomics and Integrative Biology (CSIR-IGIB) | NCDC/CSIR-IGIB | Pramod Kumar, Rajesh Pandey, Pooja Sharma, Mahesh Dhar, Vivekanand A, Bharathram Uppili, Himanshu Vashisht, Saruchi Wadhwa, Nishu Tyagi, Uma Sharma, Priyanka Singh, Hemlata Lall, Meena Datta, Poonam Gupta, Nidhi Saini, Aarti Tewari, Bibhash Nandi, Dhirendra Kumar, Satyabrata Bag, Varun Jaiswal, Hema Gogia, Preeti Madan, Simrita Singh, Prateek Singh, Debasis Dash, Mitali Mukerji, Manju |
| hCoV-19/India/NCDC-02310/2020 | EPI_ISL_435085 | 3/29/2020 | National Centre for Disease control (NCDC), CSIR-Institute of Genomics and Integrative Biology (CSIR-IGIB) | NCDC/CSIR-IGIB | Pramod Kumar, Rajesh Pandey, Pooja Sharma, Mahesh Dhar, Vivekanand A, Bharathram Uppili, Himanshu Vashisht, Saruchi Wadhwa, Nishu Tyagi, Uma Sharma, Priyanka Singh, Hemlata Lall, Meena Datta, Poonam Gupta, Nidhi Saini, Aarti Tewari, Bibhash Nandi, Dhirendra Kumar, Satyabrata Bag, Varun Jaiswal, Hema Gogia, Preeti Madan, Simrita Singh, Prateek Singh, Debasis Dash, Mitali Mukerji, Manju |
| hCoV-19/India/NCDC-02315/2020 | EPI_ISL_435086 | 3/29/2020 | National Centre for Disease control (NCDC), CSIR-Institute of Genomics and Integrative Biology (CSIR-IGIB) | NCDC/CSIR-IGIB | Pramod Kumar, Rajesh Pandey, Pooja Sharma, Mahesh Dhar, Vivekanand A, Bharathram Uppili, Himanshu Vashisht, Saruchi Wadhwa, Nishu Tyagi, Uma Sharma, Priyanka Singh, Hemlata Lall, Meena Datta, Poonam Gupta, Nidhi Saini, Aarti Tewari, Bibhash Nandi, Dhirendra Kumar, Satyabrata Bag, Varun Jaiswal, Hema Gogia, Preeti Madan, Simrita Singh, Prateek Singh, Debasis Dash, Mitali Mukerji, Manju |
| hCoV-19/India/NCDC-02326/2020 | EPI_ISL_435090 | 3/29/2020 | National Centre for Disease control (NCDC), CSIR-Institute of Genomics and Integrative Biology (CSIR-IGIB) | NCDC/CSIR-IGIB | Pramod Kumar, Rajesh Pandey, Pooja Sharma, Mahesh Dhar, Vivekanand A, Bharathram Uppili, Himanshu Vashisht, Saruchi Wadhwa, Nishu Tyagi, Uma Sharma, Priyanka Singh, Hemlata Lall, Meena Datta, Poonam Gupta, Nidhi Saini, Aarti Tewari, Bibhash Nandi, Dhirendra Kumar, Satyabrata Bag, Varun Jaiswal, Hema Gogia, Preeti Madan, Simrita Singh, Prateek Singh, Debasis Dash, Mitali Mukerji, Manju |
| hCoV-19/India/NCDC-02245/2020 | EPI_ISL_435091 | 3/28/2020 | National Centre for Disease control (NCDC), CSIR-Institute of Genomics and Integrative Biology (CSIR-IGIB) | NCDC/CSIR-IGIB | Pramod Kumar, Rajesh Pandey, Pooja Sharma, Mahesh Dhar, Vivekanand A, Bharathram Uppili, Himanshu Vashisht, Saruchi Wadhwa, Nishu Tyagi, Uma Sharma, Priyanka Singh, Hemlata Lall, Meena Datta, Poonam Gupta, Nidhi Saini, Aarti Tewari, Bibhash Nandi, Dhirendra Kumar, Satyabrata Bag, Varun Jaiswal, Hema Gogia, Preeti Madan, Simrita Singh, Prateek Singh, Debasis Dash, Mitali Mukerji, Manju |
| hCoV-19/India/NCDC-02244/2020 | EPI_ISL_435092 | 3/28/2020 | National Centre for Disease control (NCDC), CSIR-Institute of Genomics and Integrative Biology (CSIR-IGIB) | NCDC/CSIR-IGIB | Pramod Kumar, Rajesh Pandey, Pooja Sharma, Mahesh Dhar, Vivekanand A, Bharathram Uppili, Himanshu Vashisht, Saruchi Wadhwa, Nishu Tyagi, Uma Sharma, Priyanka Singh, Hemlata Lall, Meena Datta, Poonam Gupta, Nidhi Saini, Aarti Tewari, Bibhash Nandi, Dhirendra Kumar, Satyabrata Bag, Varun Jaiswal, Hema Gogia, Preeti Madan, Simrita Singh, Prateek Singh, Debasis Dash, Mitali Mukerji, Manju |
| hCoV-19/India/NCDC-02312/2020 | EPI_ISL_435093 | 3/29/2020 | National Centre for Disease control (NCDC), CSIR-Institute of Genomics and Integrative Biology (CSIR-IGIB) | NCDC/CSIR-IGIB | Pramod Kumar, Rajesh Pandey, Pooja Sharma, Mahesh Dhar, Vivekanand A, Bharathram Uppili, Himanshu Vashisht, Saruchi Wadhwa, Nishu Tyagi, Uma Sharma, Priyanka Singh, Hemlata Lall, Meena Datta, Poonam Gupta, Nidhi Saini, Aarti Tewari, Bibhash Nandi, Dhirendra Kumar, Satyabrata Bag, Varun Jaiswal, Hema Gogia, Preeti Madan, Simrita Singh, Prateek Singh, Debasis Dash, Mitali Mukerji, Manju |
| hCoV-19/India/NCDC-02327/2020 | EPI_ISL_435094 | 3/29/2020 | National Centre for Disease control (NCDC), CSIR-Institute of Genomics and Integrative Biology (CSIR-IGIB) | NCDC/CSIR-IGIB | Pramod Kumar, Rajesh Pandey, Pooja Sharma, Mahesh Dhar, Vivekanand A, Bharathram Uppili, Himanshu Vashisht, Saruchi Wadhwa, Nishu Tyagi, Uma Sharma, Priyanka Singh, Hemlata Lall, Meena Datta, Poonam Gupta, Nidhi Saini, Aarti Tewari, Bibhash Nandi, Dhirendra Kumar, Satyabrata Bag, Varun Jaiswal, Hema Gogia, Preeti Madan, Simrita Singh, Prateek Singh, Debasis Dash, Mitali Mukerji, Manju |
| hCoV-19/India/NCDC-02328/2020 | EPI_ISL_435095 | 3/29/2020 | National Centre for Disease control (NCDC), CSIR-Institute of Genomics and Integrative Biology (CSIR-IGIB) | NCDC/CSIR-IGIB | Pramod Kumar, Rajesh Pandey, Pooja Sharma, Mahesh Dhar, Vivekanand A, Bharathram Uppili, Himanshu Vashisht, Saruchi Wadhwa, Nishu Tyagi, Uma Sharma, Priyanka Singh, Hemlata Lall, Meena Datta, Poonam Gupta, Nidhi Saini, Aarti Tewari, Bibhash Nandi, Dhirendra Kumar, Satyabrata Bag, Varun Jaiswal, Hema Gogia, Preeti Madan, Simrita Singh, Prateek Singh, Debasis Dash, Mitali Mukerji, Manju |
| hCoV-19/India/NCDC-02332/2020 | EPI_ISL_435096 | 3/29/2020 | National Centre for Disease control (NCDC), CSIR-Institute of Genomics and Integrative Biology (CSIR-IGIB) | NCDC/CSIR-IGIB | Pramod Kumar, Rajesh Pandey, Pooja Sharma, Mahesh Dhar, Vivekanand A, Bharathram Uppili, Himanshu Vashisht, Saruchi Wadhwa, Nishu Tyagi, Uma Sharma, Priyanka Singh, Hemlata Lall, Meena Datta, Poonam Gupta, Nidhi Saini, Aarti Tewari, Bibhash Nandi, Dhirendra Kumar, Satyabrata Bag, Varun Jaiswal, Hema Gogia, Preeti Madan, Simrita Singh, Prateek Singh, Debasis Dash, Mitali Mukerji, Manju |
| hCoV-19/India/NCDC-02252/2020 | EPI_ISL_435097 | 3/28/2020 | National Centre for Disease control (NCDC), CSIR-Institute of Genomics and Integrative Biology (CSIR-IGIB) | NCDC/CSIR-IGIB | Pramod Kumar, Rajesh Pandey, Pooja Sharma, Mahesh Dhar, Vivekanand A, Bharathram Uppili, Himanshu Vashisht, Saruchi Wadhwa, Nishu Tyagi, Uma Sharma, Priyanka Singh, Hemlata Lall, Meena Datta, Poonam Gupta, Nidhi Saini, Aarti Tewari, Bibhash Nandi, Dhirendra Kumar, Satyabrata Bag, Varun Jaiswal, Hema Gogia, Preeti Madan, Simrita Singh, Prateek Singh, Debasis Dash, Mitali Mukerji, Manju |
| hCoV-19/India/NCDC-02240/2020 | EPI_ISL_435098 | 3/28/2020 | National Centre for Disease control (NCDC), CSIR-Institute of Genomics and Integrative Biology (CSIR-IGIB) | NCDC/CSIR-IGIB | Pramod Kumar, Rajesh Pandey, Pooja Sharma, Mahesh Dhar, Vivekanand A, Bharathram Uppili, Himanshu Vashisht, Saruchi Wadhwa, Nishu Tyagi, Uma Sharma, Priyanka Singh, Hemlata Lall, Meena Datta, Poonam Gupta, Nidhi Saini, Aarti Tewari, Bibhash Nandi, Dhirendra Kumar, Satyabrata Bag, Varun Jaiswal, Hema Gogia, Preeti Madan, Simrita Singh, Prateek Singh, Debasis Dash, Mitali Mukerji, Manju |
| hCoV-19/India/NCDC-02320/2020 | EPI_ISL_435099 | 3/29/2020 | National Centre for Disease control (NCDC), CSIR-Institute of Genomics and Integrative Biology (CSIR-IGIB) | NCDC/CSIR-IGIB | Pramod Kumar, Rajesh Pandey, Pooja Sharma, Mahesh Dhar, Vivekanand A, Bharathram Uppili, Himanshu Vashisht, Saruchi Wadhwa, Nishu Tyagi, Uma Sharma, Priyanka Singh, Hemlata Lall, Meena Datta, Poonam Gupta, Nidhi Saini, Aarti Tewari, Bibhash Nandi, Dhirendra Kumar, Satyabrata Bag, Varun Jaiswal, Hema Gogia, Preeti Madan, Simrita Singh, Prateek Singh, Debasis Dash, Mitali Mukerji, Manju |
| hCoV-19/India/NCDC-02321/2020 | EPI_ISL_435100 | 3/29/2020 | National Centre for Disease control (NCDC), CSIR-Institute of Genomics and Integrative Biology (CSIR-IGIB) | NCDC/CSIR-IGIB | Pramod Kumar, Rajesh Pandey, Pooja Sharma, Mahesh Dhar, Vivekanand A, Bharathram Uppili, Himanshu Vashisht, Saruchi Wadhwa, Nishu Tyagi, Uma Sharma, Priyanka Singh, Hemlata Lall, Meena Datta, Poonam Gupta, Nidhi Saini, Aarti Tewari, Bibhash Nandi, Dhirendra Kumar, Satyabrata Bag, Varun Jaiswal, Hema Gogia, Preeti Madan, Simrita Singh, Prateek Singh, Debasis Dash, Mitali Mukerji, Manju |
| hCoV-19/India/NCDC-01441/2020 | EPI_ISL_435101 | 3/15/2020 | National Centre for Disease control (NCDC), CSIR-Institute of Genomics and Integrative Biology (CSIR-IGIB) | NCDC/CSIR-IGIB | Pramod Kumar, Rajesh Pandey, Pooja Sharma, Mahesh Dhar, Vivekanand A, Bharathram Uppili, Himanshu Vashisht, Saruchi Wadhwa, Nishu Tyagi, Uma Sharma, Priyanka Singh, Hemlata Lall, Meena Datta, Poonam Gupta, Nidhi Saini, Aarti Tewari, Bibhash Nandi, Dhirendra Kumar, Satyabrata Bag, Varun Jaiswal, Hema Gogia, Preeti Madan, Simrita Singh, Prateek Singh, Debasis Dash, Mitali Mukerji, Manju |
| hCoV-19/India/NCDC-01444/2020 | EPI_ISL_435102 | 3/15/2020 | National Centre for Disease control (NCDC), CSIR-Institute of Genomics and Integrative Biology (CSIR-IGIB) | NCDC/CSIR-IGIB | Pramod Kumar, Rajesh Pandey, Pooja Sharma, Mahesh Dhar, Vivekanand A, Bharathram Uppili, Himanshu Vashisht, Saruchi Wadhwa, Nishu Tyagi, Uma Sharma, Priyanka Singh, Hemlata Lall, Meena Datta, Poonam Gupta, Nidhi Saini, Aarti Tewari, Bibhash Nandi, Dhirendra Kumar, Satyabrata Bag, Varun Jaiswal, Hema Gogia, Preeti Madan, Simrita Singh, Prateek Singh, Debasis Dash, Mitali Mukerji, Manju |
| hCoV-19/India/NCDC-01604/2020 | EPI_ISL_435103 | 3/17/2020 | National Centre for Disease control (NCDC), CSIR-Institute of Genomics and Integrative Biology (CSIR-IGIB) | NCDC/CSIR-IGIB | Pramod Kumar, Rajesh Pandey, Pooja Sharma, Mahesh Dhar, Vivekanand A, Bharathram Uppili, Himanshu Vashisht, Saruchi Wadhwa, Nishu Tyagi, Uma Sharma, Priyanka Singh, Hemlata Lall, Meena Datta, Poonam Gupta, Nidhi Saini, Aarti Tewari, Bibhash Nandi, Dhirendra Kumar, Satyabrata Bag, Varun Jaiswal, Hema Gogia, Preeti Madan, Simrita Singh, Prateek Singh, Debasis Dash, Mitali Mukerji, Manju |
| hCoV-19/India/NCDC-01614/2020 | EPI_ISL_435104 | 3/17/2020 | National Centre for Disease control (NCDC), CSIR-Institute of Genomics and Integrative Biology (CSIR-IGIB) | NCDC/CSIR-IGIB | Pramod Kumar, Rajesh Pandey, Pooja Sharma, Mahesh Dhar, Vivekanand A, Bharathram Uppili, Himanshu Vashisht, Saruchi Wadhwa, Nishu Tyagi, Uma Sharma, Priyanka Singh, Hemlata Lall, Meena Datta, Poonam Gupta, Nidhi Saini, Aarti Tewari, Bibhash Nandi, Dhirendra Kumar, Satyabrata Bag, Varun Jaiswal, Hema Gogia, Preeti Madan, Simrita Singh, Prateek Singh, Debasis Dash, Mitali Mukerji, Manju |
| hCoV-19/India/NCDC-01616/2020 | EPI_ISL_435105 | 3/17/2020 | National Centre for Disease control (NCDC), CSIR-Institute of Genomics and Integrative Biology (CSIR-IGIB) | NCDC/CSIR-IGIB | Pramod Kumar, Rajesh Pandey, Pooja Sharma, Mahesh Dhar, Vivekanand A, Bharathram Uppili, Himanshu Vashisht, Saruchi Wadhwa, Nishu Tyagi, Uma Sharma, Priyanka Singh, Hemlata Lall, Meena Datta, Poonam Gupta, Nidhi Saini, Aarti Tewari, Bibhash Nandi, Dhirendra Kumar, Satyabrata Bag, Varun Jaiswal, Hema Gogia, Preeti Madan, Simrita Singh, Prateek Singh, Debasis Dash, Mitali Mukerji, Manju |
| hCoV-19/India/NCDC-01457/2020 | EPI_ISL_435108 | 3/15/2020 | National Centre for Disease control (NCDC), CSIR-Institute of Genomics and Integrative Biology (CSIR-IGIB) | NCDC/CSIR-IGIB | Pramod Kumar, Rajesh Pandey, Pooja Sharma, Mahesh Dhar, Vivekanand A, Bharathram Uppili, Himanshu Vashisht, Saruchi Wadhwa, Nishu Tyagi, Uma Sharma, Priyanka Singh, Hemlata Lall, Meena Datta, Poonam Gupta, Nidhi Saini, Aarti Tewari, Bibhash Nandi, Dhirendra Kumar, Satyabrata Bag, Varun Jaiswal, Hema Gogia, Preeti Madan, Simrita Singh, Prateek Singh, Debasis Dash, Mitali Mukerji, Manju |
| hCoV-19/India/NCDC-01638/2020 | EPI_ISL_435109 | 3/17/2020 | National Centre for Disease control (NCDC), CSIR-Institute of Genomics and Integrative Biology (CSIR-IGIB) | NCDC/CSIR-IGIB | Pramod Kumar, Rajesh Pandey, Pooja Sharma, Mahesh Dhar, Vivekanand A, Bharathram Uppili, Himanshu Vashisht, Saruchi Wadhwa, Nishu Tyagi, Uma Sharma, Priyanka Singh, Hemlata Lall, Meena Datta, Poonam Gupta, Nidhi Saini, Aarti Tewari, Bibhash Nandi, Dhirendra Kumar, Satyabrata Bag, Varun Jaiswal, Hema Gogia, Preeti Madan, Simrita Singh, Prateek Singh, Debasis Dash, Mitali Mukerji, Manju |
| hCoV-19/India/NCDC-02415/2020 | EPI_ISL_435111 | 3/30/2020 | National Centre for Disease control (NCDC), CSIR-Institute of Genomics and Integrative Biology (CSIR-IGIB) | NCDC/CSIR-IGIB | Pramod Kumar, Rajesh Pandey, Pooja Sharma, Mahesh Dhar, Vivekanand A, Bharathram Uppili, Himanshu Vashisht, Saruchi Wadhwa, Nishu Tyagi, Uma Sharma, Priyanka Singh, Hemlata Lall, Meena Datta, Poonam Gupta, Nidhi Saini, Aarti Tewari, Bibhash Nandi, Dhirendra Kumar, Satyabrata Bag, Varun Jaiswal, Hema Gogia, Preeti Madan, Simrita Singh, Prateek Singh, Debasis Dash, Mitali Mukerji, Manju |
| hCoV-19/India/NCDC-02251/2020 | EPI_ISL_435112 | 3/28/2020 | National Centre for Disease control (NCDC), CSIR-Institute of Genomics and Integrative Biology (CSIR-IGIB) | NCDC/CSIR-IGIB | Pramod Kumar, Rajesh Pandey, Pooja Sharma, Mahesh Dhar, Vivekanand A, Bharathram Uppili, Himanshu Vashisht, Saruchi Wadhwa, Nishu Tyagi, Uma Sharma, Priyanka Singh, Hemlata Lall, Meena Datta, Poonam Gupta, Nidhi Saini, Aarti Tewari, Bibhash Nandi, Dhirendra Kumar, Satyabrata Bag, Varun Jaiswal, Hema Gogia, Preeti Madan, Simrita Singh, Prateek Singh, Debasis Dash, Mitali Mukerji, Manju |
| hCoV-19/India/NCDC-2502/2020 | EPI_ISL_436414 | 3/31/2020 | National Centre for Disease control (NCDC) | NCDC/CSIR-IGIB | Pramod Kumar#, Rajesh Pandey#, Pooja Sharma, Mahesh S Dhar, Vivekanand A, Bharathram Uppili, Himanshu Vashisht, Saruchi Wadhwa, Nishu Tyagi, Uma Sharma, Priyanka Singh, Hemlata Lall, Meena Datta, Poonam Gupta, Nidhi Saini, Aarti Tewari, Bibhash Nandi, Dhirendra Kumar, Satyabrata Bag, Varun Jaiswal, Hema Gogia, Preeti Madan, Simrita Singh, Prateek Singh, Debasis Dash, Mitali Mukerji, Ma |
| hCoV-19/India/NCDC-2507/2020 | EPI_ISL_436415 | 3/31/2020 | National Centre for Disease control (NCDC) | NCDC/CSIR-IGIB | Pramod Kumar#, Rajesh Pandey#, Pooja Sharma, Mahesh S Dhar, Vivekanand A, Bharathram Uppili, Himanshu Vashisht, Saruchi Wadhwa, Nishu Tyagi, Uma Sharma, Priyanka Singh, Hemlata Lall, Meena Datta, Poonam Gupta, Nidhi Saini, Aarti Tewari, Bibhash Nandi, Dhirendra Kumar, Satyabrata Bag, Varun Jaiswal, Hema Gogia, Preeti Madan, Simrita Singh, Prateek Singh, Debasis Dash, Mitali Mukerji, Ma |
| hCoV-19/India/NCDC-2517/2020 | EPI_ISL_436418 | 3/31/2020 | National Centre for Disease control (NCDC) | NCDC/CSIR-IGIB | Pramod Kumar#, Rajesh Pandey#, Pooja Sharma, Mahesh S Dhar, Vivekanand A, Bharathram Uppili, Himanshu Vashisht, Saruchi Wadhwa, Nishu Tyagi, Uma Sharma, Priyanka Singh, Hemlata Lall, Meena Datta, Poonam Gupta, Nidhi Saini, Aarti Tewari, Bibhash Nandi, Dhirendra Kumar, Satyabrata Bag, Varun Jaiswal, Hema Gogia, Preeti Madan, Simrita Singh, Prateek Singh, Debasis Dash, Mitali Mukerji, Ma |
| hCoV-19/India/NCDC-2522/2020 | EPI_ISL_436420 | 3/31/2020 | National Centre for Disease control (NCDC) | NCDC/CSIR-IGIB | Pramod Kumar#, Rajesh Pandey#, Pooja Sharma, Mahesh S Dhar, Vivekanand A, Bharathram Uppili, Himanshu Vashisht, Saruchi Wadhwa, Nishu Tyagi, Uma Sharma, Priyanka Singh, Hemlata Lall, Meena Datta, Poonam Gupta, Nidhi Saini, Aarti Tewari, Bibhash Nandi, Dhirendra Kumar, Satyabrata Bag, Varun Jaiswal, Hema Gogia, Preeti Madan, Simrita Singh, Prateek Singh, Debasis Dash, Mitali Mukerji, Ma |
| hCoV-19/India/NCDC-2525/2020 | EPI_ISL_436421 | 3/31/2020 | National Centre for Disease control (NCDC) | NCDC/CSIR-IGIB | Pramod Kumar#, Rajesh Pandey#, Pooja Sharma, Mahesh S Dhar, Vivekanand A, Bharathram Uppili, Himanshu Vashisht, Saruchi Wadhwa, Nishu Tyagi, Uma Sharma, Priyanka Singh, Hemlata Lall, Meena Datta, Poonam Gupta, Nidhi Saini, Aarti Tewari, Bibhash Nandi, Dhirendra Kumar, Satyabrata Bag, Varun Jaiswal, Hema Gogia, Preeti Madan, Simrita Singh, Prateek Singh, Debasis Dash, Mitali Mukerji, Ma |
| hCoV-19/India/NCDC-2535/2020 | EPI_ISL_436422 | 3/31/2020 | National Centre for Disease control (NCDC) | NCDC/CSIR-IGIB | Pramod Kumar#, Rajesh Pandey#, Pooja Sharma, Mahesh S Dhar, Vivekanand A, Bharathram Uppili, Himanshu Vashisht, Saruchi Wadhwa, Nishu Tyagi, Uma Sharma, Priyanka Singh, Hemlata Lall, Meena Datta, Poonam Gupta, Nidhi Saini, Aarti Tewari, Bibhash Nandi, Dhirendra Kumar, Satyabrata Bag, Varun Jaiswal, Hema Gogia, Preeti Madan, Simrita Singh, Prateek Singh, Debasis Dash, Mitali Mukerji, Ma |
| hCoV-19/India/NCDC-3163/2020 | EPI_ISL_436424 | 4/5/2020 | National Centre for Disease control (NCDC) | NCDC/CSIR-IGIB | Pramod Kumar#, Rajesh Pandey#, Pooja Sharma, Mahesh S Dhar, Vivekanand A, Bharathram Uppili, Himanshu Vashisht, Saruchi Wadhwa, Nishu Tyagi, Uma Sharma, Priyanka Singh, Hemlata Lall, Meena Datta, Poonam Gupta, Nidhi Saini, Aarti Tewari, Bibhash Nandi, Dhirendra Kumar, Satyabrata Bag, Varun Jaiswal, Hema Gogia, Preeti Madan, Simrita Singh, Prateek Singh, Debasis Dash, Mitali Mukerji, Ma |
| hCoV-19/India/NCDC-3170/2020 | EPI_ISL_436425 | 4/5/2020 | National Centre for Disease control (NCDC) | NCDC/CSIR-IGIB | Pramod Kumar#, Rajesh Pandey#, Pooja Sharma, Mahesh S Dhar, Vivekanand A, Bharathram Uppili, Himanshu Vashisht, Saruchi Wadhwa, Nishu Tyagi, Uma Sharma, Priyanka Singh, Hemlata Lall, Meena Datta, Poonam Gupta, Nidhi Saini, Aarti Tewari, Bibhash Nandi, Dhirendra Kumar, Satyabrata Bag, Varun Jaiswal, Hema Gogia, Preeti Madan, Simrita Singh, Prateek Singh, Debasis Dash, Mitali Mukerji, Ma |
| hCoV-19/India/NCDC-3175/2020 | EPI_ISL_436426 | 4/5/2020 | National Centre for Disease control (NCDC) | NCDC/CSIR-IGIB | Pramod Kumar#, Rajesh Pandey#, Pooja Sharma, Mahesh S Dhar, Vivekanand A, Bharathram Uppili, Himanshu Vashisht, Saruchi Wadhwa, Nishu Tyagi, Uma Sharma, Priyanka Singh, Hemlata Lall, Meena Datta, Poonam Gupta, Nidhi Saini, Aarti Tewari, Bibhash Nandi, Dhirendra Kumar, Satyabrata Bag, Varun Jaiswal, Hema Gogia, Preeti Madan, Simrita Singh, Prateek Singh, Debasis Dash, Mitali Mukerji, Ma |
| hCoV-19/India/NCDC-3181/2020 | EPI_ISL_436428 | 4/5/2020 | National Centre for Disease control (NCDC) | NCDC/CSIR-IGIB | Pramod Kumar#, Rajesh Pandey#, Pooja Sharma, Mahesh S Dhar, Vivekanand A, Bharathram Uppili, Himanshu Vashisht, Saruchi Wadhwa, Nishu Tyagi, Uma Sharma, Priyanka Singh, Hemlata Lall, Meena Datta, Poonam Gupta, Nidhi Saini, Aarti Tewari, Bibhash Nandi, Dhirendra Kumar, Satyabrata Bag, Varun Jaiswal, Hema Gogia, Preeti Madan, Simrita Singh, Prateek Singh, Debasis Dash, Mitali Mukerji, Ma |
| hCoV-19/India/NCDC-3190/2020 | EPI_ISL_436429 | 4/5/2020 | National Centre for Disease control (NCDC) | NCDC/CSIR-IGIB | Pramod Kumar#, Rajesh Pandey#, Pooja Sharma, Mahesh S Dhar, Vivekanand A, Bharathram Uppili, Himanshu Vashisht, Saruchi Wadhwa, Nishu Tyagi, Uma Sharma, Priyanka Singh, Hemlata Lall, Meena Datta, Poonam Gupta, Nidhi Saini, Aarti Tewari, Bibhash Nandi, Dhirendra Kumar, Satyabrata Bag, Varun Jaiswal, Hema Gogia, Preeti Madan, Simrita Singh, Prateek Singh, Debasis Dash, Mitali Mukerji, Ma |
| hCoV-19/India/NCDC-3195/2020 | EPI_ISL_436430 | 4/5/2020 | National Centre for Disease control (NCDC) | NCDC/CSIR-IGIB | Pramod Kumar#, Rajesh Pandey#, Pooja Sharma, Mahesh S Dhar, Vivekanand A, Bharathram Uppili, Himanshu Vashisht, Saruchi Wadhwa, Nishu Tyagi, Uma Sharma, Priyanka Singh, Hemlata Lall, Meena Datta, Poonam Gupta, Nidhi Saini, Aarti Tewari, Bibhash Nandi, Dhirendra Kumar, Satyabrata Bag, Varun Jaiswal, Hema Gogia, Preeti Madan, Simrita Singh, Prateek Singh, Debasis Dash, Mitali Mukerji, Ma |
| hCoV-19/India/NCDC-3230/2020 | EPI_ISL_436431 | 4/5/2020 | National Centre for Disease control (NCDC) | NCDC/CSIR-IGIB | Pramod Kumar#, Rajesh Pandey#, Pooja Sharma, Mahesh S Dhar, Vivekanand A, Bharathram Uppili, Himanshu Vashisht, Saruchi Wadhwa, Nishu Tyagi, Uma Sharma, Priyanka Singh, Hemlata Lall, Meena Datta, Poonam Gupta, Nidhi Saini, Aarti Tewari, Bibhash Nandi, Dhirendra Kumar, Satyabrata Bag, Varun Jaiswal, Hema Gogia, Preeti Madan, Simrita Singh, Prateek Singh, Debasis Dash, Mitali Mukerji, Ma |
| hCoV-19/India/NCDC-3271/2020 | EPI_ISL_436433 | 4/5/2020 | National Centre for Disease control (NCDC) | NCDC/CSIR-IGIB | Pramod Kumar#, Rajesh Pandey#, Pooja Sharma, Mahesh S Dhar, Vivekanand A, Bharathram Uppili, Himanshu Vashisht, Saruchi Wadhwa, Nishu Tyagi, Uma Sharma, Priyanka Singh, Hemlata Lall, Meena Datta, Poonam Gupta, Nidhi Saini, Aarti Tewari, Bibhash Nandi, Dhirendra Kumar, Satyabrata Bag, Varun Jaiswal, Hema Gogia, Preeti Madan, Simrita Singh, Prateek Singh, Debasis Dash, Mitali Mukerji, Ma |
| hCoV-19/India/NCDC-3394/2020 | EPI_ISL_436435 | 4/9/2020 | National Centre for Disease control (NCDC) | NCDC/CSIR-IGIB | Pramod Kumar#, Rajesh Pandey#, Pooja Sharma, Mahesh S Dhar, Vivekanand A, Bharathram Uppili, Himanshu Vashisht, Saruchi Wadhwa, Nishu Tyagi, Uma Sharma, Priyanka Singh, Hemlata Lall, Meena Datta, Poonam Gupta, Nidhi Saini, Aarti Tewari, Bibhash Nandi, Dhirendra Kumar, Satyabrata Bag, Varun Jaiswal, Hema Gogia, Preeti Madan, Simrita Singh, Prateek Singh, Debasis Dash, Mitali Mukerji, Ma |
| hCoV-19/India/NCDC-3936/2020 | EPI_ISL_436440 | 4/9/2020 | National Centre for Disease control (NCDC) | NCDC/CSIR-IGIB | Pramod Kumar#, Rajesh Pandey#, Pooja Sharma, Mahesh S Dhar, Vivekanand A, Bharathram Uppili, Himanshu Vashisht, Saruchi Wadhwa, Nishu Tyagi, Uma Sharma, Priyanka Singh, Hemlata Lall, Meena Datta, Poonam Gupta, Nidhi Saini, Aarti Tewari, Bibhash Nandi, Dhirendra Kumar, Satyabrata Bag, Varun Jaiswal, Hema Gogia, Preeti Madan, Simrita Singh, Prateek Singh, Debasis Dash, Mitali Mukerji, Ma |
| hCoV-19/India/NCDC-3965/2020 | EPI_ISL_436444 | 4/9/2020 | National Centre for Disease control (NCDC) | NCDC/CSIR-IGIB | Pramod Kumar#, Rajesh Pandey#, Pooja Sharma, Mahesh S Dhar, Vivekanand A, Bharathram Uppili, Himanshu Vashisht, Saruchi Wadhwa, Nishu Tyagi, Uma Sharma, Priyanka Singh, Hemlata Lall, Meena Datta, Poonam Gupta, Nidhi Saini, Aarti Tewari, Bibhash Nandi, Dhirendra Kumar, Satyabrata Bag, Varun Jaiswal, Hema Gogia, Preeti Madan, Simrita Singh, Prateek Singh, Debasis Dash, Mitali Mukerji, Ma |
| hCoV-19/India/NCDC-3982/2020 | EPI_ISL_436445 | 4/9/2020 | National Centre for Disease control (NCDC) | NCDC/CSIR-IGIB | Pramod Kumar#, Rajesh Pandey#, Pooja Sharma, Mahesh S Dhar, Vivekanand A, Bharathram Uppili, Himanshu Vashisht, Saruchi Wadhwa, Nishu Tyagi, Uma Sharma, Priyanka Singh, Hemlata Lall, Meena Datta, Poonam Gupta, Nidhi Saini, Aarti Tewari, Bibhash Nandi, Dhirendra Kumar, Satyabrata Bag, Varun Jaiswal, Hema Gogia, Preeti Madan, Simrita Singh, Prateek Singh, Debasis Dash, Mitali Mukerji, Ma |
| hCoV-19/India/NCDC-4055/2020 | EPI_ISL_436447 | 4/10/2020 | National Centre for Disease control (NCDC) | NCDC/CSIR-IGIB | Pramod Kumar#, Rajesh Pandey#, Pooja Sharma, Mahesh S Dhar, Vivekanand A, Bharathram Uppili, Himanshu Vashisht, Saruchi Wadhwa, Nishu Tyagi, Uma Sharma, Priyanka Singh, Hemlata Lall, Meena Datta, Poonam Gupta, Nidhi Saini, Aarti Tewari, Bibhash Nandi, Dhirendra Kumar, Satyabrata Bag, Varun Jaiswal, Hema Gogia, Preeti Madan, Simrita Singh, Prateek Singh, Debasis Dash, Mitali Mukerji, Ma |
| hCoV-19/India/NCDC-4138/2020 | EPI_ISL_436448 | 4/12/2020 | National Centre for Disease control (NCDC) | NCDC/CSIR-IGIB | Pramod Kumar#, Rajesh Pandey#, Pooja Sharma, Mahesh S Dhar, Vivekanand A, Bharathram Uppili, Himanshu Vashisht, Saruchi Wadhwa, Nishu Tyagi, Uma Sharma, Priyanka Singh, Hemlata Lall, Meena Datta, Poonam Gupta, Nidhi Saini, Aarti Tewari, Bibhash Nandi, Dhirendra Kumar, Satyabrata Bag, Varun Jaiswal, Hema Gogia, Preeti Madan, Simrita Singh, Prateek Singh, Debasis Dash, Mitali Mukerji, Ma |
| hCoV-19/India/NCDC-4155/2020 | EPI_ISL_436449 | 4/12/2020 | National Centre for Disease control (NCDC) | NCDC/CSIR-IGIB | Pramod Kumar#, Rajesh Pandey#, Pooja Sharma, Mahesh S Dhar, Vivekanand A, Bharathram Uppili, Himanshu Vashisht, Saruchi Wadhwa, Nishu Tyagi, Uma Sharma, Priyanka Singh, Hemlata Lall, Meena Datta, Poonam Gupta, Nidhi Saini, Aarti Tewari, Bibhash Nandi, Dhirendra Kumar, Satyabrata Bag, Varun Jaiswal, Hema Gogia, Preeti Madan, Simrita Singh, Prateek Singh, Debasis Dash, Mitali Mukerji, Ma |
| hCoV-19/India/NCDC-4189/2020 | EPI_ISL_436450 | 4/13/2020 | National Centre for Disease control (NCDC) | NCDC/CSIR-IGIB | Pramod Kumar#, Rajesh Pandey#, Pooja Sharma, Mahesh S Dhar, Vivekanand A, Bharathram Uppili, Himanshu Vashisht, Saruchi Wadhwa, Nishu Tyagi, Uma Sharma, Priyanka Singh, Hemlata Lall, Meena Datta, Poonam Gupta, Nidhi Saini, Aarti Tewari, Bibhash Nandi, Dhirendra Kumar, Satyabrata Bag, Varun Jaiswal, Hema Gogia, Preeti Madan, Simrita Singh, Prateek Singh, Debasis Dash, Mitali Mukerji, Ma |
| hCoV-19/India/NCDC-4205/2020 | EPI_ISL_436451 | 4/13/2020 | National Centre for Disease control (NCDC) | NCDC/CSIR-IGIB | Pramod Kumar#, Rajesh Pandey#, Pooja Sharma, Mahesh S Dhar, Vivekanand A, Bharathram Uppili, Himanshu Vashisht, Saruchi Wadhwa, Nishu Tyagi, Uma Sharma, Priyanka Singh, Hemlata Lall, Meena Datta, Poonam Gupta, Nidhi Saini, Aarti Tewari, Bibhash Nandi, Dhirendra Kumar, Satyabrata Bag, Varun Jaiswal, Hema Gogia, Preeti Madan, Simrita Singh, Prateek Singh, Debasis Dash, Mitali Mukerji, Ma |
| hCoV-19/India/NCDC-4208/2020 | EPI_ISL_436452 | 4/14/2020 | National Centre for Disease control (NCDC) | NCDC/CSIR-IGIB | Pramod Kumar#, Rajesh Pandey#, Pooja Sharma, Mahesh S Dhar, Vivekanand A, Bharathram Uppili, Himanshu Vashisht, Saruchi Wadhwa, Nishu Tyagi, Uma Sharma, Priyanka Singh, Hemlata Lall, Meena Datta, Poonam Gupta, Nidhi Saini, Aarti Tewari, Bibhash Nandi, Dhirendra Kumar, Satyabrata Bag, Varun Jaiswal, Hema Gogia, Preeti Madan, Simrita Singh, Prateek Singh, Debasis Dash, Mitali Mukerji, Ma |
| hCoV-19/India/NCDC-4475/2020 | EPI_ISL_436454 | 4/13/2020 | National Centre for Disease control (NCDC) | NCDC/CSIR-IGIB | Pramod Kumar#, Rajesh Pandey#, Pooja Sharma, Mahesh S Dhar, Vivekanand A, Bharathram Uppili, Himanshu Vashisht, Saruchi Wadhwa, Nishu Tyagi, Uma Sharma, Priyanka Singh, Hemlata Lall, Meena Datta, Poonam Gupta, Nidhi Saini, Aarti Tewari, Bibhash Nandi, Dhirendra Kumar, Satyabrata Bag, Varun Jaiswal, Hema Gogia, Preeti Madan, Simrita Singh, Prateek Singh, Debasis Dash, Mitali Mukerji, Ma |
| hCoV-19/India/NCDC-4476/2020 | EPI_ISL_436455 | 4/13/2020 | National Centre for Disease control (NCDC) | NCDC/CSIR-IGIB | Pramod Kumar#, Rajesh Pandey#, Pooja Sharma, Mahesh S Dhar, Vivekanand A, Bharathram Uppili, Himanshu Vashisht, Saruchi Wadhwa, Nishu Tyagi, Uma Sharma, Priyanka Singh, Hemlata Lall, Meena Datta, Poonam Gupta, Nidhi Saini, Aarti Tewari, Bibhash Nandi, Dhirendra Kumar, Satyabrata Bag, Varun Jaiswal, Hema Gogia, Preeti Madan, Simrita Singh, Prateek Singh, Debasis Dash, Mitali Mukerji, Ma |
| hCoV-19/India/NCDC-4846/2020 | EPI_ISL_436456 | 4/20/2020 | National Centre for Disease control (NCDC) | NCDC/CSIR-IGIB | Pramod Kumar#, Rajesh Pandey#, Pooja Sharma, Mahesh S Dhar, Vivekanand A, Bharathram Uppili, Himanshu Vashisht, Saruchi Wadhwa, Nishu Tyagi, Uma Sharma, Priyanka Singh, Hemlata Lall, Meena Datta, Poonam Gupta, Nidhi Saini, Aarti Tewari, Bibhash Nandi, Dhirendra Kumar, Satyabrata Bag, Varun Jaiswal, Hema Gogia, Preeti Madan, Simrita Singh, Prateek Singh, Debasis Dash, Mitali Mukerji, Ma |
| hCoV-19/India/NCDC-4859/2020 | EPI_ISL_436457 | 4/20/2020 | National Centre for Disease control (NCDC) | NCDC/CSIR-IGIB | Pramod Kumar#, Rajesh Pandey#, Pooja Sharma, Mahesh S Dhar, Vivekanand A, Bharathram Uppili, Himanshu Vashisht, Saruchi Wadhwa, Nishu Tyagi, Uma Sharma, Priyanka Singh, Hemlata Lall, Meena Datta, Poonam Gupta, Nidhi Saini, Aarti Tewari, Bibhash Nandi, Dhirendra Kumar, Satyabrata Bag, Varun Jaiswal, Hema Gogia, Preeti Madan, Simrita Singh, Prateek Singh, Debasis Dash, Mitali Mukerji, Ma |
| hCoV-19/India/NCDC-4867/2020 | EPI_ISL_436458 | 4/16/2020 | National Centre for Disease control (NCDC) | NCDC/CSIR-IGIB | Pramod Kumar#, Rajesh Pandey#, Pooja Sharma, Mahesh S Dhar, Vivekanand A, Bharathram Uppili, Himanshu Vashisht, Saruchi Wadhwa, Nishu Tyagi, Uma Sharma, Priyanka Singh, Hemlata Lall, Meena Datta, Poonam Gupta, Nidhi Saini, Aarti Tewari, Bibhash Nandi, Dhirendra Kumar, Satyabrata Bag, Varun Jaiswal, Hema Gogia, Preeti Madan, Simrita Singh, Prateek Singh, Debasis Dash, Mitali Mukerji, Ma |
| hCoV-19/India/NCDC-4874/2020 | EPI_ISL_436459 | 4/20/2020 | National Centre for Disease control (NCDC) | NCDC/CSIR-IGIB | Pramod Kumar#, Rajesh Pandey#, Pooja Sharma, Mahesh S Dhar, Vivekanand A, Bharathram Uppili, Himanshu Vashisht, Saruchi Wadhwa, Nishu Tyagi, Uma Sharma, Priyanka Singh, Hemlata Lall, Meena Datta, Poonam Gupta, Nidhi Saini, Aarti Tewari, Bibhash Nandi, Dhirendra Kumar, Satyabrata Bag, Varun Jaiswal, Hema Gogia, Preeti Madan, Simrita Singh, Prateek Singh, Debasis Dash, Mitali Mukerji, Ma |
| hCoV-19/India/NCDC-4876/2020 | EPI_ISL_436460 | 4/20/2020 | National Centre for Disease control (NCDC) | NCDC/CSIR-IGIB | Pramod Kumar#, Rajesh Pandey#, Pooja Sharma, Mahesh S Dhar, Vivekanand A, Bharathram Uppili, Himanshu Vashisht, Saruchi Wadhwa, Nishu Tyagi, Uma Sharma, Priyanka Singh, Hemlata Lall, Meena Datta, Poonam Gupta, Nidhi Saini, Aarti Tewari, Bibhash Nandi, Dhirendra Kumar, Satyabrata Bag, Varun Jaiswal, Hema Gogia, Preeti Madan, Simrita Singh, Prateek Singh, Debasis Dash, Mitali Mukerji, Ma |
| hCoV-19/India/NCDC-4877/2020 | EPI_ISL_436461 | 4/20/2020 | National Centre for Disease control (NCDC) | NCDC/CSIR-IGIB | Pramod Kumar#, Rajesh Pandey#, Pooja Sharma, Mahesh S Dhar, Vivekanand A, Bharathram Uppili, Himanshu Vashisht, Saruchi Wadhwa, Nishu Tyagi, Uma Sharma, Priyanka Singh, Hemlata Lall, Meena Datta, Poonam Gupta, Nidhi Saini, Aarti Tewari, Bibhash Nandi, Dhirendra Kumar, Satyabrata Bag, Varun Jaiswal, Hema Gogia, Preeti Madan, Simrita Singh, Prateek Singh, Debasis Dash, Mitali Mukerji, Ma |
| hCoV-19/India/NCDC-4878/2020 | EPI_ISL_436462 | 4/16/2020 | National Centre for Disease control (NCDC) | NCDC/CSIR-IGIB | Pramod Kumar#, Rajesh Pandey#, Pooja Sharma, Mahesh S Dhar, Vivekanand A, Bharathram Uppili, Himanshu Vashisht, Saruchi Wadhwa, Nishu Tyagi, Uma Sharma, Priyanka Singh, Hemlata Lall, Meena Datta, Poonam Gupta, Nidhi Saini, Aarti Tewari, Bibhash Nandi, Dhirendra Kumar, Satyabrata Bag, Varun Jaiswal, Hema Gogia, Preeti Madan, Simrita Singh, Prateek Singh, Debasis Dash, Mitali Mukerji, Ma |
| hCoV-19/India/GBRC10/2020 | EPI_ISL_437438 | 4/24/2020 | Department of MicroBiology, Government Medical College, Surat | Gujarat Biotechnology Research Centre | Amit Kanani, Akanksha Verma, Nitin Savaliya, Raghawendra Kumar, Dinesh Kumar, Zuber Saiyed, Dipa Kinariwala, Disha Patel, Binita Aring, Neeta Khandelwal, Geeta Vaghela, Sonia Barve, Bhavesh Modi, Kairavi Joshi, Gaurishankar Shrimali, Nidhi Sood, Pranay Shah, R D Dixit, Snehal Bagatharia, Kamlesh J Upadhyay, Ramesh Pandit, Tejas Shah, Ankit Hinsu, Pritesh Sabara, Apurvasinh Puvar, Janvi Raval, |
| hCoV-19/India/GBRC12/2020 | EPI_ISL_437440 | 4/26/2020 | Department of MicroBiology, Government Medical College, Surat | Gujarat Biotechnology Research Centre | Nitin Savaliya, Raghawendra Kumar, Dinesh Kumar, Zuber Saiyed, Dipa Kinariwala, Disha Patel, Binita Aring, Neeta Khandelwal, Geeta Vaghela, Sonia Barve, Bhavesh Modi, Kairavi Joshi, Gaurishankar Shrimali, Nidhi Sood, Pranay Shah, R D Dixit, Snehal Bagatharia, Kamlesh J Upadhyay, Ramesh Pandit, Tejas Shah, Ankit Hinsu, Pritesh Sabara, Apurvasinh Puvar, Janvi Raval, Monika Gandhi, Pinal Trivedi, M |
| hCoV-19/India/GBRC16/2020 | EPI_ISL_437444 | 4/26/2020 | Department of MicroBiology, Government Medical College, Surat | Gujarat Biotechnology Research Centre | Dipa Kinariwala, Disha Patel, Binita Aring, Neeta Khandelwal, Geeta Vaghela, Sonia Barve, Bhavesh Modi, Kairavi Joshi, Gaurishankar Shrimali, Nidhi Sood, Pranay Shah, R D Dixit, Snehal Bagatharia, Kamlesh J Upadhyay, Ramesh Pandit, Tejas Shah, Ankit Hinsu, Pritesh Sabara, Apurvasinh Puvar, Janvi Raval, Monika Gandhi, Pinal Trivedi, Maharshi Pandya, Amit Kanani, Akanksha Verma, Nitin Savaliya, Ra |
| hCoV-19/India/GBRC17a/2020 | EPI_ISL_437445 | 4/26/2020 | B.J. Medical College and Civil hospital | Gujarat Biotechnology Research Centre | Disha Patel, Binita Aring, Neeta Khandelwal, Geeta Vaghela, Sonia Barve, Bhavesh Modi, Kairavi Joshi, Gaurishankar Shrimali, Nidhi Sood, Pranay Shah, R D Dixit, Snehal Bagatharia, Kamlesh J Upadhyay, Ramesh Pandit, Tejas Shah, Ankit Hinsu, Pritesh Sabara, Apurvasinh Puvar, Janvi Raval, Monika Gandhi, Pinal Trivedi, Maharshi Pandya, Amit Kanani, Akanksha Verma, Nitin Savaliya, Raghawendra Kuma |
| hCoV-19/India/GBRC17b/2020 | EPI_ISL_437446 | 4/26/2020 | B.J. Medical College and Civil hospital | Gujarat Biotechnology Research Centre | Binita Aring, Neeta Khandelwal, Geeta Vaghela, Sonia Barve, Bhavesh Modi, Kairavi Joshi, Gaurishankar Shrimali, Nidhi Sood, Pranay Shah, R D Dixit, Snehal Bagatharia, Kamlesh J Upadhyay, Ramesh Pandit, Tejas Shah, Ankit Hinsu, Pritesh Sabara, Apurvasinh Puvar, Janvi Raval, Monika Gandhi, Pinal Trivedi, Maharshi Pandya, Amit Kanani, Akanksha Verma, Nitin Savaliya, Raghawendra Kumar, Dinesh Ku |
| hCoV-19/India/GBRC18/2020 | EPI_ISL_437447 | 4/26/2020 | B.J. Medical College and Civil hospital | Gujarat Biotechnology Research Centre | Neeta Khandelwal, Geeta Vaghela, Sonia Barve, Bhavesh Modi, Kairavi Joshi, Gaurishankar Shrimali, Nidhi Sood, Pranay Shah, R D Dixit, Snehal Bagatharia, Kamlesh J Upadhyay, Ramesh Pandit, Tejas Shah, Ankit Hinsu, Pritesh Sabara, Apurvasinh Puvar, Janvi Raval, Monika Gandhi, Pinal Trivedi, Maharshi Pandya, Amit Kanani, Akanksha Verma, Nitin Savaliya, Raghawendra Kumar, Dinesh Kumar, Zuber S |
| hCoV-19/India/GBRC19/2020 | EPI_ISL_437448 | 4/26/2020 | B.J. Medical College and Civil hospital | Gujarat Biotechnology Research Centre | Geeta Vaghela, Sonia Barve, Bhavesh Modi, Kairavi Joshi, Gaurishankar Shrimali, Nidhi Sood, Pranay Shah, R D Dixit, Snehal Bagatharia, Kamlesh J Upadhyay, Ramesh Pandit, Tejas Shah, Ankit Hinsu, Pritesh Sabara, Apurvasinh Puvar, Janvi Raval, Monika Gandhi, Pinal Trivedi, Maharshi Pandya, Amit Kanani, Akanksha Verma, Nitin Savaliya, Raghawendra Kumar, Dinesh Kumar, Zuber Saiyed, Dipa Kinariw |
| hCoV-19/India/GBRC20/2020 | EPI_ISL_437449 | 4/26/2020 | B.J. Medical College and Civil hospital | Gujarat Biotechnology Research Centre | Sonia Barve, Bhavesh Modi, Kairavi Joshi, Gaurishankar Shrimali, Nidhi Sood, Pranay Shah, R D Dixit, Snehal Bagatharia, Kamlesh J Upadhyay, Ramesh Pandit, Tejas Shah, Ankit Hinsu, Pritesh Sabara, Apurvasinh Puvar, Janvi Raval, Monika Gandhi, Pinal Trivedi, Maharshi Pandya, Amit Kanani, Akanksha Verma, Nitin Savaliya, Raghawendra Kumar, Dinesh Kumar, Zuber Saiyed, Dipa Kinariwala, Disha Patel, |
| hCoV-19/India/GBRC21/2020 | EPI_ISL_437450 | 4/26/2020 | B.J. Medical College and Civil hospital | Gujarat Biotechnology Research Centre | Bhavesh Modi, Kairavi Joshi, Gaurishankar Shrimali, Nidhi Sood, Pranay Shah, R D Dixit, Snehal Bagatharia, Kamlesh J Upadhyay, Ramesh Pandit, Tejas Shah, Ankit Hinsu, Pritesh Sabara, Apurvasinh Puvar, Janvi Raval, Monika Gandhi, Pinal Trivedi, Maharshi Pandya, Amit Kanani, Akanksha Verma, Nitin Savaliya, Raghawendra Kumar, Dinesh Kumar, Zuber Saiyed, Dipa Kinariwala, Disha Patel, Binita Aring, |
| hCoV-19/India/GBRC22/2020 | EPI_ISL_437451 | 4/26/2020 | B.J. Medical College and Civil hospital | Gujarat Biotechnology Research Centre | Kairavi Joshi, Gaurishankar Shrimali, Nidhi Sood, Pranay Shah, R D Dixit, Snehal Bagatharia, Kamlesh J Upadhyay, Ramesh Pandit, Tejas Shah, Ankit Hinsu, Pritesh Sabara, Apurvasinh Puvar, Janvi Raval, Monika Gandhi, Pinal Trivedi, Maharshi Pandya, Amit Kanani, Akanksha Verma, Nitin Savaliya, Raghawendra Kumar, Dinesh Kumar, Zuber Saiyed, Dipa Kinariwala, Disha Patel, Binita Aring, Neeta Khandelw |
| hCoV-19/India/GBRC23/2020 | EPI_ISL_437452 | 4/26/2020 | B.J. Medical College and Civil hospital | Gujarat Biotechnology Research Centre | Gaurishankar Shrimali, Nidhi Sood, Pranay Shah, R D Dixit, Snehal Bagatharia, Kamlesh J Upadhyay, Ramesh Pandit, Tejas Shah, Ankit Hinsu, Pritesh Sabara, Apurvasinh Puvar, Janvi Raval, Monika Gandhi, Pinal Trivedi, Maharshi Pandya, Amit Kanani, Akanksha Verma, Nitin Savaliya, Raghawendra Kumar, Dinesh Kumar, Zuber Saiyed, Dipa Kinariwala, Disha Patel, Binita Aring, Neeta Khandelwal, Geeta Va |
| hCoV-19/India/GBRC24a/2020 | EPI_ISL_437453 | 4/26/2020 | B.J. Medical College and Civil hospital | Gujarat Biotechnology Research Centre | Nidhi Sood, Pranay Shah, R D Dixit, Snehal Bagatharia, Kamlesh J Upadhyay, Ramesh Pandit, Tejas Shah, Ankit Hinsu, Pritesh Sabara, Apurvasinh Puvar, Janvi Raval, Monika Gandhi, Pinal Trivedi, Maharshi Pandya, Amit Kanani, Akanksha Verma, Nitin Savaliya, Raghawendra Kumar, Dinesh Kumar, Zuber Saiyed, Dipa Kinariwala, Disha Patel, Binita Aring, Neeta Khandelwal, Geeta Vaghela, Sonia Barve, Bha |
| hCoV-19/India/GBRC24b/2020 | EPI_ISL_437454 | 4/26/2020 | B.J. Medical College and Civil hospital | Gujarat Biotechnology Research Centre | Pranay Shah, R D Dixit, Snehal Bagatharia, Kamlesh J Upadhyay, Ramesh Pandit, Tejas Shah, Ankit Hinsu, Pritesh Sabara, Apurvasinh Puvar, Janvi Raval, Monika Gandhi, Pinal Trivedi, Maharshi Pandya, Amit Kanani, Akanksha Verma, Nitin Savaliya, Raghawendra Kumar, Dinesh Kumar, Zuber Saiyed, Dipa Kinariwala, Disha Patel, Binita Aring, Neeta Khandelwal, Geeta Vaghela, Sonia Barve, Bhavesh Modi, |
| hCoV-19/India/S12/2020 | EPI_ISL_437539 | 4/6/2020 | ICMR-National Institute of Cholera and Enteric Diseases | National Institute of Biomedical Genomics | Arindam Maitra, Mamta Chawla Sarkar, Sreedhar Chinnaswamy, Hasina Banu, Ananya Chatterjee, Shanta Dutta, Saumitra Das |
| hCoV-19/India/GMC-KP1125/2020 | EPI_ISL_437626 | 3/24/2020 | Department of Microbiology,Gandhi Medical College and Hospital | Department of Veterinary Biotechnalogy, College of Veterinary Science, Rajendranagar, PV Narsimha Rao Telengana Veterinary University | Kalyani Putty, Muttineni Radhakrishna, Nagamani K, Thrilok Chander B, Raja Rao M, Ravikumar P, Sunitha P, Pankaj Singh D, Anand Kumar K, Amit A. Upadhyay, Steven Bosinger, Rama Amara |
| hCoV-19/India/GMC-RR1191/2020 | EPI_ISL_438138 | 3/25/2020 | Department of Microbiology,Gandhi Medical College and Hospital | Department of Microbiology, Gandhi Medical College and Hospital<br>Secendrabad, Hyderbad, India | Raja Rao Mesipogu, Muttineni Radhakrishna, Nagamani K, Thrilok Chander B, Kalyani Putty, Ravikumar P, Sunitha P, Pankaj Singh D, Anand Kumar K, Amit A. Upadhyay, Steven Bosinger, Rama Amara |
| hCoV-19/India/GMC-RK1090/2020 | EPI_ISL_438139 | 3/25/2020 | Department of Microbiology,Gandhi Medical College and Hospital,Hyderabad | Virus Research Laboratory, Department of Zoology, Osmania University, Hyderabad, India | Muttineni Radhakrishna, Nagamani K, Thrilok Chander B, Raja Rao M, Kalyani Putty, Ravikumar P, Sunitha P, Pankaj Singh D, Anand Kumar K, Amit A. Upadhyay, Steven Bosinger, Rama Amara |

| | | | | | |
|---|---|---|---|---|---|
| hCoV-19/India/GBRC25/2020 | EPI_ISL_444456 | 4/26/2020 | B.J. Medical College and Civil hospital | Gujarat Biotechnology Research Centre | R D Dixit, Snehal Bagatharia, Kamlesh J Upadhyay, Ramesh Pandit, Tejas Shah, Ankit Hinsu, Pritesh Sabara, Apurvasinh Puvar, Janvi Raval, Monika Gandhi, Pinal Trivedi, Maharshi Pandya, Amit Kanani, Akanksha Verma, Nitin Savaliya, Raghawendra Kumar, Dinesh Kumar, Zuber Saiyed, Dipa Kinariwala, Disha Patel, Binita Aring, Neeta Khandelwal, Geeta Vaghela, Sonia Barve, Bhavesh Modi, Kairavi Joshi, G |
| hCoV-19/India/GBRC26/2020 | EPI_ISL_444457 | 4/26/2020 | B.J. Medical College and Civil hospital | Gujarat Biotechnology Research Centre | Snehal Bagatharia, Kamlesh J Upadhyay, Ramesh Pandit, Tejas Shah, Ankit Hinsu, Pritesh Sabara, Apurvasinh Puvar, Janvi Raval, Monika Gandhi, Pinal Trivedi, Maharshi Pandya, Amit Kanani, Akanksha Verma, Nitin Savaliya, Raghawendra Kumar, Dinesh Kumar, Zuber Saiyed, Dipa Kinariwala, Disha Patel, Binita Aring, Neeta Khandelwal, Geeta Vaghela, Sonia Barve, Bhavesh Modi, Kairavi Joshi, Gaurishan |
| hCoV-19/India/GBRC27/2020 | EPI_ISL_444458 | 4/26/2020 | B.J. Medical College and Civil hospital | Gujarat Biotechnology Research Centre | Kamlesh J Upadhyay, Ramesh Pandit, Tejas Shah, Ankit Hinsu, Pritesh Sabara, Apurvasinh Puvar, Janvi Raval, Monika Gandhi, Pinal Trivedi, Maharshi Pandya, Amit Kanani, Akanksha Verma, Nitin Savaliya, Raghawendra Kumar, Dinesh Kumar, Zuber Saiyed, Dipa Kinariwala, Disha Patel, Binita Aring, Neeta Khandelwal, Geeta Vaghela, Sonia Barve, Bhavesh Modi, Kairavi Joshi, Gaurishankar Shrimali, Nidhi S |
| hCoV-19/India/GBRC28/2020 | EPI_ISL_444459 | 4/30/2020 | B.J. Medical College and Civil hospital | Gujarat Biotechnology Research Centre | Ramesh Pandit, Tejas Shah, Ankit Hinsu, Pritesh Sabara, Apurvasinh Puvar, Janvi Raval, Monika Gandhi, Pinal Trivedi, Maharshi Pandya, Amit Kanani, Akanksha Verma, Nitin Savaliya, Raghawendra Kumar, Dinesh Kumar, Zuber Saiyed, Dipa Kinariwala, Disha Patel, Binita Aring, Neeta Khandelwal, Geeta Vaghela, Sonia Barve, Bhavesh Modi, Kairavi Joshi, Gaurishankar Shrimali, Nidhi Sood, Pranay Shah, R D |
| hCoV-19/India/GBRC29/2020 | EPI_ISL_444460 | 4/30/2020 | B.J. Medical College and Civil hospital | Gujarat Biotechnology Research Centre | Tejas Shah, Ankit Hinsu, Pritesh Sabara, Apurvasinh Puvar, Janvi Raval, Monika Gandhi, Pinal Trivedi, Maharshi Pandya, Amit Kanani, Akanksha Verma, Nitin Savaliya, Raghawendra Kumar, Dinesh Kumar, Zuber Saiyed, Dipa Kinariwala, Disha Patel, Binita Aring, Neeta Khandelwal, Geeta Vaghela, Sonia Barve, Bhavesh Modi, Kairavi Joshi, Gaurishankar Shrimali, Nidhi Sood, Pranay Shah, R D Dixit, Snehal Ba |
| hCoV-19/India/GBRC30/2020 | EPI_ISL_444461 | 4/29/2020 | B.J. Medical College and Civil hospital | Gujarat Biotechnology Research Centre | Ankit Hinsu, Pritesh Sabara, Apurvasinh Puvar, Janvi Raval, Monika Gandhi, Pinal Trivedi, Maharshi Pandya, Amit Kanani, Akanksha Verma, Nitin Savaliya, Raghawendra Kumar, Dinesh Kumar, Zuber Saiyed, Dipa Kinariwala, Disha Patel, Binita Aring, Neeta Khandelwal, Geeta Vaghela, Sonia Barve, Bhavesh Modi, Kairavi Joshi, Gaurishankar Shrimali, Nidhi Sood, Pranay Shah, R D Dixit, Snehal Bagatharia, Ka |
| hCoV-19/India/GBRC31/2020 | EPI_ISL_444462 | 4/29/2020 | B.J. Medical College and Civil hospital | Gujarat Biotechnology Research Centre | Pritesh Sabara, Apurvasinh Puvar, Janvi Raval, Monika Gandhi, Pinal Trivedi, Maharshi Pandya, Amit Kanani, Akanksha Verma, Nitin Savaliya, Raghawendra Kumar, Dinesh Kumar, Zuber Saiyed, Dipa Kinariwala, Disha Patel, Binita Aring, Neeta Khandelwal, Geeta Vaghela, Sonia Barve, Bhavesh Modi, Kairavi Joshi, Gaurishankar Shrimali, Nidhi Sood, Pranay Shah, R D Dixit, Snehal Bagatharia, Kamlesh J Upa |
| hCoV-19/India/GBRC32/2020 | EPI_ISL_444463 | 4/29/2020 | B.J. Medical College and Civil hospital | Gujarat Biotechnology Research Centre | Apurvasinh Puvar, Janvi Raval, Monika Gandhi, Pinal Trivedi, Maharshi Pandya, Amit Kanani, Akanksha Verma, Nitin Savaliya, Raghawendra Kumar, Dinesh Kumar, Zuber Saiyed, Dipa Kinariwala, Disha Patel, Binita Aring, Neeta Khandelwal, Geeta Vaghela, Sonia Barve, Bhavesh Modi, Kairavi Joshi, Gaurishankar Shrimali, Nidhi Sood, Pranay Shah, R D Dixit, Snehal Bagatharia, Kamlesh J Upadhyay, Ramesh |
| hCoV-19/India/GBRC33/2020 | EPI_ISL_444464 | 4/29/2020 | B.J. Medical College and Civil hospital | Gujarat Biotechnology Research Centre | Janvi Raval, Monika Gandhi, Pinal Trivedi, Maharshi Pandya, Amit Kanani, Akanksha Verma, Nitin Savaliya, Raghawendra Kumar, Dinesh Kumar, Zuber Saiyed, Dipa Kinariwala, Disha Patel, Binita Aring, Neeta Khandelwal, Geeta Vaghela, Sonia Barve, Bhavesh Modi, Kairavi Joshi, Gaurishankar Shrimali, Nidhi Sood, Pranay Shah, R D Dixit, Snehal Bagatharia, Kamlesh J Upadhyay, Ramesh Pandit, Tejas Shah |
| hCoV-19/India/GBRC34/2020 | EPI_ISL_444465 | 4/29/2020 | B.J. Medical College and Civil hospital | Gujarat Biotechnology Research Centre | Monika Gandhi, Pinal Trivedi, Maharshi Pandya, Amit Kanani, Akanksha Verma, Nitin Savaliya, Raghawendra Kumar, Dinesh Kumar, Zuber Saiyed, Dipa Kinariwala, Disha Patel, Binita Aring, Neeta Khandelwal, Geeta Vaghela, Sonia Barve, Bhavesh Modi, Kairavi Joshi, Gaurishankar Shrimali, Nidhi Sood, Pranay Shah, R D Dixit, Snehal Bagatharia, Kamlesh J Upadhyay, Ramesh Pandit, Tejas Shah, Ankit Hinsu |
| hCoV-19/India/GBRC35/2020 | EPI_ISL_444466 | 4/29/2020 | B.J. Medical College and Civil hospital | Gujarat Biotechnology Research Centre | Pinal Trivedi, Maharshi Pandya, Amit Kanani, Akanksha Verma, Nitin Savaliya, Raghawendra Kumar, Dinesh Kumar, Zuber Saiyed, Dipa Kinariwala, Disha Patel, Binita Aring, Neeta Khandelwal, Geeta Vaghela, Sonia Barve, Bhavesh Modi, Kairavi Joshi, Gaurishankar Shrimali, Nidhi Sood, Pranay Shah, R D Dixit, Snehal Bagatharia, Kamlesh J Upadhyay, Ramesh Pandit, Tejas Shah, Ankit Hinsu, Pritesh Sabara |
| hCoV-19/India/GBRC36/2020 | EPI_ISL_444467 | 4/29/2020 | B.J. Medical College and Civil hospital | Gujarat Biotechnology Research Centre | Maharshi Pandya, Amit Kanani, Akanksha Verma, Nitin Savaliya, Raghawendra Kumar, Dinesh Kumar, Zuber Saiyed, Dipa Kinariwala, Disha Patel, Binita Aring, Neeta Khandelwal, Geeta Vaghela, Sonia Barve, Bhavesh Modi, Kairavi Joshi, Gaurishankar Shrimali, Nidhi Sood, Pranay Shah, R D Dixit, Snehal Bagatharia, Kamlesh J Upadhyay, Ramesh Pandit, Tejas Shah, Ankit Hinsu, Pritesh Sabara, Apurvasinh P |
| hCoV-19/India/GBRC37/2020 | EPI_ISL_444468 | 4/29/2020 | B.J. Medical College and Civil hospital | Gujarat Biotechnology Research Centre | Amit Kanani, Akanksha Verma, Nitin Savaliya, Raghawendra Kumar, Dinesh Kumar, Zuber Saiyed, Dipa Kinariwala, Disha Patel, Binita Aring, Neeta Khandelwal, Geeta Vaghela, Sonia Barve, Bhavesh Modi, Kairavi Joshi, Gaurishankar Shrimali, Nidhi Sood, Pranay Shah, R D Dixit, Snehal Bagatharia, Kamlesh J Upadhyay, Ramesh Pandit, Tejas Shah, Ankit Hinsu, Pritesh Sabara, Apurvasinh Puvar, Janvi Raval, |
| hCoV-19/India/GBRC38/2020 | EPI_ISL_444469 | 4/29/2020 | B.J. Medical College and Civil hospital | Gujarat Biotechnology Research Centre | Akanksha Verma, Nitin Savaliya, Raghawendra Kumar, Dinesh Kumar, Zuber Saiyed, Dipa Kinariwala, Disha Patel, Binita Aring, Neeta Khandelwal, Geeta Vaghela, Sonia Barve, Bhavesh Modi, Kairavi Joshi, Gaurishankar Shrimali, Nidhi Sood, Pranay Shah, R D Dixit, Snehal Bagatharia, Kamlesh J Upadhyay, Ramesh Pandit, Tejas Shah, Ankit Hinsu, Pritesh Sabara, Apurvasinh Puvar, Janvi Raval, Monika Gand |
| hCoV-19/India/GBRC39/2020 | EPI_ISL_444470 | 4/29/2020 | B.J. Medical College and Civil hospital | Gujarat Biotechnology Research Centre | Nitin Savaliya, Raghawendra Kumar, Dinesh Kumar, Zuber Saiyed, Dipa Kinariwala, Disha Patel, Binita Aring, Neeta Khandelwal, Geeta Vaghela, Sonia Barve, Bhavesh Modi, Kairavi Joshi, Gaurishankar Shrimali, Nidhi Sood, Pranay Shah, R D Dixit, Snehal Bagatharia, Kamlesh J Upadhyay, Ramesh Pandit, Tejas Shah, Ankit Hinsu, Pritesh Sabara, Apurvasinh Puvar, Janvi Raval, Monika Gandhi, Pinal Trivedi, M |
| hCoV-19/India/GBRC40/2020 | EPI_ISL_444471 | 4/29/2020 | B.J. Medical College and Civil hospital | Gujarat Biotechnology Research Centre | Raghawendra Kumar, Dinesh Kumar, Zuber Saiyed, Dipa Kinariwala, Disha Patel, Binita Aring, Neeta Khandelwal, Geeta Vaghela, Sonia Barve, Bhavesh Modi, Kairavi Joshi, Gaurishankar Shrimali, Nidhi Sood, Pranay Shah, R D Dixit, Snehal Bagatharia, Kamlesh J Upadhyay, Ramesh Pandit, Tejas Shah, Ankit Hinsu, Pritesh Sabara, Apurvasinh Puvar, Janvi Raval, Monika Gandhi, Pinal Trivedi, Maharshi Pandy |
| hCoV-19/India/GBRC41/2020 | EPI_ISL_444472 | 4/29/2020 | B.J. Medical College and Civil hospital | Gujarat Biotechnology Research Centre | Dinesh Kumar, Zuber Saiyed, Dipa Kinariwala, Disha Patel, Binita Aring, Neeta Khandelwal, Geeta Vaghela, Sonia Barve, Bhavesh Modi, Kairavi Joshi, Gaurishankar Shrimali, Nidhi Sood, Pranay Shah, R D Dixit, Snehal Bagatharia, Kamlesh J Upadhyay, Ramesh Pandit, Tejas Shah, Ankit Hinsu, Pritesh Sabara, Apurvasinh Puvar, Janvi Raval, Monika Gandhi, Pinal Trivedi, Maharshi Pandya, Amit Kanani, Akank |
| hCoV-19/India/GBRC42/2020 | EPI_ISL_444473 | 4/29/2020 | B.J. Medical College and Civil hospital | Gujarat Biotechnology Research Centre | Zuber Saiyed, Dipa Kinariwala, Disha Patel, Binita Aring, Neeta Khandelwal, Geeta Vaghela, Sonia Barve, Bhavesh Modi, Kairavi Joshi, Gaurishankar Shrimali, Nidhi Sood, Pranay Shah, R D Dixit, Snehal Bagatharia, Kamlesh J Upadhyay, Ramesh Pandit, Tejas Shah, Ankit Hinsu, Pritesh Sabara, Apurvasinh Puvar, Janvi Raval, Monika Gandhi, Pinal Trivedi, Maharshi Pandya, Amit Kanani, Akanksha Verma, Niti |
| hCoV-19/India/GBRC43/2020 | EPI_ISL_444474 | 4/29/2020 | B.J. Medical College and Civil hospital | Gujarat Biotechnology Research Centre | Dipa Kinariwala, Disha Patel, Binita Aring, Neeta Khandelwal, Geeta Vaghela, Sonia Barve, Bhavesh Modi, Kairavi Joshi, Gaurishankar Shrimali, Nidhi Sood, Pranay Shah, R D Dixit, Snehal Bagatharia, Kamlesh J Upadhyay, Ramesh Pandit, Tejas Shah, Ankit Hinsu, Pritesh Sabara, Apurvasinh Puvar, Janvi Raval, Monika Gandhi, Pinal Trivedi, Maharshi Pandya, Amit Kanani, Akanksha Verma, Nitin Savaliya, Rag |
| hCoV-19/India/GBRC44/2020 | EPI_ISL_444475 | 4/29/2020 | B.J. Medical College and Civil hospital | Gujarat Biotechnology Research Centre | Disha Patel, Binita Aring, Neeta Khandelwal, Geeta Vaghela, Sonia Barve, Bhavesh Modi, Kairavi Joshi, Gaurishankar Shrimali, Nidhi Sood, Pranay Shah, R D Dixit, Snehal Bagatharia, Kamlesh J Upadhyay, Ramesh Pandit, Tejas Shah, Ankit Hinsu, Pritesh Sabara, Apurvasinh Puvar, Janvi Raval, Monika Gandhi, Pinal Trivedi, Maharshi Pandya, Amit Kanani, Akanksha Verma, Nitin Savaliya, Raghawendra Kuma |
| hCoV-19/India/GBRC45/2020 | EPI_ISL_444476 | 4/29/2020 | B.J. Medical College and Civil hospital | Gujarat Biotechnology Research Centre | Binita Aring, Neeta Khandelwal, Geeta Vaghela, Sonia Barve, Bhavesh Modi, Kairavi Joshi, Gaurishankar Shrimali, Nidhi Sood, Pranay Shah, R D Dixit, Snehal Bagatharia, Kamlesh J Upadhyay, Ramesh Pandit, Tejas Shah, Ankit Hinsu, Pritesh Sabara, Apurvasinh Puvar, Janvi Raval, Monika Gandhi, Pinal Trivedi, Maharshi Pandya, Amit Kanani, Akanksha Verma, Nitin Savaliya, Raghawendra Kumar, Dinesh Ku |
| hCoV-19/India/GBRC46/2020 | EPI_ISL_444477 | 4/29/2020 | B.J. Medical College and Civil hospital | Gujarat Biotechnology Research Centre | Neeta Khandelwal, Geeta Vaghela, Sonia Barve, Bhavesh Modi, Kairavi Joshi, Gaurishankar Shrimali, Nidhi Sood, Pranay Shah, R D Dixit, Snehal Bagatharia, Kamlesh J Upadhyay, Ramesh Pandit, Tejas Shah, Ankit Hinsu, Pritesh Sabara, Apurvasinh Puvar, Janvi Raval, Monika Gandhi, Pinal Trivedi, Maharshi Pandya, Amit Kanani, Akanksha Verma, Nitin Savaliya, Raghawendra Kumar, Dinesh Kumar, Zuber S |
| hCoV-19/India/GBRC47/2020 | EPI_ISL_444478 | 4/30/2020 | B.J. Medical College and Civil hospital | Gujarat Biotechnology Research Centre | Geeta Vaghela, Sonia Barve, Bhavesh Modi, Kairavi Joshi, Gaurishankar Shrimali, Nidhi Sood, Pranay Shah, R D Dixit, Snehal Bagatharia, Kamlesh J Upadhyay, Ramesh Pandit, Tejas Shah, Ankit Hinsu, Pritesh Sabara, Apurvasinh Puvar, Janvi Raval, Monika Gandhi, Pinal Trivedi, Maharshi Pandya, Amit Kanani, Akanksha Verma, Nitin Savaliya, Raghawendra Kumar, Dinesh Kumar, Zuber Saiyed, Dipa Kinariw |
| hCoV-19/India/GBRC48/2020 | EPI_ISL_444479 | 4/29/2020 | B.J. Medical College and Civil hospital | Gujarat Biotechnology Research Centre | Sonia Barve, Bhavesh Modi, Kairavi Joshi, Gaurishankar Shrimali, Nidhi Sood, Pranay Shah, R D Dixit, Snehal Bagatharia, Kamlesh J Upadhyay, Ramesh Pandit, Tejas Shah, Ankit Hinsu, Pritesh Sabara, Apurvasinh Puvar, Janvi Raval, Monika Gandhi, Pinal Trivedi, Maharshi Pandya, Amit Kanani, Akanksha Verma, Nitin Savaliya, Raghawendra Kumar, Dinesh Kumar, Zuber Saiyed, Dipa Kinariwala, Disha Patel, |
| hCoV-19/India/GBRC49/2020 | EPI_ISL_444480 | 4/30/2020 | B.J. Medical College and Civil hospital | Gujarat Biotechnology Research Centre | Bhavesh Modi, Kairavi Joshi, Gaurishankar Shrimali, Nidhi Sood, Pranay Shah, R D Dixit, Snehal Bagatharia, Kamlesh J Upadhyay, Ramesh Pandit, Tejas Shah, Ankit Hinsu, Pritesh Sabara, Apurvasinh Puvar, Janvi Raval, Monika Gandhi, Pinal Trivedi, Maharshi Pandya, Amit Kanani, Akanksha Verma, Nitin Savaliya, Raghawendra Kumar, Dinesh Kumar, Zuber Saiyed, Dipa Kinariwala, Disha Patel, Binita Aring, |
| hCoV-19/India/GBRC50/2020 | EPI_ISL_444481 | 4/30/2020 | B.J. Medical College and Civil hospital | Gujarat Biotechnology Research Centre | Kairavi Joshi, Gaurishankar Shrimali, Nidhi Sood, Pranay Shah, R D Dixit, Snehal Bagatharia, Kamlesh J Upadhyay, Ramesh Pandit, Tejas Shah, Ankit Hinsu, Pritesh Sabara, Apurvasinh Puvar, Janvi Raval, Monika Gandhi, Pinal Trivedi, Maharshi Pandya, Amit Kanani, Akanksha Verma, Nitin Savaliya, Raghawendra Kumar, Dinesh Kumar, Zuber Saiyed, Dipa Kinariwala, Disha Patel, Binita Aring, Neeta Khandel |
| hCoV-19/India/GBRC51/2020 | EPI_ISL_444482 | 5/2/2020 | Gujarat Biotechnology Research Centre | Gujarat Biotechnology Research Centre | Gaurishankar Shrimali, Nidhi Sood, Pranay Shah, R D Dixit, Snehal Bagatharia, Kamlesh J Upadhyay, Ramesh Pandit, Tejas Shah, Ankit Hinsu, Pritesh Sabara, Apurvasinh Puvar, Janvi Raval, Monika Gandhi, Pinal Trivedi, Maharshi Pandya, Amit Kanani, Akanksha Verma, Nitin Savaliya, Raghawendra Kumar, Dinesh Kumar, Zuber Saiyed, Dipa Kinariwala, Disha Patel, Binita Aring, Neeta Khandelwal, Geeta Va |
| hCoV-19/India/GBRC52/2020 | EPI_ISL_444483 | 5/2/2020 | Gujarat Biotechnology Research Centre | Gujarat Biotechnology Research Centre | Nidhi Sood, Pranay Shah, R D Dixit, Snehal Bagatharia, Kamlesh J Upadhyay, Ramesh Pandit, Tejas Shah, Ankit Hinsu, Pritesh Sabara, Apurvasinh Puvar, Janvi Raval, Monika Gandhi, Pinal Trivedi, Maharshi Pandya, Amit Kanani, Akanksha Verma, Nitin Savaliya, Raghawendra Kumar, Dinesh Kumar, Zuber Saiyed, Dipa Kinariwala, Disha Patel, Binita Aring, Neeta Khandelwal, Geeta Vaghela, Sonia Barve, Bha |
| hCoV-19/India/GBRC53/2020 | EPI_ISL_444484 | 5/4/2020 | Gujarat Biotechnology Research Centre | Gujarat Biotechnology Research Centre | Pranay Shah, R D Dixit, Snehal Bagatharia, Kamlesh J Upadhyay, Ramesh Pandit, Tejas Shah, Ankit Hinsu, Pritesh Sabara, Apurvasinh Puvar, Janvi Raval, Monika Gandhi, Pinal Trivedi, Maharshi Pandya, Amit Kanani, Akanksha Verma, Nitin Savaliya, Raghawendra Kumar, Dinesh Kumar, Zuber Saiyed, Dipa Kinariwala, Disha Patel, Binita Aring, Neeta Khandelwal, Geeta Vaghela, Sonia Barve, Bhavesh Modi, K |
| hCoV-19/India/GBRC54/2020 | EPI_ISL_444485 | 5/4/2020 | Gujarat Biotechnology Research Centre | Gujarat Biotechnology Research Centre | R D Dixit, Snehal Bagatharia, Kamlesh J Upadhyay, Ramesh Pandit, Tejas Shah, Ankit Hinsu, Pritesh Sabara, Apurvasinh Puvar, Janvi Raval, Monika Gandhi, Pinal Trivedi, Maharshi Pandya, Amit Kanani, Akanksha Verma, Nitin Savaliya, Raghawendra Kumar, Dinesh Kumar, Zuber Saiyed, Dipa Kinariwala, Disha Patel, Binita Aring, Neeta Khandelwal, Geeta Vaghela, Sonia Barve, Bhavesh Modi, Kairavi Joshi, G |
| hCoV-19/India/GBRC55/2020 | EPI_ISL_444486 | 5/4/2020 | Gujarat Biotechnology Research Centre | Gujarat Biotechnology Research Centre | Snehal Bagatharia, Kamlesh J Upadhyay, Ramesh Pandit, Tejas Shah, Ankit Hinsu, Pritesh Sabara, Apurvasinh Puvar, Janvi Raval, Monika Gandhi, Pinal Trivedi, Maharshi Pandya, Amit Kanani, Akanksha Verma, Nitin Savaliya, Raghawendra Kumar, Dinesh Kumar, Zuber Saiyed, Dipa Kinariwala, Disha Patel, Binita Aring, Neeta Khandelwal, Geeta Vaghela, Sonia Barve, Bhavesh Modi, Kairavi Joshi, Gaurishan |
| hCoV-19/India/GBRC56a/2020 | EPI_ISL_447030 | 5/3/2020 | B.J. Medical College and Civil hospital | Gujarat Biotechnology Research Centre | Kamlesh J Upadhyay, Ramesh Pandit, Tejas Shah, Ankit Hinsu, Pritesh Sabara, Apurvasinh Puvar, Janvi Raval, Monika Gandhi, Pinal Trivedi, Maharshi Pandya, Amit Kanani, Akanksha Verma, Nitin Savaliya, Raghawendra Kumar, Dinesh Kumar, Zuber Saiyed, Dipa Kinariwala, Disha Patel, Binita Aring, Neeta Khandelwal, Geeta Vaghela, Sonia Barve, Bhavesh Modi, Kairavi Joshi, Gaurishankar Shrimali, Nidhi S |
| hCoV-19/India/GBRC56b/2020 | EPI_ISL_447031 | 5/3/2020 | B.J. Medical College and Civil hospital | Gujarat Biotechnology Research Centre | Ramesh Pandit, Tejas Shah, Ankit Hinsu, Pritesh Sabara, Apurvasinh Puvar, Janvi Raval, Monika Gandhi, Pinal Trivedi, Maharshi Pandya, Amit Kanani, Akanksha Verma, Nitin Savaliya, Raghawendra Kumar, Dinesh Kumar, Zuber Saiyed, Dipa Kinariwala, Disha Patel, Binita Aring, Neeta Khandelwal, Geeta Vaghela, Sonia Barve, Bhavesh Modi, Kairavi Joshi, Gaurishankar Shrimali, Nidhi Sood, Pranay Shah, R D |
| hCoV-19/India/GBRC58/2020 | EPI_ISL_447033 | 5/3/2020 | B.J. Medical College and Civil hospital | Gujarat Biotechnology Research Centre | Ankit Hinsu, Pritesh Sabara, Apurvasinh Puvar, Janvi Raval, Monika Gandhi, Pinal Trivedi, Maharshi Pandya, Amit Kanani, Akanksha Verma, Nitin Savaliya, Raghawendra Kumar, Dinesh Kumar, Zuber Saiyed, Dipa Kinariwala, Disha Patel, Binita Aring, Neeta Khandelwal, Geeta Vaghela, Sonia Barve, Bhavesh Modi, Kairavi Joshi, Gaurishankar Shrimali, Nidhi Sood, Pranay Shah, R D Dixit, Snehal Bagatharia, Ka |
| hCoV-19/India/GBRC60/2020 | EPI_ISL_447035 | 5/3/2020 | B.J. Medical College and Civil hospital | Gujarat Biotechnology Research Centre | Apurvasinh Puvar, Janvi Raval, Monika Gandhi, Pinal Trivedi, Maharshi Pandya, Amit Kanani, Akanksha Verma, Nitin Savaliya, Raghawendra Kumar, Dinesh Kumar, Zuber Saiyed, Dipa Kinariwala, Disha Patel, Binita Aring, Neeta Khandelwal, Geeta Vaghela, Sonia Barve, Bhavesh Modi, Kairavi Joshi, Gaurishankar Shrimali, Nidhi Sood, Pranay Shah, R D Dixit, Snehal Bagatharia, Kamlesh J Upadhyay, Ramesh |
| hCoV-19/India/GBRC61/2020 | EPI_ISL_447036 | 5/3/2020 | B.J. Medical College and Civil hospital | Gujarat Biotechnology Research Centre | Janvi Raval, Monika Gandhi, Pinal Trivedi, Maharshi Pandya, Amit Kanani, Akanksha Verma, Nitin Savaliya, Raghawendra Kumar, Dinesh Kumar, Zuber Saiyed, Dipa Kinariwala, Disha Patel, Binita Aring, Neeta Khandelwal, Geeta Vaghela, Sonia Barve, Bhavesh Modi, Kairavi Joshi, Gaurishankar Shrimali, Nidhi Sood, Pranay Shah, R D Dixit, Snehal Bagatharia, Kamlesh J Upadhyay, Ramesh Pandit, Tejas Shah |
| hCoV-19/India/GBRC62/2020 | EPI_ISL_447037 | 5/3/2020 | B.J. Medical College and Civil hospital | Gujarat Biotechnology Research Centre | Monika Gandhi, Pinal Trivedi, Maharshi Pandya, Amit Kanani, Akanksha Verma, Nitin Savaliya, Raghawendra Kumar, Dinesh Kumar, Zuber Saiyed, Dipa Kinariwala, Disha Patel, Binita Aring, Neeta Khandelwal, Geeta Vaghela, Sonia Barve, Bhavesh Modi, Kairavi Joshi, Gaurishankar Shrimali, Nidhi Sood, Pranay Shah, R D Dixit, Snehal Bagatharia, Kamlesh J Upadhyay, Ramesh Pandit, Tejas Shah, Ankit Hinsu |
| hCoV-19/India/GBRC63/2020 | EPI_ISL_447038 | 5/3/2020 | B.J. Medical College and Civil hospital | Gujarat Biotechnology Research Centre | Pinal Trivedi, Maharshi Pandya, Amit Kanani, Akanksha Verma, Nitin Savaliya, Raghawendra Kumar, Dinesh Kumar, Zuber Saiyed, Dipa Kinariwala, Disha Patel, Binita Aring, Neeta Khandelwal, Geeta Vaghela, Sonia Barve, Bhavesh Modi, Kairavi Joshi, Gaurishankar Shrimali, Nidhi Sood, Pranay Shah, R D Dixit, Snehal Bagatharia, Kamlesh J Upadhyay, Ramesh Pandit, Tejas Shah, Ankit Hinsu, Pritesh Sabara |
| hCoV-19/India/GBRC64/2020 | EPI_ISL_447039 | 5/3/2020 | B.J. Medical College and Civil hospital | Gujarat Biotechnology Research Centre | Maharshi Pandya, Amit Kanani, Akanksha Verma, Nitin Savaliya, Raghawendra Kumar, Dinesh Kumar, Zuber Saiyed, Dipa Kinariwala, Disha Patel, Binita Aring, Neeta Khandelwal, Geeta Vaghela, Sonia Barve, Bhavesh Modi, Kairavi Joshi, Gaurishankar Shrimali, Nidhi Sood, Pranay Shah, R D Dixit, Snehal Bagatharia, Kamlesh J Upadhyay, Ramesh Pandit, Tejas Shah, Ankit Hinsu, Pritesh Sabara, Apurvasinh P |
| hCoV-19/India/GBRC65a/2020 | EPI_ISL_447040 | 5/3/2020 | B.J. Medical College and Civil hospital | Gujarat Biotechnology Research Centre | Amit Kanani, Akanksha Verma, Nitin Savaliya, Raghawendra Kumar, Dinesh Kumar, Zuber Saiyed, Dipa Kinariwala, Disha Patel, Binita Aring, Neeta Khandelwal, Geeta Vaghela, Sonia Barve, Bhavesh Modi, Kairavi Joshi, Gaurishankar Shrimali, Nidhi Sood, Pranay Shah, R D Dixit, Snehal Bagatharia, Kamlesh J Upadhyay, Ramesh Pandit, Tejas Shah, Ankit Hinsu, Pritesh Sabara, Apurvasinh Puvar, Janvi Raval, |
| hCoV-19/India/GBRC65b/2020 | EPI_ISL_447041 | 5/3/2020 | B.J. Medical College and Civil hospital | Gujarat Biotechnology Research Centre | Akanksha Verma, Nitin Savaliya, Raghawendra Kumar, Dinesh Kumar, Zuber Saiyed, Dipa Kinariwala, Disha Patel, Binita Aring, Neeta Khandelwal, Geeta Vaghela, Sonia Barve, Bhavesh Modi, Kairavi Joshi, Gaurishankar Shrimali, Nidhi Sood, Pranay Shah, R D Dixit, Snehal Bagatharia, Kamlesh J Upadhyay, Ramesh Pandit, Tejas Shah, Ankit Hinsu, Pritesh Sabara, Apurvasinh Puvar, Janvi Raval, Monika Gand |
| hCoV-19/India/GBRC66/2020 | EPI_ISL_447042 | 5/3/2020 | B.J. Medical College and Civil hospital | Gujarat Biotechnology Research Centre | Nitin Savaliya, Raghawendra Kumar, Dinesh Kumar, Zuber Saiyed, Dipa Kinariwala, Disha Patel, Binita Aring, Neeta Khandelwal, Geeta Vaghela, Sonia Barve, Bhavesh Modi, Kairavi Joshi, Gaurishankar Shrimali, Nidhi Sood, Pranay Shah, R D Dixit, Snehal Bagatharia, Kamlesh J Upadhyay, Ramesh Pandit, Tejas Shah, Ankit Hinsu, Pritesh Sabara, Apurvasinh Puvar, Janvi Raval, Monika Gandhi, Pinal Trivedi, M |
| hCoV-19/India/GBRC67a/2020 | EPI_ISL_447043 | 5/3/2020 | B.J. Medical College and Civil hospital | Gujarat Biotechnology Research Centre | Raghawendra Kumar, Dinesh Kumar, Zuber Saiyed, Dipa Kinariwala, Disha Patel, Binita Aring, Neeta Khandelwal, Geeta Vaghela, Sonia Barve, Bhavesh Modi, Kairavi Joshi, Gaurishankar Shrimali, Nidhi Sood, Pranay Shah, R D Dixit, Snehal Bagatharia, Kamlesh J Upadhyay, Ramesh Pandit, Tejas Shah, Ankit Hinsu, Pritesh Sabara, Apurvasinh Puvar, Janvi Raval, Monika Gandhi, Pinal Trivedi, Maharshi Pandy |
| hCoV-19/India/GBRC67b/2020 | EPI_ISL_447044 | 5/3/2020 | B.J. Medical College and Civil hospital | Gujarat Biotechnology Research Centre | Dinesh Kumar, Zuber Saiyed, Dipa Kinariwala, Disha Patel, Binita Aring, Neeta Khandelwal, Geeta Vaghela, Sonia Barve, Bhavesh Modi, Kairavi Joshi, Gaurishankar Shrimali, Nidhi Sood, Pranay Shah, R D Dixit, Snehal Bagatharia, Kamlesh J Upadhyay, Ramesh Pandit, Tejas Shah, Ankit Hinsu, Pritesh Sabara, Apurvasinh Puvar, Janvi Raval, Monika Gandhi, Pinal Trivedi, Maharshi Pandya, Amit Kanani, Akank |
| hCoV-19/India/GBRC68/2020 | EPI_ISL_447045 | 5/3/2020 | B.J. Medical College and Civil hospital | Gujarat Biotechnology Research Centre | Zuber Saiyed, Dipa Kinariwala, Disha Patel, Binita Aring, Neeta Khandelwal, Geeta Vaghela, Sonia Barve, Bhavesh Modi, Kairavi Joshi, Gaurishankar Shrimali, Nidhi Sood, Pranay Shah, R D Dixit, Snehal Bagatharia, Kamlesh J Upadhyay, Ramesh Pandit, Tejas Shah, Ankit Hinsu, Pritesh Sabara, Apurvasinh Puvar, Janvi Raval, Monika Gandhi, Pinal Trivedi, Maharshi Pandya, Amit Kanani, Akanksha Verma, Niti |
| hCoV-19/India/GBRC69/2020 | EPI_ISL_447046 | 5/4/2020 | B.J. Medical College and Civil hospital | Gujarat Biotechnology Research Centre | Dipa Kinariwala, Disha Patel, Binita Aring, Neeta Khandelwal, Geeta Vaghela, Sonia Barve, Bhavesh Modi, Kairavi Joshi, Gaurishankar Shrimali, Nidhi Sood, Pranay Shah, R D Dixit, Snehal Bagatharia, Kamlesh J Upadhyay, Ramesh Pandit, Tejas Shah, Ankit Hinsu, Pritesh Sabara, Apurvasinh Puvar, Janvi Raval, Monika Gandhi, Pinal Trivedi, Maharshi Pandya, Amit Kanani, Akanksha Verma, Nitin Savaliya, Rag |
| hCoV-19/India/GBRC70/2020 | EPI_ISL_447047 | 4/29/2020 | GMERS Medical College and Hospital, Gandhinagar | Gujarat Biotechnology Research Centre | Disha Patel, Binita Aring, Neeta Khandelwal, Geeta Vaghela, Sonia Barve, Bhavesh Modi, Kairavi Joshi, Gaurishankar Shrimali, Nidhi Sood, Pranay Shah, R D Dixit, Snehal Bagatharia, Kamlesh J Upadhyay, Ramesh Pandit, Tejas Shah, Ankit Hinsu, Pritesh Sabara, Apurvasinh Puvar, Janvi Raval, Monika Gandhi, Pinal Trivedi, Maharshi Pandya, Amit Kanani, Akanksha Verma, Nitin Savaliya, Raghawendra Kuma |
| hCoV-19/India/GBRC71/2020 | EPI_ISL_447048 | 4/27/2020 | GMERS Medical College and Hospital, Gandhinagar | Gujarat Biotechnology Research Centre | Binita Aring, Neeta Khandelwal, Geeta Vaghela, Sonia Barve, Bhavesh Modi, Kairavi Joshi, Gaurishankar Shrimali, Nidhi Sood, Pranay Shah, R D Dixit, Snehal Bagatharia, Kamlesh J Upadhyay, Ramesh Pandit, Tejas Shah, Ankit Hinsu, Pritesh Sabara, Apurvasinh Puvar, Janvi Raval, Monika Gandhi, Pinal Trivedi, Maharshi Pandya, Amit Kanani, Akanksha Verma, Nitin Savaliya, Raghawendra Kumar, Dinesh Ku |
| hCoV-19/India/GBRC72a/2020 | EPI_ISL_447049 | 4/29/2020 | GMERS Medical College and Hospital, Gandhinagar | Gujarat Biotechnology Research Centre | Neeta Khandelwal, Geeta Vaghela, Sonia Barve, Bhavesh Modi, Kairavi Joshi, Gaurishankar Shrimali, Nidhi Sood, Pranay Shah, R D Dixit, Snehal Bagatharia, Kamlesh J Upadhyay, Ramesh Pandit, Tejas Shah, Ankit Hinsu, Pritesh Sabara, Apurvasinh Puvar, Janvi Raval, Monika Gandhi, Pinal Trivedi, Maharshi Pandya, Amit Kanani, Akanksha Verma, Nitin Savaliya, Raghawendra Kumar, Dinesh Kumar, Zuber S |
| hCoV-19/India/GBRC72b/2020 | EPI_ISL_447050 | 4/29/2020 | GMERS Medical College and Hospital, Gandhinagar | Gujarat Biotechnology Research Centre | Geeta Vaghela, Sonia Barve, Bhavesh Modi, Kairavi Joshi, Gaurishankar Shrimali, Nidhi Sood, Pranay Shah, R D Dixit, Snehal Bagatharia, Kamlesh J Upadhyay, Ramesh Pandit, Tejas Shah, Ankit Hinsu, Pritesh Sabara, Apurvasinh Puvar, Janvi Raval, Monika Gandhi, Pinal Trivedi, Maharshi Pandya, Amit Kanani, Akanksha Verma, Nitin Savaliya, Raghawendra Kumar, Dinesh Kumar, Zuber Saiyed, Dipa Kinariw |
| hCoV-19/India/GBRC73/2020 | EPI_ISL_447051 | 4/29/2020 | GMERS Medical College and Hospital, Gandhinagar | Gujarat Biotechnology Research Centre | Sonia Barve, Bhavesh Modi, Kairavi Joshi, Gaurishankar Shrimali, Nidhi Sood, Pranay Shah, R D Dixit, Snehal Bagatharia, Kamlesh J Upadhyay, Ramesh Pandit, Tejas Shah, Ankit Hinsu, Pritesh Sabara, Apurvasinh Puvar, Janvi Raval, Monika Gandhi, Pinal Trivedi, Maharshi Pandya, Amit Kanani, Akanksha Verma, Nitin Savaliya, Raghawendra Kumar, Dinesh Kumar, Zuber Saiyed, Dipa Kinariwala, Disha Patel, |
| hCoV-19/India/GBRC74/2020 | EPI_ISL_447052 | 5/2/2020 | GMERS Medical College and Hospital, Gandhinagar | Gujarat Biotechnology Research Centre | Bhavesh Modi, Kairavi Joshi, Gaurishankar Shrimali, Nidhi Sood, Pranay Shah, R D Dixit, Snehal Bagatharia, Kamlesh J Upadhyay, Ramesh Pandit, Tejas Shah, Ankit Hinsu, Pritesh Sabara, Apurvasinh Puvar, Janvi Raval, Monika Gandhi, Pinal Trivedi, Maharshi Pandya, Amit Kanani, Akanksha Verma, Nitin Savaliya, Raghawendra Kumar, Dinesh Kumar, Zuber Saiyed, Dipa Kinariwala, Disha Patel, Binita Aring, |
| hCoV-19/India/GBRC75/2020 | EPI_ISL_447053 | 4/29/2020 | GMERS Medical College and Hospital, Gandhinagar | Gujarat Biotechnology Research Centre | Kairavi Joshi, Gaurishankar Shrimali, Nidhi Sood, Pranay Shah, R D Dixit, Snehal Bagatharia, Kamlesh J Upadhyay, Ramesh Pandit, Tejas Shah, Ankit Hinsu, Pritesh Sabara, Apurvasinh Puvar, Janvi Raval, Monika Gandhi, Pinal Trivedi, Maharshi Pandya, Amit Kanani, Akanksha Verma, Nitin Savaliya, Raghawendra Kumar, Dinesh Kumar, Zuber Saiyed, Dipa Kinariwala, Disha Patel, Binita Aring, Neeta Khandel |
| hCoV-19/India/GBRC76/2020 | EPI_ISL_447534 | 5/5/2020 | Gujarat Biotechnology Research Centre | Gujarat Biotechnology Research Centre | Gaurishankar Shrimali, Nidhi Sood, Pranay Shah, R D Dixit, Snehal Bagatharia, Kamlesh J Upadhyay, Ramesh Pandit, Tejas Shah, Ankit Hinsu, Pritesh Sabara, Apurvasinh Puvar, Janvi Raval, Monika Gandhi, Pinal Trivedi, Maharshi Pandya, Amit Kanani, Akanksha Verma, Nitin Savaliya, Raghawendra Kumar, Dinesh Kumar, Zuber Saiyed, Dipa Kinariwala, Disha Patel, Binita Aring, Neeta Khandelwal, Geeta Va |
| hCoV-19/India/GBRC77/2020 | EPI_ISL_447535 | 5/5/2020 | Gujarat Biotechnology Research Centre | Gujarat Biotechnology Research Centre | Nidhi Sood, Pranay Shah, R D Dixit, Snehal Bagatharia, Kamlesh J Upadhyay, Ramesh Pandit, Tejas Shah, Ankit Hinsu, Pritesh Sabara, Apurvasinh Puvar, Janvi Raval, Monika Gandhi, Pinal Trivedi, Maharshi Pandya, Amit Kanani, Akanksha Verma, Nitin Savaliya, Raghawendra Kumar, Dinesh Kumar, Zuber Saiyed, Dipa Kinariwala, Disha Patel, Binita Aring, Neeta Khandelwal, Geeta Vaghela, Sonia Barve, Bha |
| hCoV-19/India/GBRC78a/2020 | EPI_ISL_447536 | 5/5/2020 | Gujarat Biotechnology Research Centre | Gujarat Biotechnology Research Centre | Pranay Shah, R D Dixit, Snehal Bagatharia, Kamlesh J Upadhyay, Ramesh Pandit, Tejas Shah, Ankit Hinsu, Pritesh Sabara, Apurvasinh Puvar, Janvi Raval, Monika Gandhi, Pinal Trivedi, Maharshi Pandya, Amit Kanani, Akanksha Verma, Nitin Savaliya, Raghawendra Kumar, Dinesh Kumar, Zuber Saiyed, Dipa Kinariwala, Disha Patel, Binita Aring, Neeta Khandelwal, Geeta Vaghela, Sonia Barve, Bhavesh Modi, K |
| hCoV-19/India/GBRC78b/2020 | EPI_ISL_447537 | 5/5/2020 | Gujarat Biotechnology Research Centre | Gujarat Biotechnology Research Centre | R D Dixit, Snehal Bagatharia, Kamlesh J Upadhyay, Ramesh Pandit, Tejas Shah, Ankit Hinsu, Pritesh Sabara, Apurvasinh Puvar, Janvi Raval, Monika Gandhi, Pinal Trivedi, Maharshi Pandya, Amit Kanani, Akanksha Verma, Nitin Savaliya, Raghawendra Kumar, Dinesh Kumar, Zuber Saiyed, Dipa Kinariwala, Disha Patel, Binita Aring, Neeta Khandelwal, Geeta Vaghela, Sonia Barve, Bhavesh Modi, Kairavi Joshi, G |
| hCoV-19/India/GBRC79a/2020 | EPI_ISL_447538 | 5/5/2020 | Gujarat Biotechnology Research Centre | Gujarat Biotechnology Research Centre | Snehal Bagatharia, Kamlesh J Upadhyay, Ramesh Pandit, Tejas Shah, Ankit Hinsu, Pritesh Sabara, Apurvasinh Puvar, Janvi Raval, Monika Gandhi, Pinal Trivedi, Maharshi Pandya, Amit Kanani, Akanksha Verma, Nitin Savaliya, Raghawendra Kumar, Dinesh Kumar, Zuber Saiyed, Dipa Kinariwala, Disha Patel, Binita Aring, Neeta Khandelwal, Geeta Vaghela, Sonia Barve, Bhavesh Modi, Kairavi Joshi, Gaurishan |
| hCoV-19/India/GBRC79b/2020 | EPI_ISL_447539 | 5/5/2020 | Gujarat Biotechnology Research Centre | Gujarat Biotechnology Research Centre | Kamlesh J Upadhyay, Ramesh Pandit, Tejas Shah, Ankit Hinsu, Pritesh Sabara, Apurvasinh Puvar, Janvi Raval, Monika Gandhi, Pinal Trivedi, Maharshi Pandya, Amit Kanani, Akanksha Verma, Nitin Savaliya, Raghawendra Kumar, Dinesh Kumar, Zuber Saiyed, Dipa Kinariwala, Disha Patel, Binita Aring, Neeta Khandelwal, Geeta Vaghela, Sonia Barve, Bhavesh Modi, Kairavi Joshi, Gaurishankar Shrimali, Nidhi S |
| hCoV-19/India/GBRC81a/2020 | EPI_ISL_447540 | 5/5/2020 | Gujarat Biotechnology Research Centre | Gujarat Biotechnology Research Centre | Ramesh Pandit, Tejas Shah, Ankit Hinsu, Pritesh Sabara, Apurvasinh Puvar, Janvi Raval, Monika Gandhi, Pinal Trivedi, Maharshi Pandya, Amit Kanani, Akanksha Verma, Nitin Savaliya, Raghawendra Kumar, Dinesh Kumar, Zuber Saiyed, Dipa Kinariwala, Disha Patel, Binita Aring, Neeta Khandelwal, Geeta Vaghela, Sonia Barve, Bhavesh Modi, Kairavi Joshi, Gaurishankar Shrimali, Nidhi Sood, Pranay Shah, R D |
| hCoV-19/India/GBRC81b/2020 | EPI_ISL_447541 | 5/5/2020 | Gujarat Biotechnology Research Centre | Gujarat Biotechnology Research Centre | Tejas Shah, Ankit Hinsu, Pritesh Sabara, Apurvasinh Puvar, Janvi Raval, Monika Gandhi, Pinal Trivedi, Maharshi Pandya, Amit Kanani, Akanksha Verma, Nitin Savaliya, Raghawendra Kumar, Dinesh Kumar, Zuber Saiyed, Dipa Kinariwala, Disha Patel, Binita Aring, Neeta Khandelwal, Geeta Vaghela, Sonia Barve, Bhavesh Modi, Kairavi Joshi, Gaurishankar Shrimali, Nidhi Sood, Pranay Shah, R D Dixit, Snehal Ba |
| hCoV-19/India/GBRC82a/2020 | EPI_ISL_447542 | 5/5/2020 | Gujarat Biotechnology Research Centre | Gujarat Biotechnology Research Centre | Ankit Hinsu, Pritesh Sabara, Apurvasinh Puvar, Janvi Raval, Monika Gandhi, Pinal Trivedi, Maharshi Pandya, Amit Kanani, Akanksha Verma, Nitin Savaliya, Raghawendra Kumar, Dinesh Kumar, Zuber Saiyed, Dipa Kinariwala, Disha Patel, Binita Aring, Neeta Khandelwal, Geeta Vaghela, Sonia Barve, Bhavesh Modi, Kairavi Joshi, Gaurishankar Shrimali, Nidhi Sood, Pranay Shah, R D Dixit, Snehal Bagatharia, Ka |
| hCoV-19/India/GBRC82b/2020 | EPI_ISL_447543 | 5/5/2020 | Gujarat Biotechnology Research Centre | Gujarat Biotechnology Research Centre | Pritesh Sabara, Apurvasinh Puvar, Janvi Raval, Monika Gandhi, Pinal Trivedi, Maharshi Pandya, Amit Kanani, Akanksha Verma, Nitin Savaliya, Raghawendra Kumar, Dinesh Kumar, Zuber Saiyed, Dipa Kinariwala, Disha Patel, Binita Aring, Neeta Khandelwal, Geeta Vaghela, Sonia Barve, Bhavesh Modi, Kairavi Joshi, Gaurishankar Shrimali, Nidhi Sood, Pranay Shah, R D Dixit, Snehal Bagatharia, Kamlesh J Upa |
| hCoV-19/India/GBRC83/2020 | EPI_ISL_447544 | 5/5/2020 | Gujarat Biotechnology Research Centre | Gujarat Biotechnology Research Centre | Apurvasinh Puvar, Janvi Raval, Monika Gandhi, Pinal Trivedi, Maharshi Pandya, Amit Kanani, Akanksha Verma, Nitin Savaliya, Raghawendra Kumar, Dinesh Kumar, Zuber Saiyed, Dipa Kinariwala, Disha Patel, Binita Aring, Neeta Khandelwal, Geeta Vaghela, Sonia Barve, Bhavesh Modi, Kairavi Joshi, Gaurishankar Shrimali, Nidhi Sood, Pranay Shah, R D Dixit, Snehal Bagatharia, Kamlesh J Upadhyay, Ramesh |
| hCoV-19/India/GBRC84/2020 | EPI_ISL_447545 | 5/5/2020 | Gujarat Biotechnology Research Centre | Gujarat Biotechnology Research Centre | Janvi Raval, Monika Gandhi, Pinal Trivedi, Maharshi Pandya, Amit Kanani, Akanksha Verma, Nitin Savaliya, Raghawendra Kumar, Dinesh Kumar, Zuber Saiyed, Dipa Kinariwala, Disha Patel, Binita Aring, Neeta Khandelwal, Geeta Vaghela, Sonia Barve, Bhavesh Modi, Kairavi Joshi, Gaurishankar Shrimali, Nidhi Sood, Pranay Shah, R D Dixit, Snehal Bagatharia, Kamlesh J Upadhyay, Ramesh Pandit, Tejas Shah |
| hCoV-19/India/GBRC85/2020 | EPI_ISL_447546 | 5/5/2020 | Gujarat Biotechnology Research Centre | Gujarat Biotechnology Research Centre | Monika Gandhi, Pinal Trivedi, Maharshi Pandya, Amit Kanani, Akanksha Verma, Nitin Savaliya, Raghawendra Kumar, Dinesh Kumar, Zuber Saiyed, Dipa Kinariwala, Disha Patel, Binita Aring, Neeta Khandelwal, Geeta Vaghela, Sonia Barve, Bhavesh Modi, Kairavi Joshi, Gaurishankar Shrimali, Nidhi Sood, Pranay Shah, R D Dixit, Snehal Bagatharia, Kamlesh J Upadhyay, Ramesh Pandit, Tejas Shah, Ankit Hinsu |
| hCoV-19/India/GBRC86/2020 | EPI_ISL_447547 | 4/28/2020 | GMERS Medical College and Hospital, Gandhinagar | Gujarat Biotechnology Research Centre | Pinal Trivedi, Maharshi Pandya, Amit Kanani, Akanksha Verma, Nitin Savaliya, Raghawendra Kumar, Dinesh Kumar, Zuber Saiyed, Dipa Kinariwala, Disha Patel, Binita Aring, Neeta Khandelwal, Geeta Vaghela, Sonia Barve, Bhavesh Modi, Kairavi Joshi, Gaurishankar Shrimali, Nidhi Sood, Pranay Shah, R D Dixit, Snehal Bagatharia, Kamlesh J Upadhyay, Ramesh Pandit, Tejas Shah, Ankit Hinsu, Pritesh Sabara |
| hCoV-19/India/GBRC87/2020 | EPI_ISL_447548 | 4/28/2020 | GMERS Medical College and Hospital, Gandhinagar | Gujarat Biotechnology Research Centre | Maharshi Pandya, Amit Kanani, Akanksha Verma, Nitin Savaliya, Raghawendra Kumar, Dinesh Kumar, Zuber Saiyed, Dipa Kinariwala, Disha Patel, Binita Aring, Neeta Khandelwal, Geeta Vaghela, Sonia Barve, Bhavesh Modi, Kairavi Joshi, Gaurishankar Shrimali, Nidhi Sood, Pranay Shah, R D Dixit, Snehal Bagatharia, Kamlesh J Upadhyay, Ramesh Pandit, Tejas Shah, Ankit Hinsu, Pritesh Sabara, Apurvasinh P |
| hCoV-19/India/GBRC88a/2020 | EPI_ISL_447549 | 5/3/2020 | GMERS Medical College and Hospital, Gandhinagar | Gujarat Biotechnology Research Centre | Amit Kanani, Akanksha Verma, Nitin Savaliya, Raghawendra Kumar, Dinesh Kumar, Zuber Saiyed, Dipa Kinariwala, Disha Patel, Binita Aring, Neeta Khandelwal, Geeta Vaghela, Sonia Barve, Bhavesh Modi, Kairavi Joshi, Gaurishankar Shrimali, Nidhi Sood, Pranay Shah, R D Dixit, Snehal Bagatharia, Kamlesh J Upadhyay, Ramesh Pandit, Tejas Shah, Ankit Hinsu, Pritesh Sabara, Apurvasinh Puvar, Janvi Raval, |
| hCoV-19/India/GBRC88b/2020 | EPI_ISL_447550 | 5/3/2020 | GMERS Medical College and Hospital, Gandhinagar | Gujarat Biotechnology Research Centre | Akanksha Verma, Nitin Savaliya, Raghawendra Kumar, Dinesh Kumar, Zuber Saiyed, Dipa Kinariwala, Disha Patel, Binita Aring, Neeta Khandelwal, Geeta Vaghela, Sonia Barve, Bhavesh Modi, Kairavi Joshi, Gaurishankar Shrimali, Nidhi Sood, Pranay Shah, R D Dixit, Snehal Bagatharia, Kamlesh J Upadhyay, Ramesh Pandit, Tejas Shah, Ankit Hinsu, Pritesh Sabara, Apurvasinh Puvar, Janvi Raval, Monika Gand |
| hCoV-19/India/GBRC89/2020 | EPI_ISL_447551 | 4/25/2020 | GMERS Medical College and Hospital, Gandhinagar | Gujarat Biotechnology Research Centre | Nitin Savaliya, Raghawendra Kumar, Dinesh Kumar, Zuber Saiyed, Dipa Kinariwala, Disha Patel, Binita Aring, Neeta Khandelwal, Geeta Vaghela, Sonia Barve, Bhavesh Modi, Kairavi Joshi, Gaurishankar Shrimali, Nidhi Sood, Pranay Shah, R D Dixit, Snehal Bagatharia, Kamlesh J Upadhyay, Ramesh Pandit, Tejas Shah, Ankit Hinsu, Pritesh Sabara, Apurvasinh Puvar, Janvi Raval, Monika Gandhi, Pinal Trivedi, M |
| hCoV-19/India/GBRC90/2020 | EPI_ISL_447552 | 5/2/2020 | GMERS Medical College and Hospital, Gandhinagar | Gujarat Biotechnology Research Centre | Raghawendra Kumar, Dinesh Kumar, Zuber Saiyed, Dipa Kinariwala, Disha Patel, Binita Aring, Neeta Khandelwal, Geeta Vaghela, Sonia Barve, Bhavesh Modi, Kairavi Joshi, Gaurishankar Shrimali, Nidhi Sood, Pranay Shah, R D Dixit, Snehal Bagatharia, Kamlesh J Upadhyay, Ramesh Pandit, Tejas Shah, Ankit Hinsu, Pritesh Sabara, Apurvasinh Puvar, Janvi Raval, Monika Gandhi, Pinal Trivedi, Maharshi Pandy |
| hCoV-19/India/GBRC91/2020 | EPI_ISL_447553 | 4/27/2020 | GMERS Medical College and Hospital, Gandhinagar | Gujarat Biotechnology Research Centre | Dinesh Kumar, Zuber Saiyed, Dipa Kinariwala, Disha Patel, Binita Aring, Neeta Khandelwal, Geeta Vaghela, Sonia Barve, Bhavesh Modi, Kairavi Joshi, Gaurishankar Shrimali, Nidhi Sood, Pranay Shah, R D Dixit, Snehal Bagatharia, Kamlesh J Upadhyay, Ramesh Pandit, Tejas Shah, Ankit Hinsu, Pritesh Sabara, Apurvasinh Puvar, Janvi Raval, Monika Gandhi, Pinal Trivedi, Maharshi Pandya, Amit Kanani, Akank |
| hCoV-19/India/GBRC92/2020 | EPI_ISL_447554 | 4/26/2020 | GMERS Medical College and Hospital, Gandhinagar | Gujarat Biotechnology Research Centre | Zuber Saiyed, Dipa Kinariwala, Disha Patel, Binita Aring, Neeta Khandelwal, Geeta Vaghela, Sonia Barve, Bhavesh Modi, Kairavi Joshi, Gaurishankar Shrimali, Nidhi Sood, Pranay Shah, R D Dixit, Snehal Bagatharia, Kamlesh J Upadhyay, Ramesh Pandit, Tejas Shah, Ankit Hinsu, Pritesh Sabara, Apurvasinh Puvar, Janvi Raval, Monika Gandhi, Pinal Trivedi, Maharshi Pandya, Amit Kanani, Akanksha Verma, Niti |
| hCoV-19/India/GBRC93/2020 | EPI_ISL_447555 | 4/28/2020 | GMERS Medical College and Hospital, Gandhinagar | Gujarat Biotechnology Research Centre | Dipa Kinariwala, Disha Patel, Binita Aring, Neeta Khandelwal, Geeta Vaghela, Sonia Barve, Bhavesh Modi, Kairavi Joshi, Gaurishankar Shrimali, Nidhi Sood, Pranay Shah, R D Dixit, Snehal Bagatharia, Kamlesh J Upadhyay, Ramesh Pandit, Tejas Shah, Ankit Hinsu, Pritesh Sabara, Apurvasinh Puvar, Janvi Raval, Monika Gandhi, Pinal Trivedi, Maharshi Pandya, Amit Kanani, Akanksha Verma, Nitin Savaliya, Rag |
| hCoV-19/India/CCMB_J021/2020 | EPI_ISL_447556 | 3/30/2020 | CSIR-Centre for Cellular and Molecular Biology | CSIR-Centre for Cellular and Molecular Biology | Sofia Banu, Payel Mukherjee, Priya Singh, Dhiviya Vedagiri, Divya Gupta, Vishal Sah, Santosh Kumar Kuncha, Krishnan Harinivas Harshan, Archana Bharadwaj Siva, Karthik Bharadwaj Tallapaka, Shagufta Khan, Lamuk Zaveri, Namami Gaur, Sakshi Shambhavi, Tulasi Nagabandi, Purushotham Vodnala, Rakesh K Mishra, Divya Tej Sowpati |
| hCoV-19/India/CCMB_J009/2020 | EPI_ISL_447557 | 3/30/2020 | CSIR-Centre for Cellular and Molecular Biology | CSIR-Centre for Cellular and Molecular Biology | Payel Mukherjee, Sofia Banu, Priya Singh, Dhiviya Vedagiri, Divya Gupta, Vishal Sah, Santosh Kumar Kuncha, Krishnan Harinivas Harshan, Archana Bharadwaj Siva, Karthik Bharadwaj Tallapaka, Shagufta Khan, Lamuk Zaveri, Namami Gaur, Sakshi Shambhavi, Tulasi Nagabandi, Purushotham Vodnala, Rakesh K Mishra, Divya Tej Sowpati |
| hCoV-19/India/CCMB_J014/2020 | EPI_ISL_447558 | 3/30/2020 | CSIR-Centre for Cellular and Molecular Biology | CSIR-Centre for Cellular and Molecular Biology | Payel Mukherjee, Sofia Banu, Priya Singh, Dhiviya Vedagiri, Divya Gupta, Vishal Sah, Santosh Kumar Kuncha, Krishnan Harinivas Harshan, Archana Bharadwaj Siva, Karthik Bharadwaj Tallapaka, Shagufta Khan, Lamuk Zaveri, Namami Gaur, Sakshi Shambhavi, Tulasi Nagabandi, Purushotham Vodnala, Rakesh K Mishra, Divya Tej Sowpati |
| hCoV-19/India/CCMB_J048/2020 | EPI_ISL_447559 | 3/31/2020 | CSIR-Centre for Cellular and Molecular Biology | CSIR-Centre for Cellular and Molecular Biology | Sofia Banu, Payel Mukherjee, Priya Singh, Dhiviya Vedagiri, Divya Gupta, Vishal Sah, Santosh Kumar Kuncha, Krishnan Harinivas Harshan, Archana Bharadwaj Siva, Karthik Bharadwaj Tallapaka, Shagufta Khan, Lamuk Zaveri, Namami Gaur, Sakshi Shambhavi, Tulasi Nagabandi, Purushotham Vodnala, Rakesh K Mishra, Divya Tej Sowpati |
| hCoV-19/India/CCMB_J043/2020 | EPI_ISL_447560 | 3/31/2020 | CSIR-Centre for Cellular and Molecular Biology | CSIR-Centre for Cellular and Molecular Biology | Payel Mukherjee, Sofia Banu, Priya Singh, Dhiviya Vedagiri, Divya Gupta, Vishal Sah, Santosh Kumar Kuncha, Krishnan Harinivas Harshan, Archana Bharadwaj Siva, Karthik Bharadwaj Tallapaka, Shagufta Khan, Lamuk Zaveri, Namami Gaur, Sakshi Shambhavi, Tulasi Nagabandi, Purushotham Vodnala, Rakesh K Mishra, Divya Tej Sowpati |
| hCoV-19/India/CCMB_J067/2020 | EPI_ISL_447561 | 3/31/2020 | CSIR-Centre for Cellular and Molecular Biology | CSIR-Centre for Cellular and Molecular Biology | Shagufta Khan, Lamuk Zaveri, Namami Gaur, Sakshi Shambhavi, Tulasi Nagabandi, Purushotham Vodnala, Payel Mukherjee, Sofia Banu, Priya Singh, Dhiviya Vedagiri, Divya Gupta, Vishal Sah, Santosh Kumar Kuncha, Krishnan Harinivas Harshan, Archana Bharadwaj Siva, Karthik Bharadwaj Tallapaka, Rakesh K Mishra, Divya Tej Sowpati |
| hCoV-19/India/CCMB_J068/2020 | EPI_ISL_447562 | 3/31/2020 | CSIR-Centre for Cellular and Molecular Biology | CSIR-Centre for Cellular and Molecular Biology | Shagufta Khan, Lamuk Zaveri, Namami Gaur, Sakshi Shambhavi, Tulasi Nagabandi, Purushotham Vodnala, Payel Mukherjee, Sofia Banu, Priya Singh, Dhiviya Vedagiri, Divya Gupta, Vishal Sah, Santosh Kumar Kuncha, Krishnan Harinivas Harshan, Archana Bharadwaj Siva, Karthik Bharadwaj Tallapaka, Rakesh K Mishra, Divya Tej Sowpati |
| hCoV-19/India/CCMB_J057/2020 | EPI_ISL_447563 | 3/31/2020 | CSIR-Centre for Cellular and Molecular Biology | CSIR-Centre for Cellular and Molecular Biology | Sakshi Shambhavi, Lamuk Zaveri, Shagufta Khan, Namami Gaur, Tulasi Nagabandi, Purushotham Vodnala, Payel Mukherjee, Sofia Banu, Priya Singh, Dhiviya Vedagiri, Divya Gupta, Vishal Sah, Santosh Kumar Kuncha, Krishnan Harinivas Harshan, Archana Bharadwaj Siva, Karthik Bharadwaj Tallapaka, Rakesh K Mishra, Divya Tej Sowpati |
| hCoV-19/India/CCMB_J212/2020 | EPI_ISL_447564 | 4/1/2020 | CSIR-Centre for Cellular and Molecular Biology | CSIR-Centre for Cellular and Molecular Biology | Sofia Banu, Payel Mukherjee, Priya Singh, Dhiviya Vedagiri, Divya Gupta, Vishal Sah, Santosh Kumar Kuncha, Krishnan Harinivas Harshan, Archana Bharadwaj Siva, Karthik Bharadwaj Tallapaka, Shagufta Khan, Lamuk Zaveri, Namami Gaur, Sakshi Shambhavi, Tulasi Nagabandi, Purushotham Vodnala, Rakesh K Mishra, Divya Tej Sowpati |
| hCoV-19/India/CCMB_J278/2020 | EPI_ISL_447565 | 4/1/2020 | CSIR-Centre for Cellular and Molecular Biology | CSIR-Centre for Cellular and Molecular Biology | Sofia Banu, Payel Mukherjee, Priya Singh, Dhiviya Vedagiri, Divya Gupta, Vishal Sah, Santosh Kumar Kuncha, Krishnan Harinivas Harshan, Archana Bharadwaj Siva, Karthik Bharadwaj Tallapaka, Shagufta Khan, Lamuk Zaveri, Namami Gaur, Sakshi Shambhavi, Tulasi Nagabandi, Purushotham Vodnala, Rakesh K Mishra, Divya Tej Sowpati |
| hCoV-19/India/CCMB_J131/2020 | EPI_ISL_447566 | 4/1/2020 | CSIR-Centre for Cellular and Molecular Biology | CSIR-Centre for Cellular and Molecular Biology | Sofia Banu, Payel Mukherjee, Priya Singh, Dhiviya Vedagiri, Divya Gupta, Vishal Sah, Santosh Kumar Kuncha, Krishnan Harinivas Harshan, Archana Bharadwaj Siva, Karthik Bharadwaj Tallapaka, Shagufta Khan, Lamuk Zaveri, Namami Gaur, Sakshi Shambhavi, Tulasi Nagabandi, Purushotham Vodnala, Rakesh K Mishra, Divya Tej Sowpati |
| hCoV-19/India/CCMB_J304/2020 | EPI_ISL_447567 | 4/1/2020 | CSIR-Centre for Cellular and Molecular Biology | CSIR-Centre for Cellular and Molecular Biology | Payel Mukherjee, Sofia Banu, Priya Singh, Dhiviya Vedagiri, Divya Gupta, Vishal Sah, Santosh Kumar Kuncha, Krishnan Harinivas Harshan, Archana Bharadwaj Siva, Karthik Bharadwaj Tallapaka, Shagufta Khan, Lamuk Zaveri, Namami Gaur, Sakshi Shambhavi, Tulasi Nagabandi, Purushotham Vodnala, Rakesh K Mishra, Divya Tej Sowpati |
| hCoV-19/India/CCMB_J166/2020 | EPI_ISL_447568 | 4/1/2020 | CSIR-Centre for Cellular and Molecular Biology | CSIR-Centre for Cellular and Molecular Biology | Shagufta Khan, Lamuk Zaveri, Namami Gaur, Sakshi Shambhavi, Tulasi Nagabandi, Purushotham Vodnala, Payel Mukherjee, Sofia Banu, Priya Singh, Dhiviya Vedagiri, Divya Gupta, Vishal Sah, Santosh Kumar Kuncha, Krishnan Harinivas Harshan, Archana Bharadwaj Siva, Karthik Bharadwaj Tallapaka, Rakesh K Mishra, Divya Tej Sowpati |
| hCoV-19/India/CCMB_J206/2020 | EPI_ISL_447569 | 4/1/2020 | CSIR-Centre for Cellular and Molecular Biology | CSIR-Centre for Cellular and Molecular Biology | Shagufta Khan, Lamuk Zaveri, Namami Gaur, Sakshi Shambhavi, Tulasi Nagabandi, Purushotham Vodnala, Payel Mukherjee, Sofia Banu, Priya Singh, Dhiviya Vedagiri, Divya Gupta, Vishal Sah, Santosh Kumar Kuncha, Krishnan Harinivas Harshan, Archana Bharadwaj Siva, Karthik Bharadwaj Tallapaka, Rakesh K Mishra, Divya Tej Sowpati |
| hCoV-19/India/CCMB_J207/2020 | EPI_ISL_447570 | 4/1/2020 | CSIR-Centre for Cellular and Molecular Biology | CSIR-Centre for Cellular and Molecular Biology | Lamuk Zaveri, Shagufta Khan, Namami Gaur, Sakshi Shambhavi, Tulasi Nagabandi, Purushotham Vodnala, Payel Mukherjee, Sofia Banu, Priya Singh, Dhiviya Vedagiri, Divya Gupta, Vishal Sah, Santosh Kumar Kuncha, Krishnan Harinivas Harshan, Archana Bharadwaj Siva, Karthik Bharadwaj Tallapaka, Rakesh K Mishra, Divya Tej Sowpati |
| hCoV-19/India/CCMB_J224/2020 | EPI_ISL_447571 | 4/1/2020 | CSIR-Centre for Cellular and Molecular Biology | CSIR-Centre for Cellular and Molecular Biology | Lamuk Zaveri, Shagufta Khan, Namami Gaur, Sakshi Shambhavi, Tulasi Nagabandi, Purushotham Vodnala, Payel Mukherjee, Sofia Banu, Priya Singh, Dhiviya Vedagiri, Divya Gupta, Vishal Sah, Santosh Kumar Kuncha, Krishnan Harinivas Harshan, Archana Bharadwaj Siva, Karthik Bharadwaj Tallapaka, Rakesh K Mishra, Divya Tej Sowpati |
| hCoV-19/India/CCMB_J230/2020 | EPI_ISL_447572 | 4/1/2020 | CSIR-Centre for Cellular and Molecular Biology | CSIR-Centre for Cellular and Molecular Biology | Lamuk Zaveri, Shagufta Khan, Namami Gaur, Sakshi Shambhavi, Tulasi Nagabandi, Purushotham Vodnala, Payel Mukherjee, Sofia Banu, Priya Singh, Dhiviya Vedagiri, Divya Gupta, Vishal Sah, Santosh Kumar Kuncha, Krishnan Harinivas Harshan, Archana Bharadwaj Siva, Karthik Bharadwaj Tallapaka, Rakesh K Mishra, Divya Tej Sowpati |
| hCoV-19/India/CCMB_J118/2020 | EPI_ISL_447573 | 4/1/2020 | CSIR-Centre for Cellular and Molecular Biology | CSIR-Centre for Cellular and Molecular Biology | Sakshi Shambhavi, Lamuk Zaveri, Shagufta Khan, Namami Gaur, Tulasi Nagabandi, Purushotham Vodnala, Payel Mukherjee, Sofia Banu, Priya Singh, Dhiviya Vedagiri, Divya Gupta, Vishal Sah, Santosh Kumar Kuncha, Krishnan Harinivas Harshan, Archana Bharadwaj Siva, Karthik Bharadwaj Tallapaka, Rakesh K Mishra, Divya Tej Sowpati |
| hCoV-19/India/CCMB_J208/2020 | EPI_ISL_447574 | 4/1/2020 | CSIR-Centre for Cellular and Molecular Biology | CSIR-Centre for Cellular and Molecular Biology | Namami Gaur, Sakshi Shambhavi, Lamuk Zaveri, Shagufta Khan, Tulasi Nagabandi, Purushotham Vodnala, Payel Mukherjee, Sofia Banu, Priya Singh, Dhiviya Vedagiri, Divya Gupta, Vishal Sah, Santosh Kumar Kuncha, Krishnan Harinivas Harshan, Archana Bharadwaj Siva, Karthik Bharadwaj Tallapaka, Rakesh K Mishra, Divya Tej Sowpati |
| hCoV-19/India/CCMB_J458/2020 | EPI_ISL_447575 | 4/2/2020 | CSIR-Centre for Cellular and Molecular Biology | CSIR-Centre for Cellular and Molecular Biology | Sofia Banu, Payel Mukherjee, Priya Singh, Dhiviya Vedagiri, Divya Gupta, Vishal Sah, Santosh Kumar Kuncha, Krishnan Harinivas Harshan, Archana Bharadwaj Siva, Karthik Bharadwaj Tallapaka, Shagufta Khan, Lamuk Zaveri, Namami Gaur, Sakshi Shambhavi, Tulasi Nagabandi, Purushotham Vodnala, Rakesh K Mishra, Divya Tej Sowpati |
| hCoV-19/India/CCMB_J232/2020 | EPI_ISL_447576 | 4/2/2020 | CSIR-Centre for Cellular and Molecular Biology | CSIR-Centre for Cellular and Molecular Biology | Namami Gaur, Sakshi Shambhavi, Lamuk Zaveri, Shagufta Khan, Tulasi Nagabandi, Purushotham Vodnala, Payel Mukherjee, Sofia Banu, Priya Singh, Dhiviya Vedagiri, Divya Gupta, Vishal Sah, Santosh Kumar Kuncha, Krishnan Harinivas Harshan, Archana Bharadwaj Siva, Karthik Bharadwaj Tallapaka, Rakesh K Mishra, Divya Tej Sowpati |
| hCoV-19/India/CCMB_J321/2020 | EPI_ISL_447577 | 4/2/2020 | CSIR-Centre for Cellular and Molecular Biology | CSIR-Centre for Cellular and Molecular Biology | Namami Gaur, Sakshi Shambhavi, Lamuk Zaveri, Shagufta Khan, Tulasi Nagabandi, Purushotham Vodnala, Payel Mukherjee, Sofia Banu, Priya Singh, Dhiviya Vedagiri, Divya Gupta, Vishal Sah, Santosh Kumar Kuncha, Krishnan Harinivas Harshan, Archana Bharadwaj Siva, Karthik Bharadwaj Tallapaka, Rakesh K Mishra, Divya Tej Sowpati |
| hCoV-19/India/CCMB_J327/2020 | EPI_ISL_447578 | 4/2/2020 | CSIR-Centre for Cellular and Molecular Biology | CSIR-Centre for Cellular and Molecular Biology | Namami Gaur, Sakshi Shambhavi, Lamuk Zaveri, Shagufta Khan, Tulasi Nagabandi, Purushotham Vodnala, Payel Mukherjee, Sofia Banu, Priya Singh, Dhiviya Vedagiri, Divya Gupta, Vishal Sah, Santosh Kumar Kuncha, Krishnan Harinivas Harshan, Archana Bharadwaj Siva, Karthik Bharadwaj Tallapaka, Rakesh K Mishra, Divya Tej Sowpati |
| hCoV-19/India/CCMB_K128/2020 | EPI_ISL_447579 | 4/10/2020 | CSIR-Centre for Cellular and Molecular Biology | CSIR-Centre for Cellular and Molecular Biology | Tulasi Nagabandi, Namami Gaur, Sakshi Shambhavi, Lamuk Zaveri, Shagufta Khan, Purushotham Vodnala, Payel Mukherjee, Sofia Banu, Priya Singh, Dhiviya Vedagiri, Divya Gupta, Vishal Sah, Santosh Kumar Kuncha, Krishnan Harinivas Harshan, Archana Bharadwaj Siva, Karthik Bharadwaj Tallapaka, Rakesh K Mishra, Divya Tej Sowpati |
| hCoV-19/India/CCMB_K496/2020 | EPI_ISL_447580 | 4/13/2020 | CSIR-Centre for Cellular and Molecular Biology | CSIR-Centre for Cellular and Molecular Biology | Tulasi Nagabandi, Namami Gaur, Sakshi Shambhavi, Lamuk Zaveri, Shagufta Khan, Purushotham Vodnala, Payel Mukherjee, Sofia Banu, Priya Singh, Dhiviya Vedagiri, Divya Gupta, Vishal Sah, Santosh Kumar Kuncha, Krishnan Harinivas Harshan, Archana Bharadwaj Siva, Karthik Bharadwaj Tallapaka, Rakesh K Mishra, Divya Tej Sowpati |
| hCoV-19/India/CCMB_K600B/2020 | EPI_ISL_447581 | 4/14/2020 | CSIR-Centre for Cellular and Molecular Biology | CSIR-Centre for Cellular and Molecular Biology | Sakshi Shambhavi, Lamuk Zaveri, Shagufta Khan, Namami Gaur, Tulasi Nagabandi, Purushotham Vodnala, Payel Mukherjee, Sofia Banu, Priya Singh, Dhiviya Vedagiri, Divya Gupta, Vishal Sah, Santosh Kumar Kuncha, Krishnan Harinivas Harshan, Archana Bharadwaj Siva, Karthik Bharadwaj Tallapaka, Rakesh K Mishra, Divya Tej Sowpati |
| hCoV-19/India/CCMB_K601/2020 | EPI_ISL_447582 | 4/14/2020 | CSIR-Centre for Cellular and Molecular Biology | CSIR-Centre for Cellular and Molecular Biology | Tulasi Nagabandi, Namami Gaur, Sakshi Shambhavi, Lamuk Zaveri, Shagufta Khan, Purushotham Vodnala, Payel Mukherjee, Sofia Banu, Priya Singh, Dhiviya Vedagiri, Divya Gupta, Vishal Sah, Santosh Kumar Kuncha, Krishnan Harinivas Harshan, Archana Bharadwaj Siva, Karthik Bharadwaj Tallapaka, Rakesh K Mishra, Divya Tej Sowpati |
| hCoV-19/India/CCMB_L007/2020 | EPI_ISL_447583 | 4/16/2020 | CSIR-Centre for Cellular and Molecular Biology | CSIR-Centre for Cellular and Molecular Biology | Tulasi Nagabandi, Namami Gaur, Sakshi Shambhavi, Lamuk Zaveri, Shagufta Khan, Purushotham Vodnala, Payel Mukherjee, Sofia Banu, Priya Singh, Dhiviya Vedagiri, Divya Gupta, Vishal Sah, Santosh Kumar Kuncha, Krishnan Harinivas Harshan, Archana Bharadwaj Siva, Karthik Bharadwaj Tallapaka, Rakesh K Mishra, Divya Tej Sowpati |
| hCoV-19/India/CCMB_C1-12/2020 | EPI_ISL_447584 | 4/16/2020 | Tamil Nadu Veterinary and Animal Sciences University | CSIR-Centre for Cellular and Molecular Biology | K Kaveri, S Sivasubramanian, S Vennila, P Padmapriya, R Kiruba, S Magesh, G Dhinakar Raj, G Ravi Kumar, Payel Mukherjee, Tulasi Nagabandi, Namami Gaur, Sakshi Shambhavi, Lamuk Zaveri, Shagufta Khan, Purushotham Vodnala, Sofia Banu, Priya Singh, Dhiviya Vedagiri, Divya Gupta, Vishal Sah, Santosh Kumar Kuncha, Krishnan Harinivas Harshan, Archana Bharadwaj Siva, Karthik Bharadwaj Tallapak |
| hCoV-19/India/CCMB_C2-13/2020 | EPI_ISL_447585 | 4/16/2020 | Tamil Nadu Veterinary and Animal Sciences University | CSIR-Centre for Cellular and Molecular Biology | K Kaveri, S Sivasubramanian, S Vennila, P Padmapriya, R Kiruba, S Magesh, G Dhinakar Raj, G Ravi Kumar, Payel Mukherjee, Tulasi Nagabandi, Namami Gaur, Sakshi Shambhavi, Lamuk Zaveri, Shagufta Khan, Purushotham Vodnala, Sofia Banu, Priya Singh, Dhiviya Vedagiri, Divya Gupta, Vishal Sah, Santosh Kumar Kuncha, Krishnan Harinivas Harshan, Archana Bharadwaj Siva, Karthik Bharadwaj Tallapak |
| hCoV-19/India/CCMB_C3-14/2020 | EPI_ISL_447586 | 4/16/2020 | Tamil Nadu Veterinary and Animal Sciences University | CSIR-Centre for Cellular and Molecular Biology | K Kaveri, S Sivasubramanian, S Vennila, P Padmapriya, R Kiruba, S Magesh, G Dhinakar Raj, G Ravi Kumar, Payel Mukherjee, Tulasi Nagabandi, Namami Gaur, Sakshi Shambhavi, Lamuk Zaveri, Shagufta Khan, Purushotham Vodnala, Sofia Banu, Priya Singh, Dhiviya Vedagiri, Divya Gupta, Vishal Sah, Santosh Kumar Kuncha, Krishnan Harinivas Harshan, Archana Bharadwaj Siva, Karthik Bharadwaj Tallapak |
| hCoV-19/India/CCMB_C4-15/2020 | EPI_ISL_447587 | 4/20/2020 | Tamil Nadu Veterinary and Animal Sciences University | CSIR-Centre for Cellular and Molecular Biology | K Kaveri, S Sivasubramanian, S Vennila, P Padmapriya, R Kiruba, S Magesh, G Dhinakar Raj, G Ravi Kumar, Payel Mukherjee, Tulasi Nagabandi, Namami Gaur, Sakshi Shambhavi, Lamuk Zaveri, Shagufta Khan, Purushotham Vodnala, Sofia Banu, Priya Singh, Dhiviya Vedagiri, Divya Gupta, Vishal Sah, Santosh Kumar Kuncha, Krishnan Harinivas Harshan, Archana Bharadwaj Siva, Karthik Bharadwaj Tallapak |
| hCoV-19/India/CCMB_J045/2020 | EPI_ISL_447847 | 3/31/2020 | CSIR-Centre for Cellular and Molecular Biology | CSIR-Centre for Cellular and Molecular Biology | Payel Mukherjee, Sofia Banu, Priya Singh, Dhiviya Vedagiri, Divya Gupta, Vishal Sah, Santosh Kumar Kuncha, Krishnan Harinivas Harshan, Archana Bharadwaj Siva, Karthik Bharadwaj Tallapaka, Shagufta Khan, Lamuk Zaveri, Namami Gaur, Sakshi Shambhavi, Tulasi Nagabandi, Purushotham Vodnala, Rakesh K Mishra, Divya Tej Sowpati |
| hCoV-19/India/CCMB_J165/2020 | EPI_ISL_447848 | 4/1/2020 | CSIR-Centre for Cellular and Molecular Biology | CSIR-Centre for Cellular and Molecular Biology | Sofia Banu, Payel Mukherjee, Priya Singh, Dhiviya Vedagiri, Divya Gupta, Vishal Sah, Santosh Kumar Kuncha, Krishnan Harinivas Harshan, Archana Bharadwaj Siva, Karthik Bharadwaj Tallapaka, Shagufta Khan, Lamuk Zaveri, Namami Gaur, Sakshi Shambhavi, Tulasi Nagabandi, Purushotham Vodnala, Rakesh K Mishra, Divya Tej Sowpati |
| hCoV-19/India/CCMB_J107/2020 | EPI_ISL_447849 | 4/1/2020 | CSIR-Centre for Cellular and Molecular Biology | CSIR-Centre for Cellular and Molecular Biology | Shagufta Khan, Lamuk Zaveri, Namami Gaur, Sakshi Shambhavi, Tulasi Nagabandi, Purushotham Vodnala, Payel Mukherjee, Sofia Banu, Priya Singh, Dhiviya Vedagiri, Divya Gupta, Vishal Sah, Santosh Kumar Kuncha, Krishnan Harinivas Harshan, Archana Bharadwaj Siva, Karthik Bharadwaj Tallapaka, Rakesh K Mishra, Divya Tej Sowpati |
| hCoV-19/India/CCMB_J132/2020 | EPI_ISL_447850 | 4/1/2020 | CSIR-Centre for Cellular and Molecular Biology | CSIR-Centre for Cellular and Molecular Biology | Shagufta Khan, Lamuk Zaveri, Namami Gaur, Sakshi Shambhavi, Tulasi Nagabandi, Purushotham Vodnala, Payel Mukherjee, Sofia Banu, Priya Singh, Dhiviya Vedagiri, Divya Gupta, Vishal Sah, Santosh Kumar Kuncha, Krishnan Harinivas Harshan, Archana Bharadwaj Siva, Karthik Bharadwaj Tallapaka, Rakesh K Mishra, Divya Tej Sowpati |
| hCoV-19/India/CCMB_J223/2020 | EPI_ISL_447851 | 4/1/2020 | CSIR-Centre for Cellular and Molecular Biology | CSIR-Centre for Cellular and Molecular Biology | Lamuk Zaveri, Shagufta Khan, Namami Gaur, Sakshi Shambhavi, Tulasi Nagabandi, Purushotham Vodnala, Payel Mukherjee, Sofia Banu, Priya Singh, Dhiviya Vedagiri, Divya Gupta, Vishal Sah, Santosh Kumar Kuncha, Krishnan Harinivas Harshan, Archana Bharadwaj Siva, Karthik Bharadwaj Tallapaka, Rakesh K Mishra, Divya Tej Sowpati |
| hCoV-19/India/CCMB_J235/2020 | EPI_ISL_447852 | 4/1/2020 | CSIR-Centre for Cellular and Molecular Biology | CSIR-Centre for Cellular and Molecular Biology | Lamuk Zaveri, Shagufta Khan, Namami Gaur, Sakshi Shambhavi, Tulasi Nagabandi, Purushotham Vodnala, Payel Mukherjee, Sofia Banu, Priya Singh, Dhiviya Vedagiri, Divya Gupta, Vishal Sah, Santosh Kumar Kuncha, Krishnan Harinivas Harshan, Archana Bharadwaj Siva, Karthik Bharadwaj Tallapaka, Rakesh K Mishra, Divya Tej Sowpati |
| hCoV-19/India/CCMB_J247/2020 | EPI_ISL_447853 | 4/1/2020 | CSIR-Centre for Cellular and Molecular Biology | CSIR-Centre for Cellular and Molecular Biology | Namami Gaur, Sakshi Shambhavi, Lamuk Zaveri, Shagufta Khan, Tulasi Nagabandi, Purushotham Vodnala, Payel Mukherjee, Sofia Banu, Priya Singh, Dhiviya Vedagiri, Divya Gupta, Vishal Sah, Santosh Kumar Kuncha, Krishnan Harinivas Harshan, Archana Bharadwaj Siva, Karthik Bharadwaj Tallapaka, Rakesh K Mishra, Divya Tej Sowpati |
| hCoV-19/India/CCMB_J350/2020 | EPI_ISL_447854 | 4/2/2020 | CSIR-Centre for Cellular and Molecular Biology | CSIR-Centre for Cellular and Molecular Biology | Payel Mukherjee, Sofia Banu, Priya Singh, Dhiviya Vedagiri, Divya Gupta, Vishal Sah, Santosh Kumar Kuncha, Krishnan Harinivas Harshan, Archana Bharadwaj Siva, Karthik Bharadwaj Tallapaka, Shagufta Khan, Lamuk Zaveri, Namami Gaur, Sakshi Shambhavi, Tulasi Nagabandi, Purushotham Vodnala, Rakesh K Mishra, Divya Tej Sowpati |
| hCoV-19/India/CCMB_J318/2020 | EPI_ISL_447855 | 4/2/2020 | CSIR-Centre for Cellular and Molecular Biology | CSIR-Centre for Cellular and Molecular Biology | Lamuk Zaveri, Shagufta Khan, Namami Gaur, Sakshi Shambhavi, Tulasi Nagabandi, Purushotham Vodnala, Payel Mukherjee, Sofia Banu, Priya Singh, Dhiviya Vedagiri, Divya Gupta, Vishal Sah, Santosh Kumar Kuncha, Krishnan Harinivas Harshan, Archana Bharadwaj Siva, Karthik Bharadwaj Tallapaka, Rakesh K Mishra, Divya Tej Sowpati |
| hCoV-19/India/CCMB_J374/2020 | EPI_ISL_447856 | 4/2/2020 | CSIR-Centre for Cellular and Molecular Biology | CSIR-Centre for Cellular and Molecular Biology | Sakshi Shambhavi, Lamuk Zaveri, Shagufta Khan, Namami Gaur, Tulasi Nagabandi, Purushotham Vodnala, Payel Mukherjee, Sofia Banu, Priya Singh, Dhiviya Vedagiri, Divya Gupta, Vishal Sah, Santosh Kumar Kuncha, Krishnan Harinivas Harshan, Archana Bharadwaj Siva, Karthik Bharadwaj Tallapaka, Rakesh K Mishra, Divya Tej Sowpati |
| hCoV-19/India/CCMB_J390/2020 | EPI_ISL_447857 | 4/2/2020 | CSIR-Centre for Cellular and Molecular Biology | CSIR-Centre for Cellular and Molecular Biology | Sakshi Shambhavi, Lamuk Zaveri, Shagufta Khan, Namami Gaur, Tulasi Nagabandi, Purushotham Vodnala, Payel Mukherjee, Sofia Banu, Priya Singh, Dhiviya Vedagiri, Divya Gupta, Vishal Sah, Santosh Kumar Kuncha, Krishnan Harinivas Harshan, Archana Bharadwaj Siva, Karthik Bharadwaj Tallapaka, Rakesh K Mishra, Divya Tej Sowpati |
| hCoV-19/India/CCMB_J812/2020 | EPI_ISL_447858 | 4/6/2020 | CSIR-Centre for Cellular and Molecular Biology | CSIR-Centre for Cellular and Molecular Biology | Sakshi Shambhavi, Lamuk Zaveri, Shagufta Khan, Namami Gaur, Tulasi Nagabandi, Purushotham Vodnala, Payel Mukherjee, Sofia Banu, Priya Singh, Dhiviya Vedagiri, Divya Gupta, Vishal Sah, Santosh Kumar Kuncha, Krishnan Harinivas Harshan, Archana Bharadwaj Siva, Karthik Bharadwaj Tallapaka, Rakesh K Mishra, Divya Tej Sowpati |
| hCoV-19/India/CCMB_K771/2020 | EPI_ISL_447859 | 4/15/2020 | CSIR-Centre for Cellular and Molecular Biology | CSIR-Centre for Cellular and Molecular Biology | Payel Mukherjee, Sofia Banu, Priya Singh, Dhiviya Vedagiri, Divya Gupta, Vishal Sah, Santosh Kumar Kuncha, Krishnan Harinivas Harshan, Archana Bharadwaj Siva, Karthik Bharadwaj Tallapaka, Shagufta Khan, Lamuk Zaveri, Namami Gaur, Sakshi Shambhavi, Tulasi Nagabandi, Purushotham Vodnala, Rakesh K Mishra, Divya Tej Sowpati |
| hCoV-19/India/CCMB_K849/2020 | EPI_ISL_447860 | 4/16/2020 | CSIR-Centre for Cellular and Molecular Biology | CSIR-Centre for Cellular and Molecular Biology | Tulasi Nagabandi, Namami Gaur, Sakshi Shambhavi, Lamuk Zaveri, Shagufta Khan, Purushotham Vodnala, Payel Mukherjee, Sofia Banu, Priya Singh, Dhiviya Vedagiri, Divya Gupta, Vishal Sah, Santosh Kumar Kuncha, Krishnan Harinivas Harshan, Archana Bharadwaj Siva, Karthik Bharadwaj Tallapaka, Rakesh K Mishra, Divya Tej Sowpati |
| hCoV-19/India/CCMB_L149/2020 | EPI_ISL_447861 | 4/16/2020 | CSIR-Centre for Cellular and Molecular Biology | CSIR-Centre for Cellular and Molecular Biology | Tulasi Nagabandi, Namami Gaur, Sakshi Shambhavi, Lamuk Zaveri, Shagufta Khan, Purushotham Vodnala, Payel Mukherjee, Sofia Banu, Priya Singh, Dhiviya Vedagiri, Divya Gupta, Vishal Sah, Santosh Kumar Kuncha, Krishnan Harinivas Harshan, Archana Bharadwaj Siva, Karthik Bharadwaj Tallapaka, Rakesh K Mishra, Divya Tej Sowpati |
| hCoV-19/India/CCMB_J047/2020 | EPI_ISL_447862 | 3/31/2020 | CSIR-Centre for Cellular and Molecular Biology | CSIR-Centre for Cellular and Molecular Biology | Payel Mukherjee, Sofia Banu, Priya Singh, Dhiviya Vedagiri, Divya Gupta, Vishal Sah, Santosh Kumar Kuncha, Krishnan Harinivas Harshan, Archana Bharadwaj Siva, Karthik Bharadwaj Tallapaka, Shagufta Khan, Lamuk Zaveri, Namami Gaur, Sakshi Shambhavi, Tulasi Nagabandi, Purushotham Vodnala, Rakesh K Mishra, Divya Tej Sowpati |
| hCoV-19/India/CCMB_K599/2020 | EPI_ISL_447863 | 4/14/2020 | CSIR-Centre for Cellular and Molecular Biology | CSIR-Centre for Cellular and Molecular Biology | Payel Mukherjee, Sofia Banu, Priya Singh, Dhiviya Vedagiri, Divya Gupta, Vishal Sah, Santosh Kumar Kuncha, Krishnan Harinivas Harshan, Archana Bharadwaj Siva, Karthik Bharadwaj Tallapaka, Shagufta Khan, Lamuk Zaveri, Namami Gaur, Sakshi Shambhavi, Tulasi Nagabandi, Purushotham Vodnala, Rakesh K Mishra, Divya Tej Sowpati |
| hCoV-19/India/CCMB_L044/2020 | EPI_ISL_447864 | 4/16/2020 | CSIR-Centre for Cellular and Molecular Biology | CSIR-Centre for Cellular and Molecular Biology | Payel Mukherjee, Sofia Banu, Priya Singh, Dhiviya Vedagiri, Divya Gupta, Vishal Sah, Santosh Kumar Kuncha, Krishnan Harinivas Harshan, Archana Bharadwaj Siva, Karthik Bharadwaj Tallapaka, Shagufta Khan, Lamuk Zaveri, Namami Gaur, Sakshi Shambhavi, Tulasi Nagabandi, Purushotham Vodnala, Rakesh K Mishra, Divya Tej Sowpati |
| hCoV-19/India/CCMB_K499/2020 | EPI_ISL_447865 | 4/14/2020 | CSIR-Centre for Cellular and Molecular Biology | CSIR-Centre for Cellular and Molecular Biology | Sofia Banu, Payel Mukherjee, Priya Singh, Dhiviya Vedagiri, Divya Gupta, Vishal Sah, Santosh Kumar Kuncha, Krishnan Harinivas Harshan, Archana Bharadwaj Siva, Karthik Bharadwaj Tallapaka, Shagufta Khan, Lamuk Zaveri, Namami Gaur, Sakshi Shambhavi, Tulasi Nagabandi, Purushotham Vodnala, Rakesh K Mishra, Divya Tej Sowpati |
| hCoV-19/India/CCMB_K500/2020 | EPI_ISL_447866 | 4/14/2020 | CSIR-Centre for Cellular and Molecular Biology | CSIR-Centre for Cellular and Molecular Biology | Sofia Banu, Payel Mukherjee, Priya Singh, Dhiviya Vedagiri, Divya Gupta, Vishal Sah, Santosh Kumar Kuncha, Krishnan Harinivas Harshan, Archana Bharadwaj Siva, Karthik Bharadwaj Tallapaka, Shagufta Khan, Lamuk Zaveri, Namami Gaur, Sakshi Shambhavi, Tulasi Nagabandi, Purushotham Vodnala, Rakesh K Mishra, Divya Tej Sowpati |
| hCoV-19/India/CCMB_NIV1/2020 | EPI_ISL_450321 | 4/4/2020 | NIV Pune | CSIR-Centre for Cellular and Molecular Biology | Dr V A Potdar, Dr ML Choudhary,Dr Priya Abraham,V. Vipat, S. Jadhav, U. Saha, H. Kengle, A. Awhale, A. Jagtap, A. Gondhalikar, V Malik, N Srivastava, S. Digraskar, P. Malsane, S. Hundekar, K. Patel, Yogesh Balakartik, M. Kakade, S. Jadhav, R. Gunjikar, V. Awtade, S. Bhorekar, P Shinde, S. Salve, B. Minhas S. Bharadwaj, H Kaushal Y. Gurav, S. Tomar,Payel Mukherjee, Sofia Banu, Priya Singh, Dhiviya Ve |
| hCoV-19/India/CCMB_NIV2/2020 | EPI_ISL_450322 | 4/4/2020 | NIV Pune | CSIR-Centre for Cellular and Molecular Biology | Dr V A Potdar, Dr ML Choudhary,Dr Priya Abraham,V. Vipat, S. Jadhav, U. Saha, H. Kengle, A. Awhale, A. Jagtap, A. Gondhalikar, V Malik, N Srivastava, S. Digraskar, P. Malsane, S. Hundekar, K. Patel, Yogesh Balakartik, M. Kakade, S. Jadhav, R. Gunjikar, V. Awtade, S. Bhorekar, P Shinde, S. Salve, B. Minhas S. Bharadwaj, H Kaushal Y. Gurav, S. Tomar,Sofia Banu, Payel Mukherjee, Priya Singh, Dhiviya Ve |
| hCoV-19/India/CCMB_NIV3/2020 | EPI_ISL_450323 | 3/27/2020 | NIV Pune | CSIR-Centre for Cellular and Molecular Biology | Dr V A Potdar, Dr ML Choudhary,Dr Priya Abraham,V. Vipat, S. Jadhav, U. Saha, H. Kengle, A. Awhale, A. Jagtap, A. Gondhalikar, V Malik, N Srivastava, S. Digraskar, P. Malsane, S. Hundekar, K. Patel, Yogesh Balakartik, M. Kakade, S. Jadhav, R. Gunjikar, V. Awtade, S. Bhorekar, P Shinde, S. Salve, B. Minhas S. Bharadwaj, H Kaushal Y. Gurav, S. Tomar,Payel Mukherjee, Sofia Banu, Priya Singh, Dhiviya Ve |
| hCoV-19/India/CCMB_NIV4/2020 | EPI_ISL_450324 | 3/26/2020 | NIV Pune | CSIR-Centre for Cellular and Molecular Biology | Dr V A Potdar, Dr ML Choudhary,Dr Priya Abraham,V. Vipat, S. Jadhav, U. Saha, H. Kengle, A. Awhale, A. Jagtap, A. Gondhalikar, V Malik, N Srivastava, S. Digraskar, P. Malsane, S. Hundekar, K. Patel, Yogesh Balakartik, M. Kakade, S. Jadhav, R. Gunjikar, V. Awtade, S. Bhorekar, P Shinde, S. Salve, B. Minhas S. Bharadwaj, H Kaushal Y. Gurav, S. Tomar,Sofia Banu, Payel Mukherjee, Priya Singh, Dhiviya Ve |
| hCoV-19/India/CCMB_NIV5/2020 | EPI_ISL_450325 | 3/17/2020 | NIV Pune | CSIR-Centre for Cellular and Molecular Biology | Dr V A Potdar, Dr ML Choudhary,Dr Priya Abraham,V. Vipat, S. Jadhav, U. Saha, H. Kengle, A. Awhale, A. Jagtap, A. Gondhalikar, V Malik, N Srivastava, S. Digraskar, P. Malsane, S. Hundekar, K. Patel, Yogesh Balakartik, M. Kakade, S. Jadhav, R. Gunjikar, V. Awtade, S. Bhorekar, P Shinde, S. Salve, B. Minhas S. Bharadwaj, H Kaushal Y. Gurav, S. Tomar,Payel Mukherjee, Sofia Banu, Priya Singh, Dhiviya Ve |
| hCoV-19/India/CCMB_J125/2020 | EPI_ISL_450326 | 4/1/2020 | CSIR-Centre for Cellular and Molecular Biology | CSIR-Centre for Cellular and Molecular Biology | Payel Mukherjee, Sofia Banu, Priya Singh, Dhiviya Vedagiri, Divya Gupta, Vishal Sah, Santosh Kumar Kuncha, Krishnan Harinivas Harshan, Archana Bharadwaj Siva, Karthik Bharadwaj Tallapaka, Shagufta Khan, Lamuk Zaveri, Namami Gaur, Sakshi Shambhavi, Tulasi Nagabandi, Purushotham Vodnala,G. Aditya Kumar, Koushick Sivakumar, Pooja Ramesh Gupta, Rajan Kumar Jha, Shraddha Vijay Lahoti, Dee |
| hCoV-19/India/CCMB_J199/2020 | EPI_ISL_450327 | 4/1/2020 | CSIR-Centre for Cellular and Molecular Biology | CSIR-Centre for Cellular and Molecular Biology | Sofia Banu, Payel Mukherjee, Priya Singh, Dhiviya Vedagiri, Divya Gupta, Vishal Sah, Santosh Kumar Kuncha, Krishnan Harinivas Harshan, Archana Bharadwaj Siva, Karthik Bharadwaj Tallapaka, Shagufta Khan, Lamuk Zaveri, Namami Gaur, Sakshi Shambhavi, Tulasi Nagabandi, Purushotham Vodnala, Disha Nanda, Divya Das, Jotin Gogoi, Manish Bhattacharjee, Ravi Prasad Mukku, Renu Sudhakar, Somesh |
| hCoV-19/India/CCMB_J283/2020 | EPI_ISL_450328 | 4/1/2020 | CSIR-Centre for Cellular and Molecular Biology | CSIR-Centre for Cellular and Molecular Biology | Shagufta Khan, Lamuk Zaveri, Namami Gaur, Sakshi Shambhavi, Tulasi Nagabandi, Purushotham Vodnala, Payel Mukherjee, Sofia Banu, Priya Singh, Dhiviya Vedagiri, Divya Gupta, Vishal Sah, Santosh Kumar Kuncha, Krishnan Harinivas Harshan, Archana Bharadwaj Siva, Karthik Bharadwaj Tallapaka, Zeba Rizvi, Zuberwasim Sayyad, Kakade Aishwarya Arun, Amrutha H C, Ananga Ghosh, Kezia J Ann, Radh |
| hCoV-19/India/CCMB_J288/2020 | EPI_ISL_450329 | 4/1/2020 | CSIR-Centre for Cellular and Molecular Biology | CSIR-Centre for Cellular and Molecular Biology | Namami Gaur, Sakshi Shambhavi, Lamuk Zaveri, Shagufta Khan, Tulasi Nagabandi, Purushotham Vodnala, Payel Mukherjee, Sofia Banu, Priya Singh, Dhiviya Vedagiri, Divya Gupta, Vishal Sah, Santosh Kumar Kuncha, Krishnan Harinivas Harshan, Archana Bharadwaj Siva, Karthik Bharadwaj Tallapaka, Sonu Uday, Sudipta Mondal, Annapoorna P Karthyayani, Debabrata Jana, Debrya Saha, Gokulan C G, Gun |
| hCoV-19/India/CCMB_J375/2020 | EPI_ISL_450330 | 4/2/2020 | CSIR-Centre for Cellular and Molecular Biology | CSIR-Centre for Cellular and Molecular Biology | Sakshi Shambhavi, Lamuk Zaveri, Shagufta Khan, Namami Gaur, Tulasi Nagabandi, Purushotham Vodnala, Payel Mukherjee, Sofia Banu, Priya Singh, Dhiviya Vedagiri, Divya Gupta, Vishal Sah, Santosh Kumar Kuncha, Krishnan Harinivas Harshan, Archana Bharadwaj Siva, Karthik Bharadwaj Tallapaka,Preethi Jampala, Sharada Ravi Iyer, Sulagana Mukherjee, Swetha Sundar, Peddapuvala Sai Uday Kiran, Un |
| hCoV-19/India/CCMB_J220/2020 | EPI_ISL_450331 | 4/1/2020 | CSIR-Centre for Cellular and Molecular Biology | CSIR-Centre for Cellular and Molecular Biology | Tulasi Nagabandi, Namami Gaur, Sakshi Shambhavi, Lamuk Zaveri, Shagufta Khan, Purushotham Vodnala, Payel Mukherjee, Sofia Banu, Priya Singh, Dhiviya Vedagiri, Divya Gupta, Vishal Sah, Santosh Kumar Kuncha, Krishnan Harinivas Harshan, Archana Bharadwaj Siva, Karthik Bharadwaj Tallapaka,G. Aditya Kumar, Koushick Sivakumar, Pooja Ramesh Gupta, Rajan Kumar Jha, Shraddha Vijay Lahoti, Dee |
| hCoV-19/India/CCMB_J222/2020 | EPI_ISL_450332 | 4/1/2020 | CSIR-Centre for Cellular and Molecular Biology | CSIR-Centre for Cellular and Molecular Biology | Payel Mukherjee, Sofia Banu, Priya Singh, Dhiviya Vedagiri, Divya Gupta, Vishal Sah, Santosh Kumar Kuncha, Krishnan Harinivas Harshan, Archana Bharadwaj Siva, Karthik Bharadwaj Tallapaka, Shagufta Khan, Lamuk Zaveri, Namami Gaur, Sakshi Shambhavi, Tulasi Nagabandi, Purushotham Vodnala,G. Aditya Kumar, Koushick Sivakumar, Pooja Ramesh Gupta, Rajan Kumar Jha, Shraddha Vijay Lahoti, Dee |
| hCoV-19/India/GBRC94/2020 | EPI_ISL_450781 | 5/9/2020 | Government Medical College-Bhavnagar | Gujarat Biotechnology Research Centre | Kairavi Desai, Saklain Malek, Shirish Patel, Ramesh Pandit, Tejas Shah, Ankit Hinsu, Pritesh Sabara, Apurvasinh Puvar, Janvi Raval, Zarna Patel, Monika Gandhi, Pinal Trivedi, Maharshi Pandya, Amit Kanani, Nidhi Patel, Nitin Savaliya, Raghawendra Kumar, Dinesh Kumar, Zuber Saiyed, Komal Patel, Labdhi Pandya, Snehal Bagatharia, Bhavesh Modi, Gaurishankar Shrimali, R D Dixit, A M Kadri, Akanksha Ver |
| hCoV-19/India/GBRC95a/2020 | EPI_ISL_450782 | 4/11/2020 | Government Medical College-Bhavnagar | Gujarat Biotechnology Research Centre | Saklain Malek, Shirish Patel, Kairavi Desai, Tejas Shah, Ankit Hinsu, Pritesh Sabara, Apurvasinh Puvar, Janvi Raval, Zarna Patel, Monika Gandhi, Pinal Trivedi, Maharshi Pandya, Amit Kanani, Nidhi Patel, Nitin Savaliya, Raghawendra Kumar, Dinesh Kumar, Zuber Saiyed, Komal Patel, Labdhi Pandya, Snehal Bagatharia, Ramesh Pandit, Bhavesh Modi, Gaurishankar Shrimali, R D Dixit, A M Kadri, Priti Pandita, G |

| | | | | | |
|---|---|---|---|---|---|
| hCoV-19/India/GBRC95b/2020 | EPI_ISL_450783 | 4/11/2020 | Government Medical College-Bhavnagar | Gujarat Biotechnology Research Centre | Shirish Patel, Kairavi Desai, Saklain Malek, Ankit Hinsu, Pritesh Sabara, Apurvasinh Puvar, Janvi Raval, Zarna Patel, Monika Gandhi, Pinal Trivedi, Maharshi Pandya, Amit Kanani, Nidhi Patel, Nitin Savaliya, Raghawendra Kumar, Dinesh Kumar, Zuber Saiyed, Komal Patel, Labdhi Pandya, Snehal Bagatharia, Ramesh Pandit, Tejas Shah, Bhavesh Modi, Gaurishankar Shrimali, R D Dixit, A M Kadri, Neha Rajpara, |
| hCoV-19/India/GBRC96/2020 | EPI_ISL_450784 | 5/9/2020 | Government Medical College-Bhavnagar | Gujarat Biotechnology Research Centre | Zarna Patel, Ramesh Pandit, Tejas Shah, Ankit Hinsu, Pritesh Sabara, Apurvasinh Puvar, Janvi Raval, Monika Gandhi, Pinal Trivedi, Maharshi Pandya, Amit Kanani, Nidhi Patel, Nitin Savaliya, Raghawendra Kumar, Dinesh Kumar, Zuber Saiyed, Komal Patel, Labdhi Pandya, Snehal Bagatharia, Kairavi Desai, Saklain Malek, Shirish Patel, Bhavesh Modi, Gaurishankar Shrimali, R D Dixit, A M Kadri, Afzal Ansari, C |
| hCoV-19/India/GBRC97a/2020 | EPI_ISL_450785 | 5/10/2020 | Pandit Deendayal Upadhyay Government Medical College, Rajkot | Gujarat Biotechnology Research Centre | Prakash Modi, Sejul Antala, Manish Pattani, Apurvasinh Puvar, Janvi Raval, Zarna Patel, Monika Gandhi, Pinal Trivedi, Maharshi Pandya, Amit Kanani, Nidhi Patel, Nitin Savaliya, Raghawendra Kumar, Dinesh Kumar, Zuber Saiyed, Komal Patel, Labdhi Pandya, Snehal Bagatharia, Ramesh Pandit, Tejas Shah, Ankit Hinsu, Pritesh Sabara, Bhavesh Modi, Gaurishankar Shrimali, R D Dixit, A M Kadri, Neelam Natha |
| hCoV-19/India/GBRC97b/2020 | EPI_ISL_450786 | 5/10/2020 | Pandit Deendayal Upadhyay Government Medical College, Rajkot | Gujarat Biotechnology Research Centre | Sejul Antala, Manish Pattani, Prakash Modi, Janvi Raval, Zarna Patel, Monika Gandhi, Pinal Trivedi, Maharshi Pandya, Amit Kanani, Nidhi Patel, Nitin Savaliya, Raghawendra Kumar, Dinesh Kumar, Zuber Saiyed, Komal Patel, Labdhi Pandya, Snehal Bagatharia, Ramesh Pandit, Tejas Shah, Ankit Hinsu, Pritesh Sabara, Apurvasinh Puvar, Bhavesh Modi, Gaurishankar Shrimali, R D Dixit, A M Kadri, Armi Chaudha |
| hCoV-19/India/GBRC98/2020 | EPI_ISL_450787 | 5/11/2020 | Pandit Deendayal Upadhyay Government Medical College, Rajkot | Gujarat Biotechnology Research Centre | Manish Pattani, Prakash Modi, Sejul Antala, Zarna Patel, Monika Gandhi, Pinal Trivedi, Maharshi Pandya, Amit Kanani, Nidhi Patel, Nitin Savaliya, Raghawendra Kumar, Dinesh Kumar, Zuber Saiyed, Komal Patel, Labdhi Pandya, Snehal Bagatharia, Ramesh Pandit, Tejas Shah, Ankit Hinsu, Pritesh Sabara, Apurvasinh Puvar, Janvi Raval, Bhavesh Modi, Gaurishankar Shrimali, R D Dixit, A M Kadri, Bhavya Jinda |
| hCoV-19/India/GBRC99/2020 | EPI_ISL_450788 | 5/6/2020 | Pandit Deendayal Upadhyay Government Medical College, Rajkot | Gujarat Biotechnology Research Centre | Zarna Patel, Tejas Shah, Ankit Hinsu, Pritesh Sabara, Apurvasinh Puvar, Janvi Raval, Monika Gandhi, Pinal Trivedi, Maharshi Pandya, Amit Kanani, Nidhi Patel, Nitin Savaliya, Raghawendra Kumar, Dinesh Kumar, Zuber Saiyed, Komal Patel, Labdhi Pandya, Snehal Bagatharia, Prakash Modi, Sejul Antala, Manish Pattani, Ramesh Pandit, Bhavesh Modi, Gaurishankar Shrimali, R D Dixit, A M Kadri, Camellia Cha |
| hCoV-19/India/GBRC100/2020 | EPI_ISL_450789 | 5/10/2020 | Pandit Deendayal Upadhyay Government Medical College, Rajkot | Gujarat Biotechnology Research Centre | Ankit Hinsu, Pritesh Sabara, Apurvasinh Puvar, Janvi Raval, Zarna Patel, Monika Gandhi, Pinal Trivedi, Maharshi Pandya, Amit Kanani, Nidhi Patel, Nitin Savaliya, Raghawendra Kumar, Dinesh Kumar, Zuber Saiyed, Komal Patel, Labdhi Pandya, Snehal Bagatharia, Prakash Modi, Sejul Antala, Manish Pattani, Ramesh Pandit, Tejas Shah, Bhavesh Modi, Gaurishankar Shrimali, R D Dixit, A M Kadri, Siddhant Kum |
| hCoV-19/India/GBRC101/2020 | EPI_ISL_450790 | 4/28/2020 | Pandit Deendayal Upadhyay Government Medical College, Rajkot | Gujarat Biotechnology Research Centre | Zarna Patel, Pritesh Sabara, Apurvasinh Puvar, Janvi Raval, Monika Gandhi, Pinal Trivedi, Maharshi Pandya, Amit Kanani, Nidhi Patel, Nitin Savaliya, Raghawendra Kumar, Dinesh Kumar, Zuber Saiyed, Komal Patel, Labdhi Pandya, Snehal Bagatharia, Prakash Modi, Sejul Antala, Manish Pattani, Ramesh Pandit, Tejas Shah, Ankit Hinsu, Bhavesh Modi, Gaurishankar Shrimali, R D Dixit, A M Kadri, Sharmistha M |
| hCoV-19/India/GBRC102/2020 | EPI_ISL_450791 | 4/27/2020 | Pandit Deendayal Upadhyay Government Medical College, Rajkot | Gujarat Biotechnology Research Centre | Zarna Patel, Apurvasinh Puvar, Janvi Raval, Monika Gandhi, Pinal Trivedi, Maharshi Pandya, Amit Kanani, Nidhi Patel, Nitin Savaliya, Raghawendra Kumar, Dinesh Kumar, Zuber Saiyed, Komal Patel, Labdhi Pandya, Snehal Bagatharia, Prakash Modi, Sejul Antala, Manish Pattani, Ramesh Pandit, Tejas Shah, Ankit Hinsu, Pritesh Sabara, Bhavesh Modi, Gaurishankar Shrimali, R D Dixit, A M Kadri, Pooja P Dosh |
| hCoV-19/India/GBRC104/2020 | EPI_ISL_451149 | 5/7/2020 | M.P Shah Government Medocal college Jamnagar | Gujarat Biotechnology Research Centre | Janvi Raval, Zarna Patel, Monika Gandhi, Pinal Trivedi, Maharshi Pandya, Amit Kanani, Nidhi Patel, Nitin Savaliya, Raghawendra Kumar, Dinesh Kumar, Zuber Saiyed, Komal Patel, Labdhi Pandya, Snehal Bagatharia, Ramesh Pandit, Tejas Shah, Ankit Hinsu, Pritesh Sabara, Apurvasinh Puvar, Binita Aring, Bhavesh Modi, Gaurishankar Shrimali, R D Dixit, A M Kadri, Priti Pandita, Chaitanya Joshi, Madhvi Joshi, |
| hCoV-19/India/GBRC105/2020 | EPI_ISL_451150 | 5/6/2020 | M.P Shah Government Medocal college Jamnagar | Gujarat Biotechnology Research Centre | Zarna Patel, Monika Gandhi, Pinal Trivedi, Maharshi Pandya, Amit Kanani, Nidhi Patel, Nitin Savaliya, Raghawendra Kumar, Dinesh Kumar, Zuber Saiyed, Komal Patel, Labdhi Pandya, Snehal Bagatharia, Ramesh Pandit, Tejas Shah, Ankit Hinsu, Pritesh Sabara, Apurvasinh Puvar, Binita Aring, Janvi Raval, Bhavesh Modi, Gaurishankar Shrimali, R D Dixit, A M Kadri, Pragya Sharma, Chaitanya Joshi, Madhvi Jos |
| hCoV-19/India/GBRC106/2020 | EPI_ISL_451151 | 5/5/2020 | M.P Shah Government Medocal college Jamnagar | Gujarat Biotechnology Research Centre | Monika Gandhi, Pinal Trivedi, Maharshi Pandya, Amit Kanani, Nidhi Patel, Nitin Savaliya, Raghawendra Kumar, Dinesh Kumar, Zuber Saiyed, Komal Patel, Labdhi Pandya, Snehal Bagatharia, Ramesh Pandit, Tejas Shah, Ankit Hinsu, Pritesh Sabara, Apurvasinh Puvar, Binita Aring, Janvi Raval, Zarna Patel, Bhavesh Modi, Gaurishankar Shrimali, R D Dixit, A M Kadri, Neha Rajpara, Chaitanya Joshi, Madhvi Joshi |
| hCoV-19/India/GBRC107/2020 | EPI_ISL_451152 | 5/9/2020 | M.P Shah Government Medocal college Jamnagar | Gujarat Biotechnology Research Centre | Pinal Trivedi, Maharshi Pandya, Amit Kanani, Nidhi Patel, Nitin Savaliya, Raghawendra Kumar, Dinesh Kumar, Zuber Saiyed, Komal Patel, Labdhi Pandya, Snehal Bagatharia, Ramesh Pandit, Tejas Shah, Ankit Hinsu, Pritesh Sabara, Apurvasinh Puvar, Binita Aring, Janvi Raval, Zarna Patel, Monika Gandhi, Bhavesh Modi, Gaurishankar Shrimali, R D Dixit, A M Kadri, Afzal Ansari, Chaitanya Joshi, Madhvi Joshi, |
| hCoV-19/India/GBRC108/2020 | EPI_ISL_451153 | 5/10/2020 | M.P Shah Government Medocal college Jamnagar | Gujarat Biotechnology Research Centre | Maharshi Pandya, Amit Kanani, Nidhi Patel, Nitin Savaliya, Raghawendra Kumar, Dinesh Kumar, Zuber Saiyed, Komal Patel, Labdhi Pandya, Snehal Bagatharia, Ramesh Pandit, Tejas Shah, Ankit Hinsu, Pritesh Sabara, Apurvasinh Puvar, Binita Aring, Janvi Raval, Zarna Patel, Monika Gandhi, Pinal Trivedi, Bhavesh Modi, Gaurishankar Shrimali, R D Dixit, A M Kadri, Fenil Patel, Chaitanya Joshi, Madhvi Joshi, |
| hCoV-19/India/GBRC109/2020 | EPI_ISL_451154 | 5/3/2020 | Government Medical College, Vadodara | Gujarat Biotechnology Research Centre | Manish Pattani, Tanuja Javadekar , Maharshi Pandya, Amit Kanani, Nidhi Patel, Nitin Savaliya, Raghawendra Kumar, Dinesh Kumar, Zuber Saiyed, Komal Patel, Labdhi Pandya, Snehal Bagatharia, Ramesh Pandit, Tejas Shah, Ankit Hinsu, Pritesh Sabara, Apurvasinh Puvar, Janvi Raval, Zarna Patel, Monika Gandhi, Pinal Trivedi, Bhavesh Modi, Gaurishankar Shrimali, R D Dixit, A M Kadri, Neelam Nathani, Chai |
| hCoV-19/India/GBRC110/2020 | EPI_ISL_451155 | 5/3/2020 | Government Medical College, Vadodara | Gujarat Biotechnology Research Centre | Tanuja Javadekar , Manish Pattani, Amit Kanani, Nidhi Patel, Nitin Savaliya, Raghawendra Kumar, Dinesh Kumar, Zuber Saiyed, Komal Patel, Labdhi Pandya, Snehal Bagatharia, Ramesh Pandit, Tejas Shah, Ankit Hinsu, Pritesh Sabara, Apurvasinh Puvar, Janvi Raval, Zarna Patel, Monika Gandhi, Pinal Trivedi, Maharshi Pandya, Bhavesh Modi, Gaurishankar Shrimali, R D Dixit, A M Kadri, Armi Chaudhari, Chait |
| hCoV-19/India/GBRC111/2020 | EPI_ISL_451156 | 5/1/2020 | Government Medical College, Vadodara | Gujarat Biotechnology Research Centre | Amit Kanani, Nidhi Patel, Nitin Savaliya, Raghawendra Kumar, Dinesh Kumar, Zuber Saiyed, Komal Patel, Labdhi Pandya, Snehal Bagatharia, Ramesh Pandit, Tejas Shah, Ankit Hinsu, Pritesh Sabara, Apurvasinh Puvar, Janvi Raval, Zarna Patel, Monika Gandhi, Pinal Trivedi, Maharshi Pandya, Manish Pattani, Tanuja Javadekar , Bhavesh Modi, Gaurishankar Shrimali, R D Dixit, A M Kadri, Bhavya Jindal, Chaitar |
| hCoV-19/India/GBRC112/2020 | EPI_ISL_451157 | 5/2/2020 | Government Medical College, Vadodara | Gujarat Biotechnology Research Centre | Nidhi Patel, Nitin Savaliya, Raghawendra Kumar, Dinesh Kumar, Zuber Saiyed, Komal Patel, Labdhi Pandya, Snehal Bagatharia, Ramesh Pandit, Tejas Shah, Ankit Hinsu, Pritesh Sabara, Apurvasinh Puvar, Janvi Raval, Zarna Patel, Monika Gandhi, Pinal Trivedi, Maharshi Pandya, Manish Pattani, Tanuja Javadekar , Amit Kanani, Bhavesh Modi, Gaurishankar Shrimali, R D Dixit, A M Kadri, Camellia Chakraborty, |
| hCoV-19/India/GBRC113/2020 | EPI_ISL_451158 | 5/3/2020 | Government Medical College, Vadodara | Gujarat Biotechnology Research Centre | Nitin Savaliya, Raghawendra Kumar, Dinesh Kumar, Zuber Saiyed, Komal Patel, Labdhi Pandya, Snehal Bagatharia, Ramesh Pandit, Tejas Shah, Ankit Hinsu, Pritesh Sabara, Apurvasinh Puvar, Janvi Raval, Zarna Patel, Monika Gandhi, Pinal Trivedi, Maharshi Pandya, Manish Pattani, Tanuja Javadekar , Amit Kanani, Nidhi Patel, Bhavesh Modi, Gaurishankar Shrimali, R D Dixit, A M Kadri, Siddhant Kumar, Chai |
| hCoV-19/India/GBRC114/2020 | EPI_ISL_451159 | 5/3/2020 | Government Medical College, Vadodara | Gujarat Biotechnology Research Centre | Raghawendra Kumar, Dinesh Kumar, Zuber Saiyed, Komal Patel, Labdhi Pandya, Snehal Bagatharia, Ramesh Pandit, Tejas Shah, Ankit Hinsu, Pritesh Sabara, Apurvasinh Puvar, Janvi Raval, Zarna Patel, Monika Gandhi, Pinal Trivedi, Maharshi Pandya, Manish Pattani, Tanuja Javadekar , Amit Kanani, Nidhi Patel, Nitin Savaliya, Bhavesh Modi, Gaurishankar Shrimali, R D Dixit, A M Kadri, Sharmistha Majumdar |
| hCoV-19/India/GBRC115/2020 | EPI_ISL_451160 | 5/3/2020 | Government Medical College, Vadodara | Gujarat Biotechnology Research Centre | Dinesh Kumar, Zuber Saiyed, Komal Patel, Labdhi Pandya, Snehal Bagatharia, Ramesh Pandit, Tejas Shah, Ankit Hinsu, Pritesh Sabara, Apurvasinh Puvar, Janvi Raval, Zarna Patel, Monika Gandhi, Pinal Trivedi, Maharshi Pandya, Manish Pattani, Tanuja Javadekar , Amit Kanani, Nidhi Patel, Nitin Savaliya, Raghawendra Kumar, Bhavesh Modi, Gaurishankar Shrimali, R D Dixit, A M Kadri, Pooja P Doshi, Chaita |
| hCoV-19/India/GBRC116/2020 | EPI_ISL_451161 | 5/3/2020 | Government Medical College, Vadodara | Gujarat Biotechnology Research Centre | Zuber Saiyed, Komal Patel, Labdhi Pandya, Snehal Bagatharia, Ramesh Pandit, Tejas Shah, Ankit Hinsu, Pritesh Sabara, Apurvasinh Puvar, Janvi Raval, Zarna Patel, Monika Gandhi, Pinal Trivedi, Maharshi Pandya, Manish Pattani, Tanuja Javadekar , Amit Kanani, Nidhi Patel, Nitin Savaliya, Raghawendra Kumar, Dinesh Kumar, Bhavesh Modi, Gaurishankar Shrimali, R D Dixit, A M Kadri, Akanksha Verma, Cha |
| hCoV-19/India/GBRC117/2020 | EPI_ISL_451162 | 5/3/2020 | Government Medical College, Vadodara | Gujarat Biotechnology Research Centre | Komal Patel, Labdhi Pandya, Snehal Bagatharia, Ramesh Pandit, Tejas Shah, Ankit Hinsu, Pritesh Sabara, Apurvasinh Puvar, Janvi Raval, Zarna Patel, Monika Gandhi, Pinal Trivedi, Maharshi Pandya, Manish Pattani, Tanuja Javadekar , Amit Kanani, Nidhi Patel, Nitin Savaliya, Raghawendra Kumar, Dinesh Kumar, Zuber Saiyed, Bhavesh Modi, Gaurishankar Shrimali, R D Dixit, A M Kadri, Priti Pandita, Chaitany |
| hCoV-19/India/GBRC118/2020 | EPI_ISL_451163 | 5/2/2020 | Government Medical College, Vadodara | Gujarat Biotechnology Research Centre | Labdhi Pandya, Snehal Bagatharia, Ramesh Pandit, Tejas Shah, Ankit Hinsu, Pritesh Sabara, Apurvasinh Puvar, Janvi Raval, Zarna Patel, Monika Gandhi, Pinal Trivedi, Maharshi Pandya, Manish Pattani, Tanuja Javadekar , Amit Kanani, Nidhi Patel, Nitin Savaliya, Raghawendra Kumar, Dinesh Kumar, Zuber Saiyed, Komal Patel, Bhavesh Modi, Gaurishankar Shrimali, R D Dixit, A M Kadri, Pragya Sharma, Chait |
| hCoV-19/India/GBRC103/2020 | EPI_ISL_451666 | 5/4/2020 | M.P Shah Government Medocal college Jamnagar | Gujarat Biotechnology Research Centre | Binita Aring, Janvi Raval, Zarna Patel, Monika Gandhi, Pinal Trivedi, Maharshi Pandya, Amit Kanani, Nidhi Patel, Nitin Savaliya, Raghawendra Kumar, Dinesh Kumar, Zuber Saiyed, Komal Patel, Labdhi Pandya, Snehal Bagatharia, Ramesh Pandit, Tejas Shah, Ankit Hinsu, Pritesh Sabara, Apurvasinh Puvar, Bhavesh Modi, Gaurishankar Shrimali, R D Dixit, A M Kadri, Akanksha Verma, Chaitanya Joshi, Madhvi Jo |
| hCoV-19/India/NIV-10380/2020 | EPI_ISL_452192 | 4/16/2020 | NIV Influenza | NIV Influenza | Potdar V |
| hCoV-19/India/NIV-10419/2020 | EPI_ISL_452193 | 4/16/2020 | NIV Influenza | NIV Influenza | Potdar V |
| hCoV-19/India/NIV-11677/2020 | EPI_ISL_452194 | 4/19/2020 | NIV Influenza | NIV Influenza | Potdar V |
| hCoV-19/India/NIV-11681/2020 | EPI_ISL_452195 | 4/19/2020 | NIV Influenza | NIV Influenza | Potdar V |
| hCoV-19/India/NIV-11683/2020 | EPI_ISL_452196 | 4/19/2020 | NIV Influenza | NIV Influenza | Potdar V |
| hCoV-19/India/NIV-11687/2020 | EPI_ISL_452197 | 4/19/2020 | NIV Influenza | NIV Influenza | Potdar V |
| hCoV-19/India/NIV-11855/2020 | EPI_ISL_452198 | 4/20/2020 | NIV Influenza | NIV Influenza | Potdar V |
| hCoV-19/India/NIV-12019/2020 | EPI_ISL_452199 | 4/20/2020 | NIV Influenza | NIV Influenza | Potdar V |
| hCoV-19/India/NIV-12024/2020 | EPI_ISL_452200 | 4/20/2020 | NIV Influenza | NIV Influenza | Potdar V |
| hCoV-19/India/NIV-12067/2020 | EPI_ISL_452201 | 4/20/2020 | NIV Influenza | NIV Influenza | Potdar V |
| hCoV-19/India/NIV-2810/2020 | EPI_ISL_452202 | 3/16/2020 | NIV Influenza | NIV Influenza | Potdar V |
| hCoV-19/India/NIV-4367/2020 | EPI_ISL_452203 | 3/22/2020 | NIV Influenza | NIV Influenza | Potdar V |
| hCoV-19/India/NIV-5109/2020 | EPI_ISL_452204 | 3/26/2020 | NIV Influenza | NIV Influenza | Potdar V |
| hCoV-19/India/NIV-5119/2020 | EPI_ISL_452205 | 3/26/2020 | NIV Influenza | NIV Influenza | Potdar V |
| hCoV-19/India/NIV-6311/2020 | EPI_ISL_452206 | 3/30/2020 | NIV Influenza | NIV Influenza | Potdar V |
| hCoV-19/India/NIV-7439/2020 | EPI_ISL_452207 | 4/5/2020 | NIV Influenza | NIV Influenza | Potdar V |
| hCoV-19/India/NIV-7511/2020 | EPI_ISL_452208 | 4/5/2020 | NIV Influenza | NIV Influenza | Potdar V |
| hCoV-19/India/NIV-7786/2020 | EPI_ISL_452209 | 4/6/2020 | NIV Influenza | NIV Influenza | Potdar V |
| hCoV-19/India/NIV-7830/2020 | EPI_ISL_452210 | 4/6/2020 | NIV Influenza | NIV Influenza | Potdar V |
| hCoV-19/India/NIV-9157/2020 | EPI_ISL_452211 | 4/13/2020 | NIV Influenza | NIV Influenza | Potdar V |
| hCoV-19/India/NIV-9768/2020 | EPI_ISL_452212 | 4/15/2020 | NIV Influenza | NIV Influenza | Potdar V |
| hCoV-19/India/NIV-981/2020 | EPI_ISL_452213 | 3/10/2020 | NIV Influenza | NIV Influenza | Potdar V |
| hCoV-19/India/NIV-QA-709/2020 | EPI_ISL_452214 | 4/7/2020 | NIV Influenza | NIV Influenza | Potdar V |
| hCoV-19/India/NIV-QC-797/2020 | EPI_ISL_452215 | 4/22/2020 | NIV Influenza | NIV Influenza | Potdar V |
| hCoV-19/India/NIV-QC-802/2020 | EPI_ISL_452216 | 4/27/2020 | NIV Influenza | NIV Influenza | Potdar V |
| hCoV-19/India/NIV-QC-800/2020 | EPI_ISL_452217 | 4/26/2020 | NIV Influenza | NIV Influenza | Potdar V |
| hCoV-19/India/NIHSAD-0905-216/2020 | EPI_ISL_452794 | 5/8/2020 | ICAR-National Institute of High Security Animal Diseases | ICAR-National Institute of High Security Animal Diseases | Anamika Mishra, Ashutosh Aasdev, Sandeep Bhatia, Harshad Murugkar, Chakradhar Tosh, Niranjan Mishra, Shanmugasundaram Nagarajan, Katherukamem Rajukumar, Richa Sood, G Venkatesh, Atul Kumar Pateriya, Manoj Kumar, Shashi Bhushan Sudhakar, Fateh Singh, Sethil Kumar D, Senmannan Kalaiyarasu, Pradeep Gandhale, Naveen Kumar, Chandan Kumar Dubey, Sushil Tripathi, Sandeep Kumar Jh |
| hCoV-19/India/NIHSAD-1005-52/2020 | EPI_ISL_452795 | 5/9/2020 | ICAR-National Institute of High Security Animal Diseases | ICAR-National Institute of High Security Animal Diseases | Anamika Mishra, Ashutosh Aasdev, Sandeep Bhatia, Harshad Murugkar, Chakradhar Tosh, Niranjan Mishra, Shanmugasundaram Nagarajan, Katherukamem Rajukumar, Richa Sood, G Venkatesh, Atul Kumar Pateriya, Manoj Kumar, Shashi Bhushan Sudhakar, Fateh Singh, Sethil Kumar D, Senmannan Kalaiyarasu, Pradeep Gandhale, Naveen Kumar, Chandan Kumar Dubey, Sushil Tripathi, Sandeep Kumar Jh |
| hCoV-19/India/NIV-12058/2020 | EPI_ISL_454524 | 4/20/2020 | NIV Influenza | NIV Influenza | Potdar V |
| hCoV-19/India/NIV-1738/2020 | EPI_ISL_454525 | 3/12/2020 | NIV Influenza | NIV Influenza | Potdar V |
| hCoV-19/India/NIV-1804/2020 | EPI_ISL_454526 | 3/13/2020 | NIV Influenza | NIV Influenza | Potdar V |
| hCoV-19/India/NIV-2387/2020 | EPI_ISL_454527 | 3/13/2020 | NIV Influenza | NIV Influenza | Potdar V |
| hCoV-19/India/NIV-3472/2020 | EPI_ISL_454528 | 3/17/2020 | NIV Influenza | NIV Influenza | Potdar V |
| hCoV-19/India/NIV-4130/2020 | EPI_ISL_454529 | 3/21/2020 | NIV Influenza | NIV Influenza | Potdar V |
| hCoV-19/India/NIV-4271/2020 | EPI_ISL_454530 | 3/22/2020 | NIV Influenza | NIV Influenza | Potdar V |
| hCoV-19/India/NIV-4369/2020 | EPI_ISL_454531 | 3/22/2020 | NIV Influenza | NIV Influenza | Potdar V |
| hCoV-19/India/NIV-4379/2020 | EPI_ISL_454532 | 3/23/2020 | NIV Influenza | NIV Influenza | Potdar V |
| hCoV-19/India/NIV-4438/2020 | EPI_ISL_454533 | 3/23/2020 | NIV Influenza | NIV Influenza | Potdar V |
| hCoV-19/India/NIV-5102/2020 | EPI_ISL_454536 | 3/26/2020 | NIV Influenza | NIV Influenza | Potdar V |
| hCoV-19/India/NIV-5104/2020 | EPI_ISL_454537 | 3/26/2020 | NIV Influenza | NIV Influenza | Potdar V |
| hCoV-19/India/NIV-6216/2020 | EPI_ISL_454540 | 4/1/2020 | NIV Influenza | NIV Influenza | Potdar V |
| hCoV-19/India/NIV-6315/2020 | EPI_ISL_454542 | 3/30/2020 | NIV Influenza | NIV Influenza | Potdar V |
| hCoV-19/India/NIV-6423/2020 | EPI_ISL_454543 | 3/31/2020 | NIV Influenza | NIV Influenza | Potdar V |
| hCoV-19/India/NIV-7044/2020 | EPI_ISL_454547 | 4/4/2020 | NIV Influenza | NIV Influenza | Potdar V |
| hCoV-19/India/NIV-7558/2020 | EPI_ISL_454549 | 4/5/2020 | NIV Influenza | NIV Influenza | Potdar V |
| hCoV-19/India/NIV-7893/2020 | EPI_ISL_454551 | 4/7/2020 | NIV Influenza | NIV Influenza | Potdar V |
| hCoV-19/India/NIV-9127/2020 | EPI_ISL_454556 | 4/12/2020 | NIV Influenza | NIV Influenza | Potdar V |
| hCoV-19/India/NIV-9154/2020 | EPI_ISL_454557 | 4/13/2020 | NIV Influenza | NIV Influenza | Potdar V |
| hCoV-19/India/NIV-701_3/2020 | EPI_ISL_454560 | 4/6/2020 | NIV Influenza | NIV Influenza | Potdar V |
| hCoV-19/India/NIV-QA-707/2020 | EPI_ISL_454563 | 4/10/2020 | NIV Influenza | NIV Influenza | Potdar V |
| hCoV-19/India/NIV-QA-708/2020 | EPI_ISL_454564 | 4/4/2020 | NIV Influenza | NIV Influenza | Potdar V |
| hCoV-19/India/NIV-QA-710/2020 | EPI_ISL_454565 | 3/31/2020 | NIV Influenza | NIV Influenza | Potdar V |
| hCoV-19/India/NIV-QC-795/2020 | EPI_ISL_454566 | 4/21/2020 | NIV Influenza | NIV Influenza | Potdar V |
| hCoV-19/India/NIV-QC-796/2020 | EPI_ISL_454567 | 4/22/2020 | NIV Influenza | NIV Influenza | Potdar V |
| hCoV-19/India/NIV-QC-798/2020 | EPI_ISL_454568 | 4/25/2020 | NIV Influenza | NIV Influenza | Potdar V |
| hCoV-19/India/NIV-QC-799/2020 | EPI_ISL_454569 | 4/26/2020 | NIV Influenza | NIV Influenza | Potdar V |
| hCoV-19/India/NIV-QC-801/2020 | EPI_ISL_454570 | 4/26/2020 | NIV Influenza | NIV Influenza | Potdar V |
| hCoV-19/India/SMSCOV109/2020 | EPI_ISL_454830 | 4/23/2020 | SMS Medical College, Jaipur | CSIR Institute of Genomics and Integrative Biology | Sudhir Bhandari, Rahul Bhoyar, Mohammed Imran, Mohit Divakar, Disha Sharma, Anshul Kumar, Bani Jolly, Rahul Sahlot, Abhinav Jain, Paras Sehgal, Gyan Ranjan, Vinod Scaria, Sridhar Sivasubbu, Sandeep K Mathur |
| hCoV-19/India/SMSCOV141/2020 | EPI_ISL_454831 | 4/29/2020 | SMS Medical College, Jaipur | CSIR Institute of Genomics and Integrative Biology | Sudhir Bhandari, Rahul Bhoyar, Mohammed Imran, Mohit Divakar, Disha Sharma, Anshul Kumar, Bani Jolly, Rahul Sahlot, Abhinav Jain, Paras Sehgal, Gyan Ranjan, Vinod Scaria, Sridhar Sivasubbu, Sandeep K Mathur |
| hCoV-19/India/SMSCOV161/2020 | EPI_ISL_454832 | 4/21/2020 | SMS Medical College, Jaipur | CSIR Institute of Genomics and Integrative Biology | Sudhir Bhandari, Rahul Bhoyar, Mohammed Imran, Mohit Divakar, Disha Sharma, Anshul Kumar, Bani Jolly, Rahul Sahlot, Abhinav Jain, Paras Sehgal, Gyan Ranjan, Vinod Scaria, Sridhar Sivasubbu, Sandeep K Mathur |
| hCoV-19/India/SMSCOV175/2020 | EPI_ISL_454833 | 4/27/2020 | SMS Medical College, Jaipur | CSIR Institute of Genomics and Integrative Biology | Sudhir Bhandari, Rahul Bhoyar, Mohammed Imran, Mohit Divakar, Disha Sharma, Anshul Kumar, Bani Jolly, Rahul Sahlot, Abhinav Jain, Paras Sehgal, Gyan Ranjan, Vinod Scaria, Sridhar Sivasubbu, Sandeep K Mathur |
| hCoV-19/India/THSTI-BAL_231/2020 | EPI_ISL_454862 | 4/11/2020 | Translational Health Science and Technology Institute -ESIC medical college and hospital, Faridabad | THSTI Bioassay laboratory | Saurabh Kumar, Jigme Wangchuk, Anil Kumar Pandey, Asim Das, Guruprasad R. Medigeshi |
| hCoV-19/India/THSTI-BAL_914/2020 | EPI_ISL_454866 | 4/16/2020 | Translational Health Science and Technology Institute -ESIC medical college and hospital, Faridabad | THSTI Bioassay laboratory | Saurabh Kumar, Jigme Wangchuk, Anil Kumar Pandey, Asim Das, Guruprasad R. Medigeshi |
| hCoV-19/India/THSTI-BAL_918/2020 | EPI_ISL_454867 | 4/16/2020 | Translational Health Science and Technology Institute -ESIC medical college and hospital, Faridabad | THSTI Bioassay laboratory | Saurabh Kumar, Jigme Wangchuk, Anil Kumar Pandey, Asim Das, Guruprasad R. Medigeshi |
| hCoV-19/India/GBRC119/2020 | EPI_ISL_455016 | 4/27/2020 | Pandit Deendayal Upadhyay Government Medical College, Rajkot | Gujarat Biotechnology Research Centre | Prakash Modi, Sejul Antala, Manish Pattani, Ramesh Pandit, Tejas Shah, Ankit Hinsu, Pritesh Sabara, Apurvasinh Puvar, Janvi Raval, Zarna Patel, Monika Gandhi, Pinal Trivedi, Maharshi Pandya, Amit Kanani, Nidhi Patel, Nitin Savaliya, Raghawendra Kumar, Dinesh Kumar, Zuber Saiyed, Komal Patel, Labdhi Pandya, Snehal Bagatharia, Afzal Ansari, Bhavesh Modi, Gaurishankar Shrimali, R D Dixit, A M Kadri, U |
| hCoV-19/India/GBRC120/2020 | EPI_ISL_455017 | 5/2/2020 | Government Medical College, Vadodara | Gujarat Biotechnology Research Centre | Tanuja Javadekar , R N Daveshwar, Ramesh Pandit, Tejas Shah, Ankit Hinsu, Pritesh Sabara, Apurvasinh Puvar, Janvi Raval, Zarna Patel, Monika Gandhi, Pinal Trivedi, Maharshi Pandya, Amit Kanani, Nidhi Patel, Nitin Savaliya, Raghawendra Kumar, Dinesh Kumar, Zuber Saiyed, Komal Patel, Labdhi Pandya, Snehal Bagatharia, Fenil Patel, Bhavesh Modi, Gaurishankar Shrimali, R D Dixit, A M Kadri, Umang M |
| hCoV-19/India/GBRC121/2020 | EPI_ISL_455018 | 5/2/2020 | Government Medical College, Vadodara | Gujarat Biotechnology Research Centre | R N Daveshwar, Ramesh Pandit, Tejas Shah, Ankit Hinsu, Pritesh Sabara, Apurvasinh Puvar, Janvi Raval, Zarna Patel, Monika Gandhi, Pinal Trivedi, Maharshi Pandya, Amit Kanani, Nidhi Patel, Nitin Savaliya, Raghawendra Kumar, Dinesh Kumar, Zuber Saiyed, Komal Patel, Labdhi Pandya, Snehal Bagatharia, Tanuja Javadekar , Neelam Nathani, Bhavesh Modi, Gaurishankar Shrimali, R D Dixit, A M Kadri, Uma |
| hCoV-19/India/GBRC122/2020 | EPI_ISL_455019 | 5/2/2020 | Government Medical College, Vadodara | Gujarat Biotechnology Research Centre | Ramesh Pandit, Tejas Shah, Ankit Hinsu, Pritesh Sabara, Apurvasinh Puvar, Janvi Raval, Zarna Patel, Monika Gandhi, Pinal Trivedi, Maharshi Pandya, Amit Kanani, Nidhi Patel, Nitin Savaliya, Raghawendra Kumar, Dinesh Kumar, Zuber Saiyed, Komal Patel, Labdhi Pandya, Snehal Bagatharia, Tanuja Javadekar , R N Daveshwar, Armi Chaudhari, Bhavesh Modi, Gaurishankar Shrimali, R D Dixit, A M Kadri, Uma |
| hCoV-19/India/GBRC123/2020 | EPI_ISL_455020 | 5/2/2020 | Government Medical College, Vadodara | Gujarat Biotechnology Research Centre | Tejas Shah, Ankit Hinsu, Pritesh Sabara, Apurvasinh Puvar, Janvi Raval, Zarna Patel, Monika Gandhi, Pinal Trivedi, Maharshi Pandya, Amit Kanani, Nidhi Patel, Nitin Savaliya, Raghawendra Kumar, Dinesh Kumar, Zuber Saiyed, Komal Patel, Labdhi Pandya, Snehal Bagatharia, Tanuja Javadekar , R N Daveshwar, Ramesh Pandit, Bhavya Jindal, Bhavesh Modi, Gaurishankar Shrimali, R D Dixit, A M Kadri, Uman |
| hCoV-19/India/GBRC124/2020 | EPI_ISL_455021 | 5/2/2020 | Government Medical College, Vadodara | Gujarat Biotechnology Research Centre | Ankit Hinsu, Pritesh Sabara, Apurvasinh Puvar, Janvi Raval, Zarna Patel, Monika Gandhi, Pinal Trivedi, Maharshi Pandya, Amit Kanani, Nidhi Patel, Nitin Savaliya, Raghawendra Kumar, Dinesh Kumar, Zuber Saiyed, Komal Patel, Labdhi Pandya, Snehal Bagatharia, Tanuja Javadekar , R N Daveshwar, Ramesh Pandit, Tejas Shah, Camellia Chakraborty, Bhavesh Modi, Gaurishankar Shrimali, R D Dixit, A M Kadr |
| hCoV-19/India/GBRC125/2020 | EPI_ISL_455022 | 5/2/2020 | Government Medical College, Vadodara | Gujarat Biotechnology Research Centre | Pritesh Sabara, Apurvasinh Puvar, Janvi Raval, Zarna Patel, Monika Gandhi, Pinal Trivedi, Maharshi Pandya, Amit Kanani, Nidhi Patel, Nitin Savaliya, Raghawendra Kumar, Dinesh Kumar, Zuber Saiyed, Komal Patel, Labdhi Pandya, Snehal Bagatharia, Tanuja Javadekar , R N Daveshwar, Ramesh Pandit, Tejas Shah, Ankit Hinsu, Siddhant Kumar, Bhavesh Modi, Gaurishankar Shrimali, R D Dixit, A M Kadri, Uma |
| hCoV-19/India/GBRC126/2020 | EPI_ISL_455023 | 5/2/2020 | Government Medical College, Vadodara | Gujarat Biotechnology Research Centre | Apurvasinh Puvar, Janvi Raval, Zarna Patel, Monika Gandhi, Pinal Trivedi, Maharshi Pandya, Amit Kanani, Nidhi Patel, Nitin Savaliya, Raghawendra Kumar, Dinesh Kumar, Zuber Saiyed, Komal Patel, Labdhi Pandya, Snehal Bagatharia, Tanuja Javadekar , R N Daveshwar, Ramesh Pandit, Tejas Shah, Ankit Hinsu, Pritesh Sabara, Priyanka P Vatsa, Bhavesh Modi, Gaurishankar Shrimali, R D Dixit, A M Kadri, Un |
| hCoV-19/India/GBRC127/2020 | EPI_ISL_455024 | 5/2/2020 | Government Medical College, Vadodara | Gujarat Biotechnology Research Centre | Janvi Raval, Zarna Patel, Monika Gandhi, Pinal Trivedi, Maharshi Pandya, Amit Kanani, Nidhi Patel, Nitin Savaliya, Raghawendra Kumar, Dinesh Kumar, Zuber Saiyed, Komal Patel, Labdhi Pandya, Snehal Bagatharia, Tanuja Javadekar , R N Daveshwar, Ramesh Pandit, Tejas Shah, Ankit Hinsu, Pritesh Sabara, Apurvasinh Puvar, Pooja P Doshi, Bhavesh Modi, Gaurishankar Shrimali, R D Dixit, A M Kadri, Uman |
| hCoV-19/India/GBRC128/2020 | EPI_ISL_455025 | 5/2/2020 | Government Medical College, Vadodara | Gujarat Biotechnology Research Centre | Zarna Patel, Monika Gandhi, Pinal Trivedi, Maharshi Pandya, Amit Kanani, Nidhi Patel, Nitin Savaliya, Raghawendra Kumar, Dinesh Kumar, Zuber Saiyed, Komal Patel, Labdhi Pandya, Snehal Bagatharia, Tanuja Javadekar , R N Daveshwar, Ramesh Pandit, Tejas Shah, Ankit Hinsu, Pritesh Sabara, Apurvasinh Puvar, Janvi Raval, Akanksha Verma, Bhavesh Modi, Gaurishankar Shrimali, R D Dixit, A M Kadri, Un |
| hCoV-19/India/GBRC129/2020 | EPI_ISL_455026 | 5/2/2020 | Government Medical College, Vadodara | Gujarat Biotechnology Research Centre | Monika Gandhi, Pinal Trivedi, Maharshi Pandya, Amit Kanani, Nidhi Patel, Nitin Savaliya, Raghawendra Kumar, Dinesh Kumar, Zuber Saiyed, Komal Patel, Labdhi Pandya, Snehal Bagatharia, Tanuja Javadekar , R N Daveshwar, Ramesh Pandit, Tejas Shah, Ankit Hinsu, Pritesh Sabara, Apurvasinh Puvar, Janvi Raval, Zarna Patel, Priti Pandita, Bhavesh Modi, Gaurishankar Shrimali, R D Dixit, A M Kadri, Umang |
| hCoV-19/India/GBRC130/2020 | EPI_ISL_455027 | 5/2/2020 | Government Medical College, Vadodara | Gujarat Biotechnology Research Centre | Pinal Trivedi, Maharshi Pandya, Amit Kanani, Nidhi Patel, Nitin Savaliya, Raghawendra Kumar, Dinesh Kumar, Zuber Saiyed, Komal Patel, Labdhi Pandya, Snehal Bagatharia, Tanuja Javadekar , R N Daveshwar, Ramesh Pandit, Tejas Shah, Ankit Hinsu, Pritesh Sabara, Apurvasinh Puvar, Janvi Raval, Zarna Patel, Monika Gandhi, Pragya Sharma, Bhavesh Modi, Gaurishankar Shrimali, R D Dixit, A M Kadri, Uma |
| hCoV-19/India/RMRC49/2020 | EPI_ISL_455478 | 4/9/2020 | REGIONAL VRDL,ICMR-RMRC BBSR | Immunogenomics group, Institute of Life Sciences, Bhubaneswar | Sunil Raghav, Jyotirmayee Turuk, Arup Ghosh, Atimukta Jha, Viplov K. Biswas, Swati Madhulika, Manasi Priyadarshini, Shuchi Smita, Jaya Singh Khastri, Rupesh Dash, Soma Chattopadhyay, Ghulam Hussain Syed, Shanti Senapati, Tushar K. Beuria, Debdutta Bhattacharya, Rajeeb Swain, Punit Prasad, COVID-19 team of ILS & RMRC, Orissa COVID-19 study group, Sanghamitra Pati, Ajay Parida |
| hCoV-19/India/S13/2020 | EPI_ISL_455643 | 3/30/2020 | ICMR-National Institute of Cholera and Enteric Diseases | National Institute of Biomedical Genomics | Arindam Maitra, Mamta Chawla Sarkar, Sreedhar Chinnaswamy, Hasina Banu, Ananya Chatterjee, Shanta Dutta, Saumitra Das |
| hCoV-19/India/S14/2020 | EPI_ISL_455644 | 4/2/2020 | ICMR-National Institute of Cholera and Enteric Diseases | National Institute of Biomedical Genomics | Arindam Maitra, Mamta Chawla Sarkar, Sreedhar Chinnaswamy, Hasina Banu, Ananya Chatterjee, Shanta Dutta, Saumitra Das |
| hCoV-19/India/S15/2020 | EPI_ISL_455645 | 4/3/2020 | ICMR-National Institute of Cholera and Enteric Diseases | National Institute of Biomedical Genomics | Arindam Maitra, Mamta Chawla Sarkar, Sreedhar Chinnaswamy, Hasina Banu, Ananya Chatterjee, Shanta Dutta, Saumitra Das |
| hCoV-19/India/S16/2020 | EPI_ISL_455646 | 4/3/2020 | ICMR-National Institute of Cholera and Enteric Diseases | National Institute of Biomedical Genomics | Arindam Maitra, Mamta Chawla Sarkar, Sreedhar Chinnaswamy, Hasina Banu, Ananya Chatterjee, Shanta Dutta, Saumitra Das |
| hCoV-19/India/S17/2020 | EPI_ISL_455647 | 4/19/2020 | ICMR-National Institute of Cholera and Enteric Diseases | National Institute of Biomedical Genomics | Arindam Maitra, Mamta Chawla Sarkar, Sreedhar Chinnaswamy, Hasina Banu, Ananya Chatterjee, Shanta Dutta, Saumitra Das |
| hCoV-19/India/S18/2020 | EPI_ISL_455648 | 4/20/2020 | ICMR-National Institute of Cholera and Enteric Diseases | National Institute of Biomedical Genomics | Arindam Maitra, Mamta Chawla Sarkar, Sreedhar Chinnaswamy, Hasina Banu, Ananya Chatterjee, Shanta Dutta, Saumitra Das |
| hCoV-19/India/S19/2020 | EPI_ISL_455649 | 4/20/2020 | ICMR-National Institute of Cholera and Enteric Diseases | National Institute of Biomedical Genomics | Arindam Maitra, Mamta Chawla Sarkar, Sreedhar Chinnaswamy, Hasina Banu, Ananya Chatterjee, Shanta Dutta, Saumitra Das |
| hCoV-19/India/S20/2020 | EPI_ISL_455650 | 4/20/2020 | ICMR-National Institute of Cholera and Enteric Diseases | National Institute of Biomedical Genomics | Arindam Maitra, Mamta Chawla Sarkar, Sreedhar Chinnaswamy, Hasina Banu, Ananya Chatterjee, Shanta Dutta, Saumitra Das |
| hCoV-19/India/S21/2020 | EPI_ISL_455651 | 4/21/2020 | ICMR-National Institute of Cholera and Enteric Diseases | National Institute of Biomedical Genomics | Arindam Maitra, Mamta Chawla Sarkar, Sreedhar Chinnaswamy, Hasina Banu, Ananya Chatterjee, Shanta Dutta, Saumitra Das |
| hCoV-19/India/S22/2020 | EPI_ISL_455652 | 4/21/2020 | ICMR-National Institute of Cholera and Enteric Diseases | National Institute of Biomedical Genomics | Arindam Maitra, Mamta Chawla Sarkar, Sreedhar Chinnaswamy, Hasina Banu, Ananya Chatterjee, Shanta Dutta, Saumitra Das |
| hCoV-19/India/S23/2020 | EPI_ISL_455653 | 4/21/2020 | ICMR-National Institute of Cholera and Enteric Diseases | National Institute of Biomedical Genomics | Arindam Maitra, Mamta Chawla Sarkar, Sreedhar Chinnaswamy, Hasina Banu, Ananya Chatterjee, Shanta Dutta, Saumitra Das |
| hCoV-19/India/S25/2020 | EPI_ISL_455655 | 4/30/2020 | ICMR-National Institute of Cholera and Enteric Diseases | National Institute of Biomedical Genomics | Arindam Maitra, Mamta Chawla Sarkar, Sreedhar Chinnaswamy, Hasina Banu, Ananya Chatterjee, Shanta Dutta, Saumitra Das |
| hCoV-19/India/S27/2020 | EPI_ISL_455656 | 4/30/2020 | ICMR-National Institute of Cholera and Enteric Diseases | National Institute of Biomedical Genomics | Arindam Maitra, Mamta Chawla Sarkar, Sreedhar Chinnaswamy, Hasina Banu, Ananya Chatterjee, Shanta Dutta, Saumitra Das |
| hCoV-19/India/S29/2020 | EPI_ISL_455658 | 4/30/2020 | ICMR-National Institute of Cholera and Enteric Diseases | National Institute of Biomedical Genomics | Arindam Maitra, Mamta Chawla Sarkar, Sreedhar Chinnaswamy, Hasina Banu, Ananya Chatterjee, Shanta Dutta, Saumitra Das |
| hCoV-19/India/S30/2020 | EPI_ISL_455659 | 5/1/2020 | ICMR-National Institute of Cholera and Enteric Diseases | National Institute of Biomedical Genomics | Arindam Maitra, Mamta Chawla Sarkar, Sreedhar Chinnaswamy, Hasina Banu, Ananya Chatterjee, Shanta Dutta, Saumitra Das |
| hCoV-19/India/S31/2020 | EPI_ISL_455660 | 5/1/2020 | ICMR-National Institute of Cholera and Enteric Diseases | National Institute of Biomedical Genomics | Arindam Maitra, Mamta Chawla Sarkar, Sreedhar Chinnaswamy, Hasina Banu, Ananya Chatterjee, Shanta Dutta, Saumitra Das |
| hCoV-19/India/S33/2020 | EPI_ISL_455662 | 5/2/2020 | ICMR-National Institute of Cholera and Enteric Diseases | National Institute of Biomedical Genomics | Arindam Maitra, Mamta Chawla Sarkar, Sreedhar Chinnaswamy, Hasina Banu, Ananya Chatterjee, Shanta Dutta, Saumitra Das |
| hCoV-19/India/S34/2020 | EPI_ISL_455663 | 5/2/2020 | ICMR-National Institute of Cholera and Enteric Diseases | National Institute of Biomedical Genomics | Arindam Maitra, Mamta Chawla Sarkar, Sreedhar Chinnaswamy, Hasina Banu, Ananya Chatterjee, Shanta Dutta, Saumitra Das |
| hCoV-19/India/S46/2020 | EPI_ISL_455667 | 5/3/2020 | ICMR-National Institute of Cholera and Enteric Diseases | National Institute of Biomedical Genomics | Arindam Maitra, Mamta Chawla Sarkar, Sreedhar Chinnaswamy, Hasina Banu, Ananya Chatterjee, Shanta Dutta, Saumitra Das |
| hCoV-19/India/S48/2020 | EPI_ISL_455668 | 5/5/2020 | ICMR-National Institute of Cholera and Enteric Diseases | National Institute of Biomedical Genomics | Arindam Maitra, Mamta Chawla Sarkar, Sreedhar Chinnaswamy, Hasina Banu, Ananya Chatterjee, Shanta Dutta, Saumitra Das |
| hCoV-19/India/S49/2020 | EPI_ISL_455669 | 5/3/2020 | ICMR-National Institute of Cholera and Enteric Diseases | National Institute of Biomedical Genomics | Arindam Maitra, Mamta Chawla Sarkar, Sreedhar Chinnaswamy, Hasina Banu, Ananya Chatterjee, Shanta Dutta, Saumitra Das |
| hCoV-19/India/S50/2020 | EPI_ISL_455670 | 5/3/2020 | ICMR-National Institute of Cholera and Enteric Diseases | National Institute of Biomedical Genomics | Arindam Maitra, Mamta Chawla Sarkar, Sreedhar Chinnaswamy, Hasina Banu, Ananya Chatterjee, Shanta Dutta, Saumitra Das |
| hCoV-19/India/S51/2020 | EPI_ISL_455671 | 5/3/2020 | ICMR-National Institute of Cholera and Enteric Diseases | National Institute of Biomedical Genomics | Arindam Maitra, Mamta Chawla Sarkar, Sreedhar Chinnaswamy, Hasina Banu, Ananya Chatterjee, Shanta Dutta, Saumitra Das |
| hCoV-19/India/S52/2020 | EPI_ISL_455672 | 5/3/2020 | ICMR-National Institute of Cholera and Enteric Diseases | National Institute of Biomedical Genomics | Arindam Maitra, Mamta Chawla Sarkar, Sreedhar Chinnaswamy, Hasina Banu, Ananya Chatterjee, Shanta Dutta, Saumitra Das |
| hCoV-19/India/S53/2020 | EPI_ISL_455673 | 5/2/2020 | ICMR-National Institute of Cholera and Enteric Diseases | National Institute of Biomedical Genomics | Arindam Maitra, Mamta Chawla Sarkar, Sreedhar Chinnaswamy, Hasina Banu, Ananya Chatterjee, Shanta Dutta, Saumitra Das |
| hCoV-19/India/S55/2020 | EPI_ISL_455674 | 5/3/2020 | ICMR-National Institute of Cholera and Enteric Diseases | National Institute of Biomedical Genomics | Arindam Maitra, Mamta Chawla Sarkar, Sreedhar Chinnaswamy, Hasina Banu, Ananya Chatterjee, Shanta Dutta, Saumitra Das |
| hCoV-19/India/S56/2020 | EPI_ISL_455675 | 5/3/2020 | ICMR-National Institute of Cholera and Enteric Diseases | National Institute of Biomedical Genomics | Arindam Maitra, Mamta Chawla Sarkar, Sreedhar Chinnaswamy, Hasina Banu, Ananya Chatterjee, Shanta Dutta, Saumitra Das |
| hCoV-19/India/S58/2020 | EPI_ISL_455676 | 5/3/2020 | ICMR-National Institute of Cholera and Enteric Diseases | National Institute of Biomedical Genomics | Arindam Maitra, Mamta Chawla Sarkar, Sreedhar Chinnaswamy, Hasina Banu, Ananya Chatterjee, Shanta Dutta, Saumitra Das |
| hCoV-19/India/S61/2020 | EPI_ISL_455678 | 4/30/2020 | ICMR-National Institute of Cholera and Enteric Diseases | National Institute of Biomedical Genomics | Arindam Maitra, Mamta Chawla Sarkar, Sreedhar Chinnaswamy, Hasina Banu, Ananya Chatterjee, Shanta Dutta, Saumitra Das |
| hCoV-19/India/S62/2020 | EPI_ISL_455679 | 5/4/2020 | ICMR-National Institute of Cholera and Enteric Diseases | National Institute of Biomedical Genomics | Arindam Maitra, Mamta Chawla Sarkar, Sreedhar Chinnaswamy, Hasina Banu, Ananya Chatterjee, Shanta Dutta, Saumitra Das |
| hCoV-19/India/RMRC103/2020 | EPI_ISL_455749 | 5/6/2020 | REGIONAL VRDL,ICMR-RMRC BBSR | Immunogenomics lab, Institute of Life Sciences, Bhubaneswar | Sunil Raghav, Jyotirmayee Turuk, Arup Ghosh, Atimukta Jha, Viplov K. Biswas, Swati Madhulika, Manasi Priyadarshini, Shuchi Smita, Jaya Singh Khastri, Rupesh Dash, Soma Chattopadhyay, Ghulam Hussain Syed, Shanti Senapati, Tushar K. Beuria, Debdutta Bhattacharya, Rajeeb Swain, Punit Prasad, COVID-19 team of ILS & RMRC, Orissa COVID-19 study group, Sanghamitra Pati, Ajay Parida |
| hCoV-19/India/RMRC112/2020 | EPI_ISL_455754 | 5/6/2020 | REGIONAL VRDL,ICMR-RMRC BBSR | Immunogenomics lab, Institute of Life Sciences, Bhubaneswar | Sunil Raghav, Jyotirmayee Turuk, Arup Ghosh, Atimukta Jha, Viplov K. Biswas, Swati Madhulika, Manasi Priyadarshini, Shuchi Smita, Jaya Singh Khastri, Rupesh Dash, Soma Chattopadhyay, Ghulam Hussain Syed, Shanti Senapati, Tushar K. Beuria, Debdutta Bhattacharya, Rajeeb Swain, Punit Prasad, COVID-19 team of ILS & RMRC, Orissa COVID-19 study group, Sanghamitra Pati, Ajay Parida |
| hCoV-19/India/RMRC154/2020 | EPI_ISL_455755 | 5/7/2020 | REGIONAL VRDL,ICMR-RMRC BBSR | Immunogenomics lab, Institute of Life Sciences, Bhubaneswar | Sunil Raghav, Jyotirmayee Turuk, Arup Ghosh, Atimukta Jha, Viplov K. Biswas, Swati Madhulika, Manasi Priyadarshini, Shuchi Smita, Jaya Singh Khastri, Rupesh Dash, Soma Chattopadhyay, Ghulam Hussain Syed, Shanti Senapati, Tushar K. Beuria, Debdutta Bhattacharya, Rajeeb Swain, Punit Prasad, COVID-19 team of ILS & RMRC, Orissa COVID-19 study group, Sanghamitra Pati, Ajay Parida |
| hCoV-19/India/RMRC160/2020 | EPI_ISL_455761 | 5/7/2020 | REGIONAL VRDL,ICMR-RMRC BBSR | Immunogenomics lab, Institute of Life Sciences, Bhubaneswar | Sunil Raghav, Jyotirmayee Turuk, Arup Ghosh, Atimukta Jha, Viplov K. Biswas, Swati Madhulika, Manasi Priyadarshini, Shuchi Smita, Jaya Singh Khastri, Rupesh Dash, Soma Chattopadhyay, Ghulam Hussain Syed, Shanti Senapati, Tushar K. Beuria, Debdutta Bhattacharya, Rajeeb Swain, Punit Prasad, COVID-19 team of ILS & RMRC, Orissa COVID-19 study group, Sanghamitra Pati, Ajay Parida |
| hCoV-19/India/RMRC163/2020 | EPI_ISL_455763 | 5/7/2020 | REGIONAL VRDL,ICMR-RMRC BBSR | Immunogenomics lab, Institute of Life Sciences, Bhubaneswar | Sunil Raghav, Jyotirmayee Turuk, Arup Ghosh, Atimukta Jha, Viplov K. Biswas, Swati Madhulika, Manasi Priyadarshini, Shuchi Smita, Jaya Singh Khastri, Rupesh Dash, Soma Chattopadhyay, Ghulam Hussain Syed, Shanti Senapati, Tushar K. Beuria, Debdutta Bhattacharya, Rajeeb Swain, Punit Prasad, COVID-19 team of ILS & RMRC, Orissa COVID-19 study group, Sanghamitra Pati, Ajay Parida |
| hCoV-19/India/RMRC164/2020 | EPI_ISL_455764 | 5/7/2020 | REGIONAL VRDL,ICMR-RMRC BBSR | Immunogenomics lab, Institute of Life Sciences, Bhubaneswar | Sunil Raghav, Jyotirmayee Turuk, Arup Ghosh, Atimukta Jha, Viplov K. Biswas, Swati Madhulika, Manasi Priyadarshini, Shuchi Smita, Jaya Singh Khastri, Rupesh Dash, Soma Chattopadhyay, Ghulam Hussain Syed, Shanti Senapati, Tushar K. Beuria, Debdutta Bhattacharya, Rajeeb Swain, Punit Prasad, COVID-19 team of ILS & RMRC, Orissa COVID-19 study group, Sanghamitra Pati, Ajay Parida |
| hCoV-19/India/RMRC165/2020 | EPI_ISL_455765 | 5/7/2020 | REGIONAL VRDL,ICMR-RMRC BBSR | Immunogenomics lab, Institute of Life Sciences, Bhubaneswar | Sunil Raghav, Jyotirmayee Turuk, Arup Ghosh, Atimukta Jha, Viplov K. Biswas, Swati Madhulika, Manasi Priyadarshini, Shuchi Smita, Jaya Singh Khastri, Rupesh Dash, Soma Chattopadhyay, Ghulam Hussain Syed, Shanti Senapati, Tushar K. Beuria, Debdutta Bhattacharya, Rajeeb Swain, Punit Prasad, COVID-19 team of ILS & RMRC, Orissa COVID-19 study group, Sanghamitra Pati, Ajay Parida |
| hCoV-19/India/RMRC166/2020 | EPI_ISL_455766 | 5/7/2020 | REGIONAL VRDL,ICMR-RMRC BBSR | Immunogenomics lab, Institute of Life Sciences, Bhubaneswar | Sunil Raghav, Jyotirmayee Turuk, Arup Ghosh, Atimukta Jha, Viplov K. Biswas, Swati Madhulika, Manasi Priyadarshini, Shuchi Smita, Jaya Singh Khastri, Rupesh Dash, Soma Chattopadhyay, Ghulam Hussain Syed, Shanti Senapati, Tushar K. Beuria, Debdutta Bhattacharya, Rajeeb Swain, Punit Prasad, COVID-19 team of ILS & RMRC, Orissa COVID-19 study group, Sanghamitra Pati, Ajay Parida |
| hCoV-19/India/RMRC167/2020 | EPI_ISL_455767 | 5/7/2020 | REGIONAL VRDL,ICMR-RMRC BBSR | Immunogenomics lab, Institute of Life Sciences, Bhubaneswar | Sunil Raghav, Jyotirmayee Turuk, Arup Ghosh, Atimukta Jha, Viplov K. Biswas, Swati Madhulika, Manasi Priyadarshini, Shuchi Smita, Jaya Singh Khastri, Rupesh Dash, Soma Chattopadhyay, Ghulam Hussain Syed, Shanti Senapati, Tushar K. Beuria, Debdutta Bhattacharya, Rajeeb Swain, Punit Prasad, COVID-19 team of ILS & RMRC, Orissa COVID-19 study group, Sanghamitra Pati, Ajay Parida |

| | | | | | |
|---|---|---|---|---|---|
| hCoV-19/India/RMRC169/2020 | EPI_ISL_455768 | 5/7/2020 | REGIONAL VRDL,ICMR-RMRC BBSR | Immunogenomics lab, Institute of Life Sciences, Bhubaneswar | Sunil Raghav, Jyotirmayee Turuk, Arup Ghosh, Atimukta Jha, Viplov K. Biswas, Swati Madhulika, Manasi Priyadarshini, Shuchi Smita, Jaya Singh Khastri, Rupesh Dash, Soma Chattopadhyay, Ghulam Hussain Syed, Shanti Senapati, Tushar K. Beuria, Debdutta Bhattacharya, Rajeeb Swain, Punit Prasad, COVID-19 team of ILS & RMRC, Orissa COVID-19 study group, Sanghamitra Pati, Ajay Parida |
| hCoV-19/India/RMRC171/2020 | EPI_ISL_455770 | 5/7/2020 | REGIONAL VRDL,ICMR-RMRC BBSR | Immunogenomics lab, Institute of Life Sciences, Bhubaneswar | Sunil Raghav, Jyotirmayee Turuk, Arup Ghosh, Atimukta Jha, Viplov K. Biswas, Swati Madhulika, Manasi Priyadarshini, Shuchi Smita, Jaya Singh Khastri, Rupesh Dash, Soma Chattopadhyay, Ghulam Hussain Syed, Shanti Senapati, Tushar K. Beuria, Debdutta Bhattacharya, Rajeeb Swain, Punit Prasad, COVID-19 team of ILS & RMRC, Orissa COVID-19 study group, Sanghamitra Pati, Ajay Parida |
| hCoV-19/India/RMRC24/2020 | EPI_ISL_455775 | 4/4/2020 | REGIONAL VRDL,ICMR-RMRC BBSR | Immunogenomics lab, Institute of Life Sciences, Bhubaneswar | Sunil Raghav, Jyotirmayee Turuk, Arup Ghosh, Atimukta Jha, Viplov K. Biswas, Swati Madhulika, Manasi Priyadarshini, Shuchi Smita, Jaya Singh Khastri, Rupesh Dash, Soma Chattopadhyay, Ghulam Hussain Syed, Shanti Senapati, Tushar K. Beuria, Debdutta Bhattacharya, Rajeeb Swain, Punit Prasad, COVID-19 team of ILS & RMRC, Orissa COVID-19 study group, Sanghamitra Pati, Ajay Parida |
| hCoV-19/India/RMRC27/2020 | EPI_ISL_455776 | 4/4/2020 | REGIONAL VRDL,ICMR-RMRC BBSR | Immunogenomics lab, Institute of Life Sciences, Bhubaneswar | Sunil Raghav, Jyotirmayee Turuk, Arup Ghosh, Atimukta Jha, Viplov K. Biswas, Swati Madhulika, Manasi Priyadarshini, Shuchi Smita, Jaya Singh Khastri, Rupesh Dash, Soma Chattopadhyay, Ghulam Hussain Syed, Shanti Senapati, Tushar K. Beuria, Debdutta Bhattacharya, Rajeeb Swain, Punit Prasad, COVID-19 team of ILS & RMRC, Orissa COVID-19 study group, Sanghamitra Pati, Ajay Parida |
| hCoV-19/India/RMRC28/2020 | EPI_ISL_455777 | 4/4/2020 | REGIONAL VRDL,ICMR-RMRC BBSR | Immunogenomics lab, Institute of Life Sciences, Bhubaneswar | Sunil Raghav, Jyotirmayee Turuk, Arup Ghosh, Atimukta Jha, Viplov K. Biswas, Swati Madhulika, Manasi Priyadarshini, Shuchi Smita, Jaya Singh Khastri, Rupesh Dash, Soma Chattopadhyay, Ghulam Hussain Syed, Shanti Senapati, Tushar K. Beuria, Debdutta Bhattacharya, Rajeeb Swain, Punit Prasad, COVID-19 team of ILS & RMRC, Orissa COVID-19 study group, Sanghamitra Pati, Ajay Parida |
| hCoV-19/India/RMRC30/2020 | EPI_ISL_455778 | 4/4/2020 | REGIONAL VRDL,ICMR-RMRC BBSR | Immunogenomics lab, Institute of Life Sciences, Bhubaneswar | Sunil Raghav, Jyotirmayee Turuk, Arup Ghosh, Atimukta Jha, Viplov K. Biswas, Swati Madhulika, Manasi Priyadarshini, Shuchi Smita, Jaya Singh Khastri, Rupesh Dash, Soma Chattopadhyay, Ghulam Hussain Syed, Shanti Senapati, Tushar K. Beuria, Debdutta Bhattacharya, Rajeeb Swain, Punit Prasad, COVID-19 team of ILS & RMRC, Orissa COVID-19 study group, Sanghamitra Pati, Ajay Parida |
| hCoV-19/India/RMRC38/2020 | EPI_ISL_455779 | 4/4/2020 | REGIONAL VRDL,ICMR-RMRC BBSR | Immunogenomics lab, Institute of Life Sciences, Bhubaneswar | Sunil Raghav, Jyotirmayee Turuk, Arup Ghosh, Atimukta Jha, Viplov K. Biswas, Swati Madhulika, Manasi Priyadarshini, Shuchi Smita, Jaya Singh Khastri, Rupesh Dash, Soma Chattopadhyay, Ghulam Hussain Syed, Shanti Senapati, Tushar K. Beuria, Debdutta Bhattacharya, Rajeeb Swain, Punit Prasad, COVID-19 team of ILS & RMRC, Orissa COVID-19 study group, Sanghamitra Pati, Ajay Parida |
| hCoV-19/India/RMRC46/2020 | EPI_ISL_455780 | 4/8/2020 | REGIONAL VRDL,ICMR-RMRC BBSR | Immunogenomics lab, Institute of Life Sciences, Bhubaneswar | Sunil Raghav, Jyotirmayee Turuk, Arup Ghosh, Atimukta Jha, Viplov K. Biswas, Swati Madhulika, Manasi Priyadarshini, Shuchi Smita, Jaya Singh Khastri, Rupesh Dash, Soma Chattopadhyay, Ghulam Hussain Syed, Shanti Senapati, Tushar K. Beuria, Debdutta Bhattacharya, Rajeeb Swain, Punit Prasad, COVID-19 team of ILS & RMRC, Orissa COVID-19 study group, Sanghamitra Pati, Ajay Parida |
| hCoV-19/India/RMRC6/2020 | EPI_ISL_455783 | 4/2/2020 | REGIONAL VRDL,ICMR-RMRC BBSR | Immunogenomics lab, Institute of Life Sciences, Bhubaneswar | Sunil Raghav, Jyotirmayee Turuk, Arup Ghosh, Atimukta Jha, Viplov K. Biswas, Swati Madhulika, Manasi Priyadarshini, Shuchi Smita, Jaya Singh Khastri, Rupesh Dash, Soma Chattopadhyay, Ghulam Hussain Syed, Shanti Senapati, Tushar K. Beuria, Debdutta Bhattacharya, Rajeeb Swain, Punit Prasad, COVID-19 team of ILS & RMRC, Orissa COVID-19 study group, Sanghamitra Pati, Ajay Parida |
| hCoV-19/India/RMRC90/2020 | EPI_ISL_455786 | 5/4/2020 | REGIONAL VRDL,ICMR-RMRC BBSR | Immunogenomics lab, Institute of Life Sciences, Bhubaneswar | Sunil Raghav, Jyotirmayee Turuk, Arup Ghosh, Atimukta Jha, Viplov K. Biswas, Swati Madhulika, Manasi Priyadarshini, Shuchi Smita, Jaya Singh Khastri, Rupesh Dash, Soma Chattopadhyay, Ghulam Hussain Syed, Shanti Senapati, Tushar K. Beuria, Debdutta Bhattacharya, Rajeeb Swain, Punit Prasad, COVID-19 team of ILS & RMRC, Orissa COVID-19 study group, Sanghamitra Pati, Ajay Parida |
| hCoV-19/India/RMRC92/2020 | EPI_ISL_455787 | 5/4/2020 | REGIONAL VRDL,ICMR-RMRC BBSR | Immunogenomics lab, Institute of Life Sciences, Bhubaneswar | Sunil Raghav, Jyotirmayee Turuk, Arup Ghosh, Atimukta Jha, Viplov K. Biswas, Swati Madhulika, Manasi Priyadarshini, Shuchi Smita, Jaya Singh Khastri, Rupesh Dash, Soma Chattopadhyay, Ghulam Hussain Syed, Shanti Senapati, Tushar K. Beuria, Debdutta Bhattacharya, Rajeeb Swain, Punit Prasad, COVID-19 team of ILS & RMRC, Orissa COVID-19 study group, Sanghamitra Pati, Ajay Parida |
| hCoV-19/India/CCMB_C12/2020 | EPI_ISL_458030 | 4/26/2020 | King Institute of Preventive Medicine & Research | CSIR-Centre for Cellular and Molecular Biology | K.Kaveri,S.Sivasubramanian,S.Vennila,P.Padmapriya,R.Kiruba,S.Magesh,G. Dhinakar Raj, G. Ravikumar, P. Azhahianambi,K Thangaraj,Payel Mukherjee, Sofia Banu, Priya Singh, Dhiviya Vedagiri, Divya Gupta, Vishal Sah, Santosh Kumar Kuncha, Krishnan Harinivas Harshan, Archana Bharadwaj Siva, Karthik Bharadwaj Tallapaka, Shagufta Khan, Lamuk Zaveri, Namami Gaur, Sakshi Shambhavi, Tulasi Nagab |
| hCoV-19/India/CCMB_C6/2020 | EPI_ISL_458031 | 4/26/2020 | King Institute of Preventive Medicine & Research | CSIR-Centre for Cellular and Molecular Biology | K.Kaveri,S.Sivasubramanian,S.Vennila,P.Padmapriya,R.Kiruba,S.Magesh,G. Dhinakar Raj, G. Ravikumar, P. Azhahianambi,K Thangaraj,Payel Mukherjee, Sofia Banu, Priya Singh, Dhiviya Vedagiri, Divya Gupta, Vishal Sah, Santosh Kumar Kuncha, Krishnan Harinivas Harshan, Archana Bharadwaj Siva, Karthik Bharadwaj Tallapaka, Shagufta Khan, Lamuk Zaveri, Namami Gaur, Sakshi Shambhavi, Tulasi Nagab |
| hCoV-19/India/CCMB_C8/2020 | EPI_ISL_458032 | 4/27/2020 | King Institute of Preventive Medicine & Research | CSIR-Centre for Cellular and Molecular Biology | K.Kaveri,S.Sivasubramanian,S.Vennila,P.Padmapriya,R.Kiruba,S.Magesh,G. Dhinakar Raj, G. Ravikumar, P. Azhahianambi,K Thangaraj,Payel Mukherjee, Sofia Banu, Priya Singh, Dhiviya Vedagiri, Divya Gupta, Vishal Sah, Santosh Kumar Kuncha, Krishnan Harinivas Harshan, Archana Bharadwaj Siva, Karthik Bharadwaj Tallapaka, Shagufta Khan, Lamuk Zaveri, Namami Gaur, Sakshi Shambhavi, Tulasi Nagab |
| hCoV-19/India/CCMB_C5/2020 | EPI_ISL_458033 | 4/29/2020 | King Institute of Preventive Medicine & Research | CSIR-Centre for Cellular and Molecular Biology | K.Kaveri,S.Sivasubramanian,S.Vennila,P.Padmapriya,R.Kiruba,S.Magesh,G. Dhinakar Raj, G. Ravikumar, P. Azhahianambi,K Thangaraj,Payel Mukherjee, Sofia Banu, Priya Singh, Dhiviya Vedagiri, Divya Gupta, Vishal Sah, Santosh Kumar Kuncha, Krishnan Harinivas Harshan, Archana Bharadwaj Siva, Karthik Bharadwaj Tallapaka, Shagufta Khan, Lamuk Zaveri, Namami Gaur, Sakshi Shambhavi, Tulasi Nagab |
| hCoV-19/India/CCMB_C11/2020 | EPI_ISL_458034 | 4/29/2020 | King Institute of Preventive Medicine & Research | CSIR-Centre for Cellular and Molecular Biology | K.Kaveri,S.Sivasubramanian,S.Vennila,P.Padmapriya,R.Kiruba,S.Magesh,G. Dhinakar Raj, G. Ravikumar, P. Azhahianambi,K Thangaraj,Payel Mukherjee, Sofia Banu, Priya Singh, Dhiviya Vedagiri, Divya Gupta, Vishal Sah, Santosh Kumar Kuncha, Krishnan Harinivas Harshan, Archana Bharadwaj Siva, Karthik Bharadwaj Tallapaka, Shagufta Khan, Lamuk Zaveri, Namami Gaur, Sakshi Shambhavi, Tulasi Nagab |
| hCoV-19/India/CCMB_C13/2020 | EPI_ISL_458035 | 4/29/2020 | King Institute of Preventive Medicine & Research | CSIR-Centre for Cellular and Molecular Biology | K.Kaveri,S.Sivasubramanian,S.Vennila,P.Padmapriya,R.Kiruba,S.Magesh,G. Dhinakar Raj, G. Ravikumar, P. Azhahianambi,K Thangaraj,Payel Mukherjee, Sofia Banu, Priya Singh, Dhiviya Vedagiri, Divya Gupta, Vishal Sah, Santosh Kumar Kuncha, Krishnan Harinivas Harshan, Archana Bharadwaj Siva, Karthik Bharadwaj Tallapaka, Shagufta Khan, Lamuk Zaveri, Namami Gaur, Sakshi Shambhavi, Tulasi Nagab |
| hCoV-19/India/CCMB_C10/2020 | EPI_ISL_458036 | 4/29/2020 | King Institute of Preventive Medicine & Research | CSIR-Centre for Cellular and Molecular Biology | K.Kaveri,S.Sivasubramanian,S.Vennila,P.Padmapriya,R.Kiruba,S.Magesh,G. Dhinakar Raj, G. Ravikumar, P. Azhahianambi,K Thangaraj,Payel Mukherjee, Sofia Banu, Priya Singh, Dhiviya Vedagiri, Divya Gupta, Vishal Sah, Santosh Kumar Kuncha, Krishnan Harinivas Harshan, Archana Bharadwaj Siva, Karthik Bharadwaj Tallapaka, Shagufta Khan, Lamuk Zaveri, Namami Gaur, Sakshi Shambhavi, Tulasi Nagab |
| hCoV-19/India/CCMB_C7/2020 | EPI_ISL_458037 | 4/29/2020 | King Institute of Preventive Medicine & Research | CSIR-Centre for Cellular and Molecular Biology | K.Kaveri,S.Sivasubramanian,S.Vennila,P.Padmapriya,R.Kiruba,S.Magesh,G. Dhinakar Raj, G. Ravikumar, P. Azhahianambi,K Thangaraj,Payel Mukherjee, Sofia Banu, Priya Singh, Dhiviya Vedagiri, Divya Gupta, Vishal Sah, Santosh Kumar Kuncha, Krishnan Harinivas Harshan, Archana Bharadwaj Siva, Karthik Bharadwaj Tallapaka, Shagufta Khan, Lamuk Zaveri, Namami Gaur, Sakshi Shambhavi, Tulasi Nagab |
| hCoV-19/India/CCMB_C9/2020 | EPI_ISL_458038 | 4/29/2020 | King Institute of Preventive Medicine & Research | CSIR-Centre for Cellular and Molecular Biology | K.Kaveri,S.Sivasubramanian,S.Vennila,P.Padmapriya,R.Kiruba,S.Magesh,G. Dhinakar Raj, G. Ravikumar, P. Azhahianambi,K Thangaraj,Payel Mukherjee, Sofia Banu, Priya Singh, Dhiviya Vedagiri, Divya Gupta, Vishal Sah, Santosh Kumar Kuncha, Krishnan Harinivas Harshan, Archana Bharadwaj Siva, Karthik Bharadwaj Tallapaka, Shagufta Khan, Lamuk Zaveri, Namami Gaur, Sakshi Shambhavi, Tulasi Nagab |
| hCoV-19/India/CCMB_C16/2020 | EPI_ISL_458039 | 5/6/2020 | King Institute of Preventive Medicine & Research | CSIR-Centre for Cellular and Molecular Biology | K.Kaveri,S.Sivasubramanian,S.Vennila,P.Padmapriya,R.Kiruba,S.Magesh,G. Dhinakar Raj, G. Ravikumar, P. Azhahianambi,K Thangaraj,Payel Mukherjee, Sofia Banu, Priya Singh, Dhiviya Vedagiri, Divya Gupta, Vishal Sah, Santosh Kumar Kuncha, Krishnan Harinivas Harshan, Archana Bharadwaj Siva, Karthik Bharadwaj Tallapaka, Shagufta Khan, Lamuk Zaveri, Namami Gaur, Sakshi Shambhavi, Tulasi Nagab |
| hCoV-19/India/CCMB_C19/2020 | EPI_ISL_458040 | 5/6/2020 | King Institute of Preventive Medicine & Research | CSIR-Centre for Cellular and Molecular Biology | K.Kaveri,S.Sivasubramanian,S.Vennila,P.Padmapriya,R.Kiruba,S.Magesh,G. Dhinakar Raj, G. Ravikumar, P. Azhahianambi,K Thangaraj,Payel Mukherjee, Sofia Banu, Priya Singh, Dhiviya Vedagiri, Divya Gupta, Vishal Sah, Santosh Kumar Kuncha, Krishnan Harinivas Harshan, Archana Bharadwaj Siva, Karthik Bharadwaj Tallapaka, Shagufta Khan, Lamuk Zaveri, Namami Gaur, Sakshi Shambhavi, Tulasi Nagab |
| hCoV-19/India/CCMB_C15/2020 | EPI_ISL_458041 | 5/6/2020 | King Institute of Preventive Medicine & Research | CSIR-Centre for Cellular and Molecular Biology | K.Kaveri,S.Sivasubramanian,S.Vennila,P.Padmapriya,R.Kiruba,S.Magesh,G. Dhinakar Raj, G. Ravikumar, P. Azhahianambi,K Thangaraj,Payel Mukherjee, Sofia Banu, Priya Singh, Dhiviya Vedagiri, Divya Gupta, Vishal Sah, Santosh Kumar Kuncha, Krishnan Harinivas Harshan, Archana Bharadwaj Siva, Karthik Bharadwaj Tallapaka, Shagufta Khan, Lamuk Zaveri, Namami Gaur, Sakshi Shambhavi, Tulasi Nagab |
| hCoV-19/India/CCMB_C18/2020 | EPI_ISL_458042 | 5/6/2020 | King Institute of Preventive Medicine & Research | CSIR-Centre for Cellular and Molecular Biology | K.Kaveri,S.Sivasubramanian,S.Vennila,P.Padmapriya,R.Kiruba,S.Magesh,G. Dhinakar Raj, G. Ravikumar, P. Azhahianambi,K Thangaraj,Payel Mukherjee, Sofia Banu, Priya Singh, Dhiviya Vedagiri, Divya Gupta, Vishal Sah, Santosh Kumar Kuncha, Krishnan Harinivas Harshan, Archana Bharadwaj Siva, Karthik Bharadwaj Tallapaka, Shagufta Khan, Lamuk Zaveri, Namami Gaur, Sakshi Shambhavi, Tulasi Nagab |
| hCoV-19/India/CCMB_C14/2020 | EPI_ISL_458043 | 5/6/2020 | King Institute of Preventive Medicine & Research | CSIR-Centre for Cellular and Molecular Biology | K.Kaveri,S.Sivasubramanian,S.Vennila,P.Padmapriya,R.Kiruba,S.Magesh,G. Dhinakar Raj, G. Ravikumar, P. Azhahianambi,K Thangaraj,Payel Mukherjee, Sofia Banu, Priya Singh, Dhiviya Vedagiri, Divya Gupta, Vishal Sah, Santosh Kumar Kuncha, Krishnan Harinivas Harshan, Archana Bharadwaj Siva, Karthik Bharadwaj Tallapaka, Shagufta Khan, Lamuk Zaveri, Namami Gaur, Sakshi Shambhavi, Tulasi Nagab |
| hCoV-19/India/CCMB_C17/2020 | EPI_ISL_458044 | 5/6/2020 | King Institute of Preventive Medicine & Research | CSIR-Centre for Cellular and Molecular Biology | K.Kaveri,S.Sivasubramanian,S.Vennila,P.Padmapriya,R.Kiruba,S.Magesh,G. Dhinakar Raj, G. Ravikumar, P. Azhahianambi,K Thangaraj,Payel Mukherjee, Sofia Banu, Priya Singh, Dhiviya Vedagiri, Divya Gupta, Vishal Sah, Santosh Kumar Kuncha, Krishnan Harinivas Harshan, Archana Bharadwaj Siva, Karthik Bharadwaj Tallapaka, Shagufta Khan, Lamuk Zaveri, Namami Gaur, Sakshi Shambhavi, Tulasi Nagab |
| hCoV-19/India/CCMB_L1017/2020 | EPI_ISL_458045 | 5/11/2020 | CSIR-Centre for Cellular and Molecular Biology | CSIR-Centre for Cellular and Molecular Biology | K.Kaveri,S.Sivasubramanian,S.Vennila,P.Padmapriya,R.Kiruba,S.Magesh,G. Dhinakar Raj, G. Ravikumar, P. Azhahianambi,K Thangaraj,Payel Mukherjee, Sofia Banu, Priya Singh, Dhiviya Vedagiri, Divya Gupta, Vishal Sah, Santosh Kumar Kuncha, Krishnan Harinivas Harshan, Archana Bharadwaj Siva, Karthik Bharadwaj Tallapaka, Shagufta Khan, Lamuk Zaveri, Namami Gaur, Sakshi Shambhavi, Tulasi Nagab |
| hCoV-19/India/CCMB_L1021/2020 | EPI_ISL_458046 | 5/11/2020 | CSIR-Centre for Cellular and Molecular Biology | CSIR-Centre for Cellular and Molecular Biology | K.Kaveri,S.Sivasubramanian,S.Vennila,P.Padmapriya,R.Kiruba,S.Magesh,G. Dhinakar Raj, G. Ravikumar, P. Azhahianambi,K Thangaraj,Payel Mukherjee, Sofia Banu, Priya Singh, Dhiviya Vedagiri, Divya Gupta, Vishal Sah, Santosh Kumar Kuncha, Krishnan Harinivas Harshan, Archana Bharadwaj Siva, Karthik Bharadwaj Tallapaka, Shagufta Khan, Lamuk Zaveri, Namami Gaur, Sakshi Shambhavi, Tulasi Nagab |
| hCoV-19/India/CCMB_L1022/2020 | EPI_ISL_458047 | 5/11/2020 | CSIR-Centre for Cellular and Molecular Biology | CSIR-Centre for Cellular and Molecular Biology | K.Kaveri,S.Sivasubramanian,S.Vennila,P.Padmapriya,R.Kiruba,S.Magesh,G. Dhinakar Raj, G. Ravikumar, P. Azhahianambi,K Thangaraj,Payel Mukherjee, Sofia Banu, Priya Singh, Dhiviya Vedagiri, Divya Gupta, Vishal Sah, Santosh Kumar Kuncha, Krishnan Harinivas Harshan, Archana Bharadwaj Siva, Karthik Bharadwaj Tallapaka, Shagufta Khan, Lamuk Zaveri, Namami Gaur, Sakshi Shambhavi, Tulasi Nagab |
| hCoV-19/India/CCMB_L1024/2020 | EPI_ISL_458048 | 5/11/2020 | CSIR-Centre for Cellular and Molecular Biology | CSIR-Centre for Cellular and Molecular Biology | K.Kaveri,S.Sivasubramanian,S.Vennila,P.Padmapriya,R.Kiruba,S.Magesh,G. Dhinakar Raj, G. Ravikumar, P. Azhahianambi,K Thangaraj,Payel Mukherjee, Sofia Banu, Priya Singh, Dhiviya Vedagiri, Divya Gupta, Vishal Sah, Santosh Kumar Kuncha, Krishnan Harinivas Harshan, Archana Bharadwaj Siva, Karthik Bharadwaj Tallapaka, Shagufta Khan, Lamuk Zaveri, Namami Gaur, Sakshi Shambhavi, Tulasi Nagab |
| hCoV-19/India/CCMB_L1025/2020 | EPI_ISL_458049 | 5/11/2020 | CSIR-Centre for Cellular and Molecular Biology | CSIR-Centre for Cellular and Molecular Biology | K.Kaveri,S.Sivasubramanian,S.Vennila,P.Padmapriya,R.Kiruba,S.Magesh,G. Dhinakar Raj, G. Ravikumar, P. Azhahianambi,K Thangaraj,Payel Mukherjee, Sofia Banu, Priya Singh, Dhiviya Vedagiri, Divya Gupta, Vishal Sah, Santosh Kumar Kuncha, Krishnan Harinivas Harshan, Archana Bharadwaj Siva, Karthik Bharadwaj Tallapaka, Shagufta Khan, Lamuk Zaveri, Namami Gaur, Sakshi Shambhavi, Tulasi Nagab |
| hCoV-19/India/CCMB_L1026/2020 | EPI_ISL_458050 | 5/11/2020 | CSIR-Centre for Cellular and Molecular Biology | CSIR-Centre for Cellular and Molecular Biology | K.Kaveri,S.Sivasubramanian,S.Vennila,P.Padmapriya,R.Kiruba,S.Magesh,G. Dhinakar Raj, G. Ravikumar, P. Azhahianambi,K Thangaraj,Payel Mukherjee, Sofia Banu, Priya Singh, Dhiviya Vedagiri, Divya Gupta, Vishal Sah, Santosh Kumar Kuncha, Krishnan Harinivas Harshan, Archana Bharadwaj Siva, Karthik Bharadwaj Tallapaka, Shagufta Khan, Lamuk Zaveri, Namami Gaur, Sakshi Shambhavi, Tulasi Nagab |
| hCoV-19/India/CCMB_L1062/2020 | EPI_ISL_458051 | 5/12/2020 | CSIR-Centre for Cellular and Molecular Biology | CSIR-Centre for Cellular and Molecular Biology | K.Kaveri,S.Sivasubramanian,S.Vennila,P.Padmapriya,R.Kiruba,S.Magesh,G. Dhinakar Raj, G. Ravikumar, P. Azhahianambi,K Thangaraj,Payel Mukherjee, Sofia Banu, Priya Singh, Dhiviya Vedagiri, Divya Gupta, Vishal Sah, Santosh Kumar Kuncha, Krishnan Harinivas Harshan, Archana Bharadwaj Siva, Karthik Bharadwaj Tallapaka, Shagufta Khan, Lamuk Zaveri, Namami Gaur, Sakshi Shambhavi, Tulasi Nagab |
| hCoV-19/India/CCMB_L1063/2020 | EPI_ISL_458052 | 5/12/2020 | CSIR-Centre for Cellular and Molecular Biology | CSIR-Centre for Cellular and Molecular Biology | K.Kaveri,S.Sivasubramanian,S.Vennila,P.Padmapriya,R.Kiruba,S.Magesh,G. Dhinakar Raj, G. Ravikumar, P. Azhahianambi,K Thangaraj,Payel Mukherjee, Sofia Banu, Priya Singh, Dhiviya Vedagiri, Divya Gupta, Vishal Sah, Santosh Kumar Kuncha, Krishnan Harinivas Harshan, Archana Bharadwaj Siva, Karthik Bharadwaj Tallapaka, Shagufta Khan, Lamuk Zaveri, Namami Gaur, Sakshi Shambhavi, Tulasi Nagab |
| hCoV-19/India/CCMB_L1064/2020 | EPI_ISL_458053 | 5/12/2020 | CSIR-Centre for Cellular and Molecular Biology | CSIR-Centre for Cellular and Molecular Biology | K.Kaveri,S.Sivasubramanian,S.Vennila,P.Padmapriya,R.Kiruba,S.Magesh,G. Dhinakar Raj, G. Ravikumar, P. Azhahianambi,K Thangaraj,Payel Mukherjee, Sofia Banu, Priya Singh, Dhiviya Vedagiri, Divya Gupta, Vishal Sah, Santosh Kumar Kuncha, Krishnan Harinivas Harshan, Archana Bharadwaj Siva, Karthik Bharadwaj Tallapaka, Shagufta Khan, Lamuk Zaveri, Namami Gaur, Sakshi Shambhavi, Tulasi Nagab |
| hCoV-19/India/CCMB_L1073/2020 | EPI_ISL_458054 | 5/13/2020 | CSIR-Centre for Cellular and Molecular Biology | CSIR-Centre for Cellular and Molecular Biology | K.Kaveri,S.Sivasubramanian,S.Vennila,P.Padmapriya,R.Kiruba,S.Magesh,G. Dhinakar Raj, G. Ravikumar, P. Azhahianambi,K Thangaraj,Payel Mukherjee, Sofia Banu, Priya Singh, Dhiviya Vedagiri, Divya Gupta, Vishal Sah, Santosh Kumar Kuncha, Krishnan Harinivas Harshan, Archana Bharadwaj Siva, Karthik Bharadwaj Tallapaka, Shagufta Khan, Lamuk Zaveri, Namami Gaur, Sakshi Shambhavi, Tulasi Nagab |
| hCoV-19/India/CCMB_L1082/2020 | EPI_ISL_458055 | 5/13/2020 | CSIR-Centre for Cellular and Molecular Biology | CSIR-Centre for Cellular and Molecular Biology | K.Kaveri,S.Sivasubramanian,S.Vennila,P.Padmapriya,R.Kiruba,S.Magesh,G. Dhinakar Raj, G. Ravikumar, P. Azhahianambi,K Thangaraj,Payel Mukherjee, Sofia Banu, Priya Singh, Dhiviya Vedagiri, Divya Gupta, Vishal Sah, Santosh Kumar Kuncha, Krishnan Harinivas Harshan, Archana Bharadwaj Siva, Karthik Bharadwaj Tallapaka, Shagufta Khan, Lamuk Zaveri, Namami Gaur, Sakshi Shambhavi, Tulasi Nagab |
| hCoV-19/India/CCMB_L1085/2020 | EPI_ISL_458056 | 5/13/2020 | CSIR-Centre for Cellular and Molecular Biology | CSIR-Centre for Cellular and Molecular Biology | K.Kaveri,S.Sivasubramanian,S.Vennila,P.Padmapriya,R.Kiruba,S.Magesh,G. Dhinakar Raj, G. Ravikumar, P. Azhahianambi,K Thangaraj,Payel Mukherjee, Sofia Banu, Priya Singh, Dhiviya Vedagiri, Divya Gupta, Vishal Sah, Santosh Kumar Kuncha, Krishnan Harinivas Harshan, Archana Bharadwaj Siva, Karthik Bharadwaj Tallapaka, Shagufta Khan, Lamuk Zaveri, Namami Gaur, Sakshi Shambhavi, Tulasi Nagab |
| hCoV-19/India/CCMB_L1089/2020 | EPI_ISL_458057 | 5/13/2020 | CSIR-Centre for Cellular and Molecular Biology | CSIR-Centre for Cellular and Molecular Biology | K.Kaveri,S.Sivasubramanian,S.Vennila,P.Padmapriya,R.Kiruba,S.Magesh,G. Dhinakar Raj, G. Ravikumar, P. Azhahianambi,K Thangaraj,Payel Mukherjee, Sofia Banu, Priya Singh, Dhiviya Vedagiri, Divya Gupta, Vishal Sah, Santosh Kumar Kuncha, Krishnan Harinivas Harshan, Archana Bharadwaj Siva, Karthik Bharadwaj Tallapaka, Shagufta Khan, Lamuk Zaveri, Namami Gaur, Sakshi Shambhavi, Tulasi Nagab |
| hCoV-19/India/CCMB_L1090/2020 | EPI_ISL_458058 | 5/13/2020 | CSIR-Centre for Cellular and Molecular Biology | CSIR-Centre for Cellular and Molecular Biology | K.Kaveri,S.Sivasubramanian,S.Vennila,P.Padmapriya,R.Kiruba,S.Magesh,G. Dhinakar Raj, G. Ravikumar, P. Azhahianambi,K Thangaraj,Payel Mukherjee, Sofia Banu, Priya Singh, Dhiviya Vedagiri, Divya Gupta, Vishal Sah, Santosh Kumar Kuncha, Krishnan Harinivas Harshan, Archana Bharadwaj Siva, Karthik Bharadwaj Tallapaka, Shagufta Khan, Lamuk Zaveri, Namami Gaur, Sakshi Shambhavi, Tulasi Nagab |
| hCoV-19/India/CCMB_L1100/2020 | EPI_ISL_458059 | 5/13/2020 | CSIR-Centre for Cellular and Molecular Biology | CSIR-Centre for Cellular and Molecular Biology | K.Kaveri,S.Sivasubramanian,S.Vennila,P.Padmapriya,R.Kiruba,S.Magesh,G. Dhinakar Raj, G. Ravikumar, P. Azhahianambi,K Thangaraj,Payel Mukherjee, Sofia Banu, Priya Singh, Dhiviya Vedagiri, Divya Gupta, Vishal Sah, Santosh Kumar Kuncha, Krishnan Harinivas Harshan, Archana Bharadwaj Siva, Karthik Bharadwaj Tallapaka, Shagufta Khan, Lamuk Zaveri, Namami Gaur, Sakshi Shambhavi, Tulasi Nagab |
| hCoV-19/India/CCMB_L1103/2020 | EPI_ISL_458060 | 5/13/2020 | CSIR-Centre for Cellular and Molecular Biology | CSIR-Centre for Cellular and Molecular Biology | K.Kaveri,S.Sivasubramanian,S.Vennila,P.Padmapriya,R.Kiruba,S.Magesh,G. Dhinakar Raj, G. Ravikumar, P. Azhahianambi,K Thangaraj,Payel Mukherjee, Sofia Banu, Priya Singh, Dhiviya Vedagiri, Divya Gupta, Vishal Sah, Santosh Kumar Kuncha, Krishnan Harinivas Harshan, Archana Bharadwaj Siva, Karthik Bharadwaj Tallapaka, Shagufta Khan, Lamuk Zaveri, Namami Gaur, Sakshi Shambhavi, Tulasi Nagab |
| hCoV-19/India/CCMB_L1108/2020 | EPI_ISL_458061 | 5/13/2020 | CSIR-Centre for Cellular and Molecular Biology | CSIR-Centre for Cellular and Molecular Biology | K.Kaveri,S.Sivasubramanian,S.Vennila,P.Padmapriya,R.Kiruba,S.Magesh,G. Dhinakar Raj, G. Ravikumar, P. Azhahianambi,K Thangaraj,Payel Mukherjee, Sofia Banu, Priya Singh, Dhiviya Vedagiri, Divya Gupta, Vishal Sah, Santosh Kumar Kuncha, Krishnan Harinivas Harshan, Archana Bharadwaj Siva, Karthik Bharadwaj Tallapaka, Shagufta Khan, Lamuk Zaveri, Namami Gaur, Sakshi Shambhavi, Tulasi Nagab |
| hCoV-19/India/CCMB_L299/2020 | EPI_ISL_458062 | 4/20/2020 | CSIR-Centre for Cellular and Molecular Biology | CSIR-Centre for Cellular and Molecular Biology | K.Kaveri,S.Sivasubramanian,S.Vennila,P.Padmapriya,R.Kiruba,S.Magesh,G. Dhinakar Raj, G. Ravikumar, P. Azhahianambi,K Thangaraj,Payel Mukherjee, Sofia Banu, Priya Singh, Dhiviya Vedagiri, Divya Gupta, Vishal Sah, Santosh Kumar Kuncha, Krishnan Harinivas Harshan, Archana Bharadwaj Siva, Karthik Bharadwaj Tallapaka, Shagufta Khan, Lamuk Zaveri, Namami Gaur, Sakshi Shambhavi, Tulasi Nagab |
| hCoV-19/India/CCMB_L301/2020 | EPI_ISL_458063 | 4/20/2020 | CSIR-Centre for Cellular and Molecular Biology | CSIR-Centre for Cellular and Molecular Biology | K.Kaveri,S.Sivasubramanian,S.Vennila,P.Padmapriya,R.Kiruba,S.Magesh,G. Dhinakar Raj, G. Ravikumar, P. Azhahianambi,K Thangaraj,Payel Mukherjee, Sofia Banu, Priya Singh, Dhiviya Vedagiri, Divya Gupta, Vishal Sah, Santosh Kumar Kuncha, Krishnan Harinivas Harshan, Archana Bharadwaj Siva, Karthik Bharadwaj Tallapaka, Shagufta Khan, Lamuk Zaveri, Namami Gaur, Sakshi Shambhavi, Tulasi Nagab |
| hCoV-19/India/CCMB_L988/2020 | EPI_ISL_458064 | 5/11/2020 | CSIR-Centre for Cellular and Molecular Biology | CSIR-Centre for Cellular and Molecular Biology | K.Kaveri,S.Sivasubramanian,S.Vennila,P.Padmapriya,R.Kiruba,S.Magesh,G. Dhinakar Raj, G. Ravikumar, P. Azhahianambi,K Thangaraj,Payel Mukherjee, Sofia Banu, Priya Singh, Dhiviya Vedagiri, Divya Gupta, Vishal Sah, Santosh Kumar Kuncha, Krishnan Harinivas Harshan, Archana Bharadwaj Siva, Karthik Bharadwaj Tallapaka, Shagufta Khan, Lamuk Zaveri, Namami Gaur, Sakshi Shambhavi, Tulasi Nagab |
| hCoV-19/India/CCMB_L989/2020 | EPI_ISL_458065 | 5/11/2020 | CSIR-Centre for Cellular and Molecular Biology | CSIR-Centre for Cellular and Molecular Biology | K.Kaveri,S.Sivasubramanian,S.Vennila,P.Padmapriya,R.Kiruba,S.Magesh,G. Dhinakar Raj, G. Ravikumar, P. Azhahianambi,K Thangaraj,Payel Mukherjee, Sofia Banu, Priya Singh, Dhiviya Vedagiri, Divya Gupta, Vishal Sah, Santosh Kumar Kuncha, Krishnan Harinivas Harshan, Archana Bharadwaj Siva, Karthik Bharadwaj Tallapaka, Shagufta Khan, Lamuk Zaveri, Namami Gaur, Sakshi Shambhavi, Tulasi Nagab |
| hCoV-19/India/CCMB_O1/2020 | EPI_ISL_458066 | 4/13/2020 | Osmania Medical College | CSIR-Centre for Cellular and Molecular Biology | K.Kaveri,S.Sivasubramanian,S.Vennila,P.Padmapriya,R.Kiruba,S.Magesh,G. Dhinakar Raj, G. Ravikumar, P. Azhahianambi,K Thangaraj,Payel Mukherjee, Sofia Banu, Priya Singh, Dhiviya Vedagiri, Divya Gupta, Vishal Sah, Santosh Kumar Kuncha, Krishnan Harinivas Harshan, Archana Bharadwaj Siva, Karthik Bharadwaj Tallapaka, Shagufta Khan, Lamuk Zaveri, Namami Gaur, Sakshi Shambhavi, Tulasi Nagab |
| hCoV-19/India/CCMB_O2/2020 | EPI_ISL_458067 | 4/15/2020 | Osmania Medical College | CSIR-Centre for Cellular and Molecular Biology | K.Kaveri,S.Sivasubramanian,S.Vennila,P.Padmapriya,R.Kiruba,S.Magesh,G. Dhinakar Raj, G. Ravikumar, P. Azhahianambi,K Thangaraj,Payel Mukherjee, Sofia Banu, Priya Singh, Dhiviya Vedagiri, Divya Gupta, Vishal Sah, Santosh Kumar Kuncha, Krishnan Harinivas Harshan, Archana Bharadwaj Siva, Karthik Bharadwaj Tallapaka, Shagufta Khan, Lamuk Zaveri, Namami Gaur, Sakshi Shambhavi, Tulasi Nagab |
| hCoV-19/India/CCMB_O3/2020 | EPI_ISL_458068 | 4/14/2020 | Osmania Medical College | CSIR-Centre for Cellular and Molecular Biology | K.Kaveri,S.Sivasubramanian,S.Vennila,P.Padmapriya,R.Kiruba,S.Magesh,G. Dhinakar Raj, G. Ravikumar, P. Azhahianambi,K Thangaraj,Payel Mukherjee, Sofia Banu, Priya Singh, Dhiviya Vedagiri, Divya Gupta, Vishal Sah, Santosh Kumar Kuncha, Krishnan Harinivas Harshan, Archana Bharadwaj Siva, Karthik Bharadwaj Tallapaka, Shagufta Khan, Lamuk Zaveri, Namami Gaur, Sakshi Shambhavi, Tulasi Nagab |
| hCoV-19/India/CCMB_O4/2020 | EPI_ISL_458069 | 4/3/2020 | Osmania Medical College | CSIR-Centre for Cellular and Molecular Biology | K.Kaveri,S.Sivasubramanian,S.Vennila,P.Padmapriya,R.Kiruba,S.Magesh,G. Dhinakar Raj, G. Ravikumar, P. Azhahianambi,K Thangaraj,Payel Mukherjee, Sofia Banu, Priya Singh, Dhiviya Vedagiri, Divya Gupta, Vishal Sah, Santosh Kumar Kuncha, Krishnan Harinivas Harshan, Archana Bharadwaj Siva, Karthik Bharadwaj Tallapaka, Shagufta Khan, Lamuk Zaveri, Namami Gaur, Sakshi Shambhavi, Tulasi Nagab |
| hCoV-19/India/CCMB_K272/2020 | EPI_ISL_458070 | 4/12/2020 | CSIR-Centre for Cellular and Molecular Biology | CSIR-Centre for Cellular and Molecular Biology | K.Kaveri,S.Sivasubramanian,S.Vennila,P.Padmapriya,R.Kiruba,S.Magesh,G. Dhinakar Raj, G. Ravikumar, P. Azhahianambi,K Thangaraj,Payel Mukherjee, Sofia Banu, Priya Singh, Dhiviya Vedagiri, Divya Gupta, Vishal Sah, Santosh Kumar Kuncha, Krishnan Harinivas Harshan, Archana Bharadwaj Siva, Karthik Bharadwaj Tallapaka, Shagufta Khan, Lamuk Zaveri, Namami Gaur, Sakshi Shambhavi, Tulasi Nagab |
| hCoV-19/India/CCMB_J130/2020 | EPI_ISL_458071 | 4/1/2020 | CSIR-Centre for Cellular and Molecular Biology | CSIR-Centre for Cellular and Molecular Biology | K.Kaveri,S.Sivasubramanian,S.Vennila,P.Padmapriya,R.Kiruba,S.Magesh,G. Dhinakar Raj, G. Ravikumar, P. Azhahianambi,K Thangaraj,Payel Mukherjee, Sofia Banu, Priya Singh, Dhiviya Vedagiri, Divya Gupta, Vishal Sah, Santosh Kumar Kuncha, Krishnan Harinivas Harshan, Archana Bharadwaj Siva, Karthik Bharadwaj Tallapaka, Shagufta Khan, Lamuk Zaveri, Namami Gaur, Sakshi Shambhavi, Tulasi Nagab |
| hCoV-19/India/CCMB_J812_P1/2020 | EPI_ISL_458072 | 4/8/2020 | CSIR-Centre for Cellular and Molecular Biology | CSIR-Centre for Cellular and Molecular Biology | K.Kaveri,S.Sivasubramanian,S.Vennila,P.Padmapriya,R.Kiruba,S.Magesh,G. Dhinakar Raj, G. Ravikumar, P. Azhahianambi,K Thangaraj,Payel Mukherjee, Sofia Banu, Priya Singh, Dhiviya Vedagiri, Divya Gupta, Vishal Sah, Santosh Kumar Kuncha, Krishnan Harinivas Harshan, Archana Bharadwaj Siva, Karthik Bharadwaj Tallapaka, Shagufta Khan, Lamuk Zaveri, Namami Gaur, Sakshi Shambhavi, Tulasi Nagab |
| hCoV-19/India/CCMB_L276_P1/2020 | EPI_ISL_458073 | 4/20/2020 | CSIR-Centre for Cellular and Molecular Biology | CSIR-Centre for Cellular and Molecular Biology | K.Kaveri,S.Sivasubramanian,S.Vennila,P.Padmapriya,R.Kiruba,S.Magesh,G. Dhinakar Raj, G. Ravikumar, P. Azhahianambi,K Thangaraj,Payel Mukherjee, Sofia Banu, Priya Singh, Dhiviya Vedagiri, Divya Gupta, Vishal Sah, Santosh Kumar Kuncha, Krishnan Harinivas Harshan, Archana Bharadwaj Siva, Karthik Bharadwaj Tallapaka, Shagufta Khan, Lamuk Zaveri, Namami Gaur, Sakshi Shambhavi, Tulasi Nagab |
| hCoV-19/India/CCMB_L301_P1/2020 | EPI_ISL_458074 | 4/20/2020 | CSIR-Centre for Cellular and Molecular Biology | CSIR-Centre for Cellular and Molecular Biology | K.Kaveri,S.Sivasubramanian,S.Vennila,P.Padmapriya,R.Kiruba,S.Magesh,G. Dhinakar Raj, G. Ravikumar, P. Azhahianambi,K Thangaraj,Payel Mukherjee, Sofia Banu, Priya Singh, Dhiviya Vedagiri, Divya Gupta, Vishal Sah, Santosh Kumar Kuncha, Krishnan Harinivas Harshan, Archana Bharadwaj Siva, Karthik Bharadwaj Tallapaka, Shagufta Khan, Lamuk Zaveri, Namami Gaur, Sakshi Shambhavi, Tulasi Nagab |
| hCoV-19/India/CCMB_O2_P1/2020 | EPI_ISL_458075 | 4/15/2020 | CSIR-Centre for Cellular and Molecular Biology | CSIR-Centre for Cellular and Molecular Biology | K.Kaveri,S.Sivasubramanian,S.Vennila,P.Padmapriya,R.Kiruba,S.Magesh,G. Dhinakar Raj, G. Ravikumar, P. Azhahianambi,K Thangaraj,Payel Mukherjee, Sofia Banu, Priya Singh, Dhiviya Vedagiri, Divya Gupta, Vishal Sah, Santosh Kumar Kuncha, Krishnan Harinivas Harshan, Archana Bharadwaj Siva, Karthik Bharadwaj Tallapaka, Shagufta Khan, Lamuk Zaveri, Namami Gaur, Sakshi Shambhavi, Tulasi Nagab |
| hCoV-19/India/CCMB_L302_P1/2020 | EPI_ISL_458076 | 4/20/2020 | CSIR-Centre for Cellular and Molecular Biology | CSIR-Centre for Cellular and Molecular Biology | K.Kaveri,S.Sivasubramanian,S.Vennila,P.Padmapriya,R.Kiruba,S.Magesh,G. Dhinakar Raj, G. Ravikumar, P. Azhahianambi,K Thangaraj,Payel Mukherjee, Sofia Banu, Priya Singh, Dhiviya Vedagiri, Divya Gupta, Vishal Sah, Santosh Kumar Kuncha, Krishnan Harinivas Harshan, Archana Bharadwaj Siva, Karthik Bharadwaj Tallapaka, Shagufta Khan, Lamuk Zaveri, Namami Gaur, Sakshi Shambhavi, Tulasi Nagab |
| hCoV-19/India/CCMB_L040_P1/2020 | EPI_ISL_458077 | 4/18/2020 | CSIR-Centre for Cellular and Molecular Biology | CSIR-Centre for Cellular and Molecular Biology | K.Kaveri,S.Sivasubramanian,S.Vennila,P.Padmapriya,R.Kiruba,S.Magesh,G. Dhinakar Raj, G. Ravikumar, P. Azhahianambi,K Thangaraj,Payel Mukherjee, Sofia Banu, Priya Singh, Dhiviya Vedagiri, Divya Gupta, Vishal Sah, Santosh Kumar Kuncha, Krishnan Harinivas Harshan, Archana Bharadwaj Siva, Karthik Bharadwaj Tallapaka, Shagufta Khan, Lamuk Zaveri, Namami Gaur, Sakshi Shambhavi, Tulasi Nagab |
| hCoV-19/India/CCMB_J300/2020 | EPI_ISL_458080 | 4/3/2020 | CSIR-Centre for Cellular and Molecular Biology | CSIR-Centre for Cellular and Molecular Biology | K.Kaveri,S.Sivasubramanian,S.Vennila,P.Padmapriya,R.Kiruba,S.Magesh,G. Dhinakar Raj, G. Ravikumar, P. Azhahianambi,K Thangaraj,Payel Mukherjee, Sofia Banu, Priya Singh, Dhiviya Vedagiri, Divya Gupta, Vishal Sah, Santosh Kumar Kuncha, Krishnan Harinivas Harshan, Archana Bharadwaj Siva, Karthik Bharadwaj Tallapaka, Shagufta Khan, Lamuk Zaveri, Namami Gaur, Sakshi Shambhavi, Tulasi Nagab |
| hCoV-19/India/GBRC131/2020 | EPI_ISL_458086 | 5/24/2020 | B.J. Medical College and Civil hospital | Gujarat Biotechnology Research Centre | K.Kaveri,S.Sivasubramanian,S.Vennila,P.Padmapriya,R.Kiruba,S.Magesh,G. Dhinakar Raj, G. Ravikumar, P. Azhahianambi,K Thangaraj,Payel Mukherjee, Sofia Banu, Priya Singh, Dhiviya Vedagiri, Divya Gupta, Vishal Sah, Santosh Kumar Kuncha, Krishnan Harinivas Harshan, Archana Bharadwaj Siva, Karthik Bharadwaj Tallapaka, Shagufta Khan, Lamuk Zaveri, Namami Gaur, Sakshi Shambhavi, Tulasi Nagab |
| hCoV-19/India/GBRC132/2020 | EPI_ISL_458087 | 5/24/2020 | B.J. Medical College and Civil hospital | Gujarat Biotechnology Research Centre | K.Kaveri,S.Sivasubramanian,S.Vennila,P.Padmapriya,R.Kiruba,S.Magesh,G. Dhinakar Raj, G. Ravikumar, P. Azhahianambi,K Thangaraj,Payel Mukherjee, Sofia Banu, Priya Singh, Dhiviya Vedagiri, Divya Gupta, Vishal Sah, Santosh Kumar Kuncha, Krishnan Harinivas Harshan, Archana Bharadwaj Siva, Karthik Bharadwaj Tallapaka, Shagufta Khan, Lamuk Zaveri, Namami Gaur, Sakshi Shambhavi, Tulasi Nagab |
| hCoV-19/India/GBRC133/2020 | EPI_ISL_458088 | 5/24/2020 | B.J. Medical College and Civil hospital | Gujarat Biotechnology Research Centre | K.Kaveri,S.Sivasubramanian,S.Vennila,P.Padmapriya,R.Kiruba,S.Magesh,G. Dhinakar Raj, G. Ravikumar, P. Azhahianambi,K Thangaraj,Payel Mukherjee, Sofia Banu, Priya Singh, Dhiviya Vedagiri, Divya Gupta, Vishal Sah, Santosh Kumar Kuncha, Krishnan Harinivas Harshan, Archana Bharadwaj Siva, Karthik Bharadwaj Tallapaka, Shagufta Khan, Lamuk Zaveri, Namami Gaur, Sakshi Shambhavi, Tulasi Nagab |
| hCoV-19/India/GBRC134/2020 | EPI_ISL_458089 | 5/24/2020 | B.J. Medical College and Civil hospital | Gujarat Biotechnology Research Centre | K.Kaveri,S.Sivasubramanian,S.Vennila,P.Padmapriya,R.Kiruba,S.Magesh,G. Dhinakar Raj, G. Ravikumar, P. Azhahianambi,K Thangaraj,Payel Mukherjee, Sofia Banu, Priya Singh, Dhiviya Vedagiri, Divya Gupta, Vishal Sah, Santosh Kumar Kuncha, Krishnan Harinivas Harshan, Archana Bharadwaj Siva, Karthik Bharadwaj Tallapaka, Shagufta Khan, Lamuk Zaveri, Namami Gaur, Sakshi Shambhavi, Tulasi Nagab |
| hCoV-19/India/GBRC135a/2020 | EPI_ISL_458090 | 5/24/2020 | B.J. Medical College and Civil hospital | Gujarat Biotechnology Research Centre | K.Kaveri,S.Sivasubramanian,S.Vennila,P.Padmapriya,R.Kiruba,S.Magesh,G. Dhinakar Raj, G. Ravikumar, P. Azhahianambi,K Thangaraj,Payel Mukherjee, Sofia Banu, Priya Singh, Dhiviya Vedagiri, Divya Gupta, Vishal Sah, Santosh Kumar Kuncha, Krishnan Harinivas Harshan, Archana Bharadwaj Siva, Karthik Bharadwaj Tallapaka, Shagufta Khan, Lamuk Zaveri, Namami Gaur, Sakshi Shambhavi, Tulasi Nagab |
| hCoV-19/India/GBRC135b/2020 | EPI_ISL_458091 | 5/24/2020 | B.J. Medical College and Civil hospital | Gujarat Biotechnology Research Centre | K.Kaveri,S.Sivasubramanian,S.Vennila,P.Padmapriya,R.Kiruba,S.Magesh,G. Dhinakar Raj, G. Ravikumar, P. Azhahianambi,K Thangaraj,Payel Mukherjee, Sofia Banu, Priya Singh, Dhiviya Vedagiri, Divya Gupta, Vishal Sah, Santosh Kumar Kuncha, Krishnan Harinivas Harshan, Archana Bharadwaj Siva, Karthik Bharadwaj Tallapaka, Shagufta Khan, Lamuk Zaveri, Namami Gaur, Sakshi Shambhavi, Tulasi Nagab |
| hCoV-19/India/GBRC136a/2020 | EPI_ISL_458092 | 5/24/2020 | B.J. Medical College and Civil hospital | Gujarat Biotechnology Research Centre | K.Kaveri,S.Sivasubramanian,S.Vennila,P.Padmapriya,R.Kiruba,S.Magesh,G. Dhinakar Raj, G. Ravikumar, P. Azhahianambi,K Thangaraj,Payel Mukherjee, Sofia Banu, Priya Singh, Dhiviya Vedagiri, Divya Gupta, Vishal Sah, Santosh Kumar Kuncha, Krishnan Harinivas Harshan, Archana Bharadwaj Siva, Karthik Bharadwaj Tallapaka, Shagufta Khan, Lamuk Zaveri, Namami Gaur, Sakshi Shambhavi, Tulasi Nagab |
| hCoV-19/India/GBRC136b/2020 | EPI_ISL_458093 | 5/24/2020 | B.J. Medical College and Civil hospital | Gujarat Biotechnology Research Centre | K.Kaveri,S.Sivasubramanian,S.Vennila,P.Padmapriya,R.Kiruba,S.Magesh,G. Dhinakar Raj, G. Ravikumar, P. Azhahianambi,K Thangaraj,Payel Mukherjee, Sofia Banu, Priya Singh, Dhiviya Vedagiri, Divya Gupta, Vishal Sah, Santosh Kumar Kuncha, Krishnan Harinivas Harshan, Archana Bharadwaj Siva, Karthik Bharadwaj Tallapaka, Shagufta Khan, Lamuk Zaveri, Namami Gaur, Sakshi Shambhavi, Tulasi Nagab |
| hCoV-19/India/GBRC137a/2020 | EPI_ISL_458094 | 5/24/2020 | B.J. Medical College and Civil hospital | Gujarat Biotechnology Research Centre | K.Kaveri,S.Sivasubramanian,S.Vennila,P.Padmapriya,R.Kiruba,S.Magesh,G. Dhinakar Raj, G. Ravikumar, P. Azhahianambi,K Thangaraj,Payel Mukherjee, Sofia Banu, Priya Singh, Dhiviya Vedagiri, Divya Gupta, Vishal Sah, Santosh Kumar Kuncha, Krishnan Harinivas Harshan, Archana Bharadwaj Siva, Karthik Bharadwaj Tallapaka, Shagufta Khan, Lamuk Zaveri, Namami Gaur, Sakshi Shambhavi, Tulasi Nagab |
| hCoV-19/India/GBRC137b/2020 | EPI_ISL_458095 | 5/24/2020 | B.J. Medical College and Civil hospital | Gujarat Biotechnology Research Centre | K.Kaveri,S.Sivasubramanian,S.Vennila,P.Padmapriya,R.Kiruba,S.Magesh,G. Dhinakar Raj, G. Ravikumar, P. Azhahianambi,K Thangaraj,Payel Mukherjee, Sofia Banu, Priya Singh, Dhiviya Vedagiri, Divya Gupta, Vishal Sah, Santosh Kumar Kuncha, Krishnan Harinivas Harshan, Archana Bharadwaj Siva, Karthik Bharadwaj Tallapaka, Shagufta Khan, Lamuk Zaveri, Namami Gaur, Sakshi Shambhavi, Tulasi Nagab |
| hCoV-19/India/GBRC138/2020 | EPI_ISL_458096 | 5/24/2020 | B.J. Medical College and Civil hospital | Gujarat Biotechnology Research Centre | K.Kaveri,S.Sivasubramanian,S.Vennila,P.Padmapriya,R.Kiruba,S.Magesh,G. Dhinakar Raj, G. Ravikumar, P. Azhahianambi,K Thangaraj,Payel Mukherjee, Sofia Banu, Priya Singh, Dhiviya Vedagiri, Divya Gupta, Vishal Sah, Santosh Kumar Kuncha, Krishnan Harinivas Harshan, Archana Bharadwaj Siva, Karthik Bharadwaj Tallapaka, Shagufta Khan, Lamuk Zaveri, Namami Gaur, Sakshi Shambhavi, Tulasi Nagab |
| hCoV-19/India/GBRC139a/2020 | EPI_ISL_458097 | 5/24/2020 | B.J. Medical College and Civil hospital | Gujarat Biotechnology Research Centre | K.Kaveri,S.Sivasubramanian,S.Vennila,P.Padmapriya,R.Kiruba,S.Magesh,G. Dhinakar Raj, G. Ravikumar, P. Azhahianambi,K Thangaraj,Payel Mukherjee, Sofia Banu, Priya Singh, Dhiviya Vedagiri, Divya Gupta, Vishal Sah, Santosh Kumar Kuncha, Krishnan Harinivas Harshan, Archana Bharadwaj Siva, Karthik Bharadwaj Tallapaka, Shagufta Khan, Lamuk Zaveri, Namami Gaur, Sakshi Shambhavi, Tulasi Nagab |
| hCoV-19/India/GBRC139b/2020 | EPI_ISL_458098 | 5/24/2020 | B.J. Medical College and Civil hospital | Gujarat Biotechnology Research Centre | K.Kaveri,S.Sivasubramanian,S.Vennila,P.Padmapriya,R.Kiruba,S.Magesh,G. Dhinakar Raj, G. Ravikumar, P. Azhahianambi,K Thangaraj,Payel Mukherjee, Sofia Banu, Priya Singh, Dhiviya Vedagiri, Divya Gupta, Vishal Sah, Santosh Kumar Kuncha, Krishnan Harinivas Harshan, Archana Bharadwaj Siva, Karthik Bharadwaj Tallapaka, Shagufta Khan, Lamuk Zaveri, Namami Gaur, Sakshi Shambhavi, Tulasi Nagab |
| hCoV-19/India/GBRC140/2020 | EPI_ISL_458099 | 5/24/2020 | B.J. Medical College and Civil hospital | Gujarat Biotechnology Research Centre | K.Kaveri,S.Sivasubramanian,S.Vennila,P.Padmapriya,R.Kiruba,S.Magesh,G. Dhinakar Raj, G. Ravikumar, P. Azhahianambi,K Thangaraj,Payel Mukherjee, Sofia Banu, Priya Singh, Dhiviya Vedagiri, Divya Gupta, Vishal Sah, Santosh Kumar Kuncha, Krishnan Harinivas Harshan, Archana Bharadwaj Siva, Karthik Bharadwaj Tallapaka, Shagufta Khan, Lamuk Zaveri, Namami Gaur, Sakshi Shambhavi, Tulasi Nagab |
| hCoV-19/India/GBRC141a/2020 | EPI_ISL_458100 | 5/24/2020 | B.J. Medical College and Civil hospital | Gujarat Biotechnology Research Centre | K.Kaveri,S.Sivasubramanian,S.Vennila,P.Padmapriya,R.Kiruba,S.Magesh,G. Dhinakar Raj, G. Ravikumar, P. Azhahianambi,K Thangaraj,Payel Mukherjee, Sofia Banu, Priya Singh, Dhiviya Vedagiri, Divya Gupta, Vishal Sah, Santosh Kumar Kuncha, Krishnan Harinivas Harshan, Archana Bharadwaj Siva, Karthik Bharadwaj Tallapaka, Shagufta Khan, Lamuk Zaveri, Namami Gaur, Sakshi Shambhavi, Tulasi Nagab |
| hCoV-19/India/GBRC141b/2020 | EPI_ISL_458101 | 5/24/2020 | B.J. Medical College and Civil hospital | Gujarat Biotechnology Research Centre | K.Kaveri,S.Sivasubramanian,S.Vennila,P.Padmapriya,R.Kiruba,S.Magesh,G. Dhinakar Raj, G. Ravikumar, P. Azhahianambi,K Thangaraj,Payel Mukherjee, Sofia Banu, Priya Singh, Dhiviya Vedagiri, Divya Gupta, Vishal Sah, Santosh Kumar Kuncha, Krishnan Harinivas Harshan, Archana Bharadwaj Siva, Karthik Bharadwaj Tallapaka, Shagufta Khan, Lamuk Zaveri, Namami Gaur, Sakshi Shambhavi, Tulasi Nagab |
| hCoV-19/India/GBRC142/2020 | EPI_ISL_458102 | 5/24/2020 | B.J. Medical College and Civil hospital | Gujarat Biotechnology Research Centre | K.Kaveri,S.Sivasubramanian,S.Vennila,P.Padmapriya,R.Kiruba,S.Magesh,G. Dhinakar Raj, G. Ravikumar, P. Azhahianambi,K Thangaraj,Payel Mukherjee, Sofia Banu, Priya Singh, Dhiviya Vedagiri, Divya Gupta, Vishal Sah, Santosh Kumar Kuncha, Krishnan Harinivas Harshan, Archana Bharadwaj Siva, Karthik Bharadwaj Tallapaka, Shagufta Khan, Lamuk Zaveri, Namami Gaur, Sakshi Shambhavi, Tulasi Nagab |
| hCoV-19/India/GBRC143/2020 | EPI_ISL_458103 | 5/5/2020 | Gujarat Biotechnology Research Centre | Gujarat Biotechnology Research Centre | K.Kaveri,S.Sivasubramanian,S.Vennila,P.Padmapriya,R.Kiruba,S.Magesh,G. Dhinakar Raj, G. Ravikumar, P. Azhahianambi,K Thangaraj,Payel Mukherjee, Sofia Banu, Priya Singh, Dhiviya Vedagiri, Divya Gupta, Vishal Sah, Santosh Kumar Kuncha, Krishnan Harinivas Harshan, Archana Bharadwaj Siva, Karthik Bharadwaj Tallapaka, Shagufta Khan, Lamuk Zaveri, Namami Gaur, Sakshi Shambhavi, Tulasi Nagab |
| hCoV-19/India/GBRC144/2020 | EPI_ISL_458104 | 5/6/2020 | Gujarat Biotechnology Research Centre | Gujarat Biotechnology Research Centre | K.Kaveri,S.Sivasubramanian,S.Vennila,P.Padmapriya,R.Kiruba,S.Magesh,G. Dhinakar Raj, G. Ravikumar, P. Azhahianambi,K Thangaraj,Payel Mukherjee, Sofia Banu, Priya Singh, Dhiviya Vedagiri, Divya Gupta, Vishal Sah, Santosh Kumar Kuncha, Krishnan Harinivas Harshan, Archana Bharadwaj Siva, Karthik Bharadwaj Tallapaka, Shagufta Khan, Lamuk Zaveri, Namami Gaur, Sakshi Shambhavi, Tulasi Nagab |
| hCoV-19/India/GBRC145/2020 | EPI_ISL_458105 | 5/6/2020 | Gujarat Biotechnology Research Centre | Gujarat Biotechnology Research Centre | K.Kaveri,S.Sivasubramanian,S.Vennila,P.Padmapriya,R.Kiruba,S.Magesh,G. Dhinakar Raj, G. Ravikumar, P. Azhahianambi,K Thangaraj,Payel Mukherjee, Sofia Banu, Priya Singh, Dhiviya Vedagiri, Divya Gupta, Vishal Sah, Santosh Kumar Kuncha, Krishnan Harinivas Harshan, Archana Bharadwaj Siva, Karthik Bharadwaj Tallapaka, Shagufta Khan, Lamuk Zaveri, Namami Gaur, Sakshi Shambhavi, Tulasi Nagab |
| hCoV-19/India/GBRC146a/2020 | EPI_ISL_458106 | 5/16/2020 | Gujarat Biotechnology Research Centre | Gujarat Biotechnology Research Centre | K.Kaveri,S.Sivasubramanian,S.Vennila,P.Padmapriya,R.Kiruba,S.Magesh,G. Dhinakar Raj, G. Ravikumar, P. Azhahianambi,K Thangaraj,Payel Mukherjee, Sofia Banu, Priya Singh, Dhiviya Vedagiri, Divya Gupta, Vishal Sah, Santosh Kumar Kuncha, Krishnan Harinivas Harshan, Archana Bharadwaj Siva, Karthik Bharadwaj Tallapaka, Shagufta Khan, Lamuk Zaveri, Namami Gaur, Sakshi Shambhavi, Tulasi Nagab |
| hCoV-19/India/GBRC146b/2020 | EPI_ISL_458107 | 5/16/2020 | Gujarat Biotechnology Research Centre | Gujarat Biotechnology Research Centre | K.Kaveri,S.Sivasubramanian,S.Vennila,P.Padmapriya,R.Kiruba,S.Magesh,G. Dhinakar Raj, G. Ravikumar, P. Azhahianambi,K Thangaraj,Payel Mukherjee, Sofia Banu, Priya Singh, Dhiviya Vedagiri, Divya Gupta, Vishal Sah, Santosh Kumar Kuncha, Krishnan Harinivas Harshan, Archana Bharadwaj Siva, Karthik Bharadwaj Tallapaka, Shagufta Khan, Lamuk Zaveri, Namami Gaur, Sakshi Shambhavi, Tulasi Nagab |
| hCoV-19/India/GBRC147/2020 | EPI_ISL_458108 | 5/16/2020 | Gujarat Biotechnology Research Centre | Gujarat Biotechnology Research Centre | K.Kaveri,S.Sivasubramanian,S.Vennila,P.Padmapriya,R.Kiruba,S.Magesh,G. Dhinakar Raj, G. Ravikumar, P. Azhahianambi,K Thangaraj,Payel Mukherjee, Sofia Banu, Priya Singh, Dhiviya Vedagiri, Divya Gupta, Vishal Sah, Santosh Kumar Kuncha, Krishnan Harinivas Harshan, Archana Bharadwaj Siva, Karthik Bharadwaj Tallapaka, Shagufta Khan, Lamuk Zaveri, Namami Gaur, Sakshi Shambhavi, Tulasi Nagab |
| hCoV-19/India/GBRC148a/2020 | EPI_ISL_458109 | 5/17/2020 | Gujarat Biotechnology Research Centre | Gujarat Biotechnology Research Centre | K.Kaveri,S.Sivasubramanian,S.Vennila,P.Padmapriya,R.Kiruba,S.Magesh,G. Dhinakar Raj, G. Ravikumar, P. Azhahianambi,K Thangaraj,Payel Mukherjee, Sofia Banu, Priya Singh, Dhiviya Vedagiri, Divya Gupta, Vishal Sah, Santosh Kumar Kuncha, Krishnan Harinivas Harshan, Archana Bharadwaj Siva, Karthik Bharadwaj Tallapaka, Shagufta Khan, Lamuk Zaveri, Namami Gaur, Sakshi Shambhavi, Tulasi Nagab |
| hCoV-19/India/GBRC148b/2020 | EPI_ISL_458110 | 5/17/2020 | Gujarat Biotechnology Research Centre | Gujarat Biotechnology Research Centre | K.Kaveri,S.Sivasubramanian,S.Vennila,P.Padmapriya,R.Kiruba,S.Magesh,G. Dhinakar Raj, G. Ravikumar, P. Azhahianambi,K Thangaraj,Payel Mukherjee, Sofia Banu, Priya Singh, Dhiviya Vedagiri, Divya Gupta, Vishal Sah, Santosh Kumar Kuncha, Krishnan Harinivas Harshan, Archana Bharadwaj Siva, Karthik Bharadwaj Tallapaka, Shagufta Khan, Lamuk Zaveri, Namami Gaur, Sakshi Shambhavi, Tulasi Nagab |
| hCoV-19/India/GBRC149/2020 | EPI_ISL_458111 | 5/17/2020 | Gujarat Biotechnology Research Centre | Gujarat Biotechnology Research Centre | K.Kaveri,S.Sivasubramanian,S.Vennila,P.Padmapriya,R.Kiruba,S.Magesh,G. Dhinakar Raj, G. Ravikumar, P. Azhahianambi,K Thangaraj,Payel Mukherjee, Sofia Banu, Priya Singh, Dhiviya Vedagiri, Divya Gupta, Vishal Sah, Santosh Kumar Kuncha, Krishnan Harinivas Harshan, Archana Bharadwaj Siva, Karthik Bharadwaj Tallapaka, Shagufta Khan, Lamuk Zaveri, Namami Gaur, Sakshi Shambhavi, Tulasi Nagab |
| hCoV-19/India/GBRC150a/2020 | EPI_ISL_458112 | 5/18/2020 | Gujarat Biotechnology Research Centre | Gujarat Biotechnology Research Centre | K.Kaveri,S.Sivasubramanian,S.Vennila,P.Padmapriya,R.Kiruba,S.Magesh,G. Dhinakar Raj, G. Ravikumar, P. Azhahianambi,K Thangaraj,Payel Mukherjee, Sofia Banu, Priya Singh, Dhiviya Vedagiri, Divya Gupta, Vishal Sah, Santosh Kumar Kuncha, Krishnan Harinivas Harshan, Archana Bharadwaj Siva, Karthik Bharadwaj Tallapaka, Shagufta Khan, Lamuk Zaveri, Namami Gaur, Sakshi Shambhavi, Tulasi Nagab |
| hCoV-19/India/GBRC150b/2020 | EPI_ISL_458113 | 5/18/2020 | Gujarat Biotechnology Research Centre | Gujarat Biotechnology Research Centre | K.Kaveri,S.Sivasubramanian,S.Vennila,P.Padmapriya,R.Kiruba,S.Magesh,G. Dhinakar Raj, G. Ravikumar, P. Azhahianambi,K Thangaraj,Payel Mukherjee, Sofia Banu, Priya Singh, Dhiviya Vedagiri, Divya Gupta, Vishal Sah, Santosh Kumar Kuncha, Krishnan Harinivas Harshan, Archana Bharadwaj Siva, Karthik Bharadwaj Tallapaka, Shagufta Khan, Lamuk Zaveri, Namami Gaur, Sakshi Shambhavi, Tulasi Nagab |
| hCoV-19/India/CCMB_J714/2020 | EPI_ISL_458298 | 4/6/2020 | CSIR-Centre for Cellular and Molecular Biology | CSIR-Centre for Cellular and Molecular Biology | Sakshi Shambhavi, Lamuk Zaveri, Shagufta Khan, Namami Gaur, Tulasi Nagabandi, Purushotham Vodnala, Payel Mukherjee, Sofia Banu, Priya Singh, Dhiviya Vedagiri, Divya Gupta, Vishal Sah, Santosh Kumar Kuncha, Krishnan Harinivas Harshan, Archana Bharadwaj Siva, Karthik Bharadwaj Tallapaka, Deepak Kumar, Devi Prasad Vijayashankar, Disha Nanda, Divya Das, Jotin Gogoi, Manish Bhattacharjee, R |
| hCoV-19/India/MaxCov0003_CSIR-IGIB | EPI_ISL_459911 | 5/9/2020 | Devki Devi Foundation, a unit of Max Healthcare | CSIR-IGIB/Max | Rajesh Pandey#, Samreen Siddiqui, Pooja Sharma, Bansidhar Tarai, Vivekanand A, Bharathram Uppili, Saruchi Wadhwa, Nishu Tyagi, Mitali Mukerji, Poonam Das, Sujeet Jha, Mohammed Faruq, Vinita Jha, Anurag Agrawal |
| hCoV-19/India/MaxCov0010_CSIR-IGIB | EPI_ISL_459913 | 5/3/2020 | Devki Devi Foundation, a unit of Max Healthcare | CSIR-IGIB/Max | Rajesh Pandey#, Samreen Siddiqui, Pooja Sharma, Bansidhar Tarai, Vivekanand A, Bharathram Uppili, Saruchi Wadhwa, Nishu Tyagi, Mitali Mukerji, Bansidhar Tarai, Poonam Das, Sujeet Jha, Mohammed Faruq, Vinita Jha, Anurag Agrawal |
| hCoV-19/India/MaxCov0012_CSIR-IGIB | EPI_ISL_459914 | 5/1/2020 | Devki Devi Foundation, a unit of Max Healthcare | CSIR-IGIB/Max | Rajesh Pandey#, Samreen Siddiqui, Pooja Sharma, Bansidhar Tarai, Vivekanand A, Bharathram Uppili, Saruchi Wadhwa, Nishu Tyagi, Mitali Mukerji, Bansidhar Tarai, Poonam Das, Sujeet Jha, Mohammed Faruq, Vinita Jha, Anurag Agrawal |
| hCoV-19/India/MaxCov0013_CSIR-IGIB | EPI_ISL_459915 | 5/12/2020 | Devki Devi Foundation, a unit of Max Healthcare | CSIR-IGIB/Max | Rajesh Pandey#, Samreen Siddiqui, Pooja Sharma, Bansidhar Tarai, Vivekanand A, Bharathram Uppili, Saruchi Wadhwa, Nishu Tyagi, Mitali Mukerji, Bansidhar Tarai, Poonam Das, Sujeet Jha, Mohammed Faruq, Vinita Jha, Anurag Agrawal |
| hCoV-19/India/MaxCov0014_CSIR-IGIB | EPI_ISL_459916 | 5/12/2020 | Devki Devi Foundation, a unit of Max Healthcare | CSIR-IGIB/Max | Rajesh Pandey#, Samreen Siddiqui, Pooja Sharma, Bansidhar Tarai, Vivekanand A, Bharathram Uppili, Saruchi Wadhwa, Nishu Tyagi, Mitali Mukerji, Bansidhar Tarai, Poonam Das, Sujeet Jha, Mohammed Faruq, Vinita Jha, Anurag Agrawal |
| hCoV-19/India/MaxCov0015_CSIR-IGIB | EPI_ISL_459917 | 5/8/2020 | Devki Devi Foundation, a unit of Max Healthcare | CSIR-IGIB/Max | Rajesh Pandey#, Samreen Siddiqui, Pooja Sharma, Bansidhar Tarai, Vivekanand A, Bharathram Uppili, Saruchi Wadhwa, Nishu Tyagi, Mitali Mukerji, Bansidhar Tarai, Poonam Das, Sujeet Jha, Mohammed Faruq, Vinita Jha, Anurag Agrawal |
| hCoV-19/India/MaxCov0016_CSIR-IGIB | EPI_ISL_459918 | 5/9/2020 | Devki Devi Foundation, a unit of Max Healthcare | CSIR-IGIB/Max | Rajesh Pandey#, Samreen Siddiqui, Pooja Sharma, Bansidhar Tarai, Vivekanand A, Bharathram Uppili, Saruchi Wadhwa, Nishu Tyagi, Mitali Mukerji, Bansidhar Tarai, Poonam Das, Sujeet Jha, Mohammed Faruq, Vinita Jha, Anurag Agrawal |
| hCoV-19/India/MaxCov0017_CSIR-IGIB | EPI_ISL_459919 | 5/21/2020 | Devki Devi Foundation, a unit of Max Healthcare | CSIR-IGIB/Max | Rajesh Pandey#, Samreen Siddiqui, Pooja Sharma, Bansidhar Tarai, Vivekanand A, Bharathram Uppili, Saruchi Wadhwa, Nishu Tyagi, Mitali Mukerji, Bansidhar Tarai, Poonam Das, Sujeet Jha, Mohammed Faruq, Vinita Jha, Anurag Agrawal |
| hCoV-19/India/MaxCov0018_CSIR-IGIB | EPI_ISL_459920 | 5/21/2020 | Devki Devi Foundation, a unit of Max Healthcare | CSIR-IGIB/Max | Rajesh Pandey#, Samreen Siddiqui, Pooja Sharma, Bansidhar Tarai, Vivekanand A, Bharathram Uppili, Saruchi Wadhwa, Nishu Tyagi, Mitali Mukerji, Bansidhar Tarai, Poonam Das, Sujeet Jha, Mohammed Faruq, Vinita Jha, Anurag Agrawal |
| hCoV-19/India/MaxCov0019_CSIR-IGIB | EPI_ISL_459921 | 5/8/2020 | Devki Devi Foundation, a unit of Max Healthcare | CSIR-IGIB/Max | Rajesh Pandey#, Samreen Siddiqui, Pooja Sharma, Bansidhar Tarai, Vivekanand A, Bharathram Uppili, Saruchi Wadhwa, Nishu Tyagi, Mitali Mukerji, Bansidhar Tarai, Poonam Das, Sujeet Jha, Mohammed Faruq, Vinita Jha, Anurag Agrawal |
| hCoV-19/India/MaxCov0020_CSIR-IGIB | EPI_ISL_459922 | 5/10/2020 | Devki Devi Foundation, a unit of Max Healthcare | CSIR-IGIB/Max | Rajesh Pandey#, Samreen Siddiqui, Pooja Sharma, Bansidhar Tarai, Vivekanand A, Bharathram Uppili, Saruchi Wadhwa, Nishu Tyagi, Mitali Mukerji, Bansidhar Tarai, Poonam Das, Sujeet Jha, Mohammed Faruq, Vinita Jha, Anurag Agrawal |
| hCoV-19/India/MaxCov0021_CSIR-IGIB | EPI_ISL_459923 | 5/8/2020 | Devki Devi Foundation, a unit of Max Healthcare | CSIR-IGIB/Max | Rajesh Pandey#, Samreen Siddiqui, Pooja Sharma, Bansidhar Tarai, Vivekanand A, Bharathram Uppili, Saruchi Wadhwa, Nishu Tyagi, Mitali Mukerji, Bansidhar Tarai, Poonam Das, Sujeet Jha, Mohammed Faruq, Vinita Jha, Anurag Agrawal |
| hCoV-19/India/MaxCov0022_CSIR-IGIB | EPI_ISL_459924 | 5/8/2020 | Devki Devi Foundation, a unit of Max Healthcare | CSIR-IGIB/Max | Rajesh Pandey#, Samreen Siddiqui, Pooja Sharma, Bansidhar Tarai, Vivekanand A, Bharathram Uppili, Saruchi Wadhwa, Nishu Tyagi, Mitali Mukerji, Bansidhar Tarai, Poonam Das, Sujeet Jha, Mohammed Faruq, Vinita Jha, Anurag Agrawal |
| hCoV-19/India/MaxCov0024_CSIR-IGIB | EPI_ISL_459926 | 2020 | Devki Devi Foundation, a unit of Max Healthcare | CSIR-IGIB/Max | Rajesh Pandey#, Samreen Siddiqui, Pooja Sharma, Bansidhar Tarai, Vivekanand A, Bharathram Uppili, Saruchi Wadhwa, Nishu Tyagi, Mitali Mukerji, Bansidhar Tarai, Poonam Das, Sujeet Jha, Mohammed Faruq, Vinita Jha, Anurag Agrawal |
| hCoV-19/India/MaxCov0025_CSIR-IGIB | EPI_ISL_459927 | 2020 | Devki Devi Foundation, a unit of Max Healthcare | CSIR-IGIB/Max | Rajesh Pandey#, Samreen Siddiqui, Pooja Sharma, Bansidhar Tarai, Vivekanand A, Bharathram Uppili, Saruchi Wadhwa, Nishu Tyagi, Mitali Mukerji, Bansidhar Tarai, Poonam Das, Sujeet Jha, Mohammed Faruq, Vinita Jha, Anurag Agrawal |
| hCoV-19/India/MaxCov0030_CSIR-IGIB | EPI_ISL_459932 | 5/13/2020 | Devki Devi Foundation, a unit of Max Healthcare | CSIR-IGIB/Max | Rajesh Pandey#, Samreen Siddiqui, Pooja Sharma, Bansidhar Tarai, Vivekanand A, Bharathram Uppili, Saruchi Wadhwa, Nishu Tyagi, Mitali Mukerji, Bansidhar Tarai, Poonam Das, Sujeet Jha, Mohammed Faruq, Vinita Jha, Anurag Agrawal |
| hCoV-19/India/MaxCov0031_CSIR-IGIB | EPI_ISL_459933 | 5/5/2020 | Devki Devi Foundation, a unit of Max Healthcare | CSIR-IGIB/Max | Rajesh Pandey#, Samreen Siddiqui, Pooja Sharma, Bansidhar Tarai, Vivekanand A, Bharathram Uppili, Saruchi Wadhwa, Nishu Tyagi, Mitali Mukerji, Bansidhar Tarai, Poonam Das, Sujeet Jha, Mohammed Faruq, Vinita Jha, Anurag Agrawal |
| hCoV-19/India/MaxCov0032_CSIR-IGIB | EPI_ISL_459934 | 5/5/2020 | Devki Devi Foundation, a unit of Max Healthcare | CSIR-IGIB/Max | Rajesh Pandey#, Samreen Siddiqui, Pooja Sharma, Bansidhar Tarai, Vivekanand A, Bharathram Uppili, Saruchi Wadhwa, Nishu Tyagi, Mitali Mukerji, Bansidhar Tarai, Poonam Das, Sujeet Jha, Mohammed Faruq, Vinita Jha, Anurag Agrawal |
| hCoV-19/India/MaxCov0033_CSIR-IGIB | EPI_ISL_459935 | 5/6/2020 | Devki Devi Foundation, a unit of Max Healthcare | CSIR-IGIB/Max | Rajesh Pandey#, Samreen Siddiqui, Pooja Sharma, Bansidhar Tarai, Vivekanand A, Bharathram Uppili, Saruchi Wadhwa, Nishu Tyagi, Mitali Mukerji, Bansidhar Tarai, Poonam Das, Sujeet Jha, Mohammed Faruq, Vinita Jha, Anurag Agrawal |
| hCoV-19/India/MaxCov0035_CSIR-IGIB | EPI_ISL_459937 | 5/10/2020 | Devki Devi Foundation, a unit of Max Healthcare | CSIR-IGIB/Max | Rajesh Pandey#, Samreen Siddiqui, Pooja Sharma, Bansidhar Tarai, Vivekanand A, Bharathram Uppili, Saruchi Wadhwa, Nishu Tyagi, Mitali Mukerji, Bansidhar Tarai, Poonam Das, Sujeet Jha, Mohammed Faruq, Vinita Jha, Anurag Agrawal |
| hCoV-19/India/MaxCov0038_CSIR-IGIB | EPI_ISL_459940 | 5/10/2020 | Devki Devi Foundation, a unit of Max Healthcare | CSIR-IGIB/Max | Rajesh Pandey#, Samreen Siddiqui, Pooja Sharma, Bansidhar Tarai, Vivekanand A, Bharathram Uppili, Saruchi Wadhwa, Nishu Tyagi, Mitali Mukerji, Bansidhar Tarai, Poonam Das, Sujeet Jha, Mohammed Faruq, Vinita Jha, Anurag Agrawal |
| hCoV-19/India/MaxCov0039_CSIR-IGIB | EPI_ISL_459941 | 5/11/2020 | Devki Devi Foundation, a unit of Max Healthcare | CSIR-IGIB/Max | Rajesh Pandey#, Samreen Siddiqui, Pooja Sharma, Bansidhar Tarai, Vivekanand A, Bharathram Uppili, Saruchi Wadhwa, Nishu Tyagi, Mitali Mukerji, Bansidhar Tarai, Poonam Das, Sujeet Jha, Mohammed Faruq, Vinita Jha, Anurag Agrawal |
| hCoV-19/India/MaxCov0040_CSIR-IGIB | EPI_ISL_459942 | 4/27/2020 | Devki Devi Foundation, a unit of Max Healthcare | CSIR-IGIB/Max | Rajesh Pandey#, Samreen Siddiqui, Pooja Sharma, Bansidhar Tarai, Vivekanand A, Bharathram Uppili, Saruchi Wadhwa, Nishu Tyagi, Mitali Mukerji, Bansidhar Tarai, Poonam Das, Sujeet Jha, Mohammed Faruq, Vinita Jha, Anurag Agrawal |
| hCoV-19/India/MaxCov0041_CSIR-IGIB | EPI_ISL_459943 | 5/1/2020 | Devki Devi Foundation, a unit of Max Healthcare | CSIR-IGIB/Max | Rajesh Pandey#, Samreen Siddiqui, Pooja Sharma, Bansidhar Tarai, Vivekanand A, Bharathram Uppili, Saruchi Wadhwa, Nishu Tyagi, Mitali Mukerji, Bansidhar Tarai, Poonam Das, Sujeet Jha, Mohammed Faruq, Vinita Jha, Anurag Agrawal |
| hCoV-19/India/GBRC151a/2020 | EPI_ISL_461478 | 5/2/2020 | Government Medical College, Vadodara | Gujarat Biotechnology Research Centre | Fenil Patel, Nidhi Patel, Nitin Savaliya, Raghawendra Kumar, Dinesh Kumar, Zuber Saiyed, Komal Patel, Labdhi Pandya, Snehal Bagatharia, Tanuja Javadekar , R N Daveshwar, Tejas Shah, Ankit Hinsu, Pritesh Sabara, Apurvasinh Puvar, Janvi Raval, Zarna Patel, Monika Gandhi, Pinal Trivedi, Maharshi Pandya, R D Dixit, A M Kadri, Harsh Bakshi, Chaitanya Joshi, Madhvi Joshi, |
| hCoV-19/India/GBRC151b/2020 | EPI_ISL_461479 | 5/2/2020 | Government Medical College, Vadodara | Gujarat Biotechnology Research Centre | Neelam Nathani, Nitin Savaliya, Raghawendra Kumar, Dinesh Kumar, Zuber Saiyed, Komal Patel, Labdhi Pandya, Snehal Bagatharia, Tanuja Javadekar , R N Daveshwar, Tejas Shah, Ankit Hinsu, Pritesh Sabara, Apurvasinh Puvar, Janvi Raval, Zarna Patel, Monika Gandhi, Pinal Trivedi, Maharshi Pandya, Nidhi Patel, R D Dixit, A M Kadri, Harsh Bakshi, Chaitanya Joshi, Madhvi Joshi, |
| hCoV-19/India/GBRC152/2020 | EPI_ISL_461480 | 5/2/2020 | Government Medical College, Vadodara | Gujarat Biotechnology Research Centre | Armi Chaudhari, Raghawendra Kumar, Dinesh Kumar, Zuber Saiyed, Komal Patel, Labdhi Pandya, Snehal Bagatharia, Tanuja Javadekar , R N Daveshwar, Tejas Shah, Ankit Hinsu, Pritesh Sabara, Apurvasinh Puvar, Janvi Raval, Zarna Patel, Monika Gandhi, Pinal Trivedi, Maharshi Pandya, Nidhi Patel, Nitin Savaliya, R D Dixit, A M Kadri, Harsh Bakshi, Chaitanya Joshi, Madhvi Joshi, |
| hCoV-19/India/GBRC153a/2020 | EPI_ISL_461481 | 4/27/2020 | Pandit Deendayal Upadhyay Government Medical College, Rajkot | Gujarat Biotechnology Research Centre | Bhavya Jindal, Dinesh Kumar, Zuber Saiyed, Komal Patel, Labdhi Pandya, Snehal Bagatharia, Prakash Modi, Sejul Antala, Manish Pattani, Tejas Shah, Ankit Hinsu, Pritesh Sabara, Apurvasinh Puvar, Janvi Raval, Zarna Patel, Monika Gandhi, Pinal Trivedi, Maharshi Pandya, Nidhi Patel, Nitin Savaliya, Raghawendra Kumar, R D Dixit, A M Kadri, Harsh Bakshi, Chaitanya Joshi, Madhvi Joshi |
| hCoV-19/India/GBRC153b/2020 | EPI_ISL_461482 | 4/27/2020 | Pandit Deendayal Upadhyay Government Medical College, Rajkot | Gujarat Biotechnology Research Centre | Anjali Rajwar, Zuber Saiyed, Komal Patel, Labdhi Pandya, Snehal Bagatharia, Prakash Modi, Sejul Antala, Manish Pattani, Tejas Shah, Ankit Hinsu, Pritesh Sabara, Apurvasinh Puvar, Janvi Raval, Zarna Patel, Monika Gandhi, Pinal Trivedi, Maharshi Pandya, Nidhi Patel, Nitin Savaliya, Raghawendra Kumar, Dinesh Kumar, R D Dixit, A M Kadri, Harsh Bakshi, Chaitanya Joshi, Madhvi Joshi |
| hCoV-19/India/GBRC154/2020 | EPI_ISL_461483 | 5/27/2020 | B.J. Medical College and Civil hospital | Gujarat Biotechnology Research Centre | Dipeshwari Shewale, Komal Patel, Labdhi Pandya, Snehal Bagatharia, Pranay Shah, Kamlesh J Upadhyay, Tejas Shah, Ankit Hinsu, Pritesh Sabara, Apurvasinh Puvar, Janvi Raval, Zarna Patel, Monika Gandhi, Pinal Trivedi, Maharshi Pandya, Nidhi Patel, Nitin Savaliya, Raghawendra Kumar, Dinesh Kumar, Zuber Saiyed, R D Dixit, A M Kadri, Harsh Bakshi, Chaitanya Joshi, Madhvi Joshi, |
| hCoV-19/India/GBRC155/2020 | EPI_ISL_461484 | 5/27/2020 | B.J. Medical College and Civil hospital | Gujarat Biotechnology Research Centre | Priyanka P Vatsa, Labdhi Pandya, Snehal Bagatharia, Pranay Shah, Kamlesh J Upadhyay, Tejas Shah, Ankit Hinsu, Pritesh Sabara, Apurvasinh Puvar, Janvi Raval, Zarna Patel, Monika Gandhi, Pinal Trivedi, Maharshi Pandya, Nidhi Patel, Nitin Savaliya, Raghawendra Kumar, Dinesh Kumar, Zuber Saiyed, Komal Patel, R D Dixit, A M Kadri, Harsh Bakshi, Chaitanya Joshi, Madhvi Joshi, |
| hCoV-19/India/GBRC156/2020 | EPI_ISL_461485 | 5/27/2020 | B.J. Medical College and Civil hospital | Gujarat Biotechnology Research Centre | Pooja P Doshi, Snehal Bagatharia, Pranay Shah, Kamlesh J Upadhyay, Tejas Shah, Ankit Hinsu, Pritesh Sabara, Apurvasinh Puvar, Janvi Raval, Zarna Patel, Monika Gandhi, Pinal Trivedi, Maharshi Pandya, Nidhi Patel, Nitin Savaliya, Raghawendra Kumar, Dinesh Kumar, Zuber Saiyed, Komal Patel, Labdhi Pandya, R D Dixit, A M Kadri, Harsh Bakshi, Chaitanya Joshi, Madhvi Joshi, |
| hCoV-19/India/GBRC157a/2020 | EPI_ISL_461486 | 5/27/2020 | B.J. Medical College and Civil hospital | Gujarat Biotechnology Research Centre | Akanksha Verma, Pranay Shah, Kamlesh J Upadhyay, Tejas Shah, Ankit Hinsu, Pritesh Sabara, Apurvasinh Puvar, Janvi Raval, Zarna Patel, Monika Gandhi, Pinal Trivedi, Maharshi Pandya, Nidhi Patel, Nitin Savaliya, Raghawendra Kumar, Dinesh Kumar, Zuber Saiyed, Komal Patel, Labdhi Pandya, Snehal Bagatharia, R D Dixit, A M Kadri, Harsh Bakshi, Chaitanya Joshi, Madhvi Joshi, |
| hCoV-19/India/GBRC157b/2020 | EPI_ISL_461487 | 5/27/2020 | B.J. Medical College and Civil hospital | Gujarat Biotechnology Research Centre | Priti Pandita, Kamlesh J Upadhyay, Tejas Shah, Ankit Hinsu, Pritesh Sabara, Apurvasinh Puvar, Janvi Raval, Zarna Patel, Monika Gandhi, Pinal Trivedi, Maharshi Pandya, Nidhi Patel, Nitin Savaliya, Raghawendra Kumar, Dinesh Kumar, Zuber Saiyed, Komal Patel, Labdhi Pandya, Snehal Bagatharia, Pranay Shah, R D Dixit, A M Kadri, Harsh Bakshi, Chaitanya Joshi, Madhvi Joshi, |
| hCoV-19/India/GBRC158/2020 | EPI_ISL_461488 | 5/27/2020 | B.J. Medical College and Civil hospital | Gujarat Biotechnology Research Centre | Pragya Sharma, Tejas Shah, Ankit Hinsu, Pritesh Sabara, Apurvasinh Puvar, Janvi Raval, Zarna Patel, Monika Gandhi, Pinal Trivedi, Maharshi Pandya, Nidhi Patel, Nitin Savaliya, Raghawendra Kumar, Dinesh Kumar, Zuber Saiyed, Komal Patel, Labdhi Pandya, Snehal Bagatharia, Pranay Shah, Kamlesh J Upadhyay, R D Dixit, A M Kadri, Harsh Bakshi, Chaitanya Joshi, Madhvi Joshi, |
| hCoV-19/India/GBRC159a/2020 | EPI_ISL_461489 | 5/27/2020 | B.J. Medical College and Civil hospital | Gujarat Biotechnology Research Centre | Neha Rajpara, Ankit Hinsu, Pritesh Sabara, Apurvasinh Puvar, Janvi Raval, Zarna Patel, Monika Gandhi, Pinal Trivedi, Maharshi Pandya, Nidhi Patel, Nitin Savaliya, Raghawendra Kumar, Dinesh Kumar, Zuber Saiyed, Komal Patel, Labdhi Pandya, Snehal Bagatharia, Pranay Shah, Kamlesh J Upadhyay, Tejas Shah, R D Dixit, A M Kadri, Harsh Bakshi, Chaitanya Joshi, Madhvi Joshi, |
| hCoV-19/India/GBRC159b/2020 | EPI_ISL_461490 | 5/27/2020 | B.J. Medical College and Civil hospital | Gujarat Biotechnology Research Centre | Afzal Ansari, Pritesh Sabara, Apurvasinh Puvar, Janvi Raval, Zarna Patel, Monika Gandhi, Pinal Trivedi, Maharshi Pandya, Nidhi Patel, Nitin Savaliya, Raghawendra Kumar, Dinesh Kumar, Zuber Saiyed, Komal Patel, Labdhi Pandya, Snehal Bagatharia, Pranay Shah, Kamlesh J Upadhyay, Tejas Shah, Ankit Hinsu, R D Dixit, A M Kadri, Harsh Bakshi, Chaitanya Joshi, Madhvi Joshi, |
| hCoV-19/India/GBRC160a/2020 | EPI_ISL_461491 | 5/27/2020 | B.J. Medical College and Civil hospital | Gujarat Biotechnology Research Centre | Fenil Patel, Apurvasinh Puvar, Janvi Raval, Zarna Patel, Monika Gandhi, Pinal Trivedi, Maharshi Pandya, Nidhi Patel, Nitin Savaliya, Raghawendra Kumar, Dinesh Kumar, Zuber Saiyed, Komal Patel, Labdhi Pandya, Snehal Bagatharia, Pranay Shah, Kamlesh J Upadhyay, Tejas Shah, Ankit Hinsu, Pritesh Sabara, R D Dixit, A M Kadri, Harsh Bakshi, Chaitanya Joshi, Madhvi Joshi, |
| hCoV-19/India/GBRC160b/2020 | EPI_ISL_461492 | 5/27/2020 | B.J. Medical College and Civil hospital | Gujarat Biotechnology Research Centre | Neelam Nathani, Janvi Raval, Zarna Patel, Monika Gandhi, Pinal Trivedi, Maharshi Pandya, Nidhi Patel, Nitin Savaliya, Raghawendra Kumar, Dinesh Kumar, Zuber Saiyed, Komal Patel, Labdhi Pandya, Snehal Bagatharia, Pranay Shah, Kamlesh J Upadhyay, Tejas Shah, Ankit Hinsu, Pritesh Sabara, Apurvasinh Puvar, R D Dixit, A M Kadri, Harsh Bakshi, Chaitanya Joshi, Madhvi Joshi, |
| hCoV-19/India/GBRC161a/2020 | EPI_ISL_461493 | 5/27/2020 | B.J. Medical College and Civil hospital | Gujarat Biotechnology Research Centre | Armi Chaudhari, Zarna Patel, Monika Gandhi, Pinal Trivedi, Maharshi Pandya, Nidhi Patel, Nitin Savaliya, Raghawendra Kumar, Dinesh Kumar, Zuber Saiyed, Komal Patel, Labdhi Pandya, Snehal Bagatharia, Pranay Shah, Kamlesh J Upadhyay, Tejas Shah, Ankit Hinsu, Pritesh Sabara, Apurvasinh Puvar, Janvi Raval, R D Dixit, A M Kadri, Harsh Bakshi, Chaitanya Joshi, Madhvi Joshi, |
| hCoV-19/India/GBRC161b/2020 | EPI_ISL_461494 | 5/27/2020 | B.J. Medical College and Civil hospital | Gujarat Biotechnology Research Centre | Bhavya Jindal, Monika Gandhi, Pinal Trivedi, Maharshi Pandya, Nidhi Patel, Nitin Savaliya, Raghawendra Kumar, Dinesh Kumar, Zuber Saiyed, Komal Patel, Labdhi Pandya, Snehal Bagatharia, Pranay Shah, Kamlesh J Upadhyay, Tejas Shah, Ankit Hinsu, Pritesh Sabara, Apurvasinh Puvar, Janvi Raval, Zarna Patel, R D Dixit, A M Kadri, Harsh Bakshi, Chaitanya Joshi, Madhvi Joshi, |
| hCoV-19/India/GBRC162a/2020 | EPI_ISL_461495 | 5/27/2020 | B.J. Medical College and Civil hospital | Gujarat Biotechnology Research Centre | Anjali Rajwar, Pinal Trivedi, Maharshi Pandya, Nidhi Patel, Nitin Savaliya, Raghawendra Kumar, Dinesh Kumar, Zuber Saiyed, Komal Patel, Labdhi Pandya, Snehal Bagatharia, Pranay Shah, Kamlesh J Upadhyay, Tejas Shah, Ankit Hinsu, Pritesh Sabara, Apurvasinh Puvar, Janvi Raval, Zarna Patel, Monika Gandhi, R D Dixit, A M Kadri, Harsh Bakshi, Chaitanya Joshi, Madhvi Joshi, |
| hCoV-19/India/GBRC162b/2020 | EPI_ISL_461496 | 5/27/2020 | B.J. Medical College and Civil hospital | Gujarat Biotechnology Research Centre | Dipeshwari Shewale, Maharshi Pandya, Nidhi Patel, Nitin Savaliya, Raghawendra Kumar, Dinesh Kumar, Zuber Saiyed, Komal Patel, Labdhi Pandya, Snehal Bagatharia, Pranay Shah, Kamlesh J Upadhyay, Tejas Shah, Ankit Hinsu, Pritesh Sabara, Apurvasinh Puvar, Janvi Raval, Zarna Patel, Monika Gandhi, Pinal Trivedi, R D Dixit, A M Kadri, Harsh Bakshi, Chaitanya Joshi, Madhvi Joshi, |
| hCoV-19/India/GBRC163a/2020 | EPI_ISL_461497 | 5/27/2020 | B.J. Medical College and Civil hospital | Gujarat Biotechnology Research Centre | Priyanka P Vatsa, Nidhi Patel, Nitin Savaliya, Raghawendra Kumar, Dinesh Kumar, Zuber Saiyed, Komal Patel, Labdhi Pandya, Snehal Bagatharia, Pranay Shah, Kamlesh J Upadhyay, Tejas Shah, Ankit Hinsu, Pritesh Sabara, Apurvasinh Puvar, Janvi Raval, Zarna Patel, Monika Gandhi, Pinal Trivedi, Maharshi Pandya, R D Dixit, A M Kadri, Harsh Bakshi, Chaitanya Joshi, Madhvi Joshi, |
| hCoV-19/India/GBRC163b/2020 | EPI_ISL_461498 | 5/27/2020 | B.J. Medical College and Civil hospital | Gujarat Biotechnology Research Centre | Pooja P Doshi, Nitin Savaliya, Raghawendra Kumar, Dinesh Kumar, Zuber Saiyed, Komal Patel, Labdhi Pandya, Snehal Bagatharia, Pranay Shah, Kamlesh J Upadhyay, Tejas Shah, Ankit Hinsu, Pritesh Sabara, Apurvasinh Puvar, Janvi Raval, Zarna Patel, Monika Gandhi, Pinal Trivedi, Maharshi Pandya, Nidhi Patel, R D Dixit, A M Kadri, Harsh Bakshi, Chaitanya Joshi, Madhvi Joshi, |
| hCoV-19/India/GBRC164/2020 | EPI_ISL_461499 | 5/27/2020 | B.J. Medical College and Civil hospital | Gujarat Biotechnology Research Centre | Akanksha Verma, Raghawendra Kumar, Dinesh Kumar, Zuber Saiyed, Komal Patel, Labdhi Pandya, Snehal Bagatharia, Pranay Shah, Kamlesh J Upadhyay, Tejas Shah, Ankit Hinsu, Pritesh Sabara, Apurvasinh Puvar, Janvi Raval, Zarna Patel, Monika Gandhi, Pinal Trivedi, Maharshi Pandya, Nidhi Patel, Nitin Savaliya, R D Dixit, A M Kadri, Harsh Bakshi, Chaitanya Joshi, Madhvi Joshi, |
| hCoV-19/India/GBRC165/2020 | EPI_ISL_461500 | 5/27/2020 | B.J. Medical College and Civil hospital | Gujarat Biotechnology Research Centre | Priti Pandita, Dinesh Kumar, Zuber Saiyed, Komal Patel, Labdhi Pandya, Snehal Bagatharia, Pranay Shah, Kamlesh J Upadhyay, Tejas Shah, Ankit Hinsu, Pritesh Sabara, Apurvasinh Puvar, Janvi Raval, Zarna Patel, Monika Gandhi, Pinal Trivedi, Maharshi Pandya, Nidhi Patel, Nitin Savaliya, Raghawendra Kumar, R D Dixit, A M Kadri, Harsh Bakshi, Chaitanya Joshi, Madhvi Joshi, |
| hCoV-19/India/GBRC166a/2020 | EPI_ISL_461501 | 5/27/2020 | B.J. Medical College and Civil hospital | Gujarat Biotechnology Research Centre | Pragya Sharma, Zuber Saiyed, Komal Patel, Labdhi Pandya, Snehal Bagatharia, Pranay Shah, Kamlesh J Upadhyay, Tejas Shah, Ankit Hinsu, Pritesh Sabara, Apurvasinh Puvar, Janvi Raval, Zarna Patel, Monika Gandhi, Pinal Trivedi, Maharshi Pandya, Nidhi Patel, Nitin Savaliya, Raghawendra Kumar, Dinesh Kumar, R D Dixit, A M Kadri, Harsh Bakshi, Chaitanya Joshi, Madhvi Joshi, |
| hCoV-19/India/GBRC166b/2020 | EPI_ISL_461502 | 5/27/2020 | B.J. Medical College and Civil hospital | Gujarat Biotechnology Research Centre | Neha Rajpara, Komal Patel, Labdhi Pandya, Snehal Bagatharia, Pranay Shah, Kamlesh J Upadhyay, Tejas Shah, Ankit Hinsu, Pritesh Sabara, Apurvasinh Puvar, Janvi Raval, Zarna Patel, Monika Gandhi, Pinal Trivedi, Maharshi Pandya, Nidhi Patel, Nitin Savaliya, Raghawendra Kumar, Dinesh Kumar, Zuber Saiyed, R D Dixit, A M Kadri, Harsh Bakshi, Chaitanya Joshi, Madhvi Joshi, |
| hCoV-19/India/GBRC167a/2020 | EPI_ISL_461503 | 5/27/2020 | B.J. Medical College and Civil hospital | Gujarat Biotechnology Research Centre | Afzal Ansari, Labdhi Pandya, Snehal Bagatharia, Pranay Shah, Kamlesh J Upadhyay, Tejas Shah, Ankit Hinsu, Pritesh Sabara, Apurvasinh Puvar, Janvi Raval, Zarna Patel, Monika Gandhi, Pinal Trivedi, Maharshi Pandya, Nidhi Patel, Nitin Savaliya, Raghawendra Kumar, Dinesh Kumar, Zuber Saiyed, Komal Patel, R D Dixit, A M Kadri, Harsh Bakshi, Chaitanya Joshi, Madhvi Joshi, |
| hCoV-19/India/GBRC167b/2020 | EPI_ISL_461504 | 5/27/2020 | B.J. Medical College and Civil hospital | Gujarat Biotechnology Research Centre | Snehal Bagatharia, Pranay Shah, Kamlesh J Upadhyay, Tejas Shah, Ankit Hinsu, Pritesh Sabara, Apurvasinh Puvar, Janvi Raval, Zarna Patel, Monika Gandhi, Pinal Trivedi, Maharshi Pandya, Nidhi Patel, Nitin Savaliya, Raghawendra Kumar, Dinesh Kumar, Zuber Saiyed, Komal Patel, Labdhi Pandya, Fenil Patel, R D Dixit, A M Kadri, Harsh Bakshi, Chaitanya Joshi, Madhvi Joshi, |
| hCoV-19/India/GBRC168a/2020 | EPI_ISL_461505 | 5/27/2020 | B.J. Medical College and Civil hospital | Gujarat Biotechnology Research Centre | Pranay Shah, Kamlesh J Upadhyay, Tejas Shah, Ankit Hinsu, Pritesh Sabara, Apurvasinh Puvar, Janvi Raval, Zarna Patel, Monika Gandhi, Pinal Trivedi, Maharshi Pandya, Nidhi Patel, Nitin Savaliya, Raghawendra Kumar, Dinesh Kumar, Zuber Saiyed, Komal Patel, Labdhi Pandya, Snehal Bagatharia, Neelam Nathani, R D Dixit, A M Kadri, Harsh Bakshi, Chaitanya Joshi, Madhvi Joshi, |
| hCoV-19/India/GBRC168b/2020 | EPI_ISL_461506 | 5/27/2020 | B.J. Medical College and Civil hospital | Gujarat Biotechnology Research Centre | Kamlesh J Upadhyay, Tejas Shah, Ankit Hinsu, Pritesh Sabara, Apurvasinh Puvar, Janvi Raval, Zarna Patel, Monika Gandhi, Pinal Trivedi, Maharshi Pandya, Nidhi Patel, Nitin Savaliya, Raghawendra Kumar, Dinesh Kumar, Zuber Saiyed, Komal Patel, Labdhi Pandya, Snehal Bagatharia, Pranay Shah, Armi Chaudhari, R D Dixit, A M Kadri, Harsh Bakshi, Chaitanya Joshi, Madhvi Joshi, |
| hCoV-19/India/ILSCV11165/2020 | EPI_ISL_463010 | 5/7/2020 | Institute of Life Sciences, Bhubaneswar | Immunogenomics lab, Institute of Life Sciences, Bhubaneswar | Sunil Raghav, Arup Ghosh, Atimukta Jha, Viplov K. Biswas, Swati Madhulika, Manasi Priyadarshini, Shuchi Smita, Kaushik Sen, Hiren G. Dodia, Deepak Singh, Jeky Chawla, Shamima Ansari, Rupesh Dash, Soma Chattopadhyay, Ghulam Hussain Syed, Shanti Senapati, Tushar K. Beuria, Rajeeb Swain, Punit Prasad, ILS COVID-19 TEAM, Orissa COVID-19 Study Group, Ajay Parida |
| hCoV-19/India/ILSCV12742/2020 | EPI_ISL_463013 | 5/9/2020 | Institute of Life Sciences, Bhubaneswar | Immunogenomics lab, Institute of Life Sciences, Bhubaneswar | Sunil Raghav, Arup Ghosh, Atimukta Jha, Viplov K. Biswas, Swati Madhulika, Manasi Priyadarshini, Shuchi Smita, Kaushik Sen, Hiren G. Dodia, Deepak Singh, Jeky Chawla, Shamima Ansari, Rupesh Dash, Soma Chattopadhyay, Ghulam Hussain Syed, Shanti Senapati, Tushar K. Beuria, Rajeeb Swain, Punit Prasad, ILS COVID-19 TEAM, Orissa COVID-19 Study Group, Ajay Parida |
| hCoV-19/India/ILSCV12743/2020 | EPI_ISL_463014 | 5/9/2020 | Institute of Life Sciences, Bhubaneswar | Immunogenomics lab, Institute of Life Sciences, Bhubaneswar | Sunil Raghav, Arup Ghosh, Atimukta Jha, Viplov K. Biswas, Swati Madhulika, Manasi Priyadarshini, Shuchi Smita, Kaushik Sen, Hiren G. Dodia, Deepak Singh, Jeky Chawla, Shamima Ansari, Rupesh Dash, Soma Chattopadhyay, Ghulam Hussain Syed, Shanti Senapati, Tushar K. Beuria, Rajeeb Swain, Punit Prasad, ILS COVID-19 TEAM, Orissa COVID-19 Study Group, Ajay Parida |

| | | | | | |
|---|---|---|---|---|---|
| hCoV-19/India/ILSCV12752/2020 | EPI_ISL_463017 | 5/9/2020 | Institute of Life Sciences, Bhubaneswar | Immunogenomics lab, Institute of Life Sciences, Bhubaneswar | Sunil Raghav, Arup Ghosh, Atimukta Jha, Viplov K. Biswas, Swati Madhulika, Manasi Priyadarshini, Shuchi Smita, Kaushik Sen, Hiren G. Dodia, Deepak Singh, Jeky Chawla, Shamima Ansari, Rupesh Dash, Soma Chattopadhyay, Ghulam Hussain Syed, Shanti Senapati, Tushar K. Beuria, Rajeeb Swain, Punit Prasad, ILS COVID-19 TEAM, Orissa COVID-19 Study Group, Ajay Parida |
| hCoV-19/India/ILSCV12762/2020 | EPI_ISL_463018 | 5/9/2020 | Institute of Life Sciences, Bhubaneswar | Immunogenomics lab, Institute of Life Sciences, Bhubaneswar | Sunil Raghav, Arup Ghosh, Atimukta Jha, Viplov K. Biswas, Swati Madhulika, Manasi Priyadarshini, Shuchi Smita, Kaushik Sen, Hiren G. Dodia, Deepak Singh, Jeky Chawla, Shamima Ansari, Rupesh Dash, Soma Chattopadhyay, Ghulam Hussain Syed, Shanti Senapati, Tushar K. Beuria, Rajeeb Swain, Punit Prasad, ILS COVID-19 TEAM, Orissa COVID-19 Study Group, Ajay Parida |
| hCoV-19/India/ILSCV13676/2020 | EPI_ISL_463019 | 5/9/2020 | Institute of Life Sciences, Bhubaneswar | Immunogenomics lab, Institute of Life Sciences, Bhubaneswar | Sunil Raghav, Arup Ghosh, Atimukta Jha, Viplov K. Biswas, Swati Madhulika, Manasi Priyadarshini, Shuchi Smita, Kaushik Sen, Hiren G. Dodia, Deepak Singh, Jeky Chawla, Shamima Ansari, Rupesh Dash, Soma Chattopadhyay, Ghulam Hussain Syed, Shanti Senapati, Tushar K. Beuria, Rajeeb Swain, Punit Prasad, ILS COVID-19 TEAM, Orissa COVID-19 Study Group, Ajay Parida |
| hCoV-19/India/ILSCV13715/2020 | EPI_ISL_463022 | 5/9/2020 | Institute of Life Sciences, Bhubaneswar | Immunogenomics lab, Institute of Life Sciences, Bhubaneswar | Sunil Raghav, Arup Ghosh, Atimukta Jha, Viplov K. Biswas, Swati Madhulika, Manasi Priyadarshini, Shuchi Smita, Kaushik Sen, Hiren G. Dodia, Deepak Singh, Jeky Chawla, Shamima Ansari, Rupesh Dash, Soma Chattopadhyay, Ghulam Hussain Syed, Shanti Senapati, Tushar K. Beuria, Rajeeb Swain, Punit Prasad, ILS COVID-19 TEAM, Orissa COVID-19 Study Group, Ajay Parida |
| hCoV-19/India/ILSCV13907/2020 | EPI_ISL_463024 | 5/10/2020 | Institute of Life Sciences, Bhubaneswar | Immunogenomics lab, Institute of Life Sciences, Bhubaneswar | Sunil Raghav, Arup Ghosh, Atimukta Jha, Viplov K. Biswas, Swati Madhulika, Manasi Priyadarshini, Shuchi Smita, Kaushik Sen, Hiren G. Dodia, Deepak Singh, Jeky Chawla, Shamima Ansari, Rupesh Dash, Soma Chattopadhyay, Ghulam Hussain Syed, Shanti Senapati, Tushar K. Beuria, Rajeeb Swain, Punit Prasad, ILS COVID-19 TEAM, Orissa COVID-19 Study Group, Ajay Parida |
| hCoV-19/India/ILSCV13921/2020 | EPI_ISL_463025 | 5/10/2020 | Institute of Life Sciences, Bhubaneswar | Immunogenomics lab, Institute of Life Sciences, Bhubaneswar | Sunil Raghav, Arup Ghosh, Atimukta Jha, Viplov K. Biswas, Swati Madhulika, Manasi Priyadarshini, Shuchi Smita, Kaushik Sen, Hiren G. Dodia, Deepak Singh, Jeky Chawla, Shamima Ansari, Rupesh Dash, Soma Chattopadhyay, Ghulam Hussain Syed, Shanti Senapati, Tushar K. Beuria, Rajeeb Swain, Punit Prasad, ILS COVID-19 TEAM, Orissa COVID-19 Study Group, Ajay Parida |
| hCoV-19/India/ILSCV13929/2020 | EPI_ISL_463026 | 5/10/2020 | Institute of Life Sciences, Bhubaneswar | Immunogenomics lab, Institute of Life Sciences, Bhubaneswar | Sunil Raghav, Arup Ghosh, Atimukta Jha, Viplov K. Biswas, Swati Madhulika, Manasi Priyadarshini, Shuchi Smita, Kaushik Sen, Hiren G. Dodia, Deepak Singh, Jeky Chawla, Shamima Ansari, Rupesh Dash, Soma Chattopadhyay, Ghulam Hussain Syed, Shanti Senapati, Tushar K. Beuria, Rajeeb Swain, Punit Prasad, ILS COVID-19 TEAM, Orissa COVID-19 Study Group, Ajay Parida |
| hCoV-19/India/ILSCV14004/2020 | EPI_ISL_463027 | 5/9/2020 | Institute of Life Sciences, Bhubaneswar | Immunogenomics lab, Institute of Life Sciences, Bhubaneswar | Sunil Raghav, Arup Ghosh, Atimukta Jha, Viplov K. Biswas, Swati Madhulika, Manasi Priyadarshini, Shuchi Smita, Kaushik Sen, Hiren G. Dodia, Deepak Singh, Jeky Chawla, Shamima Ansari, Rupesh Dash, Soma Chattopadhyay, Ghulam Hussain Syed, Shanti Senapati, Tushar K. Beuria, Rajeeb Swain, Punit Prasad, ILS COVID-19 TEAM, Orissa COVID-19 Study Group, Ajay Parida |
| hCoV-19/India/ILSCV14262/2020 | EPI_ISL_463029 | 5/11/2020 | Institute of Life Sciences, Bhubaneswar | Immunogenomics lab, Institute of Life Sciences, Bhubaneswar | Sunil Raghav, Arup Ghosh, Atimukta Jha, Viplov K. Biswas, Swati Madhulika, Manasi Priyadarshini, Shuchi Smita, Kaushik Sen, Hiren G. Dodia, Deepak Singh, Jeky Chawla, Shamima Ansari, Rupesh Dash, Soma Chattopadhyay, Ghulam Hussain Syed, Shanti Senapati, Tushar K. Beuria, Rajeeb Swain, Punit Prasad, ILS COVID-19 TEAM, Orissa COVID-19 Study Group, Ajay Parida |
| hCoV-19/India/ILSCV14374/2020 | EPI_ISL_463031 | 5/10/2020 | Institute of Life Sciences, Bhubaneswar | Immunogenomics lab, Institute of Life Sciences, Bhubaneswar | Sunil Raghav, Arup Ghosh, Atimukta Jha, Viplov K. Biswas, Swati Madhulika, Manasi Priyadarshini, Shuchi Smita, O. P. Shriwas, Priyanka Mohapatra, Satya Ranjan Sahu, Aliva Minz, Debyashrita Barik, Rupesh Dash, Soma Chattopadhyay, Ghulam Hussain Syed, Shanti Senapati, Tushar K. Beuria, Rajeeb Swain, Punit Prasad, ILS COVID-19 TEAM, Orissa COVID-19 Study Group, Ajay Parida |
| hCoV-19/India/ILSCV14743/2020 | EPI_ISL_463032 | 5/10/2020 | Institute of Life Sciences, Bhubaneswar | Immunogenomics lab, Institute of Life Sciences, Bhubaneswar | Sunil Raghav, Arup Ghosh, Atimukta Jha, Viplov K. Biswas, Swati Madhulika, Manasi Priyadarshini, Shuchi Smita, O. P. Shriwas, Priyanka Mohapatra, Satya Ranjan Sahu, Aliva Minz, Debyashrita Barik, Rupesh Dash, Soma Chattopadhyay, Ghulam Hussain Syed, Shanti Senapati, Tushar K. Beuria, Rajeeb Swain, Punit Prasad, ILS COVID-19 TEAM, Orissa COVID-19 Study Group, Ajay Parida |
| hCoV-19/India/ILSCV14831/2020 | EPI_ISL_463033 | 5/10/2020 | Institute of Life Sciences, Bhubaneswar | Immunogenomics lab, Institute of Life Sciences, Bhubaneswar | Sunil Raghav, Arup Ghosh, Atimukta Jha, Viplov K. Biswas, Swati Madhulika, Manasi Priyadarshini, Shuchi Smita, O. P. Shriwas, Priyanka Mohapatra, Satya Ranjan Sahu, Aliva Minz, Debyashrita Barik, Rupesh Dash, Soma Chattopadhyay, Ghulam Hussain Syed, Shanti Senapati, Tushar K. Beuria, Rajeeb Swain, Punit Prasad, ILS COVID-19 TEAM, Orissa COVID-19 Study Group, Ajay Parida |
| hCoV-19/India/ILSCV14847/2020 | EPI_ISL_463034 | 5/10/2020 | Institute of Life Sciences, Bhubaneswar | Immunogenomics lab, Institute of Life Sciences, Bhubaneswar | Sunil Raghav, Arup Ghosh, Atimukta Jha, Viplov K. Biswas, Swati Madhulika, Manasi Priyadarshini, Shuchi Smita, O. P. Shriwas, Priyanka Mohapatra, Satya Ranjan Sahu, Aliva Minz, Debyashrita Barik, Rupesh Dash, Soma Chattopadhyay, Ghulam Hussain Syed, Shanti Senapati, Tushar K. Beuria, Rajeeb Swain, Punit Prasad, ILS COVID-19 TEAM, Orissa COVID-19 Study Group, Ajay Parida |
| hCoV-19/India/ILSCV15547/2020 | EPI_ISL_463036 | 5/11/2020 | Institute of Life Sciences, Bhubaneswar | Immunogenomics lab, Institute of Life Sciences, Bhubaneswar | Sunil Raghav, Arup Ghosh, Atimukta Jha, Viplov K. Biswas, Swati Madhulika, Manasi Priyadarshini, Shuchi Smita, O. P. Shriwas, Priyanka Mohapatra, Satya Ranjan Sahu, Aliva Minz, Debyashrita Barik, Rupesh Dash, Soma Chattopadhyay, Ghulam Hussain Syed, Shanti Senapati, Tushar K. Beuria, Rajeeb Swain, Punit Prasad, ILS COVID-19 TEAM, Orissa COVID-19 Study Group, Ajay Parida |
| hCoV-19/India/ILSCV15552/2020 | EPI_ISL_463037 | 5/11/2020 | Institute of Life Sciences, Bhubaneswar | Immunogenomics lab, Institute of Life Sciences, Bhubaneswar | Sunil Raghav, Arup Ghosh, Atimukta Jha, Viplov K. Biswas, Swati Madhulika, Manasi Priyadarshini, Shuchi Smita, O. P. Shriwas, Priyanka Mohapatra, Satya Ranjan Sahu, Aliva Minz, Debyashrita Barik, Rupesh Dash, Soma Chattopadhyay, Ghulam Hussain Syed, Shanti Senapati, Tushar K. Beuria, Rajeeb Swain, Punit Prasad, ILS COVID-19 TEAM, Orissa COVID-19 Study Group, Ajay Parida |
| hCoV-19/India/ILSCV15907/2020 | EPI_ISL_463038 | 5/11/2020 | Institute of Life Sciences, Bhubaneswar | Immunogenomics lab, Institute of Life Sciences, Bhubaneswar | Sunil Raghav, Arup Ghosh, Atimukta Jha, Viplov K. Biswas, Swati Madhulika, Manasi Priyadarshini, Shuchi Smita, O. P. Shriwas, Priyanka Mohapatra, Satya Ranjan Sahu, Aliva Minz, Debyashrita Barik, Rupesh Dash, Soma Chattopadhyay, Ghulam Hussain Syed, Shanti Senapati, Tushar K. Beuria, Rajeeb Swain, Punit Prasad, ILS COVID-19 TEAM, Orissa COVID-19 Study Group, Ajay Parida |
| hCoV-19/India/ILSCV15917/2020 | EPI_ISL_463039 | 5/11/2020 | Institute of Life Sciences, Bhubaneswar | Immunogenomics lab, Institute of Life Sciences, Bhubaneswar | Sunil Raghav, Arup Ghosh, Atimukta Jha, Viplov K. Biswas, Swati Madhulika, Manasi Priyadarshini, Shuchi Smita, O. P. Shriwas, Priyanka Mohapatra, Satya Ranjan Sahu, Aliva Minz, Debyashrita Barik, Rupesh Dash, Soma Chattopadhyay, Ghulam Hussain Syed, Shanti Senapati, Tushar K. Beuria, Rajeeb Swain, Punit Prasad, ILS COVID-19 TEAM, Orissa COVID-19 Study Group, Ajay Parida |
| hCoV-19/India/ILSCV16509/2020 | EPI_ISL_463049 | 5/12/2020 | Institute of Life Sciences, Bhubaneswar | Immunogenomics lab, Institute of Life Sciences, Bhubaneswar | Sunil Raghav, Arup Ghosh, Atimukta Jha, Viplov K. Biswas, Swati Madhulika, Manasi Priyadarshini, Shuchi Smita, O. P. Shriwas, Priyanka Mohapatra, Satya Ranjan Sahu, Aliva Minz, Debyashrita Barik, Rupesh Dash, Soma Chattopadhyay, Ghulam Hussain Syed, Shanti Senapati, Tushar K. Beuria, Rajeeb Swain, Punit Prasad, ILS COVID-19 TEAM, Orissa COVID-19 Study Group, Ajay Parida |
| hCoV-19/India/ILSCV16512/2020 | EPI_ISL_463051 | 5/12/2020 | Institute of Life Sciences, Bhubaneswar | Immunogenomics lab, Institute of Life Sciences, Bhubaneswar | Sunil Raghav, Arup Ghosh, Atimukta Jha, Viplov K. Biswas, Swati Madhulika, Manasi Priyadarshini, Shuchi Smita, O. P. Shriwas, Priyanka Mohapatra, Satya Ranjan Sahu, Aliva Minz, Debyashrita Barik, Rupesh Dash, Soma Chattopadhyay, Ghulam Hussain Syed, Shanti Senapati, Tushar K. Beuria, Rajeeb Swain, Punit Prasad, ILS COVID-19 TEAM, Orissa COVID-19 Study Group, Ajay Parida |
| hCoV-19/India/ILSCV16514/2020 | EPI_ISL_463052 | 5/12/2020 | Institute of Life Sciences, Bhubaneswar | Immunogenomics lab, Institute of Life Sciences, Bhubaneswar | Sunil Raghav, Arup Ghosh, Atimukta Jha, Viplov K. Biswas, Swati Madhulika, Manasi Priyadarshini, Shuchi Smita, Sifu Agarwal, Sanchari Chatterjee, Avula Kiran, Parej Nath, Supriya Suman, Rina Yadav, Rupesh Dash, Soma Chattopadhyay, Ghulam Hussain Syed, Shanti Senapati, Tushar K. Beuria, Rajeeb Swain, Punit Prasad, ILS COVID-19 TEAM, Orissa COVID-19 Study Group, Ajay Parida |
| hCoV-19/India/ILSCV16809/2020 | EPI_ISL_463054 | 5/11/2020 | Institute of Life Sciences, Bhubaneswar | Immunogenomics lab, Institute of Life Sciences, Bhubaneswar | Sunil Raghav, Arup Ghosh, Atimukta Jha, Viplov K. Biswas, Swati Madhulika, Manasi Priyadarshini, Shuchi Smita, Sifu Agarwal, Sanchari Chatterjee, Avula Kiran, Parej Nath, Supriya Suman, Rina Yadav, Rupesh Dash, Soma Chattopadhyay, Ghulam Hussain Syed, Shanti Senapati, Tushar K. Beuria, Rajeeb Swain, Punit Prasad, ILS COVID-19 TEAM, Orissa COVID-19 Study Group, Ajay Parida |
| hCoV-19/India/ILSCV17568/2020 | EPI_ISL_463056 | 5/12/2020 | Institute of Life Sciences, Bhubaneswar | Immunogenomics lab, Institute of Life Sciences, Bhubaneswar | Sunil Raghav, Arup Ghosh, Atimukta Jha, Viplov K. Biswas, Swati Madhulika, Manasi Priyadarshini, Shuchi Smita, Sifu Agarwal, Sanchari Chatterjee, Avula Kiran, Parej Nath, Supriya Suman, Rina Yadav, Rupesh Dash, Soma Chattopadhyay, Ghulam Hussain Syed, Shanti Senapati, Tushar K. Beuria, Rajeeb Swain, Punit Prasad, ILS COVID-19 TEAM, Orissa COVID-19 Study Group, Ajay Parida |
| hCoV-19/India/ILSCV17741/2020 | EPI_ISL_463058 | 5/13/2020 | Institute of Life Sciences, Bhubaneswar | Immunogenomics lab, Institute of Life Sciences, Bhubaneswar | Sunil Raghav, Arup Ghosh, Atimukta Jha, Viplov K. Biswas, Swati Madhulika, Manasi Priyadarshini, Shuchi Smita, Sifu Agarwal, Sanchari Chatterjee, Avula Kiran, Parej Nath, Supriya Suman, Rina Yadav, Rupesh Dash, Soma Chattopadhyay, Ghulam Hussain Syed, Shanti Senapati, Tushar K. Beuria, Rajeeb Swain, Punit Prasad, ILS COVID-19 TEAM, Orissa COVID-19 Study Group, Ajay Parida |
| hCoV-19/India/ILSCV18525/2020 | EPI_ISL_463060 | 5/13/2020 | Institute of Life Sciences, Bhubaneswar | Immunogenomics lab, Institute of Life Sciences, Bhubaneswar | Sunil Raghav, Arup Ghosh, Atimukta Jha, Viplov K. Biswas, Swati Madhulika, Manasi Priyadarshini, Shuchi Smita, Sifu Agarwal, Sanchari Chatterjee, Avula Kiran, Parej Nath, Supriya Suman, Rina Yadav, Rupesh Dash, Soma Chattopadhyay, Ghulam Hussain Syed, Shanti Senapati, Tushar K. Beuria, Rajeeb Swain, Punit Prasad, ILS COVID-19 TEAM, Orissa COVID-19 Study Group, Ajay Parida |
| hCoV-19/India/ILSCV18534/2020 | EPI_ISL_463061 | 5/13/2020 | Institute of Life Sciences, Bhubaneswar | Immunogenomics lab, Institute of Life Sciences, Bhubaneswar | Sunil Raghav, Arup Ghosh, Atimukta Jha, Viplov K. Biswas, Swati Madhulika, Manasi Priyadarshini, Shuchi Smita, Sifu Agarwal, Sanchari Chatterjee, Avula Kiran, Parej Nath, Supriya Suman, Rina Yadav, Rupesh Dash, Soma Chattopadhyay, Ghulam Hussain Syed, Shanti Senapati, Tushar K. Beuria, Rajeeb Swain, Punit Prasad, ILS COVID-19 TEAM, Orissa COVID-19 Study Group, Ajay Parida |
| hCoV-19/India/ILSCV19048/2020 | EPI_ISL_463064 | 5/16/2020 | Institute of Life Sciences, Bhubaneswar | Immunogenomics lab, Institute of Life Sciences, Bhubaneswar | Sunil Raghav, Arup Ghosh, Atimukta Jha, Viplov K. Biswas, Swati Madhulika, Manasi Priyadarshini, Shuchi Smita, Sifu Agarwal, Sanchari Chatterjee, Avula Kiran, Parej Nath, Supriya Suman, Rina Yadav, Rupesh Dash, Soma Chattopadhyay, Ghulam Hussain Syed, Shanti Senapati, Tushar K. Beuria, Rajeeb Swain, Punit Prasad, ILS COVID-19 TEAM, Orissa COVID-19 Study Group, Ajay Parida |
| hCoV-19/India/ILSCV19192/2020 | EPI_ISL_463067 | 5/14/2020 | Institute of Life Sciences, Bhubaneswar | Immunogenomics lab, Institute of Life Sciences, Bhubaneswar | Sunil Raghav, Arup Ghosh, Atimukta Jha, Viplov K. Biswas, Swati Madhulika, Manasi Priyadarshini, Shuchi Smita, Sifu Agarwal, Sanchari Chatterjee, Avula Kiran, Parej Nath, Supriya Suman, Rina Yadav, Rupesh Dash, Soma Chattopadhyay, Ghulam Hussain Syed, Shanti Senapati, Tushar K. Beuria, Rajeeb Swain, Punit Prasad, ILS COVID-19 TEAM, Orissa COVID-19 Study Group, Ajay Parida |
| hCoV-19/India/ILSCV20165/2020 | EPI_ISL_463073 | 5/5/2020 | Institute of Life Sciences, Bhubaneswar | Immunogenomics lab, Institute of Life Sciences, Bhubaneswar | Sunil Raghav, Arup Ghosh, Atimukta Jha, Viplov K. Biswas, Swati Madhulika, Manasi Priyadarshini, Shuchi Smita, Kautilya Kumar Jena, Sandhya Suranjika, Neha Singh, Eshna Laha, Saiket De, Rupesh Dash, Soma Chattopadhyay, Ghulam Hussain Syed, Shanti Senapati, Tushar K. Beuria, Rajeeb Swain, Punit Prasad, ILS COVID-19 TEAM, Orissa COVID-19 Study Group, Ajay Parida |
| hCoV-19/India/ILSCV20166/2020 | EPI_ISL_463074 | 5/5/2020 | Institute of Life Sciences, Bhubaneswar | Immunogenomics lab, Institute of Life Sciences, Bhubaneswar | Sunil Raghav, Arup Ghosh, Atimukta Jha, Viplov K. Biswas, Swati Madhulika, Manasi Priyadarshini, Shuchi Smita, Kautilya Kumar Jena, Sandhya Suranjika, Neha Singh, Eshna Laha, Saiket De, Rupesh Dash, Soma Chattopadhyay, Ghulam Hussain Syed, Shanti Senapati, Tushar K. Beuria, Rajeeb Swain, Punit Prasad, ILS COVID-19 TEAM, Orissa COVID-19 Study Group, Ajay Parida |
| hCoV-19/India/ILSCV20408/2020 | EPI_ISL_463076 | 5/15/2020 | Institute of Life Sciences, Bhubaneswar | Immunogenomics lab, Institute of Life Sciences, Bhubaneswar | Sunil Raghav, Arup Ghosh, Atimukta Jha, Viplov K. Biswas, Swati Madhulika, Manasi Priyadarshini, Shuchi Smita, Kautilya Kumar Jena, Sandhya Suranjika, Neha Singh, Eshna Laha, Saiket De, Rupesh Dash, Soma Chattopadhyay, Ghulam Hussain Syed, Shanti Senapati, Tushar K. Beuria, Rajeeb Swain, Punit Prasad, ILS COVID-19 TEAM, Orissa COVID-19 Study Group, Ajay Parida |
| hCoV-19/India/ILSCV20557/2020 | EPI_ISL_463078 | 5/15/2020 | Institute of Life Sciences, Bhubaneswar | Immunogenomics lab, Institute of Life Sciences, Bhubaneswar | Sunil Raghav, Arup Ghosh, Atimukta Jha, Viplov K. Biswas, Swati Madhulika, Manasi Priyadarshini, Shuchi Smita, Kautilya Kumar Jena, Sandhya Suranjika, Neha Singh, Eshna Laha, Saiket De, Rupesh Dash, Soma Chattopadhyay, Ghulam Hussain Syed, Shanti Senapati, Tushar K. Beuria, Rajeeb Swain, Punit Prasad, ILS COVID-19 TEAM, Orissa COVID-19 Study Group, Ajay Parida |
| hCoV-19/India/ILSCV21721/2020 | EPI_ISL_463085 | 5/16/2020 | Institute of Life Sciences, Bhubaneswar | Immunogenomics lab, Institute of Life Sciences, Bhubaneswar | Sunil Raghav, Arup Ghosh, Atimukta Jha, Viplov K. Biswas, Swati Madhulika, Manasi Priyadarshini, Shuchi Smita, Kautilya Kumar Jena, Sandhya Suranjika, Neha Singh, Eshna Laha, Saiket De, Rupesh Dash, Soma Chattopadhyay, Ghulam Hussain Syed, Shanti Senapati, Tushar K. Beuria, Rajeeb Swain, Punit Prasad, ILS COVID-19 TEAM, Orissa COVID-19 Study Group, Ajay Parida |
| hCoV-19/India/ILSCV24694/2020 | EPI_ISL_463087 | 5/18/2020 | Institute of Life Sciences, Bhubaneswar | Immunogenomics lab, Institute of Life Sciences, Bhubaneswar | Sunil Raghav, Arup Ghosh, Atimukta Jha, Viplov K. Biswas, Swati Madhulika, Manasi Priyadarshini, Shuchi Smita, Kautilya Kumar Jena, Sandhya Suranjika, Neha Singh, Eshna Laha, Saiket De, Rupesh Dash, Soma Chattopadhyay, Ghulam Hussain Syed, Shanti Senapati, Tushar K. Beuria, Rajeeb Swain, Punit Prasad, ILS COVID-19 TEAM, Orissa COVID-19 Study Group, Ajay Parida |
| hCoV-19/India/ILSCV24917/2020 | EPI_ISL_463088 | 5/18/2020 | Institute of Life Sciences, Bhubaneswar | Immunogenomics lab, Institute of Life Sciences, Bhubaneswar | Sunil Raghav, Arup Ghosh, Atimukta Jha, Viplov K. Biswas, Swati Madhulika, Manasi Priyadarshini, Shuchi Smita, Kautilya Kumar Jena, Sandhya Suranjika, Neha Singh, Eshna Laha, Saiket De, Rupesh Dash, Soma Chattopadhyay, Ghulam Hussain Syed, Shanti Senapati, Tushar K. Beuria, Rajeeb Swain, Punit Prasad, ILS COVID-19 TEAM, Orissa COVID-19 Study Group, Ajay Parida |
| hCoV-19/India/ILSCV24918/2020 | EPI_ISL_463089 | 5/18/2020 | Institute of Life Sciences, Bhubaneswar | Immunogenomics lab, Institute of Life Sciences, Bhubaneswar | Sunil Raghav, Arup Ghosh, Atimukta Jha, Viplov K. Biswas, Swati Madhulika, Manasi Priyadarshini, Shuchi Smita, Kautilya Kumar Jena, Sandhya Suranjika, Neha Singh, Eshna Laha, Saiket De, Rupesh Dash, Soma Chattopadhyay, Ghulam Hussain Syed, Shanti Senapati, Tushar K. Beuria, Rajeeb Swain, Punit Prasad, ILS COVID-19 TEAM, Orissa COVID-19 Study Group, Ajay Parida |
| hCoV-19/India/GBRC169a/2020 | EPI_ISL_467029 | 6/5/2020 | GMERS Medical College and Hospital, Gandhinagar | Gujarat Biotechnology Research Centre | Seema Bhatt, Gaurishankar Shrimali, Bhavesh Modi, Bharti Rajani, Tejas Shah, Ankit Hinsu, Pritesh Sabara, Apurvasinh Puvar, Janvi Raval, Zarna Patel, Monika Gandhi, Pinal Trivedi, Maharshi Pandya, Nidhi Patel, Nitin Savaliya, Raghawendra Kumar, Dinesh Kumar, Zuber Saiyed, Komal Patel, Labdhi Pandya, Snehal Bagatharia, Bhavya Jindal, R D Dixit, A M Kadri, Harsh Bakshi, Chaitanya Joshi, Madhvi Josh |
| hCoV-19/India/GBRC169b/2020 | EPI_ISL_467030 | 6/5/2020 | GMERS Medical College and Hospital, Gandhinagar | Gujarat Biotechnology Research Centre | Gaurishankar Shrimali, Bhavesh Modi, Bharti Rajani, Tejas Shah, Ankit Hinsu, Pritesh Sabara, Apurvasinh Puvar, Janvi Raval, Zarna Patel, Monika Gandhi, Pinal Trivedi, Maharshi Pandya, Nidhi Patel, Nitin Savaliya, Raghawendra Kumar, Dinesh Kumar, Zuber Saiyed, Komal Patel, Labdhi Pandya, Snehal Bagatharia, Seema Bhatt, Priyanka P Vatsa, R D Dixit, A M Kadri, Harsh Bakshi, Chaitanya Joshi, Madhvi J |
| hCoV-19/India/GBRC170a/2020 | EPI_ISL_467031 | 6/5/2020 | GMERS Medical College and Hospital, Gandhinagar | Gujarat Biotechnology Research Centre | Bhavesh Modi, Bharti Rajani, Tejas Shah, Ankit Hinsu, Pritesh Sabara, Apurvasinh Puvar, Janvi Raval, Zarna Patel, Monika Gandhi, Pinal Trivedi, Maharshi Pandya, Nidhi Patel, Nitin Savaliya, Raghawendra Kumar, Dinesh Kumar, Zuber Saiyed, Komal Patel, Labdhi Pandya, Snehal Bagatharia, Seema Bhatt, Gaurishankar Shrimali, Pooja P Doshi, R D Dixit, A M Kadri, Harsh Bakshi, Chaitanya Joshi, Madhvi Josh |
| hCoV-19/India/GBRC170b/2020 | EPI_ISL_467032 | 6/5/2020 | GMERS Medical College and Hospital, Gandhinagar | Gujarat Biotechnology Research Centre | Bharti Rajani, Tejas Shah, Ankit Hinsu, Pritesh Sabara, Apurvasinh Puvar, Janvi Raval, Zarna Patel, Monika Gandhi, Pinal Trivedi, Maharshi Pandya, Nidhi Patel, Nitin Savaliya, Raghawendra Kumar, Dinesh Kumar, Zuber Saiyed, Komal Patel, Labdhi Pandya, Snehal Bagatharia, Seema Bhatt, Gaurishankar Shrimali, Bhavesh Modi, Akanksha Verma, R D Dixit, A M Kadri, Harsh Bakshi, Chaitanya Joshi, Madhvi J |
| hCoV-19/India/GBRC171/2020 | EPI_ISL_467033 | 6/5/2020 | GMERS Medical College and Hospital, Gandhinagar | Gujarat Biotechnology Research Centre | Tejas Shah, Ankit Hinsu, Pritesh Sabara, Apurvasinh Puvar, Janvi Raval, Zarna Patel, Monika Gandhi, Pinal Trivedi, Maharshi Pandya, Nidhi Patel, Nitin Savaliya, Raghawendra Kumar, Dinesh Kumar, Zuber Saiyed, Komal Patel, Labdhi Pandya, Snehal Bagatharia, Seema Bhatt, Gaurishankar Shrimali, Bhavesh Modi, Bharti Rajani, Priti Pandita, R D Dixit, A M Kadri, Harsh Bakshi, Chaitanya Joshi, Madhvi Joshi |
| hCoV-19/India/GBRC172/2020 | EPI_ISL_467034 | 6/5/2020 | GMERS Medical College and Hospital, Gandhinagar | Gujarat Biotechnology Research Centre | Ankit Hinsu, Pritesh Sabara, Apurvasinh Puvar, Janvi Raval, Zarna Patel, Monika Gandhi, Pinal Trivedi, Maharshi Pandya, Nidhi Patel, Nitin Savaliya, Raghawendra Kumar, Dinesh Kumar, Zuber Saiyed, Komal Patel, Labdhi Pandya, Snehal Bagatharia, Seema Bhatt, Gaurishankar Shrimali, Bhavesh Modi, Bharti Rajani, Tejas Shah, Pragya Sharma, R D Dixit, A M Kadri, Harsh Bakshi, Chaitanya Joshi, Madhvi Jos |
| hCoV-19/India/GBRC173a/2020 | EPI_ISL_467035 | 6/5/2020 | GMERS Medical College and Hospital, Gandhinagar | Gujarat Biotechnology Research Centre | Pritesh Sabara, Apurvasinh Puvar, Janvi Raval, Zarna Patel, Monika Gandhi, Pinal Trivedi, Maharshi Pandya, Nidhi Patel, Nitin Savaliya, Raghawendra Kumar, Dinesh Kumar, Zuber Saiyed, Komal Patel, Labdhi Pandya, Snehal Bagatharia, Seema Bhatt, Gaurishankar Shrimali, Bhavesh Modi, Bharti Rajani, Tejas Shah, Ankit Hinsu, Neha Rajpara, R D Dixit, A M Kadri, Harsh Bakshi, Chaitanya Joshi, Madhvi Josh |
| hCoV-19/India/GBRC173b/2020 | EPI_ISL_467036 | 6/5/2020 | GMERS Medical College and Hospital, Gandhinagar | Gujarat Biotechnology Research Centre | Apurvasinh Puvar, Janvi Raval, Zarna Patel, Monika Gandhi, Pinal Trivedi, Maharshi Pandya, Nidhi Patel, Nitin Savaliya, Raghawendra Kumar, Dinesh Kumar, Zuber Saiyed, Komal Patel, Labdhi Pandya, Snehal Bagatharia, Seema Bhatt, Gaurishankar Shrimali, Bhavesh Modi, Bharti Rajani, Tejas Shah, Ankit Hinsu, Pritesh Sabara, Afzal Ansari, R D Dixit, A M Kadri, Harsh Bakshi, Chaitanya Joshi, Madhvi Joshi |
| hCoV-19/India/GBRC174a/2020 | EPI_ISL_467037 | 6/5/2020 | GMERS Medical College and Hospital, Gandhinagar | Gujarat Biotechnology Research Centre | Janvi Raval, Zarna Patel, Monika Gandhi, Pinal Trivedi, Maharshi Pandya, Nidhi Patel, Nitin Savaliya, Raghawendra Kumar, Dinesh Kumar, Zuber Saiyed, Komal Patel, Labdhi Pandya, Snehal Bagatharia, Seema Bhatt, Gaurishankar Shrimali, Bhavesh Modi, Bharti Rajani, Tejas Shah, Ankit Hinsu, Pritesh Sabara, Apurvasinh Puvar, Fenil Patel, R D Dixit, A M Kadri, Harsh Bakshi, Chaitanya Joshi, Madhvi Joshi |
| hCoV-19/India/GBRC174b/2020 | EPI_ISL_467038 | 6/5/2020 | GMERS Medical College and Hospital, Gandhinagar | Gujarat Biotechnology Research Centre | Zarna Patel, Monika Gandhi, Pinal Trivedi, Maharshi Pandya, Nidhi Patel, Nitin Savaliya, Raghawendra Kumar, Dinesh Kumar, Zuber Saiyed, Komal Patel, Labdhi Pandya, Snehal Bagatharia, Seema Bhatt, Gaurishankar Shrimali, Bhavesh Modi, Bharti Rajani, Tejas Shah, Ankit Hinsu, Pritesh Sabara, Apurvasinh Puvar, Janvi Raval, Neelam Nathani, R D Dixit, A M Kadri, Harsh Bakshi, Chaitanya Joshi, Madhvi Jo |
| hCoV-19/India/GBRC175/2020 | EPI_ISL_467039 | 6/5/2020 | Government Medical College, Vadodara | Gujarat Biotechnology Research Centre | Meenakshi Shah, Neena Doshi, Varsha Godbole, Tejas Shah, Ankit Hinsu, Pritesh Sabara, Apurvasinh Puvar, Janvi Raval, Zarna Patel, Monika Gandhi, Pinal Trivedi, Maharshi Pandya, Nidhi Patel, Nitin Savaliya, Raghawendra Kumar, Dinesh Kumar, Zuber Saiyed, Komal Patel, Labdhi Pandya, Snehal Bagatharia, Armi Chaudhari, R D Dixit, A M Kadri, Harsh Bakshi, Chaitanya Joshi, Madhvi Joshi, |
| hCoV-19/India/GBRC176/2020 | EPI_ISL_467040 | 6/5/2020 | Government Medical College, Vadodara | Gujarat Biotechnology Research Centre | Neena Doshi, Varsha Godbole, Tejas Shah, Ankit Hinsu, Pritesh Sabara, Apurvasinh Puvar, Janvi Raval, Zarna Patel, Monika Gandhi, Pinal Trivedi, Maharshi Pandya, Nidhi Patel, Nitin Savaliya, Raghawendra Kumar, Dinesh Kumar, Zuber Saiyed, Komal Patel, Labdhi Pandya, Snehal Bagatharia, Meenakshi Shah, Bhavya Jindal, R D Dixit, A M Kadri, Harsh Bakshi, Chaitanya Joshi, Madhvi Joshi, |
| hCoV-19/India/GBRC177a/2020 | EPI_ISL_467041 | 6/3/2020 | B.J. Medical College and Civil hospital | Gujarat Biotechnology Research Centre | Monika Gandhi, Pinal Trivedi, Maharshi Pandya, Nidhi Patel, Nitin Savaliya, Raghawendra Kumar, Dinesh Kumar, Zuber Saiyed, Komal Patel, Labdhi Pandya, Snehal Bagatharia, Pranay Shah, Kamlesh J Upadhyay, Nirav Mungalpara, Tejas Shah, Ankit Hinsu, Pritesh Sabara, Apurvasinh Puvar, Janvi Raval, Zarna Patel, Priyanka P Vatsa, R D Dixit, A M Kadri, Harsh Bakshi, Chaitanya Joshi, Madhvi Joshi, |
| hCoV-19/India/GBRC177b/2020 | EPI_ISL_467042 | 6/3/2020 | B.J. Medical College and Civil hospital | Gujarat Biotechnology Research Centre | Pinal Trivedi, Maharshi Pandya, Nidhi Patel, Nitin Savaliya, Raghawendra Kumar, Dinesh Kumar, Zuber Saiyed, Komal Patel, Labdhi Pandya, Snehal Bagatharia, Pranay Shah, Kamlesh J Upadhyay, Nirav Mungalpara, Tejas Shah, Ankit Hinsu, Pritesh Sabara, Apurvasinh Puvar, Janvi Raval, Zarna Patel, Monika Gandhi, Pooja P Doshi, R D Dixit, A M Kadri, Harsh Bakshi, Chaitanya Joshi, Madhvi Joshi, |
| hCoV-19/India/GBRC178a/2020 | EPI_ISL_467043 | 6/3/2020 | B.J. Medical College and Civil hospital | Gujarat Biotechnology Research Centre | Maharshi Pandya, Nidhi Patel, Nitin Savaliya, Raghawendra Kumar, Dinesh Kumar, Zuber Saiyed, Komal Patel, Labdhi Pandya, Snehal Bagatharia, Pranay Shah, Kamlesh J Upadhyay, Nirav Mungalpara, Tejas Shah, Ankit Hinsu, Pritesh Sabara, Apurvasinh Puvar, Janvi Raval, Zarna Patel, Monika Gandhi, Pinal Trivedi, Akanksha Verma, R D Dixit, A M Kadri, Harsh Bakshi, Chaitanya Joshi, Madhvi Joshi, |
| hCoV-19/India/GBRC178b/2020 | EPI_ISL_467044 | 6/3/2020 | B.J. Medical College and Civil hospital | Gujarat Biotechnology Research Centre | Nidhi Patel, Nitin Savaliya, Raghawendra Kumar, Dinesh Kumar, Zuber Saiyed, Komal Patel, Labdhi Pandya, Snehal Bagatharia, Pranay Shah, Kamlesh J Upadhyay, Nirav Mungalpara, Tejas Shah, Ankit Hinsu, Pritesh Sabara, Apurvasinh Puvar, Janvi Raval, Zarna Patel, Monika Gandhi, Pinal Trivedi, Maharshi Pandya, Priti Pandita, R D Dixit, A M Kadri, Harsh Bakshi, Chaitanya Joshi, Madhvi Joshi, |
| hCoV-19/India/GBRC179a/2020 | EPI_ISL_467045 | 6/3/2020 | B.J. Medical College and Civil hospital | Gujarat Biotechnology Research Centre | Nitin Savaliya, Raghawendra Kumar, Dinesh Kumar, Zuber Saiyed, Komal Patel, Labdhi Pandya, Snehal Bagatharia, Pranay Shah, Kamlesh J Upadhyay, Nirav Mungalpara, Tejas Shah, Ankit Hinsu, Pritesh Sabara, Apurvasinh Puvar, Janvi Raval, Zarna Patel, Monika Gandhi, Pinal Trivedi, Maharshi Pandya, Nidhi Patel, Pragya Sharma, R D Dixit, A M Kadri, Harsh Bakshi, Chaitanya Joshi, Madhvi Joshi, |
| hCoV-19/India/GBRC179b/2020 | EPI_ISL_467046 | 6/3/2020 | B.J. Medical College and Civil hospital | Gujarat Biotechnology Research Centre | Raghawendra Kumar, Dinesh Kumar, Zuber Saiyed, Komal Patel, Labdhi Pandya, Snehal Bagatharia, Pranay Shah, Kamlesh J Upadhyay, Nirav Mungalpara, Tejas Shah, Ankit Hinsu, Pritesh Sabara, Apurvasinh Puvar, Janvi Raval, Zarna Patel, Monika Gandhi, Pinal Trivedi, Maharshi Pandya, Nidhi Patel, Nitin Savaliya, Neha Rajpara, R D Dixit, A M Kadri, Harsh Bakshi, Chaitanya Joshi, Madhvi Joshi, |
| hCoV-19/India/GBRC180a/2020 | EPI_ISL_467047 | 6/3/2020 | B.J. Medical College and Civil hospital | Gujarat Biotechnology Research Centre | Dinesh Kumar, Zuber Saiyed, Komal Patel, Labdhi Pandya, Snehal Bagatharia, Pranay Shah, Kamlesh J Upadhyay, Nirav Mungalpara, Tejas Shah, Ankit Hinsu, Pritesh Sabara, Apurvasinh Puvar, Janvi Raval, Zarna Patel, Monika Gandhi, Pinal Trivedi, Maharshi Pandya, Nidhi Patel, Nitin Savaliya, Raghawendra Kumar, Afzal Ansari, R D Dixit, A M Kadri, Harsh Bakshi, Chaitanya Joshi, Madhvi Joshi, |
| hCoV-19/India/GBRC180b/2020 | EPI_ISL_467048 | 6/3/2020 | B.J. Medical College and Civil hospital | Gujarat Biotechnology Research Centre | Zuber Saiyed, Komal Patel, Labdhi Pandya, Snehal Bagatharia, Pranay Shah, Kamlesh J Upadhyay, Nirav Mungalpara, Tejas Shah, Ankit Hinsu, Pritesh Sabara, Apurvasinh Puvar, Janvi Raval, Zarna Patel, Monika Gandhi, Pinal Trivedi, Maharshi Pandya, Nidhi Patel, Nitin Savaliya, Raghawendra Kumar, Dinesh Kumar, Fenil Patel, R D Dixit, A M Kadri, Harsh Bakshi, Chaitanya Joshi, Madhvi Joshi, |
| hCoV-19/India/GBRC181a/2020 | EPI_ISL_467049 | 6/3/2020 | B.J. Medical College and Civil hospital | Gujarat Biotechnology Research Centre | Komal Patel, Labdhi Pandya, Snehal Bagatharia, Pranay Shah, Kamlesh J Upadhyay, Nirav Mungalpara, Tejas Shah, Ankit Hinsu, Pritesh Sabara, Apurvasinh Puvar, Janvi Raval, Zarna Patel, Monika Gandhi, Pinal Trivedi, Maharshi Pandya, Nidhi Patel, Nitin Savaliya, Raghawendra Kumar, Dinesh Kumar, Zuber Saiyed, Neelam Nathani, R D Dixit, A M Kadri, Harsh Bakshi, Chaitanya Joshi, Madhvi Joshi, |
| hCoV-19/India/GBRC181b/2020 | EPI_ISL_467050 | 6/3/2020 | B.J. Medical College and Civil hospital | Gujarat Biotechnology Research Centre | Labdhi Pandya, Snehal Bagatharia, Pranay Shah, Kamlesh J Upadhyay, Nirav Mungalpara, Tejas Shah, Ankit Hinsu, Pritesh Sabara, Apurvasinh Puvar, Janvi Raval, Zarna Patel, Monika Gandhi, Pinal Trivedi, Maharshi Pandya, Nidhi Patel, Nitin Savaliya, Raghawendra Kumar, Dinesh Kumar, Zuber Saiyed, Komal Patel, Armi Chaudhari, R D Dixit, A M Kadri, Harsh Bakshi, Chaitanya Joshi, Madhvi Joshi, |
| hCoV-19/India/GBRC182a/2020 | EPI_ISL_467051 | 6/3/2020 | B.J. Medical College and Civil hospital | Gujarat Biotechnology Research Centre | Snehal Bagatharia, Pranay Shah, Kamlesh J Upadhyay, Nirav Mungalpara, Tejas Shah, Ankit Hinsu, Pritesh Sabara, Apurvasinh Puvar, Janvi Raval, Zarna Patel, Monika Gandhi, Pinal Trivedi, Maharshi Pandya, Nidhi Patel, Nitin Savaliya, Raghawendra Kumar, Dinesh Kumar, Zuber Saiyed, Komal Patel, Labdhi Pandya, Bhavya Jindal, R D Dixit, A M Kadri, Harsh Bakshi, Chaitanya Joshi, Madhvi Joshi, |
| hCoV-19/India/GBRC182b/2020 | EPI_ISL_467052 | 6/3/2020 | B.J. Medical College and Civil hospital | Gujarat Biotechnology Research Centre | Pranay Shah, Kamlesh J Upadhyay, Nirav Mungalpara, Tejas Shah, Ankit Hinsu, Pritesh Sabara, Apurvasinh Puvar, Janvi Raval, Zarna Patel, Monika Gandhi, Pinal Trivedi, Maharshi Pandya, Nidhi Patel, Nitin Savaliya, Raghawendra Kumar, Dinesh Kumar, Zuber Saiyed, Komal Patel, Labdhi Pandya, Snehal Bagatharia, Priyanka P Vatsa, R D Dixit, A M Kadri, Harsh Bakshi, Chaitanya Joshi, Madhvi Joshi, |
| hCoV-19/India/GBRC183a/2020 | EPI_ISL_467053 | 6/3/2020 | B.J. Medical College and Civil hospital | Gujarat Biotechnology Research Centre | Kamlesh J Upadhyay, Nirav Mungalpara, Tejas Shah, Ankit Hinsu, Pritesh Sabara, Apurvasinh Puvar, Janvi Raval, Zarna Patel, Monika Gandhi, Pinal Trivedi, Maharshi Pandya, Nidhi Patel, Nitin Savaliya, Raghawendra Kumar, Dinesh Kumar, Zuber Saiyed, Komal Patel, Labdhi Pandya, Snehal Bagatharia, Pranay Shah, Pooja P Doshi, R D Dixit, A M Kadri, Harsh Bakshi, Chaitanya Joshi, Madhvi Joshi, |
| hCoV-19/India/GBRC183b/2020 | EPI_ISL_467054 | 6/3/2020 | B.J. Medical College and Civil hospital | Gujarat Biotechnology Research Centre | Nirav Mungalpara, Tejas Shah, Ankit Hinsu, Pritesh Sabara, Apurvasinh Puvar, Janvi Raval, Zarna Patel, Monika Gandhi, Pinal Trivedi, Maharshi Pandya, Nidhi Patel, Nitin Savaliya, Raghawendra Kumar, Dinesh Kumar, Zuber Saiyed, Komal Patel, Labdhi Pandya, Snehal Bagatharia, Pranay Shah, Kamlesh J Upadhyay, Akanksha Verma, R D Dixit, A M Kadri, Harsh Bakshi, Chaitanya Joshi, Madhvi Joshi, |
| hCoV-19/India/GBRC184/2020 | **EPI_ISL_469024** | 5/27/2020 | B.J. Medical College and Civil hospital | Gujarat Biotechnology Research Centre | Tejas Shah, Ankit Hinsu, Pritesh Sabara, Apurvasinh Puvar, Janvi Raval, Zarna Patel, Monika Gandhi, Pinal Trivedi, Maharshi Pandya, Nidhi Patel, Nitin Savaliya, Raghawendra Kumar, Dinesh Kumar, Zuber Saiyed, Komal Patel, Labdhi Pandya, Snehal Bagatharia, Pranay Shah, Kamlesh J Upadhyay, Nirav Mungalpara, Priti Pandita, R D Dixit, A M Kadri, Harsh Bakshi, Chaitanya Joshi, Madhvi Joshi |
| hCoV-19/India/GBRC185a/2020 | **EPI_ISL_469025** | 6/3/2020 | B.J. Medical College and Civil hospital | Gujarat Biotechnology Research Centre | Ankit Hinsu, Pritesh Sabara, Apurvasinh Puvar, Janvi Raval, Zarna Patel, Monika Gandhi, Pinal Trivedi, Maharshi Pandya, Nidhi Patel, Nitin Savaliya, Raghawendra Kumar, Dinesh Kumar, Zuber Saiyed, Komal Patel, Labdhi Pandya, Snehal Bagatharia, Pranay Shah, Kamlesh J Upadhyay, Nirav Mungalpara, Tejas Shah, Pragya Sharma, R D Dixit, A M Kadri, Harsh Bakshi, Chaitanya Joshi, Madhvi Joshi |
| hCoV-19/India/GBRC185b/2020 | **EPI_ISL_469026** | 6/3/2020 | B.J. Medical College and Civil hospital | Gujarat Biotechnology Research Centre | Pritesh Sabara, Apurvasinh Puvar, Janvi Raval, Zarna Patel, Monika Gandhi, Pinal Trivedi, Maharshi Pandya, Nidhi Patel, Nitin Savaliya, Raghawendra Kumar, Dinesh Kumar, Zuber Saiyed, Komal Patel, Labdhi Pandya, Snehal Bagatharia, Pranay Shah, Kamlesh J Upadhyay, Nirav Mungalpara, Tejas Shah, Ankit Hinsu, Neha Rajpara, R D Dixit, A M Kadri, Harsh Bakshi, Chaitanya Joshi, Madhvi Joshi |
| hCoV-19/India/GBRC186a/2020 | **EPI_ISL_469027** | 6/3/2020 | B.J. Medical College and Civil hospital | Gujarat Biotechnology Research Centre | Apurvasinh Puvar, Janvi Raval, Zarna Patel, Monika Gandhi, Pinal Trivedi, Maharshi Pandya, Nidhi Patel, Nitin Savaliya, Raghawendra Kumar, Dinesh Kumar, Zuber Saiyed, Komal Patel, Labdhi Pandya, Snehal Bagatharia, Pranay Shah, Kamlesh J Upadhyay, Nirav Mungalpara, Tejas Shah, Ankit Hinsu, Pritesh Sabara, Afzal Ansari, R D Dixit, A M Kadri, Harsh Bakshi, Chaitanya Joshi, Madhvi Joshi |
| hCoV-19/India/GBRC186b/2020 | **EPI_ISL_469028** | 6/3/2020 | B.J. Medical College and Civil hospital | Gujarat Biotechnology Research Centre | Janvi Raval, Zarna Patel, Monika Gandhi, Pinal Trivedi, Maharshi Pandya, Nidhi Patel, Nitin Savaliya, Raghawendra Kumar, Dinesh Kumar, Zuber Saiyed, Komal Patel, Labdhi Pandya, Snehal Bagatharia, Pranay Shah, Kamlesh J Upadhyay, Nirav Mungalpara, Tejas Shah, Ankit Hinsu, Pritesh Sabara, Apurvasinh Puvar, Fenil Patel, R D Dixit, A M Kadri, Harsh Bakshi, Chaitanya Joshi, Madhvi Joshi |
| hCoV-19/India/GBRC187a/2020 | **EPI_ISL_469029** | 6/5/2020 | Government Medical College, Vadodara | Gujarat Biotechnology Research Centre | Zarna Patel, Monika Gandhi, Pinal Trivedi, Maharshi Pandya, Nidhi Patel, Nitin Savaliya, Raghawendra Kumar, Dinesh Kumar, Zuber Saiyed, Komal Patel, Labdhi Pandya, Snehal Bagatharia, Meenakshi Shah, Neena Doshi, Varsha Godbole, Tejas Shah, Ankit Hinsu, Pritesh Sabara, Apurvasinh Puvar, Janvi Raval, Neelam Nathani, R D Dixit, A M Kadri, Harsh Bakshi, Chaitanya Joshi, Madhvi Joshi |
| hCoV-19/India/GBRC187b/2020 | **EPI_ISL_469030** | 6/5/2020 | Government Medical College, Vadodara | Gujarat Biotechnology Research Centre | Monika Gandhi, Pinal Trivedi, Maharshi Pandya, Nidhi Patel, Nitin Savaliya, Raghawendra Kumar, Dinesh Kumar, Zuber Saiyed, Komal Patel, Labdhi Pandya, Snehal Bagatharia, Meenakshi Shah, Neena Doshi, Varsha Godbole, Tejas Shah, Ankit Hinsu, Pritesh Sabara, Apurvasinh Puvar, Janvi Raval, Zarna Patel, Armi Chaudhari, R D Dixit, A M Kadri, Harsh Bakshi, Chaitanya Joshi, Madhvi Joshi |
| hCoV-19/India/GBRC188/2020 | **EPI_ISL_469031** | 6/5/2020 | Government Medical College, Vadodara | Gujarat Biotechnology Research Centre | Pinal Trivedi, Maharshi Pandya, Nidhi Patel, Nitin Savaliya, Raghawendra Kumar, Dinesh Kumar, Zuber Saiyed, Komal Patel, Labdhi Pandya, Snehal Bagatharia, Meenakshi Shah, Neena Doshi, Varsha Godbole, Tejas Shah, Ankit Hinsu, Pritesh Sabara, Apurvasinh Puvar, Janvi Raval, Zarna Patel, Monika Gandhi, Bhavya Jindal, R D Dixit, A M Kadri, Harsh Bakshi, Chaitanya Joshi, Madhvi Joshi |
| hCoV-19/India/GBRC189/2020 | **EPI_ISL_469032** | 6/2/2020 | Government Medical College, Vadodara | Gujarat Biotechnology Research Centre | Maharshi Pandya, Nidhi Patel, Nitin Savaliya, Raghawendra Kumar, Dinesh Kumar, Zuber Saiyed, Komal Patel, Labdhi Pandya, Snehal Bagatharia, Meenakshi Shah, Neena Doshi, Varsha Godbole, Tejas Shah, Ankit Hinsu, Pritesh Sabara, Apurvasinh Puvar, Janvi Raval, Zarna Patel, Monika Gandhi, Pinal Trivedi, Pragya Sharma, R D Dixit, A M Kadri, Harsh Bakshi, Chaitanya Joshi, Madhvi Joshi |
| hCoV-19/India/GBRC190/2020 | **EPI_ISL_469033** | 6/5/2020 | Government Medical College, Vadodara | Gujarat Biotechnology Research Centre | Nidhi Patel, Nitin Savaliya, Raghawendra Kumar, Dinesh Kumar, Zuber Saiyed, Komal Patel, Labdhi Pandya, Snehal Bagatharia, Meenakshi Shah, Neena Doshi, Varsha Godbole, Tejas Shah, Ankit Hinsu, Pritesh Sabara, Apurvasinh Puvar, Janvi Raval, Zarna Patel, Monika Gandhi, Pinal Trivedi, Maharshi Pandya, Priyanka P Vatsa, R D Dixit, A M Kadri, Harsh Bakshi, Chaitanya Joshi, Madhvi Joshi |
| hCoV-19/India/GBRC191a/2020 | **EPI_ISL_469034** | 6/5/2020 | Government Medical College, Vadodara | Gujarat Biotechnology Research Centre | Nitin Savaliya, Raghawendra Kumar, Dinesh Kumar, Zuber Saiyed, Komal Patel, Labdhi Pandya, Snehal Bagatharia, Meenakshi Shah, Neena Doshi, Varsha Godbole, Tejas Shah, Ankit Hinsu, Pritesh Sabara, Apurvasinh Puvar, Janvi Raval, Zarna Patel, Monika Gandhi, Pinal Trivedi, Maharshi Pandya, Nidhi Patel, Pooja P Doshi, R D Dixit, A M Kadri, Harsh Bakshi, Chaitanya Joshi, Madhvi Joshi |
| hCoV-19/India/GBRC191b/2020 | **EPI_ISL_469035** | 6/5/2020 | Government Medical College, Vadodara | Gujarat Biotechnology Research Centre | Raghawendra Kumar, Dinesh Kumar, Zuber Saiyed, Komal Patel, Labdhi Pandya, Snehal Bagatharia, Meenakshi Shah, Neena Doshi, Varsha Godbole, Tejas Shah, Ankit Hinsu, Pritesh Sabara, Apurvasinh Puvar, Janvi Raval, Zarna Patel, Monika Gandhi, Pinal Trivedi, Maharshi Pandya, Nidhi Patel, Nitin Savaliya, Akanksha Verma, R D Dixit, A M Kadri, Harsh Bakshi, Chaitanya Joshi, Madhvi Joshi |
| hCoV-19/India/GBRC192/2020 | **EPI_ISL_469036** | 6/5/2020 | Government Medical College, Vadodara | Gujarat Biotechnology Research Centre | Dinesh Kumar, Zuber Saiyed, Komal Patel, Labdhi Pandya, Snehal Bagatharia, Meenakshi Shah, Neena Doshi, Varsha Godbole, Tejas Shah, Ankit Hinsu, Pritesh Sabara, Apurvasinh Puvar, Janvi Raval, Zarna Patel, Monika Gandhi, Pinal Trivedi, Maharshi Pandya, Nidhi Patel, Nitin Savaliya, Raghawendra Kumar, Priti Pandita, R D Dixit, A M Kadri, Harsh Bakshi, Chaitanya Joshi, Madhvi Joshi |
| hCoV-19/India/GBRC193a/2020 | **EPI_ISL_469037** | 6/3/2020 | GMERS Medical College & Hospital | Gujarat Biotechnology Research Centre | Zuber Saiyed, Komal Patel, Labdhi Pandya, Snehal Bagatharia, Meenakshi Shah, Neena Doshi, Varsha Godbole, Tejas Shah, Ankit Hinsu, Pritesh Sabara, Apurvasinh Puvar, Janvi Raval, Zarna Patel, Monika Gandhi, Pinal Trivedi, Maharshi Pandya, Nidhi Patel, Nitin Savaliya, Raghawendra Kumar, Dinesh Kumar, Pragya Sharma, R D Dixit, A M Kadri, Harsh Bakshi, Chaitanya Joshi, Madhvi Joshi |
| hCoV-19/India/GBRC193b/2020 | **EPI_ISL_469038** | 6/3/2020 | GMERS Medical College & Hospital | Gujarat Biotechnology Research Centre | Komal Patel, Labdhi Pandya, Snehal Bagatharia, Meenakshi Shah, Neena Doshi, Varsha Godbole, Tejas Shah, Ankit Hinsu, Pritesh Sabara, Apurvasinh Puvar, Janvi Raval, Zarna Patel, Monika Gandhi, Pinal Trivedi, Maharshi Pandya, Nidhi Patel, Nitin Savaliya, Raghawendra Kumar, Dinesh Kumar, Zuber Saiyed, Neha Rajpara, R D Dixit, A M Kadri, Harsh Bakshi, Chaitanya Joshi, Madhvi Joshi |
| hCoV-19/India/GBRC195a/2020 | **EPI_ISL_469039** | 6/8/2020 | GMERS Medical College & Hospital | Gujarat Biotechnology Research Centre | Labdhi Pandya, Snehal Bagatharia, Meenakshi Shah, Neena Doshi, Varsha Godbole, Tejas Shah, Ankit Hinsu, Pritesh Sabara, Apurvasinh Puvar, Janvi Raval, Zarna Patel, Monika Gandhi, Pinal Trivedi, Maharshi Pandya, Nidhi Patel, Nitin Savaliya, Raghawendra Kumar, Dinesh Kumar, Zuber Saiyed, Komal Patel, Afzal Ansari, R D Dixit, A M Kadri, Harsh Bakshi, Chaitanya Joshi, Madhvi Joshi |
| hCoV-19/India/GBRC195b/2020 | **EPI_ISL_469040** | 6/8/2020 | GMERS Medical College & Hospital | Gujarat Biotechnology Research Centre | Snehal Bagatharia, Meenakshi Shah, Neena Doshi, Varsha Godbole, Tejas Shah, Ankit Hinsu, Pritesh Sabara, Apurvasinh Puvar, Janvi Raval, Zarna Patel, Monika Gandhi, Pinal Trivedi, Maharshi Pandya, Nidhi Patel, Nitin Savaliya, Raghawendra Kumar, Dinesh Kumar, Zuber Saiyed, Komal Patel, Labdhi Pandya, Fenil Patel, R D Dixit, A M Kadri, Harsh Bakshi, Chaitanya Joshi, Madhvi Joshi |
| hCoV-19/India/GBRC196/2020 | **EPI_ISL_469041** | 6/8/2020 | GMERS Medical College & Hospital | Gujarat Biotechnology Research Centre | Meenakshi Shah, Neena Doshi, Varsha Godbole, Tejas Shah, Ankit Hinsu, Pritesh Sabara, Apurvasinh Puvar, Janvi Raval, Zarna Patel, Monika Gandhi, Pinal Trivedi, Maharshi Pandya, Nidhi Patel, Nitin Savaliya, Raghawendra Kumar, Dinesh Kumar, Zuber Saiyed, Komal Patel, Labdhi Pandya, Snehal Bagatharia, Neelam Nathani, R D Dixit, A M Kadri, Harsh Bakshi, Chaitanya Joshi, Madhvi Joshi |
| hCoV-19/India/GBRC197/2020 | **EPI_ISL_469042** | 6/8/2020 | GMERS Medical College & Hospital | Gujarat Biotechnology Research Centre | Neena Doshi, Varsha Godbole, Tejas Shah, Ankit Hinsu, Pritesh Sabara, Apurvasinh Puvar, Janvi Raval, Zarna Patel, Monika Gandhi, Pinal Trivedi, Maharshi Pandya, Nidhi Patel, Nitin Savaliya, Raghawendra Kumar, Dinesh Kumar, Zuber Saiyed, Komal Patel, Labdhi Pandya, Snehal Bagatharia, Meenakshi Shah, Armi Chaudhari, R D Dixit, A M Kadri, Harsh Bakshi, Chaitanya Joshi, Madhvi Joshi |
| hCoV-19/India/GBRC198/2020 | **EPI_ISL_469043** | 6/8/2020 | Dr. N. D. Desai Medical College & Hospital | Gujarat Biotechnology Research Centre | J G Buch, Jigar Gusani, Supreet Prabhu, Tejas Shah, Ankit Hinsu, Pritesh Sabara, Apurvasinh Puvar, Janvi Raval, Zarna Patel, Monika Gandhi, Pinal Trivedi, Maharshi Pandya, Nidhi Patel, Nitin Savaliya, Raghawendra Kumar, Dinesh Kumar, Zuber Saiyed, Komal Patel, Labdhi Pandya, Snehal Bagatharia, Bhavya Jindal, R D Dixit, A M Kadri, Harsh Bakshi, Chaitanya Joshi, Madhvi Joshi |
| hCoV-19/India/GBRC199/2020 | **EPI_ISL_469044** | 6/8/2020 | Dr. N. D. Desai Medical College & Hospital | Gujarat Biotechnology Research Centre | Jigar Gusani, Supreet Prabhu, Tejas Shah, Ankit Hinsu, Pritesh Sabara, Apurvasinh Puvar, Janvi Raval, Zarna Patel, Monika Gandhi, Pinal Trivedi, Maharshi Pandya, Nidhi Patel, Nitin Savaliya, Raghawendra Kumar, Dinesh Kumar, Zuber Saiyed, Komal Patel, Labdhi Pandya, Snehal Bagatharia, J G Buch, Neha Rajpara, R D Dixit, A M Kadri, Harsh Bakshi, Chaitanya Joshi, Madhvi Joshi |
| hCoV-19/India/GBRC200/2020 | **EPI_ISL_469045** | 6/8/2020 | Dr. N. D. Desai Medical College & Hospital | Gujarat Biotechnology Research Centre | Supreet Prabhu, Tejas Shah, Ankit Hinsu, Pritesh Sabara, Apurvasinh Puvar, Janvi Raval, Zarna Patel, Monika Gandhi, Pinal Trivedi, Maharshi Pandya, Nidhi Patel, Nitin Savaliya, Raghawendra Kumar, Dinesh Kumar, Zuber Saiyed, Komal Patel, Labdhi Pandya, Snehal Bagatharia, J G Buch, Jigar Gusani, Priyanka P Vatsa, R D Dixit, A M Kadri, Harsh Bakshi, Chaitanya Joshi, Madhvi Joshi |
| hCoV-19/India/GBRC201a/2020 | **EPI_ISL_469046** | 6/8/2020 | Dr. N. D. Desai Medical College & Hospital | Gujarat Biotechnology Research Centre | Tejas Shah, Ankit Hinsu, Pritesh Sabara, Apurvasinh Puvar, Janvi Raval, Zarna Patel, Monika Gandhi, Pinal Trivedi, Maharshi Pandya, Nidhi Patel, Nitin Savaliya, Raghawendra Kumar, Dinesh Kumar, Zuber Saiyed, Komal Patel, Labdhi Pandya, Snehal Bagatharia, J G Buch, Jigar Gusani, Supreet Prabhu, Pooja P Doshi, R D Dixit, A M Kadri, Harsh Bakshi, Chaitanya Joshi, Madhvi Joshi |
| hCoV-19/India/GBRC201b/2020 | **EPI_ISL_469047** | 6/8/2020 | Dr. N. D. Desai Medical College & Hospital | Gujarat Biotechnology Research Centre | Ankit Hinsu, Pritesh Sabara, Apurvasinh Puvar, Janvi Raval, Zarna Patel, Monika Gandhi, Pinal Trivedi, Maharshi Pandya, Nidhi Patel, Nitin Savaliya, Raghawendra Kumar, Dinesh Kumar, Zuber Saiyed, Komal Patel, Labdhi Pandya, Snehal Bagatharia, J G Buch, Jigar Gusani, Supreet Prabhu, Tejas Shah, Akanksha Verma, R D Dixit, A M Kadri, Harsh Bakshi, Chaitanya Joshi, Madhvi Joshi |
| hCoV-19/India/GBRC202/2020 | **EPI_ISL_469048** | 6/5/2020 | Banas Medical College and Research Institute | Gujarat Biotechnology Research Centre | Radhika Khara, Sunil R Joshi, Viren s Doshi, Zarna Patel, Monika Gandhi, Pinal Trivedi, Maharshi Pandya, Nidhi Patel, Nitin Savaliya, Raghawendra Kumar, Dinesh Kumar, Zuber Saiyed, Komal Patel, Labdhi Pandya, Snehal Bagatharia, Tejas Shah, Ankit Hinsu, Pritesh Sabara, Apurvasinh Puvar, Janvi Raval, Priti Pandita, R D Dixit, A M Kadri, Harsh Bakshi, Chaitanya Joshi, Madhvi Joshi |
| hCoV-19/India/CCMB_C20/2020 | **EPI_ISL_471583** | 4/29/2020 | King Institute of Preventive Medicine & Research | CSIR-Centre for Cellular and Molecular Biology | K.Kaveri,S.Sivasubramanian,S.Vennila,P.Padmapriya,R.Kiruba,S.Magesh,G. Dhinakar Raj, G. Ravikumar, P. Azhahianambi,K Thangaraj,Payel Mukherjee, Sofia Banu, Priya Singh, Dhiviya Vedagiri, Divya Gupta, Vishal Sah, Santosh Kumar Kuncha, Krishnan Harinivas Harshan, Archana Bharadwaj Siva, Karthik Bharadwaj Tallapaka, Shagufta Khan, Lamuk Zaveri, Namami Gaur, Sakshi Shambhavi, Tulasi Nagab |
| hCoV-19/India/CCMB_C21/2020 | **EPI_ISL_471584** | 5/6/2020 | King Institute of Preventive Medicine & Research | CSIR-Centre for Cellular and Molecular Biology | K.Kaveri,S.Sivasubramanian,S.Vennila,P.Padmapriya,R.Kiruba,S.Magesh,G. Dhinakar Raj, G. Ravikumar, P. Azhahianambi,K Thangaraj,Payel Mukherjee, Sofia Banu, Priya Singh, Dhiviya Vedagiri, Divya Gupta, Vishal Sah, Santosh Kumar Kuncha, Krishnan Harinivas Harshan, Archana Bharadwaj Siva, Karthik Bharadwaj Tallapaka, Shagufta Khan, Lamuk Zaveri, Namami Gaur, Sakshi Shambhavi, Tulasi Nagab |
| hCoV-19/India/CCMB_J812_P2/2020 | **EPI_ISL_471585** | 4/8/2020 | CSIR-Centre for Cellular and Molecular Biology | CSIR-Centre for Cellular and Molecular Biology | Dhiviya Vedagiri, Divya Gupta, Vishal Sah, Payel Mukherjee, Sofia Banu, Priya Singh, Santosh Kumar Kuncha, Archana Bharadwaj Siva, Karthik Bharadwaj Tallapaka, Shagufta Khan, Lamuk Zaveri, Namami Gaur, Sakshi Shambhavi, Tulasi Nagabandi, Purushotham Vodnala, Rakesh K Mishra, Divya Tej Sowpati, Krishnan Harinivas Harshan |
| hCoV-19/India/CCMB_L040/2020 | **EPI_ISL_471586** | 4/18/2020 | CSIR-Centre for Cellular and Molecular Biology | CSIR-Centre for Cellular and Molecular Biology | Lamuk Zaveri, Shagufta Khan, Namami Gaur, Sakshi Shambhavi, Tulasi Nagabandi, Purushotham Vodnala, Payel Mukherjee, Sofia Banu, Priya Singh, Dhiviya Vedagiri, Divya Gupta, Vishal Sah, Santosh Kumar Kuncha, Krishnan Harinivas Harshan, Archana Bharadwaj Siva, Karthik Bharadwaj Tallapaka,Zeba Rizvi, Zuberwasim Sayyad, Kakade Aishwarya Arun, Amrutha H C, Ananga Ghosh, Rakesh K Mishra, [illegible] |
| hCoV-19/India/CCMB_L040_P3/2020 | **EPI_ISL_471587** | 4/18/2020 | CSIR-Centre for Cellular and Molecular Biology | CSIR-Centre for Cellular and Molecular Biology | Dhiviya Vedagiri, Divya Gupta, Vishal Sah, Payel Mukherjee, Sofia Banu, Priya Singh, Santosh Kumar Kuncha, Archana Bharadwaj Siva, Karthik Bharadwaj Tallapaka, Shagufta Khan, Lamuk Zaveri, Namami Gaur, Sakshi Shambhavi, Tulasi Nagabandi, Purushotham Vodnala, Rakesh K Mishra, Divya Tej Sowpati, Krishnan Harinivas Harshan |
| hCoV-19/India/CCMB_L1208/2020 | **EPI_ISL_471588** | 5/21/2020 | CSIR-Centre for Cellular and Molecular Biology | CSIR-Centre for Cellular and Molecular Biology | Lamuk Zaveri, Shagufta Khan, Namami Gaur, Sakshi Shambhavi, Tulasi Nagabandi, Purushotham Vodnala, Payel Mukherjee, Sofia Banu, Priya Singh, Dhiviya Vedagiri, Divya Gupta, Vishal Sah, Santosh Kumar Kuncha, Krishnan Harinivas Harshan, Archana Bharadwaj Siva, Karthik Bharadwaj Tallapaka, Renu Sudhakar, Somesh Gorde, Gangumala Srinivas Reddy, Sujoy Deb, Swati Bayyana, Rakesh K Mishra, |
| hCoV-19/India/CCMB_L1276/2020 | **EPI_ISL_471589** | 5/21/2020 | CSIR-Centre for Cellular and Molecular Biology | CSIR-Centre for Cellular and Molecular Biology | Lamuk Zaveri, Shagufta Khan, Namami Gaur, Sakshi Shambhavi, Tulasi Nagabandi, Purushotham Vodnala, Payel Mukherjee, Sofia Banu, Priya Singh, Dhiviya Vedagiri, Divya Gupta, Vishal Sah, Santosh Kumar Kuncha, Krishnan Harinivas Harshan, Archana Bharadwaj Siva, Karthik Bharadwaj Tallapaka,Umesh Kumar, Unis Ahmad Bhat, Ajay Sarawagi, Priyanka Pant, Rajkanwar Nathawat, Rakesh K Mishra, Di |
| hCoV-19/India/CCMB_L1451/2020 | **EPI_ISL_471590** | 5/25/2020 | CSIR-Centre for Cellular and Molecular Biology | CSIR-Centre for Cellular and Molecular Biology | Lamuk Zaveri, Shagufta Khan, Namami Gaur, Sakshi Shambhavi, Tulasi Nagabandi, Purushotham Vodnala, Payel Mukherjee, Sofia Banu, Priya Singh, Dhiviya Vedagiri, Divya Gupta, Vishal Sah, Santosh Kumar Kuncha, Krishnan Harinivas Harshan, Archana Bharadwaj Siva, Karthik Bharadwaj Tallapaka,Zeba Rizvi, Zuberwasim Sayyad, Kakade Aishwarya Arun, Amrutha H C, Ananga Ghosh, Rakesh K Mishra, [illegible] |
| hCoV-19/India/CCMB_L1452/2020 | **EPI_ISL_471591** | 5/25/2020 | CSIR-Centre for Cellular and Molecular Biology | CSIR-Centre for Cellular and Molecular Biology | Namami Gaur, Sakshi Shambhavi, Lamuk Zaveri, Shagufta Khan, Tulasi Nagabandi, Purushotham Vodnala, Payel Mukherjee, Sofia Banu, Priya Singh, Dhiviya Vedagiri, Divya Gupta, Vishal Sah, Santosh Kumar Kuncha, Krishnan Harinivas Harshan, Archana Bharadwaj Siva, Karthik Bharadwaj Tallapaka, Zeba Rizvi, Zuberwasim Sayyad, Kakade Aishwarya Arun, Amrutha H C, Ananga Ghosh, Rakesh K Mishra, |
| hCoV-19/India/CCMB_L1453/2020 | **EPI_ISL_471592** | 5/25/2020 | CSIR-Centre for Cellular and Molecular Biology | CSIR-Centre for Cellular and Molecular Biology | Namami Gaur, Sakshi Shambhavi, Lamuk Zaveri, Shagufta Khan, Tulasi Nagabandi, Purushotham Vodnala, Payel Mukherjee, Sofia Banu, Priya Singh, Dhiviya Vedagiri, Divya Gupta, Vishal Sah, Santosh Kumar Kuncha, Krishnan Harinivas Harshan, Archana Bharadwaj Siva, Karthik Bharadwaj Tallapaka, Nikhil Hajirnis, Pratheusa Maccha, M Soujanya Reddy,G. Aditya Kumar, Koushick Sivakumar, Rakesh K Mis |
| hCoV-19/India/CCMB_L1454/2020 | **EPI_ISL_471593** | 5/25/2020 | CSIR-Centre for Cellular and Molecular Biology | CSIR-Centre for Cellular and Molecular Biology | Namami Gaur, Sakshi Shambhavi, Lamuk Zaveri, Shagufta Khan, Tulasi Nagabandi, Purushotham Vodnala, Payel Mukherjee, Sofia Banu, Priya Singh, Dhiviya Vedagiri, Divya Gupta, Vishal Sah, Santosh Kumar Kuncha, Krishnan Harinivas Harshan, Archana Bharadwaj Siva, Karthik Bharadwaj Tallapaka, Zeba Rizvi, Zuberwasim Sayyad, Kakade Aishwarya Arun, Amrutha H C, Ananga Ghosh, Rakesh K Mishra, |
| hCoV-19/India/CCMB_L1455/2020 | **EPI_ISL_471594** | 5/25/2020 | CSIR-Centre for Cellular and Molecular Biology | CSIR-Centre for Cellular and Molecular Biology | Namami Gaur, Sakshi Shambhavi, Lamuk Zaveri, Shagufta Khan, Tulasi Nagabandi, Purushotham Vodnala, Payel Mukherjee, Sofia Banu, Priya Singh, Dhiviya Vedagiri, Divya Gupta, Vishal Sah, Santosh Kumar Kuncha, Krishnan Harinivas Harshan, Archana Bharadwaj Siva, Karthik Bharadwaj Tallapaka,Kezia J Ann, Radhika Khandelwal, Roshan Maku Venkata, Shemin Mansuri, Sonu Uday, Rakesh K Mishra, [illegible] |
| hCoV-19/India/CCMB_L1456/2020 | **EPI_ISL_471595** | 5/25/2020 | CSIR-Centre for Cellular and Molecular Biology | CSIR-Centre for Cellular and Molecular Biology | Payel Mukherjee, Sofia Banu, Priya Singh, Dhiviya Vedagiri, Divya Gupta, Vishal Sah, Santosh Kumar Kuncha, Krishnan Harinivas Harshan, Archana Bharadwaj Siva, Karthik Bharadwaj Tallapaka, Shagufta Khan, Lamuk Zaveri, Namami Gaur, Sakshi Shambhavi, Tulasi Nagabandi, Purushotham Vodnala, G. Aditya Kumar, Koushick Sivakumar, Pooja Ramesh Gupta, Rajan Kumar Jha, Shraddha Vijay Lahoti, Rak |
| hCoV-19/India/CCMB_L1459/2020 | **EPI_ISL_471596** | 5/25/2020 | CSIR-Centre for Cellular and Molecular Biology | CSIR-Centre for Cellular and Molecular Biology | Payel Mukherjee, Sofia Banu, Priya Singh, Dhiviya Vedagiri, Divya Gupta, Vishal Sah, Santosh Kumar Kuncha, Krishnan Harinivas Harshan, Archana Bharadwaj Siva, Karthik Bharadwaj Tallapaka, Shagufta Khan, Lamuk Zaveri, Namami Gaur, Sakshi Shambhavi, Tulasi Nagabandi, Purushotham Vodnala, Gokulan C G, Gunjan Purohit, Hanuman Tulashiram Kale, Pankaj Kumar, Prachand Issarapu, Rakesh K Mis |
| hCoV-19/India/CCMB_L1460/2020 | **EPI_ISL_471597** | 5/25/2020 | CSIR-Centre for Cellular and Molecular Biology | CSIR-Centre for Cellular and Molecular Biology | Payel Mukherjee, Sofia Banu, Priya Singh, Dhiviya Vedagiri, Divya Gupta, Vishal Sah, Santosh Kumar Kuncha, Krishnan Harinivas Harshan, Archana Bharadwaj Siva, Karthik Bharadwaj Tallapaka, Shagufta Khan, Lamuk Zaveri, Namami Gaur, Sakshi Shambhavi, Tulasi Nagabandi, Purushotham Vodnala, Rakesh K Mishra, Sonu Uday, Sudipta Mondal, Annapoorna P Karthyayani, Debabrata Jana, Debrya Saha, |
| hCoV-19/India/CCMB_L1465/2020 | **EPI_ISL_471598** | 5/25/2020 | CSIR-Centre for Cellular and Molecular Biology | CSIR-Centre for Cellular and Molecular Biology | Payel Mukherjee, Sofia Banu, Priya Singh, Dhiviya Vedagiri, Divya Gupta, Vishal Sah, Santosh Kumar Kuncha, Krishnan Harinivas Harshan, Archana Bharadwaj Siva, Karthik Bharadwaj Tallapaka, Shagufta Khan, Lamuk Zaveri, Namami Gaur, Sakshi Shambhavi, Tulasi Nagabandi, Purushotham Vodnala,Deepak Kumar, Devi Prasad Vijayashankar, Disha Nanda, Divya Das, Jotin Gogoi, Manish Bhattacharjee, Ra |
| hCoV-19/India/CCMB_L1466/2020 | **EPI_ISL_471599** | 5/25/2020 | CSIR-Centre for Cellular and Molecular Biology | CSIR-Centre for Cellular and Molecular Biology | Sakshi Shambhavi, Lamuk Zaveri, Shagufta Khan, Namami Gaur, Tulasi Nagabandi, Purushotham Vodnala, Payel Mukherjee, Sofia Banu, Priya Singh, Dhiviya Vedagiri, Divya Gupta, Vishal Sah, Santosh Kumar Kuncha, Krishnan Harinivas Harshan, Archana Bharadwaj Siva, Karthik Bharadwaj Tallapaka, Deepak Kumar, Devi Prasad Vijayashankar, Disha Nanda, Divya Das, Jotin Gogoi, Manish Bhattacharjee, R |
| hCoV-19/India/CCMB_L1469/2020 | **EPI_ISL_471600** | 5/25/2020 | CSIR-Centre for Cellular and Molecular Biology | CSIR-Centre for Cellular and Molecular Biology | Sakshi Shambhavi, Lamuk Zaveri, Shagufta Khan, Namami Gaur, Tulasi Nagabandi, Purushotham Vodnala, Payel Mukherjee, Sofia Banu, Priya Singh, Dhiviya Vedagiri, Divya Gupta, Vishal Sah, Santosh Kumar Kuncha, Krishnan Harinivas Harshan, Archana Bharadwaj Siva, Karthik Bharadwaj Tallapaka, G. Aditya Kumar, Koushick Sivakumar, Pooja Ramesh Gupta, Rajan Kumar Jha, Shraddha Vijay Lahoti, Rak |
| hCoV-19/India/CCMB_L1470/2020 | **EPI_ISL_471601** | 5/25/2020 | CSIR-Centre for Cellular and Molecular Biology | CSIR-Centre for Cellular and Molecular Biology | Sakshi Shambhavi, Lamuk Zaveri, Shagufta Khan, Namami Gaur, Tulasi Nagabandi, Purushotham Vodnala, Payel Mukherjee, Sofia Banu, Priya Singh, Dhiviya Vedagiri, Divya Gupta, Vishal Sah, Santosh Kumar Kuncha, Krishnan Harinivas Harshan, Archana Bharadwaj Siva, Karthik Bharadwaj Tallapaka,Nikhil Hajirnis, Pratheusa Maccha, M Soujanya Reddy,G. Aditya Kumar, Koushick Sivakumar, Rakesh K Mis |
| hCoV-19/India/CCMB_L1471/2020 | **EPI_ISL_471602** | 5/25/2020 | CSIR-Centre for Cellular and Molecular Biology | CSIR-Centre for Cellular and Molecular Biology | Sakshi Shambhavi, Lamuk Zaveri, Shagufta Khan, Namami Gaur, Tulasi Nagabandi, Purushotham Vodnala, Payel Mukherjee, Sofia Banu, Priya Singh, Dhiviya Vedagiri, Divya Gupta, Vishal Sah, Santosh Kumar Kuncha, Krishnan Harinivas Harshan, Archana Bharadwaj Siva, Karthik Bharadwaj Tallapaka,Nikhil Hajirnis, Pratheusa Maccha, M Soujanya Reddy,G. Aditya Kumar, Koushick Sivakumar,Disha Nanda, [illegible] |
| hCoV-19/India/CCMB_L1475/2020 | **EPI_ISL_471603** | 5/25/2020 | CSIR-Centre for Cellular and Molecular Biology | CSIR-Centre for Cellular and Molecular Biology | Shagufta Khan, Lamuk Zaveri, Namami Gaur, Sakshi Shambhavi, Tulasi Nagabandi, Purushotham Vodnala, Payel Mukherjee, Sofia Banu, Priya Singh, Dhiviya Vedagiri, Divya Gupta, Vishal Sah, Santosh Kumar Kuncha, Krishnan Harinivas Harshan, Archana Bharadwaj Siva, Karthik Bharadwaj Tallapaka, Disha Nanda, Divya Das, Jotin Gogoi, Manish Bhattacharjee, Ravi Prasad Mukku, Rakesh K Mishra, Divya T |
| hCoV-19/India/CCMB_L1476/2020 | **EPI_ISL_471604** | 5/25/2020 | CSIR-Centre for Cellular and Molecular Biology | CSIR-Centre for Cellular and Molecular Biology | Shagufta Khan, Lamuk Zaveri, Namami Gaur, Sakshi Shambhavi, Tulasi Nagabandi, Purushotham Vodnala, Payel Mukherjee, Sofia Banu, Priya Singh, Dhiviya Vedagiri, Divya Gupta, Vishal Sah, Santosh Kumar Kuncha, Krishnan Harinivas Harshan, Archana Bharadwaj Siva, Karthik Bharadwaj Tallapaka, Renu Sudhakar, Somesh Gorde, Gangumala Srinivas Reddy, Sujoy Deb, Swati Bayyana, Rakesh K Mishra, |
| hCoV-19/India/CCMB_L1477/2020 | **EPI_ISL_471605** | 5/25/2020 | CSIR-Centre for Cellular and Molecular Biology | CSIR-Centre for Cellular and Molecular Biology | Shagufta Khan, Lamuk Zaveri, Namami Gaur, Sakshi Shambhavi, Tulasi Nagabandi, Purushotham Vodnala, Payel Mukherjee, Sofia Banu, Priya Singh, Dhiviya Vedagiri, Divya Gupta, Vishal Sah, Santosh Kumar Kuncha, Krishnan Harinivas Harshan, Archana Bharadwaj Siva, Karthik Bharadwaj Tallapaka,Preethi Jampala, Sharada Ravi Iyer, Sulagana Mukherjee, Swetha Sundar, Peddapuvala Sai Uday Kiran Rak |
| hCoV-19/India/CCMB_L1478/2020 | **EPI_ISL_471606** | 5/25/2020 | CSIR-Centre for Cellular and Molecular Biology | CSIR-Centre for Cellular and Molecular Biology | Shagufta Khan, Lamuk Zaveri, Namami Gaur, Sakshi Shambhavi, Tulasi Nagabandi, Purushotham Vodnala, Payel Mukherjee, Sofia Banu, Priya Singh, Dhiviya Vedagiri, Divya Gupta, Vishal Sah, Santosh Kumar Kuncha, Krishnan Harinivas Harshan, Archana Bharadwaj Siva, Karthik Bharadwaj Tallapaka,Umesh Kumar, Unis Ahmad Bhat, Ajay Sarawagi, Priyanka Pant, Rajkanwar Nathawat, Rakesh K Mishra, Di |
| hCoV-19/India/CCMB_L1484/2020 | **EPI_ISL_471607** | 5/25/2020 | CSIR-Centre for Cellular and Molecular Biology | CSIR-Centre for Cellular and Molecular Biology | Sofia Banu, Payel Mukherjee, Priya Singh, Dhiviya Vedagiri, Divya Gupta, Vishal Sah, Santosh Kumar Kuncha, Krishnan Harinivas Harshan, Archana Bharadwaj Siva, Karthik Bharadwaj Tallapaka, Shagufta Khan, Lamuk Zaveri, Namami Gaur, Sakshi Shambhavi, Tulasi Nagabandi, Purushotham Vodnala, Deepak Kumar, Devi Prasad Vijayashankar, Disha Nanda, Divya Das, Jotin Gogoi, Manish Bhattacharjee, R |
| hCoV-19/India/CCMB_L1487/2020 | **EPI_ISL_471608** | 5/25/2020 | CSIR-Centre for Cellular and Molecular Biology | CSIR-Centre for Cellular and Molecular Biology | Sofia Banu, Payel Mukherjee, Priya Singh, Dhiviya Vedagiri, Divya Gupta, Vishal Sah, Santosh Kumar Kuncha, Krishnan Harinivas Harshan, Archana Bharadwaj Siva, Karthik Bharadwaj Tallapaka, Shagufta Khan, Lamuk Zaveri, Namami Gaur, Sakshi Shambhavi, Tulasi Nagabandi, Purushotham Vodnala, Disha Nanda, Divya Das, Jotin Gogoi, Manish Bhattacharjee, Ravi Prasad Mukku, Rakesh K Mishra, Divya T |
| hCoV-19/India/CCMB_L1492/2020 | **EPI_ISL_471609** | 5/25/2020 | CSIR-Centre for Cellular and Molecular Biology | CSIR-Centre for Cellular and Molecular Biology | Sofia Banu, Payel Mukherjee, Priya Singh, Dhiviya Vedagiri, Divya Gupta, Vishal Sah, Santosh Kumar Kuncha, Krishnan Harinivas Harshan, Archana Bharadwaj Siva, Karthik Bharadwaj Tallapaka, Shagufta Khan, Lamuk Zaveri, Namami Gaur, Sakshi Shambhavi, Tulasi Nagabandi, Purushotham Vodnala, Gokulan C G, Gunjan Purohit, Hanuman Tulashiram Kale, Pankaj Kumar, Prachand Issarapu, Rakesh K Mis |
| hCoV-19/India/CCMB_L1494/2020 | **EPI_ISL_471610** | 5/25/2020 | CSIR-Centre for Cellular and Molecular Biology | CSIR-Centre for Cellular and Molecular Biology | Sofia Banu, Payel Mukherjee, Priya Singh, Dhiviya Vedagiri, Divya Gupta, Vishal Sah, Santosh Kumar Kuncha, Krishnan Harinivas Harshan, Archana Bharadwaj Siva, Karthik Bharadwaj Tallapaka, Shagufta Khan, Lamuk Zaveri, Namami Gaur, Sakshi Shambhavi, Tulasi Nagabandi, Purushotham Vodnala,Preethi Jampala, Sharada Ravi Iyer, Sulagana Mukherjee, Swetha Sundar, Peddapuvala Sai Uday Kiran, Ra |
| hCoV-19/India/CCMB_L1495/2020 | **EPI_ISL_471611** | 5/25/2020 | CSIR-Centre for Cellular and Molecular Biology | CSIR-Centre for Cellular and Molecular Biology | Tulasi Nagabandi, Namami Gaur, Sakshi Shambhavi, Lamuk Zaveri, Shagufta Khan, Purushotham Vodnala, Payel Mukherjee, Sofia Banu, Priya Singh, Dhiviya Vedagiri, Divya Gupta, Vishal Sah, Santosh Kumar Kuncha, Krishnan Harinivas Harshan, Archana Bharadwaj Siva, Karthik Bharadwaj Tallapaka,G. Aditya Kumar, Koushick Sivakumar, Pooja Ramesh Gupta, Rajan Kumar Jha, Shraddha Vijay Lahoti, Rak |
| hCoV-19/India/CCMB_L1496/2020 | **EPI_ISL_471612** | 5/25/2020 | CSIR-Centre for Cellular and Molecular Biology | CSIR-Centre for Cellular and Molecular Biology | Tulasi Nagabandi, Namami Gaur, Sakshi Shambhavi, Lamuk Zaveri, Shagufta Khan, Purushotham Vodnala, Payel Mukherjee, Sofia Banu, Priya Singh, Dhiviya Vedagiri, Divya Gupta, Vishal Sah, Santosh Kumar Kuncha, Krishnan Harinivas Harshan, Archana Bharadwaj Siva, Karthik Bharadwaj Tallapaka,Kezia J Ann, Radhika Khandelwal, Roshan Maku Venkata, Shemin Mansuri, Sonu Uday, Rakesh K Mishra, [illegible] |
| hCoV-19/India/CCMB_L1497/2020 | **EPI_ISL_471613** | 5/25/2020 | CSIR-Centre for Cellular and Molecular Biology | CSIR-Centre for Cellular and Molecular Biology | Tulasi Nagabandi, Namami Gaur, Sakshi Shambhavi, Lamuk Zaveri, Shagufta Khan, Purushotham Vodnala, Payel Mukherjee, Sofia Banu, Priya Singh, Dhiviya Vedagiri, Divya Gupta, Vishal Sah, Santosh Kumar Kuncha, Krishnan Harinivas Harshan, Archana Bharadwaj Siva, Karthik Bharadwaj Tallapaka,G. Aditya Kumar, Koushick Sivakumar, Pooja Ramesh Gupta, Rajan Kumar Jha, Shraddha Vijay Lahoti, Rak |
| hCoV-19/India/CCMB_L1499/2020 | **EPI_ISL_471614** | 5/25/2020 | CSIR-Centre for Cellular and Molecular Biology | CSIR-Centre for Cellular and Molecular Biology | Tulasi Nagabandi, Namami Gaur, Sakshi Shambhavi, Lamuk Zaveri, Shagufta Khan, Purushotham Vodnala, Payel Mukherjee, Sofia Banu, Priya Singh, Dhiviya Vedagiri, Divya Gupta, Vishal Sah, Santosh Kumar Kuncha, Krishnan Harinivas Harshan, Archana Bharadwaj Siva, Karthik Bharadwaj Tallapaka,Kezia J Ann, Radhika Khandelwal, Roshan Maku Venkata, Shemin Mansuri, Sonu Uday, Rakesh K Mishra, [illegible] |
| hCoV-19/India/CCMB_L1500/2020 | **EPI_ISL_471615** | 5/25/2020 | CSIR-Centre for Cellular and Molecular Biology | CSIR-Centre for Cellular and Molecular Biology | Lamuk Zaveri, Shagufta Khan, Namami Gaur, Sakshi Shambhavi, Tulasi Nagabandi, Purushotham Vodnala, Payel Mukherjee, Sofia Banu, Priya Singh, Dhiviya Vedagiri, Divya Gupta, Vishal Sah, Santosh Kumar Kuncha, Krishnan Harinivas Harshan, Archana Bharadwaj Siva, Karthik Bharadwaj Tallapaka,Zeba Rizvi, Zuberwasim Sayyad, Kakade Aishwarya Arun, Amrutha H C, Ananga Ghosh, Rakesh K Mishra, [illegible] |
| hCoV-19/India/CCMB_L1510/2020 | **EPI_ISL_471616** | 5/26/2020 | CSIR-Centre for Cellular and Molecular Biology | CSIR-Centre for Cellular and Molecular Biology | Lamuk Zaveri, Shagufta Khan, Namami Gaur, Sakshi Shambhavi, Tulasi Nagabandi, Purushotham Vodnala, Payel Mukherjee, Sofia Banu, Priya Singh, Dhiviya Vedagiri, Divya Gupta, Vishal Sah, Santosh Kumar Kuncha, Krishnan Harinivas Harshan, Archana Bharadwaj Siva, Karthik Bharadwaj Tallapaka, Renu Sudhakar, Somesh Gorde, Gangumala Srinivas Reddy, Sujoy Deb, Swati Bayyana, Rakesh K Mishra, |
| hCoV-19/India/CCMB_L1542/2020 | **EPI_ISL_471617** | 5/26/2020 | CSIR-Centre for Cellular and Molecular Biology | CSIR-Centre for Cellular and Molecular Biology | Lamuk Zaveri, Shagufta Khan, Namami Gaur, Sakshi Shambhavi, Tulasi Nagabandi, Purushotham Vodnala, Payel Mukherjee, Sofia Banu, Priya Singh, Dhiviya Vedagiri, Divya Gupta, Vishal Sah, Santosh Kumar Kuncha, Krishnan Harinivas Harshan, Archana Bharadwaj Siva, Karthik Bharadwaj Tallapaka,Umesh Kumar, Unis Ahmad Bhat, Ajay Sarawagi, Priyanka Pant, Rajkanwar Nathawat, Rakesh K Mishra, Di |
| hCoV-19/India/CCMB_L1551/2020 | **EPI_ISL_471618** | 5/26/2020 | CSIR-Centre for Cellular and Molecular Biology | CSIR-Centre for Cellular and Molecular Biology | Lamuk Zaveri, Shagufta Khan, Namami Gaur, Sakshi Shambhavi, Tulasi Nagabandi, Purushotham Vodnala, Payel Mukherjee, Sofia Banu, Priya Singh, Dhiviya Vedagiri, Divya Gupta, Vishal Sah, Santosh Kumar Kuncha, Krishnan Harinivas Harshan, Archana Bharadwaj Siva, Karthik Bharadwaj Tallapaka,Zeba Rizvi, Zuberwasim Sayyad, Kakade Aishwarya Arun, Amrutha H C, Ananga Ghosh, Rakesh K Mishra, [illegible] |
| hCoV-19/India/CCMB_L1554/2020 | **EPI_ISL_471619** | 5/26/2020 | CSIR-Centre for Cellular and Molecular Biology | CSIR-Centre for Cellular and Molecular Biology | Namami Gaur, Sakshi Shambhavi, Lamuk Zaveri, Shagufta Khan, Tulasi Nagabandi, Purushotham Vodnala, Payel Mukherjee, Sofia Banu, Priya Singh, Dhiviya Vedagiri, Divya Gupta, Vishal Sah, Santosh Kumar Kuncha, Krishnan Harinivas Harshan, Archana Bharadwaj Siva, Karthik Bharadwaj Tallapaka, Zeba Rizvi, Zuberwasim Sayyad, Kakade Aishwarya Arun, Amrutha H C, Ananga Ghosh, Rakesh K Mishra, |
| hCoV-19/India/CCMB_L1564/2020 | **EPI_ISL_471620** | 5/26/2020 | CSIR-Centre for Cellular and Molecular Biology | CSIR-Centre for Cellular and Molecular Biology | Namami Gaur, Sakshi Shambhavi, Lamuk Zaveri, Shagufta Khan, Tulasi Nagabandi, Purushotham Vodnala, Payel Mukherjee, Sofia Banu, Priya Singh, Dhiviya Vedagiri, Divya Gupta, Vishal Sah, Santosh Kumar Kuncha, Krishnan Harinivas Harshan, Archana Bharadwaj Siva, Karthik Bharadwaj Tallapaka, Nikhil Hajirnis, Pratheusa Maccha, M Soujanya Reddy,G. Aditya Kumar, Koushick Sivakumar, Rakesh K Mis |
| hCoV-19/India/CCMB_L1675/2020 | **EPI_ISL_471621** | 5/29/2020 | CSIR-Centre for Cellular and Molecular Biology | CSIR-Centre for Cellular and Molecular Biology | Namami Gaur, Sakshi Shambhavi, Lamuk Zaveri, Shagufta Khan, Tulasi Nagabandi, Purushotham Vodnala, Payel Mukherjee, Sofia Banu, Priya Singh, Dhiviya Vedagiri, Divya Gupta, Vishal Sah, Santosh Kumar Kuncha, Krishnan Harinivas Harshan, Archana Bharadwaj Siva, Karthik Bharadwaj Tallapaka, Zeba Rizvi, Zuberwasim Sayyad, Kakade Aishwarya Arun, Amrutha H C, Ananga Ghosh, Rakesh K Mishra, |
| hCoV-19/India/CCMB_L1684/2020 | **EPI_ISL_471622** | 5/29/2020 | CSIR-Centre for Cellular and Molecular Biology | CSIR-Centre for Cellular and Molecular Biology | Namami Gaur, Sakshi Shambhavi, Lamuk Zaveri, Shagufta Khan, Tulasi Nagabandi, Purushotham Vodnala, Payel Mukherjee, Sofia Banu, Priya Singh, Dhiviya Vedagiri, Divya Gupta, Vishal Sah, Santosh Kumar Kuncha, Krishnan Harinivas Harshan, Archana Bharadwaj Siva, Karthik Bharadwaj Tallapaka,Kezia J Ann, Radhika Khandelwal, Roshan Maku Venkata, Shemin Mansuri, Sonu Uday, Rakesh K Mishra, [illegible] |
| hCoV-19/India/CCMB_L1693/2020 | **EPI_ISL_471623** | 5/29/2020 | CSIR-Centre for Cellular and Molecular Biology | CSIR-Centre for Cellular and Molecular Biology | Payel Mukherjee, Sofia Banu, Priya Singh, Dhiviya Vedagiri, Divya Gupta, Vishal Sah, Santosh Kumar Kuncha, Krishnan Harinivas Harshan, Archana Bharadwaj Siva, Karthik Bharadwaj Tallapaka, Shagufta Khan, Lamuk Zaveri, Namami Gaur, Sakshi Shambhavi, Tulasi Nagabandi, Purushotham Vodnala, G. Aditya Kumar, Koushick Sivakumar, Pooja Ramesh Gupta, Rajan Kumar Jha, Shraddha Vijay Lahoti, Rak |
| hCoV-19/India/CCMB_L1694/2020 | **EPI_ISL_471624** | 5/29/2020 | CSIR-Centre for Cellular and Molecular Biology | CSIR-Centre for Cellular and Molecular Biology | Payel Mukherjee, Sofia Banu, Priya Singh, Dhiviya Vedagiri, Divya Gupta, Vishal Sah, Santosh Kumar Kuncha, Krishnan Harinivas Harshan, Archana Bharadwaj Siva, Karthik Bharadwaj Tallapaka, Shagufta Khan, Lamuk Zaveri, Namami Gaur, Sakshi Shambhavi, Tulasi Nagabandi, Purushotham Vodnala, Gokulan C G, Gunjan Purohit, Hanuman Tulashiram Kale, Pankaj Kumar, Prachand Issarapu, Rakesh K Mis |
| hCoV-19/India/CCMB_L1721/2020 | **EPI_ISL_471625** | 5/30/2020 | CSIR-Centre for Cellular and Molecular Biology | CSIR-Centre for Cellular and Molecular Biology | Payel Mukherjee, Sofia Banu, Priya Singh, Dhiviya Vedagiri, Divya Gupta, Vishal Sah, Santosh Kumar Kuncha, Krishnan Harinivas Harshan, Archana Bharadwaj Siva, Karthik Bharadwaj Tallapaka, Shagufta Khan, Lamuk Zaveri, Namami Gaur, Sakshi Shambhavi, Tulasi Nagabandi, Purushotham Vodnala, Rakesh K Mishra, Sonu Uday, Sudipta Mondal, Annapoorna P Karthyayani, Debabrata Jana, Debrya Saha, |
| hCoV-19/India/CCMB_L1724/2020 | **EPI_ISL_471626** | 5/30/2020 | CSIR-Centre for Cellular and Molecular Biology | CSIR-Centre for Cellular and Molecular Biology | Payel Mukherjee, Sofia Banu, Priya Singh, Dhiviya Vedagiri, Divya Gupta, Vishal Sah, Santosh Kumar Kuncha, Krishnan Harinivas Harshan, Archana Bharadwaj Siva, Karthik Bharadwaj Tallapaka, Shagufta Khan, Lamuk Zaveri, Namami Gaur, Sakshi Shambhavi, Tulasi Nagabandi, Purushotham Vodnala,Deepak Kumar, Devi Prasad Vijayashankar, Disha Nanda, Divya Das, Jotin Gogoi, Manish Bhattacharjee, Ra |
| hCoV-19/India/CCMB_L1726/2020 | **EPI_ISL_471627** | 5/30/2020 | CSIR-Centre for Cellular and Molecular Biology | CSIR-Centre for Cellular and Molecular Biology | Sakshi Shambhavi, Lamuk Zaveri, Shagufta Khan, Namami Gaur, Tulasi Nagabandi, Purushotham Vodnala, Payel Mukherjee, Sofia Banu, Priya Singh, Dhiviya Vedagiri, Divya Gupta, Vishal Sah, Santosh Kumar Kuncha, Krishnan Harinivas Harshan, Archana Bharadwaj Siva, Karthik Bharadwaj Tallapaka, Deepak Kumar, Devi Prasad Vijayashankar, Disha Nanda, Divya Das, Jotin Gogoi, Manish Bhattacharjee, R |
| hCoV-19/India/CCMB_L1728/2020 | **EPI_ISL_471628** | 5/30/2020 | CSIR-Centre for Cellular and Molecular Biology | CSIR-Centre for Cellular and Molecular Biology | Sakshi Shambhavi, Lamuk Zaveri, Shagufta Khan, Namami Gaur, Tulasi Nagabandi, Purushotham Vodnala, Payel Mukherjee, Sofia Banu, Priya Singh, Dhiviya Vedagiri, Divya Gupta, Vishal Sah, Santosh Kumar Kuncha, Krishnan Harinivas Harshan, Archana Bharadwaj Siva, Karthik Bharadwaj Tallapaka, G. Aditya Kumar, Koushick Sivakumar, Pooja Ramesh Gupta, Rajan Kumar Jha, Shraddha Vijay Lahoti, Rak |
| hCoV-19/India/CCMB_L1738/2020 | **EPI_ISL_471629** | 6/1/2020 | CSIR-Centre for Cellular and Molecular Biology | CSIR-Centre for Cellular and Molecular Biology | Sakshi Shambhavi, Lamuk Zaveri, Shagufta Khan, Namami Gaur, Tulasi Nagabandi, Purushotham Vodnala, Payel Mukherjee, Sofia Banu, Priya Singh, Dhiviya Vedagiri, Divya Gupta, Vishal Sah, Santosh Kumar Kuncha, Krishnan Harinivas Harshan, Archana Bharadwaj Siva, Karthik Bharadwaj Tallapaka,Nikhil Hajirnis, Pratheusa Maccha, M Soujanya Reddy,G. Aditya Kumar, Koushick Sivakumar, Rakesh K Mis |
| hCoV-19/India/CCMB_L1743/2020 | **EPI_ISL_471630** | 6/1/2020 | CSIR-Centre for Cellular and Molecular Biology | CSIR-Centre for Cellular and Molecular Biology | Sakshi Shambhavi, Lamuk Zaveri, Shagufta Khan, Namami Gaur, Tulasi Nagabandi, Purushotham Vodnala, Payel Mukherjee, Sofia Banu, Priya Singh, Dhiviya Vedagiri, Divya Gupta, Vishal Sah, Santosh Kumar Kuncha, Krishnan Harinivas Harshan, Archana Bharadwaj Siva, Karthik Bharadwaj Tallapaka,Nikhil Hajirnis, Pratheusa Maccha, M Soujanya Reddy,G. Aditya Kumar, Koushick Sivakumar,Disha Nanda, [illegible] |
| hCoV-19/India/CCMB_L1746/2020 | **EPI_ISL_471631** | 6/1/2020 | CSIR-Centre for Cellular and Molecular Biology | CSIR-Centre for Cellular and Molecular Biology | Shagufta Khan, Lamuk Zaveri, Namami Gaur, Sakshi Shambhavi, Tulasi Nagabandi, Purushotham Vodnala, Payel Mukherjee, Sofia Banu, Priya Singh, Dhiviya Vedagiri, Divya Gupta, Vishal Sah, Santosh Kumar Kuncha, Krishnan Harinivas Harshan, Archana Bharadwaj Siva, Karthik Bharadwaj Tallapaka, Disha Nanda, Divya Das, Jotin Gogoi, Manish Bhattacharjee, Ravi Prasad Mukku, Rakesh K Mishra, Divya T |
| hCoV-19/India/CCMB_L1762/2020 | **EPI_ISL_471632** | 6/1/2020 | CSIR-Centre for Cellular and Molecular Biology | CSIR-Centre for Cellular and Molecular Biology | Shagufta Khan, Lamuk Zaveri, Namami Gaur, Sakshi Shambhavi, Tulasi Nagabandi, Purushotham Vodnala, Payel Mukherjee, Sofia Banu, Priya Singh, Dhiviya Vedagiri, Divya Gupta, Vishal Sah, Santosh Kumar Kuncha, Krishnan Harinivas Harshan, Archana Bharadwaj Siva, Karthik Bharadwaj Tallapaka, Renu Sudhakar, Somesh Gorde, Gangumala Srinivas Reddy, Sujoy Deb, Swati Bayyana, Rakesh K Mishra, |
| hCoV-19/India/CCMB_L1781/2020 | **EPI_ISL_471633** | 6/2/2020 | CSIR-Centre for Cellular and Molecular Biology | CSIR-Centre for Cellular and Molecular Biology | Shagufta Khan, Lamuk Zaveri, Namami Gaur, Sakshi Shambhavi, Tulasi Nagabandi, Purushotham Vodnala, Payel Mukherjee, Sofia Banu, Priya Singh, Dhiviya Vedagiri, Divya Gupta, Vishal Sah, Santosh Kumar Kuncha, Krishnan Harinivas Harshan, Archana Bharadwaj Siva, Karthik Bharadwaj Tallapaka,Preethi Jampala, Sharada Ravi Iyer, Sulagana Mukherjee, Swetha Sundar, Peddapuvala Sai Uday Kiran Rak |
| hCoV-19/India/CCMB_L1806/2020 | **EPI_ISL_471634** | 6/2/2020 | CSIR-Centre for Cellular and Molecular Biology | CSIR-Centre for Cellular and Molecular Biology | Shagufta Khan, Lamuk Zaveri, Namami Gaur, Sakshi Shambhavi, Tulasi Nagabandi, Purushotham Vodnala, Payel Mukherjee, Sofia Banu, Priya Singh, Dhiviya Vedagiri, Divya Gupta, Vishal Sah, Santosh Kumar Kuncha, Krishnan Harinivas Harshan, Archana Bharadwaj Siva, Karthik Bharadwaj Tallapaka,Umesh Kumar, Unis Ahmad Bhat, Ajay Sarawagi, Priyanka Pant, Rajkanwar Nathawat, Rakesh K Mishra, Di |
| hCoV-19/India/CCMB_L1809/2020 | **EPI_ISL_471635** | 6/2/2020 | CSIR-Centre for Cellular and Molecular Biology | CSIR-Centre for Cellular and Molecular Biology | Sofia Banu, Payel Mukherjee, Priya Singh, Dhiviya Vedagiri, Divya Gupta, Vishal Sah, Santosh Kumar Kuncha, Krishnan Harinivas Harshan, Archana Bharadwaj Siva, Karthik Bharadwaj Tallapaka, Shagufta Khan, Lamuk Zaveri, Namami Gaur, Sakshi Shambhavi, Tulasi Nagabandi, Purushotham Vodnala, Deepak Kumar, Devi Prasad Vijayashankar, Disha Nanda, Divya Das, Jotin Gogoi, Manish Bhattacharjee, R |
| hCoV-19/India/CCMB_L1810/2020 | **EPI_ISL_471636** | 6/2/2020 | CSIR-Centre for Cellular and Molecular Biology | CSIR-Centre for Cellular and Molecular Biology | Sofia Banu, Payel Mukherjee, Priya Singh, Dhiviya Vedagiri, Divya Gupta, Vishal Sah, Santosh Kumar Kuncha, Krishnan Harinivas Harshan, Archana Bharadwaj Siva, Karthik Bharadwaj Tallapaka, Shagufta Khan, Lamuk Zaveri, Namami Gaur, Sakshi Shambhavi, Tulasi Nagabandi, Purushotham Vodnala, Disha Nanda, Divya Das, Jotin Gogoi, Manish Bhattacharjee, Ravi Prasad Mukku, Rakesh K Mishra, Divya T |
| hCoV-19/India/CCMB_L1811/2020 | **EPI_ISL_471637** | 6/2/2020 | CSIR-Centre for Cellular and Molecular Biology | CSIR-Centre for Cellular and Molecular Biology | Sofia Banu, Payel Mukherjee, Priya Singh, Dhiviya Vedagiri, Divya Gupta, Vishal Sah, Santosh Kumar Kuncha, Krishnan Harinivas Harshan, Archana Bharadwaj Siva, Karthik Bharadwaj Tallapaka, Shagufta Khan, Lamuk Zaveri, Namami Gaur, Sakshi Shambhavi, Tulasi Nagabandi, Purushotham Vodnala, Gokulan C G, Gunjan Purohit, Hanuman Tulashiram Kale, Pankaj Kumar, Prachand Issarapu, Rakesh K Mis |
| hCoV-19/India/CCMB_L1819/2020 | **EPI_ISL_471638** | 6/2/2020 | CSIR-Centre for Cellular and Molecular Biology | CSIR-Centre for Cellular and Molecular Biology | Sofia Banu, Payel Mukherjee, Priya Singh, Dhiviya Vedagiri, Divya Gupta, Vishal Sah, Santosh Kumar Kuncha, Krishnan Harinivas Harshan, Archana Bharadwaj Siva, Karthik Bharadwaj Tallapaka, Shagufta Khan, Lamuk Zaveri, Namami Gaur, Sakshi Shambhavi, Tulasi Nagabandi, Purushotham Vodnala,Preethi Jampala, Sharada Ravi Iyer, Sulagana Mukherjee, Swetha Sundar, Peddapuvala Sai Uday Kiran, Ra |
| hCoV-19/India/CCMB_L1822/2020 | **EPI_ISL_471639** | 6/2/2020 | CSIR-Centre for Cellular and Molecular Biology | CSIR-Centre for Cellular and Molecular Biology | Tulasi Nagabandi, Namami Gaur, Sakshi Shambhavi, Lamuk Zaveri, Shagufta Khan, Purushotham Vodnala, Payel Mukherjee, Sofia Banu, Priya Singh, Dhiviya Vedagiri, Divya Gupta, Vishal Sah, Santosh Kumar Kuncha, Krishnan Harinivas Harshan, Archana Bharadwaj Siva, Karthik Bharadwaj Tallapaka,G. Aditya Kumar, Koushick Sivakumar, Pooja Ramesh Gupta, Rajan Kumar Jha, Shraddha Vijay Lahoti, Rak |
| hCoV-19/India/CCMB_L1824/2020 | **EPI_ISL_471640** | 6/2/2020 | CSIR-Centre for Cellular and Molecular Biology | CSIR-Centre for Cellular and Molecular Biology | Tulasi Nagabandi, Namami Gaur, Sakshi Shambhavi, Lamuk Zaveri, Shagufta Khan, Purushotham Vodnala, Payel Mukherjee, Sofia Banu, Priya Singh, Dhiviya Vedagiri, Divya Gupta, Vishal Sah, Santosh Kumar Kuncha, Krishnan Harinivas Harshan, Archana Bharadwaj Siva, Karthik Bharadwaj Tallapaka,Kezia J Ann, Radhika Khandelwal, Roshan Maku Venkata, Shemin Mansuri, Sonu Uday, Rakesh K Mishra, [illegible] |
| hCoV-19/India/CCMB_L276_P3/2020 | **EPI_ISL_471641** | 4/20/2020 | CSIR-Centre for Cellular and Molecular Biology | CSIR-Centre for Cellular and Molecular Biology | Dhiviya Vedagiri, Divya Gupta, Vishal Sah, Payel Mukherjee, Sofia Banu, Priya Singh, Santosh Kumar Kuncha, Archana Bharadwaj Siva, Karthik Bharadwaj Tallapaka, Shagufta Khan, Lamuk Zaveri, Namami Gaur, Sakshi Shambhavi, Tulasi Nagabandi, Purushotham Vodnala, Rakesh K Mishra, Divya Tej Sowpati, Krishnan Harinivas Harshan |

| | | | | | |
|---|---|---|---|---|---|
| hCoV-19/India/CCMB_L301_P3/2020 | **EPI_ISL_471642** | 4/20/2020 | CSIR-Centre for Cellular and Molecular Biology | CSIR-Centre for Cellular and Molecular Biology | Dhiviya Vedagiri, Divya Gupta, Vishal Sah, Payel Mukherjee, Sofia Banu, Priya Singh, Santosh Kumar Kuncha, Archana Bharadwaj Siva, Karthik Bharadwaj Tallapaka, Shagufta Khan, Lamuk Zaveri, Namami Gaur, Sakshi Shambhavi, Tulasi Nagabandi, Purushotham Vodnala, Rakesh K Mishra, Divya Tej Sowpati, Krishnan Harinivas Harshan |
| hCoV-19/India/CCMB_L302/2020 | **EPI_ISL_471643** | 4/20/2020 | CSIR-Centre for Cellular and Molecular Biology | CSIR-Centre for Cellular and Molecular Biology | Tulasi Nagabandi, Namami Gaur, Sakshi Shambhavi, Lamuk Zaveri, Shagufta Khan, Purushotham Vodnala, Payel Mukherjee, Sofia Banu, Priya Singh, Dhiviya Vedagiri, Divya Gupta, Vishal Sah, Santosh Kumar Kuncha, Krishnan Harinivas Harshan, Archana Bharadwaj Siva, Karthik Bharadwaj Tallapaka,G. Aditya Kumar, Koushick Sivakumar, Pooja Ramesh Gupta, Rajan Kumar Jha, Shraddha Vijay Lahoti, Rak |
| hCoV-19/India/CCMB_L912/2020 | **EPI_ISL_471644** | 5/9/2020 | CSIR-Centre for Cellular and Molecular Biology | CSIR-Centre for Cellular and Molecular Biology | Tulasi Nagabandi, Namami Gaur, Sakshi Shambhavi, Lamuk Zaveri, Shagufta Khan, Purushotham Vodnala, Payel Mukherjee, Sofia Banu, Priya Singh, Dhiviya Vedagiri, Divya Gupta, Vishal Sah, Santosh Kumar Kuncha, Krishnan Harinivas Harshan, Archana Bharadwaj Siva, Karthik Bharadwaj Tallapaka,Kezia J Ann, Radhika Khandelwal, Roshan Maku Venkata, Shemin Mansuri, Sonu Uday, Rakesh K Mishra, D |
| hCoV-19/India/CCMB_O2_P4M/2020 | **EPI_ISL_471645** | 4/15/2020 | CSIR-Centre for Cellular and Molecular Biology | CSIR-Centre for Cellular and Molecular Biology | Dhiviya Vedagiri, Divya Gupta, Vishal Sah, Payel Mukherjee, Sofia Banu, Priya Singh, Santosh Kumar Kuncha, Archana Bharadwaj Siva, Karthik Bharadwaj Tallapaka, Shagufta Khan, Lamuk Zaveri, Namami Gaur, Sakshi Shambhavi, Tulasi Nagabandi, Purushotham Vodnala, Rakesh K Mishra, Divya Tej Sowpati, Krishnan Harinivas Harshan |
| hCoV-19/India/CCMB_O2_P4S/2020 | **EPI_ISL_471646** | 4/15/2020 | CSIR-Centre for Cellular and Molecular Biology | CSIR-Centre for Cellular and Molecular Biology | Dhiviya Vedagiri, Divya Gupta, Vishal Sah, Payel Mukherjee, Sofia Banu, Priya Singh, Santosh Kumar Kuncha, Archana Bharadwaj Siva, Karthik Bharadwaj Tallapaka, Shagufta Khan, Lamuk Zaveri, Namami Gaur, Sakshi Shambhavi, Tulasi Nagabandi, Purushotham Vodnala, Rakesh K Mishra, Divya Tej Sowpati, Krishnan Harinivas Harshan |

**Supplementary Data 1. Summary of the genomes of Indian isolates of SARS-CoV-2 available in public domain**
