## Supplementary Data 2 for "Analysis of the potential impact of genomic variants in SARS-CoV-2 genomes from India on molecular diagnostic assays"

| S.No. | Primers Sequence | Start | End | Probe Sequence | Start | End | Source |
|---|---|---|---|---|---|---|---|
| 1 | GTGARATGGTCATGTGTGGCGG | 15431 | 15452 | CAGGTGGAACCTCATCAGGAGATGC | 15470 | 15494 | https://www.who.int/docs/default-source/coronaviruse/protocol-v2-1.pdf?sfvrsn=a9ef618c_2 |
| | CARATGTTAAASACACTATTAGCAT | 15505 | 15530 | CCAGGTGGWACRTCATCMGGTGATGC | 15470 | 15494 | https://www.who.int/docs/default-source/coronaviruse/protocol-v2-1.pdf?sfvrsn=a9ef618c_22 |
| 2 | ACAGGTACGTTAATAGTTAATAGCG | 26269 | 26294 | ACACTAGCCATCCTTACTGCGCTTCG | 26332 | 26357 | https://www.who.int/docs/default-source/coronaviruse/protocol-v2-1.pdf?sfvrsn=a9ef618c_2 |
| | ATATTGCAGCAGTACGCACACA | 26360 | 26381 | | | | https://www.who.int/docs/default-source/coronaviruse/protocol-v2-1.pdf?sfvrsn=a9ef618c_2 |
| 3 | GGGGAACTTCTCCTGCTAGAAT | 28881 | 28902 | TTGCTGCTGCTTGACAGATT | 28934 | 28953 | https://www.who.int/docs/default-source/coronaviruse/whoinhouseassays.pdf?sfvrsn=de3a76aa_2 |
| | CAGACATTTTGCTCTCAAGCTG | 28958 | 28979 | | | | https://www.who.int/docs/default-source/coronaviruse/whoinhouseassays.pdf?sfvrsn=de3a76aa_2 |
| 4 | CCCTGTGGGTTTTACACTTAA | 13342 | 13362 | CCGTCTGCGGTATGTGGAAAGGTTATGG | 13377 | 13404 | https://www.who.int/docs/default-source/coronaviruse/whoinhouseassays.pdf?sfvrsn=de3a76aa_22 |
| | ACGATTGTGCATCAGCTGA | 13442 | 13460 | | | | https://www.who.int/docs/default-source/coronaviruse/whoinhouseassays.pdf?sfvrsn=de3a76aa_2 |
| 5 | GACCCCAAAATCAGCGAAAT | 28287 | 28306 | ACCCCGCATTACGTTTGGTGGACC | 28309 | 28332 | https://wwwnc.cdc.gov/eid/article/26/8/20-1246-t1 |
| | TCTGGTTACTGCCAGTTGAATCTG | 28335 | 28358 | | | | https://wwwnc.cdc.gov/eid/article/26/8/20-1246-t1 |
| 6 | TTACAAACATTGGCCGCAAA | 29164 | 29183 | ACAATTTGCCCCCAGCGCTTCAG | 29188 | 29210 | https://wwwnc.cdc.gov/eid/article/26/8/20-1246-t1 |
| | GCGCGACATTCCGAAGAA | 29213 | 29230 | | | | https://wwwnc.cdc.gov/eid/article/26/8/20-1246-t1 |
| 7 | GGGAGCCTTGAATACACCAAAA | 28681 | 28702 | AYCACATTGGCACCCGCAATCCTG | 28706 | 28727 | https://wwwnc.cdc.gov/eid/article/26/8/20-1246-t1 |
| | TGTAGCACGATTGCAGCATTG | 28732 | 28752 | | | | https://wwwnc.cdc.gov/eid/article/26/8/20-1246-t1 |
| 8 | TAATCAGACAAGGAACTGATTA | 29145 | 29166 | GCAAATTGTGCAATTTGCGG | 29177 | 29196 | https://www.who.int/docs/default-source/coronaviruse/peiris-protocol-16-1-20.pdf?sfvrsn=af1aac73_4 |
| | CGAAGGTGTGACTTCCATG | 29236 | 29254 | | | | https://www.who.int/docs/default-source/coronaviruse/peiris-protocol-16-1-20.pdf?sfvrsn=af1aac73_4 |
| 9 | AAATTTTGGGGACCAGGAAC | 29125 | 29144 | ATGTCGCGCATTGGCATGGA | 29222 | 29241 | https://www.who.int/docs/default-source/coronaviruse/method-niid-20200123-2.pdf?sfvrsn=fbf75320_2 |
| | TGGCAGCTGTGTAGGTCAAC | 29263 | 29282 | | | | https://www.who.int/docs/default-source/coronaviruse/method-niid-20200123-2.pdf?sfvrsn=fbf75320_2 |
| 10 | CACATTGGCACCCGCAATC | 28706 | 28724 | ACTTCCTCAAGGAACAACATTGCCA | 28753 | 28777 | https://www.ncbi.nlm.nih.gov/pmc/articles/PMC6988269/ |
| | GAGGAACGAGAAGAGGCTTG | 28814 | 28833 | | | | https://www.ncbi.nlm.nih.gov/pmc/articles/PMC6988269/ |
| 11 | CGTTTGGTGGACCCTCAGAT | 28320 | 28339 | CAACTGGCAGTAACCA | 28341 | 28356 | https://www.who.int/docs/default-source/coronaviruse/conventional-rt-pcr-followed-by-sequencing-for-detection-of-ncov-rirl-nat-inst-health-t.pdf?sfvrsn=42271c6d_4 |
| | CCCCACTGCGTTCTCCATT | 28358 | 28376 | | | | https://www.who.int/docs/default-source/coronaviruse/conventional-rt-pcr-followed-by-sequencing-for-detection-of-ncov-rirl-nat-inst-health-t.pdf?sfvrsn=42271c6d_4 |
| 12 | TGGGGYTTTACRGGTAACCT | 18778 | 18797 | TAGTTGTGATGCWATCATGACTAG | 18849 | 18872 | https://www.who.int/docs/default-source/coronaviruse/peiris-protocol-16-1-20.pdf?sfvrsn=af1aac73_4 |
| | AACRCGCTTAACAAAGCACTC | 18889 | 18909 | | | | https://www.who.int/docs/default-source/coronaviruse/peiris-protocol-16-1-20.pdf?sfvrsn=af1aac73_4 |
| 13 | CTCAGTCCAAGATGGTATTTCT | 28583 | 28604 | | | | https://academic.oup.com/cid/article/doi/10.1093/cid/ciaa201/5766416 |
| | AGCACCATAGGGAAGTCC | 28631 | 28648 | | | | https://academic.oup.com/cid/article/doi/10.1093/cid/ciaa201/5766416 |
| 14 | TCATTGTTAATGCCTATATTAACC | 14155 | 14178 | AACTGCAGAGTCACATGTTGACA | 14193 | 14215 | https://academic.oup.com/cid/article/doi/10.1093/cid/ciaa201/5766416 |
| | CACTTAATGTAAGGCTTTGTTAAG | 14220 | 14243 | | | | https://academic.oup.com/cid/article/doi/10.1093/cid/ciaa201/5766416 |
| 15 | CAAGTGGGGTAAGGCTAGACTTT | 14961 | 14983 | | | | https://www.thelancet.com/journals/lancet/article/PIIS0140-6736(20)30154-9/fulltext |
| | ACTTAGGATAATCCCAACCCAT | 15283 | 15304 | | | | https://www.thelancet.com/journals/lancet/article/PIIS0140-6736(20)30154-9/fulltext |
| 16 | CCTACTAAATTAAATGATCTCTGCTT | 22712 | 22741 | | | | https://www.thelancet.com/journals/lancet/article/PIIS0140-6736(20)30154-9/fulltext |
| | CAAGCTATAACGCAGCCTGTA | 22849 | 22869 | | | | https://www.thelancet.com/journals/lancet/article/PIIS0140-6736(20)30154-9/fulltext |
| 17 | CATGTGTGGCGGTTCACTAT | 15441 | 15460 | | | | http://www.en-journal.org/journal/view.html?uid=513&vmd=Full#T1 |
| | TGCATTAACATTGGCCGTGA | 15539 | 15558 | | | | http://www.en-journal.org/journal/view.html?uid=513&vmd=Full#T1 |
| 18 | CTACATGCACCAGCAACTGT | 23114 | 23133 | | | | http://www.en-journal.org/journal/view.html?uid=513&vmd=Full#T1 |
| | CACCTGTGCCTGTTAAACCA | 23194 | 23213 | | | | http://www.en-journal.org/journal/view.html?uid=513&vmd=Full#T1 |
| 19 | CAATGCTGCAATCGTGCTAC | 28732 | 28751 | | | | http://www.en-journal.org/journal/view.html?uid=513&vmd=Full#T1 |
| | GTTGCGACTACGTGATGAGG | 28830 | 28849 | | | | http://www.en-journal.org/journal/view.html?uid=513&vmd=Full#T1 |
| 20 | AGAATAGAGCTCGCACCGTA | 15092 | 15111 | | | | http://www.en-journal.org/journal/view.html?uid=513&vmd=Full#T1 |
| | CTCCTCTAGTGGCGGCTATT | 15174 | 15193 | | | | http://www.en-journal.org/journal/view.html?uid=513&vmd=Full#T1 |
| 21 | TCTGTGATGCCATGCGAAAT | 14015 | 14034 | | | | http://www.en-journal.org/journal/view.html?uid=513&vmd=Full#T1 |
| | ACTACCTGGCGTGGTTTGTA | 14108 | 14127 | | | | http://www.en-journal.org/journal/view.html?uid=513&vmd=Full#T1 |
| 22 | GCTGGTGCTGCAGCTTATTA | 22340 | 22359 | | | | http://www.en-journal.org/journal/view.html?uid=513&vmd=Full#T1 |
| | AGGGTCAAGTGCACAGTCTA | 22428 | 22447 | | | | http://www.en-journal.org/journal/view.html?uid=513&vmd=Full#T1 |
| 23 | TTCGGAAGAGACAGGTACGTTA | 26259 | 26280 | | | | http://www.en-journal.org/journal/view.html?uid=513&vmd=Full#T1 |
| | AGCAGTACGCACACAATCG | 26356 | 26374 | | | | http://www.en-journal.org/journal/view.html?uid=513&vmd=Full#T1 |
| 24 | GCTGCAATCGTGCTACAACT | 28736 | 28755 | | | | http://www.en-journal.org/journal/view.html?uid=513&vmd=Full#T1 |
| | TGAACTGTTGCGACTACGTG | 28836 | 28855 | | | | http://www.en-journal.org/journal/view.html?uid=513&vmd=Full#T1 |
| 26 | GCGTTCTTCGGAATGTCG | 29210 | 29227 | AACGTGGTTGACCTACACAGST | 29257 | 29276 | https://jcm.asm.org/content/58/5/e00310-20 |
| | TTGGATCTTTGTCATCCAATTTG | 29284 | 29306 | | | | https://jcm.asm.org/content/58/5/e00310-20 |
| 27 | AAACTTGTGCCCTTTTGGTG | 22561 | 22580 | CGCCACCAGATTTGCATCTG | 22591 | 22610 | https://www.sciencedirect.com/science/article/pii/S1477893920301009?via%3Dihub#bib8 |
| | TGCTGATTCTCTTCCTGTTCC | 22620 | 22640 | | | | https://www.sciencedirect.com/science/article/pii/S1477893920301009?via%3Dihub#bib88 |
| 28 | ATGGGTTGGGATTATCCTAAATGTGA | 15283 | 15308 | ATGCTTAGAATTATGGCCTCAC | 15325 | 15346 | https://www.biorxiv.org/content/10.1101/2020.04.14.041319v1.full |
| | GCAGTTGTGGCATCTCCTGATGAG | 15480 | 15503 | | | | https://www.biorxiv.org/content/10.1101/2020.04.14.041319v1.full |
| 29 | GGAAGAGACAGGTACGTTAATA | 26262 | 26283 | ACACTAGCCATCCTTACTGCGCTTCG | 26332 | 26357 | https://www.medrxiv.org/content/10.1101/2020.04.17.20067348v2.full.pdf+html |
| | AGCAGTACGCACACAATCGAA | 26354 | 26374 | | | | https://www.medrxiv.org/content/10.1101/2020.04.17.20067348v2.full.pdf+htmll |
| 30 | TCTGGTAAAGGCCAACAACAA | 28976 | 28996 | CTGTCACTAAGAAATCTGCTGCTGAGGC | 29007 | 29034 | https://www.biorxiv.org/content/10.1101/2020.04.06.028316v2.full.pdf+html |
| | TGTATGCTTTAGTGGCAGTACG | 29057 | 29078 | | | | https://www.biorxiv.org/content/10.1101/2020.04.06.028316v2.full.pdf+html |
| 31 | TGTGACATCAAGGACCTGCC | 26997 | 27016 | TGTTGCTACATCACGAACGC | 27029 | 27048 | https://www.biorxiv.org/content/10.1101/2020.04.20.052258v1.full.pdf |
| | CTGAGTCACCTGCTACACGC | 27077 | 27096 | | | | https://www.biorxiv.org/content/10.1101/2020.04.20.052258v1.full.pdf |
| 32 | TTGGCAAAATTCAAGACTCACTTT | 24354 | 24377 | | | | https://www.medrxiv.org/content/10.1101/2020.04.25.20079830v1.full.pdf |
| | TGTGGTTCATAAAAATTCCTTTGTG | 24876 | 24900 | | | | https://www.medrxiv.org/content/10.1101/2020.04.25.20079830v1.full.pdf |
| 33 | ATGAGCTTAGTCCTGTTG | 12690 | 12707 | AGATGTCTTGTGCTGCCGGTA | 12717 | 12737 | https://www.medrxiv.org/content/10.1101/2020.05.03.20072207v1 |
| | CTCCCTTTGTTGTGTTGT | 12780 | 12797 | | | | https://www.medrxiv.org/content/10.1101/2020.05.03.20072207v1 |
| 34 | GGTAACTGGTATGATTTCG | 14080 | 14098 | TCATACAAACCACGCCAGG | 14105 | 14123 | https://www.medrxiv.org/content/10.1101/2020.05.03.20072207v1 |
| | CTGGTCAAGGTTAATATAGG | 14167 | 14186 | | | | https://www.medrxiv.org/content/10.1101/2020.05.03.20072207v1 |

| | | | | | | | |
|---|---|---|---|---|---|---|---|
| 35 | CAAATTCTATGGTGGTTGGCACA | 15216 | 15238 | ATAATCCCAACCCATRAG | 15283 | 15300 | https://jcm.asm.org/content/jcm/early/2020/04/06/JCM.00557-20.full.pdf |
| | GGCATGGCTCTATCACATTTAGG | 15298 | 15320 | | | | https://jcm.asm.org/content/jcm/early/2020/04/06/JCM.00557-20.full.pdf |
| 36 | TCGGAAGAGACAGGTACGTT | 26260 | 26279 | CATCCTTACTGCGCTTCGAT | 26340 | 26359 | https://jabonline.in/admin/php/uploads/465_pdf.pdf |
| | CAATATTGCAGCAGTACGCA | 26364 | 26383 | | | | https://jabonline.in/admin/php/uploads/465_pdf.pdf |
| 37 | CTGACCAGACCGCTTCTAG | 26907 | 26925 | GACCTGCCTAAAGAAATCACT | 27009 | 27029 | https://jabonline.in/admin/php/uploads/465_pdf.pdf |
| | GAAAGCGTTCGTGATGTAGC | 27033 | 27052 | | | | https://jabonline.in/admin/php/uploads/465_pdf.pdf |
| 38 | CAAGCTTTCGGCAGACGTG | 29087 | 29105 | GCTTCAGCGTTCTTCGGAA | 29204 | 29222 | https://jabonline.in/admin/php/uploads/465_pdf.pdf |
| | TCCATGCCAATGCGCGAC | 29224 | 29241 | | | | https://jabonline.in/admin/php/uploads/465_pdf.pdf |
| 39 | TAATGGACCCCAAAATCAGC | 28282 | 28301 | | | | https://jabonline.in/admin/php/uploads/465_pdf.pdf |
| | TGCGTTCTCCATTCTGGTTA | 28351 | 28370 | | | | https://jabonline.in/admin/php/uploads/465_pdf.pdf |
| 40 | GAACTGATTACAAACATTGG | 29157 | 29176 | GCTTCAGCGTTCTTCGGAA | 29204 | 29222 | https://jabonline.in/admin/php/uploads/465_pdf.pdf |
| | CTTCCATGCCAATGCGCGACA | 29223 | 29243 | | | | https://jabonline.in/admin/php/uploads/465_pdf.pdf |
| 41 | TGCAACTGAGGGAGCCTTGA | 28672 | 28691 | | | | https://jabonline.in/admin/php/uploads/465_pdf.pdf |
| | CTTGAGGAAGTTGTAGCACG | 28744 | 28763 | | | | https://jabonline.in/admin/php/uploads/465_pdf.pdf |
| 42 | ACAGTTTATGATCCTTTGCAAC | 24968 | 24989 | | | | https://jabonline.in/admin/php/uploads/465_pdf.pdf |
| 43 | TGATGGTGGTGTCACTCGTG | 8701 | 8720 | TGGTTTAGCCAGCGTGGTGGT | 8774 | 8794 | https://virological.org/t/preliminary-in-silico-assessment-of-the-specificity-of-published-molecular-assays-and-design-of-new-assays-using-the-available-whole-genome-sequences-of-2019-ncov/343 |
| | GAAGTGGGTTTTGTCGTGCC | 8846 | 8865 | | | | https://virological.org/t/preliminary-in-silico-assessment-of-the-specificity-of-published-molecular-assays-and-design-of-new-assays-using-the-available-whole-genome-sequences-of-2019-ncov/343 |
| 44 | GCCGCTGTTGATGCACTATG | 17170 | 17189 | ACGTGCTCGTGTAGAGTGTTTTGAT | 17244 | 17268 | https://virological.org/t/preliminary-in-silico-assessment-of-the-specificity-of-published-molecular-assays-and-design-of-new-assays-using-the-available-whole-genome-sequences-of-2019-ncov/343 |
| | ATGCATTGCCTGAGACGACA | 17318 | 17337 | | | | https://virological.org/t/preliminary-in-silico-assessment-of-the-specificity-of-published-molecular-assays-and-design-of-new-assays-using-the-available-whole-genome-sequences-of-2019-ncov/343 |
| 45 | CGGATGGCTTATTGTTGGCG | 25521 | 25540 | TGCTCGTTGCTGCTGGCCTT | 25676 | 25695 | https://virological.org/t/preliminary-in-silico-assessment-of-the-specificity-of-published-molecular-assays-and-design-of-new-assays-using-the-available-whole-genome-sequences-of-2019-ncov/343 |
| | TTGGCTTTGCTGGAAATGCC | 25773 | 25792 | | | | https://virological.org/t/preliminary-in-silico-assessment-of-the-specificity-of-published-molecular-assays-and-design-of-new-assays-using-the-available-whole-genome-sequences-of-2019-ncov/343 |
| 46 | TGTCGTTGACAGGACACGAG | 148 | 167 | TTCGTCCGTGTTGCAGCCGA | 201 | 220 | https://virological.org/t/preliminary-in-silico-assessment-of-the-specificity-of-published-molecular-assays-and-design-of-new-assays-using-the-available-whole-genome-sequences-of-2019-ncov/343 |
| | CGTACGTGGCTTTGGAGACT | 346 | 365 | | | | https://virological.org/t/preliminary-in-silico-assessment-of-the-specificity-of-published-molecular-assays-and-design-of-new-assays-using-the-available-whole-genome-sequences-of-2019-ncov/343 |
| 47 | CACATTGGCACCCGCAATC | 28706 | 28724 | ACTTCCTCAAGGAACAACATTGCCA | 28753 | 28777 | https://virological.org/t/preliminary-in-silico-assessment-of-the-specificity-of-published-molecular-assays-and-design-of-new-assays-using-the-available-whole-genome-sequences-of-2019-ncov/343 |
| | CAAGCCTCTTCTCGTTCCTC | 28814 | 28833 | | | | https://virological.org/t/preliminary-in-silico-assessment-of-the-specificity-of-published-molecular-assays-and-design-of-new-assays-using-the-available-whole-genome-sequences-of-2019-ncov/343 |
| 48 | GTGARATGGTCATGTGTGGCGG | 15431 | 15452 | CAGGTGGAACCTCATCAGGAGATGC | 15470 | 15494 | https://virological.org/t/preliminary-in-silico-assessment-of-the-specificity-of-published-molecular-assays-and-design-of-new-assays-using-the-available-whole-genome-sequences-of-2019-ncov/343 |
| | TATGCTAATAGTGTSTTTAACATYTG | 15505 | 15531 | | | | https://virological.org/t/preliminary-in-silico-assessment-of-the-specificity-of-published-molecular-assays-and-design-of-new-assays-using-the-available-whole-genome-sequences-of-2019-ncov/343 |
| 49 | GTCATGTGTGGCGGTTCACT | 15439 | 15458 | CAGGTGGAACCTCATCAGGAGATGC | 15470 | 15494 | https://www.medrxiv.org/content/10.1101/2020.04.17.20067348v2.full.pdf+html |
| 50 | AAATTCTATGGTGGTTGGCACAACAT | 15217 | 15245 | TGGGTTGGGATTATC | 15284 | 15298 | http://covid-19.dnageography.com/ |
| 51 | CGCATACAGTCTTRCAGGCT | 16220 | 16239 | TTAAGATGTGGTGCTTGCATACGTAGAC | 16276 | 16303 | https://jcm.asm.org/content/early/2020/03/27/JCM.00310-20 |

**Supplementary Data 2. Curated primers and probes sequence and their genomic coordinate used in the molecular assays for detection of SARS-CoV-2.**
