## Supplementary Data 3 for "Analysis of the potential impact of genomic variants in SARS-CoV-2 genomes from India on molecular diagnostic assays"

| Primer/Probe Sequence / Description | Genomic Variant | Melting Temperature (Ref / Alt) | Gene | No of genomes with variant | Allele frequency | Cumulative No of variants and Freq of variants |
|---|---|---|---|---|---|---|
| GGGGAACTTCTCCTGCTAGAAT (28881-28902) | 28881:G:A | 54.8 / 53 | N | 93 | 0.13 | 283 (0.395) |
| | 28881:G:R | 54.8 / 53 to 54.8 | | 1 | 0.001 | |
| | 28882:G:A | 54.8 / 53 | | 93 | 0.13 | |
| | 28882:G:R | 54.8 / 53 to 54.8 | | 1 | 0.001 | |
| | 28883:G:C | 54.8 / 54.8 | | 93 | 0.13 | |
| | 28883:G:R | 54.8 / 53 to 54.8 | | 1 | 0.13 | |
| | 28890:C:G | 54.8 / 54.8 | | 1 | 0.001 | |
| ACCCCGCATTACGTTTGGTGGACC (28309-28332) | 28311:C:T | 58.8 / 57.1 | N | 191 | 0.266 | 214 (0.298) |
| | 28311:C:Y | 58.8 / 57.1 to 58.8 | | 9 | 0.013 | |
| | 28312:C:T | 58.8 / 57.1 | | 5 | 0.007 | |
| | 28326:G:T | 58.8 / 57.1 | | 9 | 0.013 | |
| TGAACTGTTGCGACTACGTG (28836-28855) | 28845:G:T | 51.8 / 49.7 | N | 2 | 0.003 | 96 (0.134) |
| | 28851:G:T | 51.8 / 49.7 | | 1 | 0.001 | |
| | 28854:C:T | 51.8 / 49.7 | | 93 | 0.13 | |
| AGGGTCAAGTGCACAGTCTA (22428-22447) | 22444:C:T | 51.8 / 49.7 | S | 88 | 0.123 | 88 (0.123) |
| TGGTTTAGCCAGCGTGGTGGT (8774-8794) | 8782:C:T | 56.3 / 54.4 | ORF1a | 35 | 0.049 | 35 (0.049) |
| TGCAACTGAGGGAGCCTTGA (28672-28691) | 28674:C:T | 53.8 / 51.8 | N | 1 | 0.001 | 28 (0.039) |
| | 28676:A:M | 53.8 / 53.8 to 55.9 | | 2 | 0.003 | |
| | 28688:T:C | 53.8 / 55.9 | | 20 | 0.028 | |
| | 28688:T:Y | 53.8 / 53.8 to 55.9 | | 2 | 0.003 | |
| | 28690:G:T | 53.8 / 51.8 | | 3 | 0.004 | |
| GGGAGCCTTGAATACACCAAAA (28681-28702) | 28688:T:C | 53 / 54.8 | N | 20 | 0.028 | 26 (0.036) |
| | 28688:T:Y | 53 / 53 to 54.8 | | 2 | 0.003 | |
| | 28690:G:T | 53 / 51.1 | | 3 | 0.004 | |
| | 28699:A:M | 53 / 53 to 54.8 | | 1 | 0.001 | |
| GTGARATGGTCATGTGTGGCGG (15431-15452) | 15435:A:R | 56.7 to 58.6 / 56.7 to 60.4 | RdRP | 19 | 0.026 | 24 (0.033) |
| | 15444:G:T | 56.3 / 54.4 | | 1 | 0.001 | |
| | 15451:G:A | 56.3 / 54.4 | | 4 | 0.006 | |
| CTACATGCACCAGCAACTGT (23114-23133) | 23116:A:T | 51.8 / 51.8 | S | 1 | 0.001 | 15 (0.021) |
| | 23116:A:W | 51.8 / 51.8 | | 4 | 0.006 | |
| | 23118:A:M | 51.8 / 51.8 to 53.8 | | 3 | 0.004 | |
| | 23120:G:K | 51.8 / 49.7 to 51.8 | | 1 | 0.001 | |
| | 23120:G:T | 51.8 / 49.7 | | 5 | 0.007 | |
| | 23123:C:T | 51.8 / 49.7 | | 1 | 0.001 | |
| CGGATGGCTTATTGTTGGCG (25521-25540) | 25528:C:T | 53.8 / 51.8 | ORF3a | 13 | 0.018 | 14 (0.020) |
| | 25528:C:Y | 53.8 / 51.8 to 53.8 | | 1 | 0.001 | |
| ACACTAGCCATCCTTACTGCGCTTCG (26332-26357) | 26332:A:T | 61.1 / 61.1 | E | 1 | 0.001 | 10 (0.014) |
| | 26336:T:C | 61.1 / 62.7 | | 1 | 0.001 | |
| | 26338:G:T | 61.1 / 59.5 | | 2 | 0.003 | |
| | 26351:CGCTTC:- | 61.1 / 51.8 | | 5 | 0.007 | |

| | | | | | | |
|---|---|---|---|---|---|---|
| | 26356:C:A | 61.1 / 59.5 | | 1 | 0.001 | |
| CGTTTGGTGGACCCTCAGAT (28320-28339) | 28326:G:T | 53.8 / 51.8 | N | 9 | 0.013 | 10 (0.014) |
| | 28337:G:T | 53.8 / 51.8 | | 1 | 0.001 | |
| AGCAGTACGCACACAATCGAA (26354-26374) | 26351:CGCTTC:- | 52.4 / 48 | E | 5 | 0.007 | 10 (0.014) |
| | 26356:C:A | 52.4 / 50.5 | | 1 | 0.001 | |
| | 26356:C:T | 52.4 / 50.5 | | 1 | 0.001 | |
| | 26358: ATTGTGTGCGTACTGCTGCAATATT GTTAACGTGAGTCTTG:- | 52.4 / 12 | | 1 | 0.001 | |
| | 26361:G:A | 52.4 / 50.5 | | 1 | 0.001 | |
| | 26370:C:T | 52.4 / 50.5 | | 1 | 0.001 | |
| TTCGTCCGTGTTGCAGCCGA (201-220) | 203:C:T | 55.9 / 53.8 | ORF1a | 1 | 0.001 | 8 (0.011) |
| | 204:G:T | 55.9 / 53.8 | | 1 | 0.001 | |
| | 218:C:T | 55.9 / 53.8 | | 2 | 0.003 | |
| | 219:G:T | 55.9 / 53.8 | | 4 | 0.006 | |
| ATATTGCAGCAGTACGCACACA (26360-26381) | 26358: ATTGTGTGCGTACTGCTGCAATATT GTTAACGTGAGTCTTG:- | 53 / 4 | E | 1 | 0.001 | 8 (0.011) |
| | 26361:G:A | 53 / 51.1 | | 1 | 0.001 | |
| | 26370:C:T | 53 / 51.1 | | 1 | 0.001 | |
| | 26375:G:A | 53 / 51.1 | | 2 | 0.003 | |
| | 26376:C:A | 53 / 51.1 | | 2 | 0.003 | |
| | 26378:A:R | 53 / 53 to 54.8 | | 1 | 0.001 | |
| CATCCTTACTGCGCTTCGAT (26340-26359) | 26351:CGCTTC:- | 51.8 / 34.4 | E | 5 | 0.007 | 7 (0.010) |
| | 26356:C:A | 51.8 / 49.7 | | 1 | 0.001 | |
| | 26358: ATTGTGTGCGTACTGCTGCAATATT GTTAACGTGAGTCTTG:- | 51.8 / 48 | | 1 | 0.001 | |
| ACGATTGTGCATCAGCTGA (13442-13460) | 13449:A:G | 48.9 / 51.1 | ORF1ab | 1 | 0.001 | 7 (0.010) |
| | 13458:C:T | 48.9 / 46.8 | | 6 | 0.008 | |
| CTAGTTACACTAGCCATCCTTACTGC (26326-26351) | 26329:G:A | 58 / 56.4 | E | 1 | 0.001 | 7 (0.010) |
| | 26330:T:G | 58 / 59.5 | | 2 | 0.003 | |
| | 26332:A:T | 58 / 58 | | 1 | 0.001 | |
| | 26336:T:C | 58 / 59.5 | | 1 | 0.001 | |
| | 26338:G:T | 58 / 56.4 | | 2 | 0.003 | |
| CAATATTGCAGCAGTACGCA (26364-26383) | 26358: ATTGTGTGCGTACTGCTGCAATATT GTTAACGTGAGTCTTG:- | 49.7 / NA | E | 1 | 0.001 | 7 (0.010) |
| | 26370:C:T | 49.7 / 47.7 | | 1 | 0.001 | |
| | 26375:G:A | 49.7 / 47.7 | | 2 | 0.003 | |
| | 26376:C:A | 49.7 / 47.7 | | 2 | 0.003 | |
| | 26378:A:R | 49.7 / 53 to 54.8 | | 1 | 0.001 | |
| CAGACATTTTGCTCTCAAGCTG (28958-28979) | 28969:C:T | 53 / 51.1 | N | 3 | 0.004 | 6 (0.008) |
| | 28975:G:C | 53 / 53 | | 1 | 0.001 | |
| | 28975:G:K | 53 / 51.1 to 53 | | 1 | 0.001 | |

| | | | | | | |
|---|---|---|---|---|---|---|
| | 28979:G:T | 53 / 51.1 | | 1 | 0.001 | |
| TTGCTGCTGCTTGACAGATT (28934-28953) | 28948:C:T | 49.7 / 47.7 | N | 5 | 0.007 | 6 (0.008) |
| | 28952:T:Y | 49.7 / 49.7 to 51.8 | | 1 | 0.001 | |
| GTCATGTGTGGCGGTTCACT (15439-15458) | 15444:G:T | 53.8 / 51.8 | RdRp | 1 | 0.001 | 5 (0.007) |
| | 15451:G:A | 53.8 / 51.8 | | 4 | 0.006 | |
| CATGTGTGGCGGTTCACTAT (15441-15460) | 15444:G:T | 51.8 / 49.7 | RdRp | 1 | 0.001 | 5 (0.007) |
| | 15451:G:A | 51.8 / 49.7 | | 4 | 0.006 | |
| GCTGGTGCTGCAGCTTATTA (22340-22359) | 22343:G:A | 51.8 / 49.7 | S | 3 | 0.004 | 5 (0.007) |
| | 22350:C:M | 51.8 / 49.7 to 51.8 | | 1 | 0.001 | |
| | 22351:A:W | 51.8 / 51.8 | | 1 | 0.001 | |
| CCGTCTGCGGTATGTGGAAAGGTTATGG (13377-13404) | 13390:G:K | 62.9 / 61.4 to 62.9 | ORF1ab | 1 | 0.001 | 5 (0.007) |
| | 13394:A:R | 62.9 / 62.9 to 64.3 | | 1 | 0.001 | |
| | 13402:T:K | 62.9 / 62.9 to 64.3 | | 3 | 0.004 | |
| AACTGCAGAGTCACATGTTGACA (14193-14215) | 14198:C:G | 53.5 / 53.5 | ORF1ab | 1 | 0.001 | 5 (0.007) |
| | 14207:A:M | 53.5 / 53.5 to 55.3 | | 3 | 0.004 | |
| | 14212:G:T | 53.5 / 51.7 | | 1 | 0.001 | |
| CAAGCTATAACGCAGCCTGTA (22849-22869) | 22855:C:A | 52.4 / 50.5 | S | 1 | 0.001 | 4 (0.006) |
| | 22858:C:A | 52.4 / 50.5 | | 1 | 0.001 | |
| | 22861:T:A | 52.4 / 52.4 | | 1 | 0.001 | |
| | 22863:T:A | 52.4 / 52.4 | | 1 | 0.001 | |
| ACAATTTGCCCCCAGCGCTTCAG (29188-29210) | 29188:A:T | 60.8 / 60.8 | N | 2 | 0.003 | 4 (0.006) |
| | 29197:C:T | 60.8 / 59.1 | | 2 | 0.003 | |
| AGCAGTACGCACACAATCG (26356-26374) | 26356:C:A | 51.1 / 48.9 | E | 1 | 0.001 | 4 (0.006) |
| | 26356:C:T | 51.1 / 48.9 | | 1 | 0.001 | |
| | 26361:G:A | 51.1 / 48.9 | | 1 | 0.001 | |
| | 26370:C:T | 51.1 / 48.9 | | 1 | 0.001 | |
| TTAAGATGTGGTGCTTGCATACGTAGAC (16276-16303) | 16281:A:R | 58.5 / 58.8 to 59.9 | RdRp | 4 | 0.006 | 4 (0.006) |
| AYCACATTGGCACCCGCAATCCTG (28706-28727) | 28712:G:R | 59.1 to 60.8 / 57.4 to 60.8 | N | 2 | 0.003 | 4 (0.006) |
| | 28720:C:T | 59.1 to 60.8 / 57.1 | | 1 | 0.001 | |
| | 28727:G:T | 59.1 to 60.8 / 57.1 | | 1 | 0.001 | |
| CGTACGTGGCTTTGGAGACT (346-365) | 347:G:R | 53.8 / 51.8 to 53.8 | NA | 1 | 0.001 | 3 (0.004) |
| | 351:G:K | 53.8 / 51.8 to 53.8 | | 1 | 0.001 | |
| | 353:G:T | 53.8 / 51.8 | | 1 | 0.001 | |
| CAAGTGGGGTAAGGCTAGACTTT (14961-14983) | 14964:A:R | 55.3 / 55.3 to 57.1 | ORF1b | 1 | 0.001 | 3 (0.004) |
| | 14974:G:K | 55.3 / 53.5 to 55.3 | | 2 | 0.003 | |
| CGCATACAGTCTTRCAGGCT (16220-16239) | 16221:G:A | 51.1 / 48.9 | RdRp | 1 | 0.001 | 3 (0.004) |
| | 16232:T:W | 51.1 / 51.8 to 53.8 | | 2 | 0.003 | |
| CACATTGGCACCCGCAATC (28706-28724) | 28712:G:R | 53.2 / 51.1 to 53.2 | N | 2 | 0.003 | 3 (0.004) |
| | 28720:C:T | 53.2 / 51.1 | | 1 | 0.001 | |
| CGAAGGTGTGACTTCCATG (29236-29254) | 29236:C:T | 51.1 / 48.9 | N | 2 | 0.003 | 3 (0.004) |
| | 29254:G:T | 51.1 / 48.9 | | 1 | 0.001 | |
| TGCTCGTTGCTGCTGGCCTT (25676-25695) | 25691:G:T | 55.9 / 53.8 | ORF3a | 1 | 0.001 | 3 (0.004) |
| | 25693:C:T | 55.9 / 53.8 | | 2 | 0.003 | |

| | | | | | | |
|---|---|---|---|---|---|---|
| GCAAATTGTGCAATTTGCGG (29177-29196) | 29177:C:T | 49.7 / 47.7 | N | 1 | 0.001 | 3 (0.004) |
| | 29188:A:T | 49.7 / 49.7 | | 2 | 0.003 | |
| CTGGTCAAGGTTAATATAGG (14167-14186) | 14181:G:A | 47.7 / 45.6 | RdRp | 2 | 0.003 | 2 (0.003) |
| CTCCTCTAGTGGCGGCTATT (15174-15193) | 15180:C:A | 53.8 / 51.8 | RdRp | 1 | 0.001 | 2 (0.003) |
| | 15180:C:M | 53.8 / 51.8 to 53.8 | | 1 | 0.001 | |
| TTGGCTTTGCTGGAAATGCC (25773-25792) | 25785:G:T | 51.8 / 49.7 | ORF3a | 1 | 0.001 | 2 (0.003) |
| | 25788:A:G | 51.8 / 53.8 | | 1 | 0.001 | |
| TAATGGACCCCAAAATCAGC (28282-28301) | 28289:C:A | 49.7 / 47.7 | N | 2 | 0.003 | 2 (0.003) |
| GACCCCAAAATCAGCGAAAT (28287-28306) | 28289:C:A | 49.7 / 47.7 | N | 2 | 0.003 | 2 (0.003) |
| CCCCACTGCGTTCTCCATT (28358-28376) | 28362:G:T | 53.2 / 51.1 | N | 1 | 0.001 | 2 (0.003) |
| | 28371:G:T | 53.2 / 51.1 | | 1 | 0.001 | |
| CTCAGTCCAAGATGGTATTTCT (28583-28604) | 28603:C:T | 51.1 / 49.2 | N | 2 | 0.003 | 2 (0.003) |
| TGTAGCACGATTGCAGCATTG (28732-28752) | 28739:G:T | 52.4 / 50.5 | N | 1 | 0.001 | 2 (0.003) |
| | 28752:A:C | 52.4 / 54.4 | | 1 | 0.001 | |
| GCTGCAATCGTGCTACAACT (28736-28755) | 28739:G:T | 51.8 / 49.7 | N | 1 | 0.001 | 2 (0.003) |
| | 28752:A:C | 51.8 / 53.8 | | 1 | 0.001 | |
| CTTGAGGAAGTTGTAGCACG (28744-28763) | 28752:A:C | 51.8 / 53.8 | N | 1 | 0.001 | 2 (0.003) |
| | 28761:A:T | 51.8 / 51.8 | | 1 | 0.001 | |
| GTTGCGACTACGTGATGAGG (28830-28849) | 28845:G:T | 53.8 / 51.8 | N | 2 | 0.003 | 2 (0.003) |
| TCTGGTAAAGGCCAACAACAA (28976-28996) | 28979:G:T | 50.5 / 48.5 | N | 1 | 0.001 | 2 (0.003) |
| | 28985:G:K | 50.5 / 48.5 to 50.5 | | 1 | 0.001 | |
| TTACAAACATTGGCCGCAAA (29164-29183) | 29171:C:T | 47.7 / 45.6 | N | 1 | 0.001 | 2 (0.003) |
| | 29177:C:T | 47.7 / 45.6 | | 1 | 0.001 | |
| CTTCCATGCCAATGCGCGACA (29223-29243) | 29236:C:T | 56.3 / 54.4 | N | 2 | 0.003 | 2 (0.003) |
| TCCATGCCAATGCGCGAC (29224-29241) | 29236:C:T | 52.6 / 50.3 | N | 2 | 0.003 | 2 (0.003) |
| TTGGATCTTTGTCATCCAATTTG (29284-29306) | 29296:C:T | 49.9 / 48.1 | N | 1 | 0.001 | 2 (0.003) |
| | 29303:C:T | 49.9 / 48.1 | | 1 | 0.001 | |
| CAGGTGGAACCTCATCAGGAGATGC (15470-15494) | 15477:A:G | 61 / 62.6 | RdRP | 1 | 0.001 | 2 (0.003) |
| | 15491:A:G | 61 / 62.6 | | 1 | 0.001 | |
| CCAGGTGGWACRTCATCMGGTGATGC (15470-15494) | 15477:A:G | 60.6 / 62.4 | RdRP | 1 | 0.001 | 2 (0.003) |
| | 15491:A:G | 60.6 / 62.4 | | 1 | 0.001 | |
| CGCCACCAGATTTGCATCTG (22591-22610) | 22603:T:C | 53.8 / 55.9 | S | 1 | 0.001 | 2 (0.003) |
| | 22604:G:T | 53.8 / 51.8 | | 1 | 0.001 | |
| ATGTCGCGCATTGGCATGGA (29222-29241) | 29236:C:T | 53.8 / 51.8 | N | 2 | 0.003 | 2 (0.003) |
| TGATGGTGGTGTCACTCGTG (8701-8720) | 8713:C:T | 53.8 / 51.8 | ORF1a | 1 | 0.001 | 1 (0.001) |
| GAAGTGGGTTTTGTCGTGCC (8846-8865) | 8856:T:Y | 53.8 / 53.8 to 55.9 | NA | 1 | 0.001 | 1 (0.001) |
| CTCCCTTTGTTGTGTTGT (12780-12797) | 12793:G:T | 45.8 / 43.5 | RdRp | 1 | 0.001 | 1 (0.001) |

| | | | | | | |
|---|---|---|---|---|---|---|
| TCTGTGATGCCATGCGAAAT (14015-14034) | 14028:G:T | 49.7 / 47.7 | RdRp | 1 | 0.001 | 1 (0.001) |
| ACTACCTGGCGTGGTTTGTA (14108-14127) | 14125:A:G | 51.8 / 53.8 | RdRp | 1 | 0.001 | 1 (0.001) |
| GCAGTTGTGGCATCTCCTGATGAG (15480-15503) | 15491:A:G | 59.1 / 60.8 | S | 1 | 0.001 | 1 (0.001) |
| GCCGCTGTTGATGCACTATG (17170-17189) | 17185:C:T | 53.8 / 51.8 | NA | 1 | 0.001 | 1 (0.001) |
| AAACTTGTGCCCTTTTGGTG (22561-22580) | 22567:G:A | 49.7 / 47.7 | S | 1 | 0.001 | 1 (0.001) |
| CCTACTAAATTAAATGATCTCTGCTTTACT (22712-22741) | 22724:A:R | 54.8 / 54.8 to 56.2 | S | 1 | 0.001 | 1 (0.001) |
| TTGGCAAAATTCAAGACTCACTTT (24354-24377) | 24367:A:G | 50.6 / 52.3 | S | 1 | 0.001 | 1 (0.001) |
| TGTGGTTCATAAAAATTCCTTTGTG (24876-24900) | 24886:T:Y | 51.1 / 51.1 to 52.8 | S | 1 | 0.001 | 1 (0.001) |
| TGTGACATCAAGGACCTGCC (26997-27016) | 27014:G:T | 53.8 / 51.8 | M | 1 | 0.001 | 1 (0.001) |
| TCTGGTTACTGCCAGTTGAATCTG (28335-28358) | 28337:G:T | 55.7 / 54 | N | 1 | 0.001 | 1 (0.001) |
| TGCGTTCTCCATTCTGGTTA (28351-28370) | 28362:G:T | 49.7 / 47.7 | N | 1 | 0.001 | 1 (0.001) |
| AGCACCATAGGGAAGTCC (28631-28648) | 28631:G:A | 50.3 / 48 | N | 1 | 0.001 | 1 (0.001) |
| CAATGCTGCAATCGTGCTAC (28732-28751) | 28739:G:T | 51.8 / 49.7 | N | 1 | 0.001 | 1 (0.001) |
| GAACTGATTACAAACATTGG (29157-29176) | 29171:C:T | 45.6 / 43.6 | N | 1 | 0.001 | 1 (0.001) |
| TGGCAGCTGTGTAGGTCAAC (29263-29282) | 29272:C:T | 53.8 / 51.8 | N | 1 | 0.001 | 1 (0.001) |
| ATGCTTAGAATTATGGCCTCAC (15325-15346) | 15346:C:A | 51.1 / 49.2 | S | 1 | 0.001 | 1 (0.001) |
| GACCTGCCTAAAGAAATCACT (27009-27029) | 27014:G:T | 50.5 / 48.5 | M | 1 | 0.001 | 1 (0.001) |
| ACTTCCTCAAGGAACAACATTGCCA (28753-28777) | 28761:A:T | 56 / 56 | N | 1 | 0.001 | 1 (0.001) |
| AACGTGGTTGACCTACACAGST (29257-29276) | 29272:C:T | 54.8 / 50.5 | N | 1 | 0.001 | 1 (0.001) |

**Supplementary Data 3. Summary of Primer and Probe sequences and genomic variants**
Calculated variant frequency, cumulative variant frequency, and melting temperature (Tm) for reference and alternate in the Indian SARS-CoV-2 isolates in the primers and probes binding sites.
