## Supplementary Method 1 for "Analysis of the potential impact of genomic variants in SARS-CoV-2 genomes from India on molecular diagnostic assays"

**Supplementary Method 1: Methodology adopted for calculating primers/probes melting temperature and Gibbs free energy with variant impact.**

Tm has been calculated for primers with nucleotide greater than 13 using a formula Tm = 64.9 +41*(yG+zC-16.4)/(wA+xT+yG+zC) where w, x, y, and z are the number of A, T, G, and C nucleotides respectively (Wallace et al. 1979)**.**

*Estimation of thermodynamic parameters for oligonucleotide duplex: Evaluating the* thermodynamics of nucleotide binding could provide a unique opportunity to reveal the stability of the oligonucleotides and their interaction with the viral genome. The interactions could be summarised as follows. In RT-PCR, the key components depend on the hybridization of viral RNA to the reverse primer in the first step of cDNA synthesis that provide the first template for further reaction (RNA-DNA interaction). After cDNA synthesis it undergoes a PCR reaction that depends on the hybridization of the cDNA template to the primer / probe (DNA-DNA interaction). Similarly, the probes bind to the amplified DNA product (DNA-DNA interaction).

To evaluate the stability of viral RNA and reverse primer as well as viral cDNA and forward primer or probe duplex upon variation at the primer / probe binding sites. We analyzed the thermodynamic parameter Gibbs free energy (ΔG) at the standard conditions (1M NaCl, 37°C, and pH=7) for the reference and variant viral genome sequences with the cognate primer and probe sequences. We have considered only variants mapping to the primer/probe target sequences with a cumulative variant frequency ≥ 1% in the SARS-CoV-2 genomes of isolates from India. ΔG was calculated based on the Watson-Crick base pair nearest neighbor model for both the duplex secondary structure. The thermodynamic parameter for both RNA and DNA has been experimentally evaluated by different labs and derived using linear regression at the standard condition (1M NaCl, 37°C, and pH=7), tabulated in **Supplementary Data 4.** $\Delta G^0_{37}$ for DNA/DNA duplex and RNA/DNA heteroduplex calculated using Eq. 1 and Eq. 2 respectively (Sugimoto et al. 1995, 1996; SantaLucia 1998; SantaLucia and Hicks 2004).

$\Delta G^0_{37}$ (DNA duplex) = $\Delta G^0_{37}$(initiation) + $\Delta G^0_{37}$(symmetry) + $\Sigma\Delta G^0_{37}$(stack) +$\Delta G^0_{AT}$ (terminal)
(1)

where $\Delta G^0_{37}$ (initiation)=1.96 kcal/mol; $\Delta G^0_{AT}$ (terminal)=0.05 kcal/mol, $\Delta G^0_{37}$ (symmetry)= 0.43 kcal/mol penalty is applied on self complementary sequences and $\Sigma\Delta G^0_{37}$(stack) is the summation of the $\Delta G^0_{37}$ of the adjacent bases.

$\Delta G^0_{37}$ (RNA/DNA duplex) = $\Sigma\Delta G^0_{37}$(stack) + $\Delta G^0_{37}$(initiation)
(2)
where $\Delta G^0_{37}$(initiation) =3.1 kcal/mol and $\Sigma\Delta G^0_{37}$(stack) is the summation of the $\Delta G^0_{37}$ of the adjacent bases.

We also evaluated the internal single mismatch as well as terminal mismatch had an impact on the thermodynamics stability of the nucleic acid secondary structure. The thermodynamic parameter for internal single mismatch and terminal dangling end mismatch for nearest neighbour base pair has been evaluated by UV absorbance and NMR experiment of DNA/DNA duplex (Allawi and SantaLucia 1997, 1998a, 1998b, 1998c) as well as for rA.dA, rC.dC, rG.dG, and rU.dT mismatches for RNA/DNA heteroduplex (Bommarito et al. 2000; Watkins et al. 2011). The RNA/DNA free energy for primer at the single internal mismatch is calculated by Eq. 3. ΔG for RNA/DNA heteroduplex for single internal mismatches other than rA.dA, rC.dC, rG.dG, and rU.dT as well as terminal mismatch for DNA/DNA duplex are not readily available. So, we have used free energy parameters by single internal mismatch and terminal dangling end mismatch of DNA/DNA duplex to predict the free energy since they had comparative thermodynamic values and follow similar trend (Peyret et al. 1999; Bommarito et al. 2000; Watkins et al. 2011).

$$\Delta G^0{}_{37}\text{(RNA mismatch trimer)} = \Delta G^0{}_{37}\text{(exp)} - \Delta G^0{}_{37}\text{(core)} + \Delta G^0{}_{37}\text{(NN)} \quad (3)$$

where $\Delta G^0{}_{37}$(exp) is the free energy for the mutated RNA/DNA hybrid sequence $\Delta G^0{}_{37}$(core) is the free energy of core complex i.e. without variation, $\Delta G^0{}_{37}$ (NN) is the free energy for Watson-Crick dimer nearest-neighbour.
